# Chemical Interrogation and Reprogramming of ATAT1-Mediated Tubulin Acetylation

**DOI:** 10.64898/2026.08.17.745310

**Authors:** Luis E. Hernández Ramírez, Aleksandar Salim, Cornelia Egoldt, Laurane Michel, Charlotte Aumeier, Sascha Hoogendoorn

## Abstract

Acetylation of *α*-tubulin K40 by *α*-tubulin acetyltransferase 1 (ATAT1) using acetyl-coenzyme A (Ac-CoA) marks stable microtubule populations, yet chemical tools to directly measure ATAT1 ligand engagement, inhibit its activity, or visualize ATAT1-mediated modification on intact microtubules remain limited. Through the development of a quantitative binding assay, we uncovered that ATAT1 can bind unnatural cofactors but fails to efficiently use them in acyl-transfer reactions. Structure-guided mutation subsequently yielded ATAT1-L163A, which successfully installed clickable handles at the native *α*-tubulin K40 site of synthetic tubulin peptides, *α*-tubulin, and intact microtubules. Cu(I)-catalyzed azide–alkyne cycloaddition enabled visualization of modified microtubules by in-gel fluorescence and microscopy. Moreover, we report a p11-CoA bisubstrate inhibitor that suppressed both native acetylation and engineered acylation. Together, these tools provide chemically controlled access to ATAT1 activity and a site-verified, clickable K40 modification on intact microtubules.

## Introduction

Microtubules are dynamic cytoskeletal polymers that organize intracellular architecture and drive essential processes including chromosome segregation, intracellular transport, cell migration, and ciliogenesis ^[1–3]^. Microtubules are assembled from *α*/*β*-tubulin heterodimers into a cylindrical structure that typically contains 13 protofilaments ^[1,4]^. Beyond the incorporation of different tubulin isotypes, microtubules are decorated with a variety of post-translational modifications (PTMs), which together have been proposed to constitute a ‘tubulin code’ that regulates the recruitment and activity of microtubule-associated proteins, motors, and modifying enzymes ^[1,2]^. One such PTM is the acetylation of lysine 40 of *α*-tubulin, a modification that marks stable microtubule structures, such as the ciliary axoneme and the midbody of dividing cells ^[5–7]^.

K40 acetylation levels are dynamic and controlled by two opposing classes of enzymes: *α*-tubulin acetyltransferase 1 (ATAT1), the major K40 *α*-tubulin acetyltransferase that uses the cofactor acetyl-coenzyme A (Ac-CoA), and the deacetylases histone deacetylase 6 (HDAC6) and sirtuin 2 (SIRT2), which remove this mark ^[8–10]^. The preferred substrates of these enzymes differ, with ATAT1 preferentially acetylating microtubules by accessing K40 within the microtubule lumen, whereas HDAC6 and SIRT2 deacetylate acetylated tubulin dimers ^[11,12]^. Structural analysis of human ATAT1 in complex with Ac-CoA reveals a Gcn5-like catalytic fold with a widened substrate-binding groove and distinct cofactor-recognition features that create an *α*-tubulin-specific binding pocket for K40 acetylation ^[11,13]^. Despite the strong association of K40 acetylation with stable microtubules, genetic loss of ATAT1 often causes surprisingly mild developmental phenotypes at the organismal level while producing clear defects in specialized processes such as sperm motility, touch sensation, neuronal morphogenesis, and ciliogenesis kinetics, supporting context-dependent functions and partial buffering by compensatory mechanisms ^[8,14]^.

Dissecting the functional consequences of ATAT1-mediated acetylation remains challenging because available tools are largely based on indirect perturbations and endpoint measurements, rather than direct target-engagement assays or methods that chemically trace ATAT1-mediated modification on intact microtubules^[6]^. Although numerous selective inhibitors have been developed for the *α*-tubulin deacetylases, selective, well-validated inhibitors of ATAT1 remain scarce ^[6,15,16]^. Biochemical studies on lysine acetyltransferases have established that bisubstrate ligands can achieve enhanced affinity, and thereby inhibition, by simultaneously engaging substrate- and cofactor-recognition elements, providing an effective strategy for their chemical inhibition^[17]^. Szyk et al. reported a bisubstrate analog targeted to ATAT1, in which they linked a 4-mer peptide corresponding to *α*-tubulin (38-41) covalently to CoA. This bisubstrate ligand was used for structure determination studies, and has a reported *IC*_50_ of 100 *µ*M for inhibition of ATAT1-mediated tubulin acetylation ^[11]^. This example illustrates that ATAT1 is amenable to inhibition by bisubstrate ligands, in line with the existence of a ternary complex between ATAT1, Ac-CoA and its tubulin substrate^[13]^. Direct methods to quantify ligand binding affinities, however, are lacking. As such, it is currently not possible to directly correlate ligand binding affinity with biochemical inhibition.

Current approaches to visualize tubulin acetylation largely rely on anti-acetylated tubulin antibodies or targeted, antibody-free proteomics ^[6,8,18]^. For other lysine acetyltransferases, engineered enzyme–cofactor pairs have enabled selective transfer of bioorthogonal acyl groups by relieving steric constraints within cofactor-binding pockets, thereby providing chemical access to enzyme-specific acylation events ^[19]^. Whether ATAT1 can recognize non-native Ac-CoA mimics, whether such recognition is sufficient for productive acyl transfer, and whether donor specificity can be engineered while preserving modification of luminal K40 on polymerized microtubules have not been established. Here, we report an integrated chemical toolbox to address these questions. A tubulin-derived peptide-CoA fluorescent tracer enabled quantitative fluorescence polarization measurements of ATAT1–ligand interactions and guided the development of a potent peptide-derived bisubstrate inhibitor with submicromolar affinity for ATAT1 that suppresses acetylation in the presence of the natural cofactor Ac-CoA. Notably, although WT-ATAT1 bound several synthetic cofactor mimics, it failed to efficiently transfer their acyl groups to tubulin, revealing that cofactor recognition and productive catalysis are not necessarily coupled. Structure-guided mutagenesis of the cofactor-binding pocket overcame this restriction, yielding the ATAT1-L163A variant, which efficiently utilized clickable Ac-CoA mimics to modify a tubulin-derived peptide, *α*-tubulin, and intact microtubules. Together, these tools provide complementary means to quantify, inhibit, and chemically redirect ATAT1 activity while enabling site-specific installation of a clickable modification at *α*-tubulin K40.

## Results

Given the modest potency of the previously reported 4-mer–CoA bisubstrate analog ^[11]^, we sought to develop a bisubstrate inhibitor that more effectively captures both substrate-recognition and cofactor-binding interactions. To investigate the contribution of the peptide part, we synthesized three peptides using standard Fmoc solid-phase peptide synthesis: a 10-mer peptide previously shown to be acetylated by ATAT1 (p10: Ac-GQMPSDKTIG-NH_2_), the 4-mer peptide included in the bisubstrate analog (p4: Ac-SDKT-NH_2_) ^[11]^, and, to probe the effect of peptide extension in the C-terminal direction rather than N-terminally to K40 as previously reported, an 11-mer corresponding to *α*-tubulin (38-48) (p11: Ac-SDKTIGGGDDS-NH_2_) (Figure 1A). We then subjected these peptides to liquid chromatography-mass spectrometry (LC-MS) analysis of ATAT1-mediated acetylation. The 11-mer was the most efficiently acetylated by ATAT1 in a time-dependent manner, as indicated by the progressive formation of product (p11-Ac) and concomitant depletion of the p11 substrate (Figure 1B, C) at both 24 and 37 °C (Figure S1), whereas little acetylation was detected for the 10-mer and none for the 4-mer (Figure S2).

**Figure 1.**
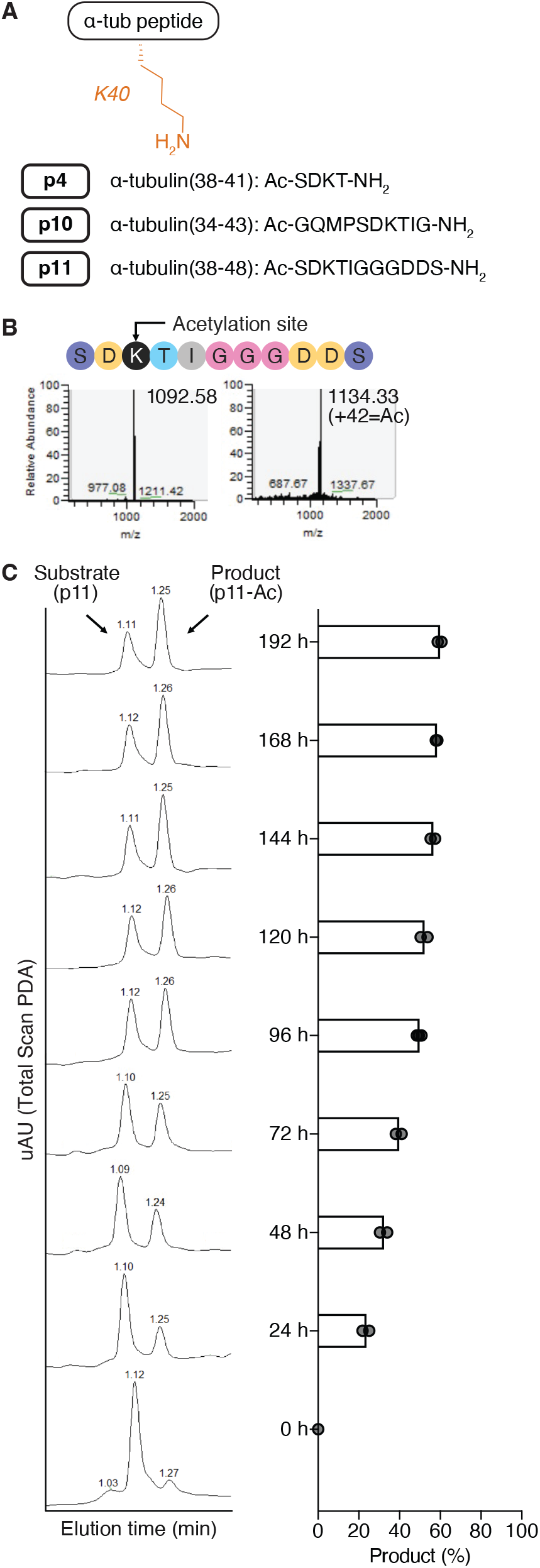
*α*-Tubulin peptide p11 enables quantitative assessment of ATAT1 acetylation activity. (A) *α*-Tubulin peptides synthesized and evaluated in this study. Amino acid sequences encompassing K40 are shown, with corresponding *α*-tubulin residue numbers indicated. (B) LC-MS assay for *α*-tubulin peptide acetylation. ESI-MS spectra of p11 substrate and acetylated p11 (p11-Ac) product, showing a 42-Da mass increase upon acetylation.Theoretical m/z values for p11 and p11-Ac are 1092.48 and 1134.49, respectively. (C) Representative LC–MS traces monitored by UV absorbance showing the time-dependent acetylation of p11 at 24 °C (375 *µ*M p11, 1.5 mM Ac-CoA, 75 *µ*M WT-ATAT1). The p11-Ac product peak was quantified from *N* = 3 independent experiments at 96 h and *N* = 2 independent experiments at all other time points.

To generate bisubstrate probes, we next covalently linked these peptides via their lysine *ε*-amino group to CoA through a short linker, yielding p4-CoA ^[11]^, p10-CoA, and p11-CoA (Figure 2A). To evaluate the binding of these probes, we set out to develop a competitive fluorescence polarization assay. For this, we designed a fluorescent tracer based on p11-CoA, in which the C-terminus of p11 was extended with a short linker and an alkyne group as a ligation handle for click conjugation with TAMRA-N_3_, yielding p11-CoA-TAMRA (Figure 2B).

**Figure 2.**
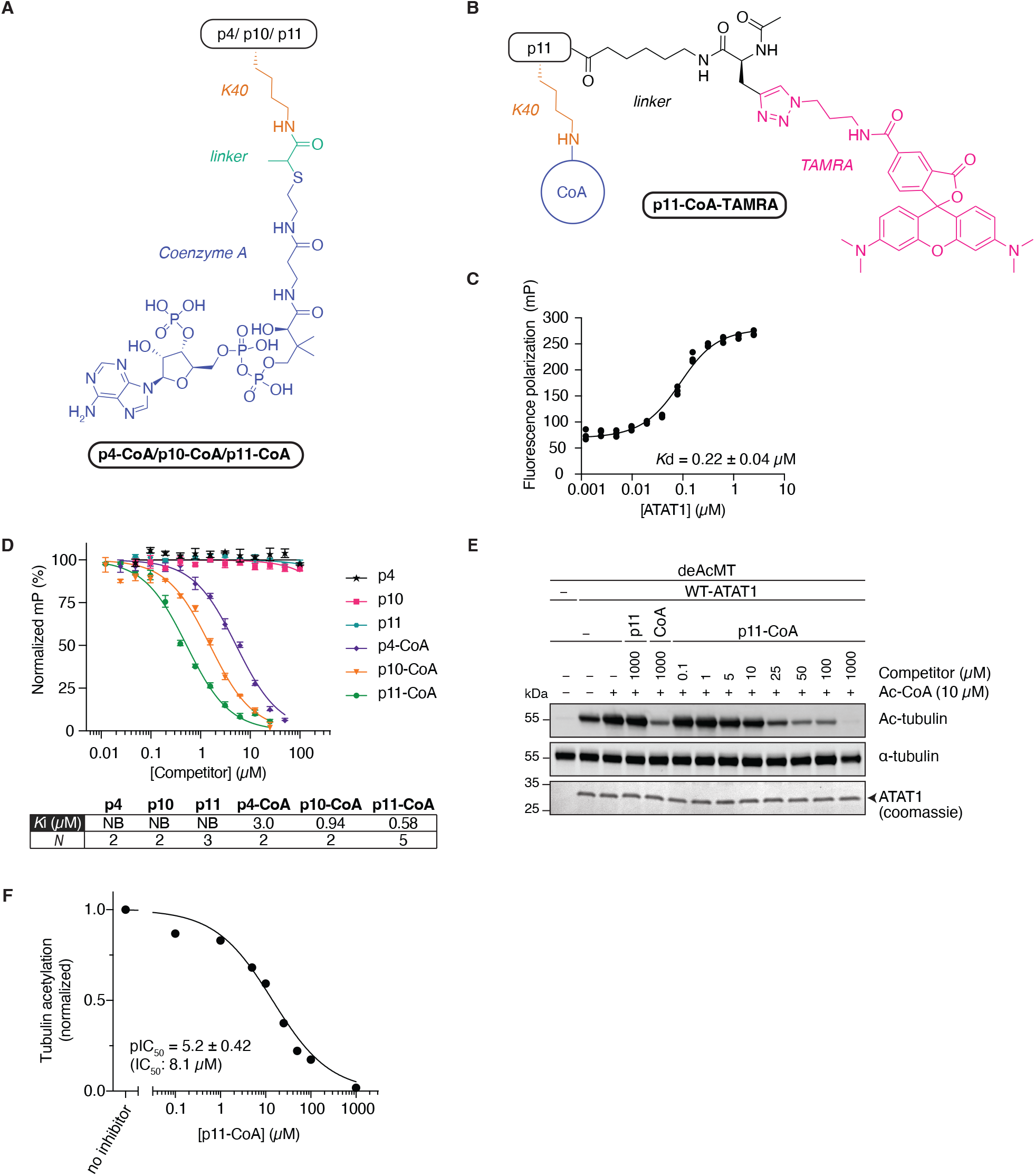
Bisubstrate probes bind ATAT1 and inhibit its acetylation activity. (A) Structures of the bisubstrate probes synthesized and evaluated in this study. (B) Structure of the bisubstrate-based tracer developed to measure ATAT1 binding affinity. (C) Representative fluorescence polarization (FP) titration of 50 nM p11-CoA-TAMRA tracer with WT-ATAT1 for 1 h at room temperature (points: technical triplicates). The mean *K*_*d*_ ± SEM from *N* = 10 independent experiments is shown. (D) Competitive FP assays to measure WT-ATAT1 binding to *α*-tubulin substrates and bisubstrate probes. Assays were performed using 500 nM WT-ATAT1 and 50 nM p11-CoA-TAMRA tracer. Representative curves (mean ± SEM of technical triplicates) and the mean *K*_*i*_ from *N* individual experiments are shown; individual fitted parameters are provided in Table S2. (E) Representative in vitro competition assay showing dose-dependent inhibition of ATAT1-mediated acetylation by p11-CoA. Deacetylated microtubules (deAcMT, 10 *µ*M), Ac-CoA (10 *µ*M), and WT-ATAT1 (2 *µ*M) were incubated with the indicated concentrations of p11-CoA at room temperature for 1 h. *α*-Tubulin acetylation was assessed by western blot. Quantification of band intensities is shown in (F). The *pIC*_50_ value (mean ± SD) was calculated from *N* = 3 independent experiments.

By titrating increasing concentrations of ATAT1 into a fixed 50 nM concentration of p11-CoA-TAMRA tracer for 1 h at room temperature, we found this tracer to be a strong binder, with a *K*_*d*_ of 220 nM for *Hs*ATAT1(2-236), validating the approach (Figure 2C). We then used p11-CoA-TAMRA in a competitive binding assay with the *α*-tubulin peptides and the bisubstrate probes, using 500 nM ATAT1 and 50 nM tracer for 1 h at room temperature (Figure 2D). We detected no binding for the peptides alone at concentrations up to 100 *µ*M. Occupation of both the substrate and cofactor pockets by the bisubstrate probes substantially enhanced the binding affinity of these probes for ATAT1, with apparent *K*_*i*_ values in the low-to submicromolar range (Figure 2D; Table S2). In line with p11 being the best substrate peptide (Figure 1B, C, Figure S2), p11-CoA was found to be the most potent binder, with an apparent *K*_*i*_ of 0.58 *µ*M (Figure 2D), and was therefore selected for all subsequent experiments.

To investigate the inhibitory potency of p11-CoA, we turned to the preferred substrate of ATAT1, microtubules. Because approximately 30% of *α*-tubulin in mammalian brain has been estimated to be acetylated at K40 ^[20,21]^, we first treated tubulin with HDAC6 to remove pre-existing acetylation before polymerizing it into microtubules (Figure 2E, first lane). We consistently found that our ATAT1 preparations co-purified with Ac-CoA: tubulin acetylation was detected even in the absence of externally added cofactor (Figure 2E, second lane; Figure S3), as was the p11-Ac product by LC-MS analysis (Figure S4). Tubulin acetylation was further increased upon addition of 10 *µ*M Ac-CoA (Figure 2E, third lane). While 1 mM p11 was unable to outcompete Ac-CoA, 1 mM CoA-SH inhibited acetylation, albeit incompletely. Gratifyingly, 1 mM p11-CoA completely blocked microtubule acetylation by ATAT1, with an *IC*_50_ of 8 *µ*M despite the presence of co-purifying Ac-CoA (Figure 2F).

Tubulin acetylation is typically studied with the use of an anti-acetylated tubulin antibody on fixed cells or by western blot. We envisioned incorporating a bioorthogonal ligation handle on microtubules as a proxy for the acetylation event, using cofactor mimics. To this end, we synthesized seven Ac-CoA mimics of different alkyl chain lengths bearing an alkyne group (3BY-CoA, 4PY-CoA, 5HY-CoA and 6HY-CoA), an azide group (3AZ-CoA and 4AZ-CoA) or a chloroacetyl group (ClAc-CoA) (Figure 3A). We then evaluated the ability of these unnatural cofactors to bind ATAT1 using our fluorescence polarization assay. The shortest alkyne derivative, 3BY-CoA, was a very poor binder (*K*_*i*_ > 50 *µ*M), as was CoA-SH, and binding affinity increased with increasing linker length for the alkyne series. The cofactors 6HY-CoA, 3AZ-CoA, 4AZ-CoA and ClAc-CoA were found to have comparable binding affinities to the natural cofactor Ac-CoA, with apparent *K*_*i*_ values in the low micromolar range (Figure 3B; Figure S5, Table S3).

**Figure 3.**
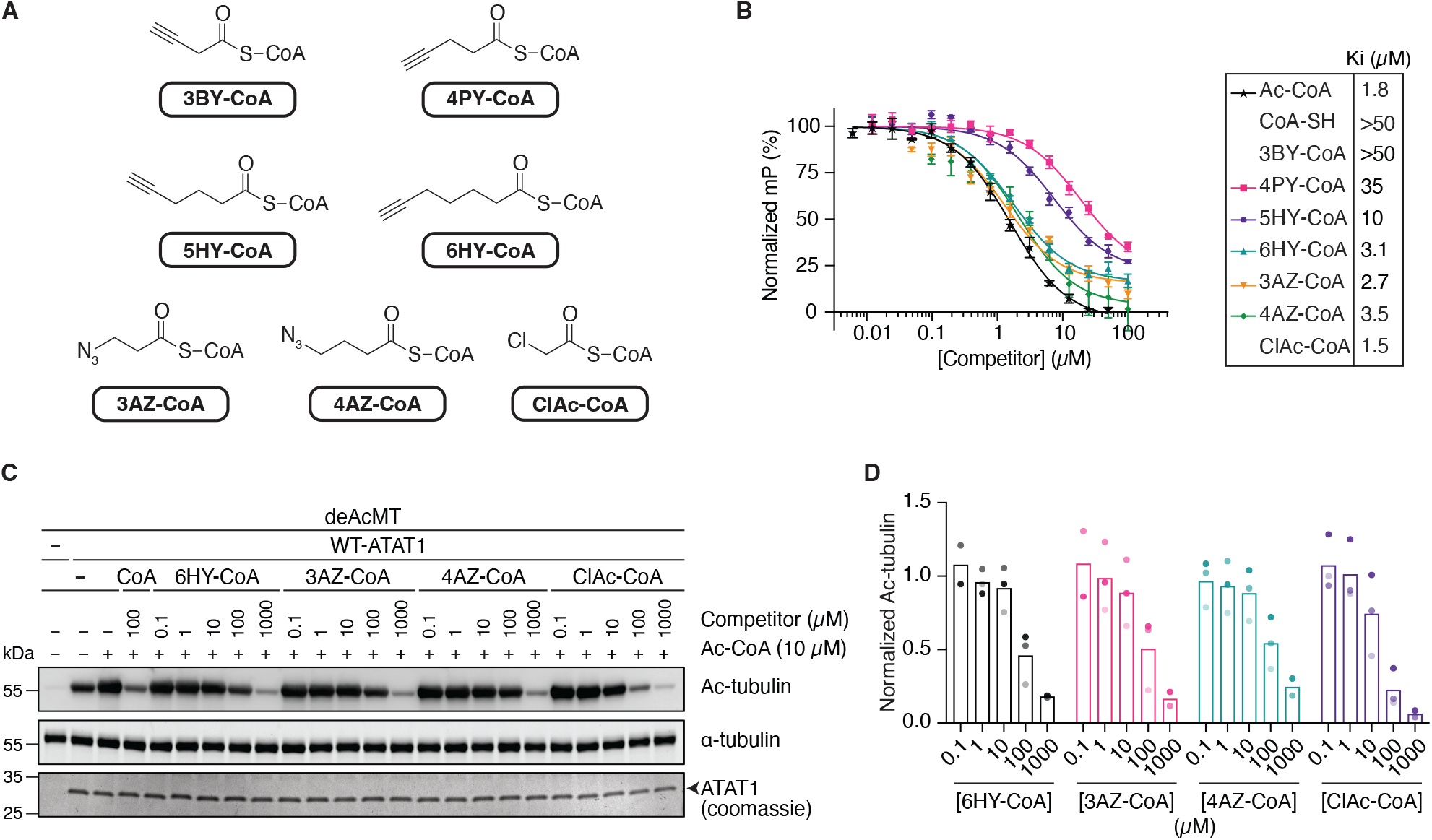
Ac-CoA mimics differentially bind WT-ATAT1 and compete with Ac-CoA-mediated acetylation. (A) Ac-CoA mimics synthesized and evaluated in this study. (B) Representative FP-based competition assays measuring WT-ATAT1 binding to Ac-CoA mimics (mean ± SEM of technical triplicates is plotted). Assays were performed using 500 nM WT-ATAT1 and 50 nM p11-CoA-TAMRA tracer. Mean *K*_*i*_ values are shown from *N* independent experiments; individual fitted parameters are provided in Table S3. *N* = 4 for Ac-CoA; *N* = 3 for CoA-SH and 4AZ-CoA; and *N* = 2 for all other Ac-CoA mimics. (C) Dose-dependent inhibition of ATAT1-mediated acetylation by Ac-CoA mimics was assessed by western blot. DeAcMT (10 *µ*M), Ac-CoA (10 *µ*M), and WT-ATAT1 (2 *µ*M) were incubated with the indicated concentrations of 6HY-, 3AZ-, 4AZ-, or ClAc-CoA at room temperature for 1 h. *α*-Tubulin acetylation was assessed by western blot. A representative blot is shown in (C) and all data quantified in (D), presented as normalized acetylated *α*-tubulin relative to total *α*-tubulin. For 6HY- and 3AZ-CoA, *N* = 3 for the 1, 10, and 100 *µ*M concentrations and *N* = 2 for the 0.1 and 1000 *µ*M concentrations. For 4AZ- and ClAc-CoA, *N* = 3 for the 0.1, 1, 10, and 100 *µ*M concentrations and *N* = 2 for the 1000 *µ*M concentration.

We next asked whether this retained cofactor binding translated into productive acyl transfer. Despite the low-micromolar binding of several mimics to WT-ATAT1, LC– MS analysis revealed essentially no transfer to the p11 substrate, with only a minor product detected for ClAc-CoA (Figure S6; Figure S7). We did observe, however, that both the UV absorbance and mass peaks corresponding to acetylated p11—generated by ATAT1-catalyzed transfer of the acetyl group from co-purifying Ac-CoA—were diminished or below the detection limit in the presence of the cofactor mimics (Figure S6; Figure S7), indicating that these mimics may act as inhibitors.

To investigate this further, we incubated deacetylated microtubules with ATAT1, Ac-CoA, and increasing concentrations of the most potent binding cofactor mimics, 6HY-CoA, 3AZ-CoA, 4AZ-CoA, and ClAc-CoA. As shown in Figure 3C, D, these cofactors indeed dose-dependently competed with Ac-CoA for ATAT1 binding, thereby preventing microtubule acetylation. Consistent with these results, 6HY-CoA also inhibited ATAT1-mediated acetylation of deacetylated tubulin (Figure S8). Despite its favorable binding and competition properties, and the minor levels of p11 acylation detected by LC-MS, further characterization of ClAc-CoA was limited by its chemical instability under the assay conditions, with the cofactor rapidly degrading at 37°C and substantially decreasing in abundance at room temperature, whereas Ac-CoA remained stable (Figure S9); additional supplementation increased product formation but not sufficiently to render ClAc-CoA a practically useful acyl donor (Figure S10).

Together, the measurable binding and competitive inhibition by the Ac-CoA mimics, yet negligible productive transfer by WT-ATAT1, indicated that cofactor recognition itself was not the principal barrier to their use as acyl donors. We therefore examined the crystal structure of ATAT1 bound to Ac-CoA (PDB 4IF5) ^[22]^, together with previously reported structural and mutagenesis studies ^[13]^, to identify residues whose mutation might relieve constraints on productive accommodation of the extended acyl groups. Analogous to the bump-and-hole approach ^[23]^, in which an enzyme active site is engineered to accommodate a modified substrate, cofactor, or inhibitor, we reasoned that a similar strategy could generate a productive enzyme–cofactor pair. Based on inspection of the crystal structure, we selected L163, I156, and I121 as candidate residues for generating a ‘hole’, as their side chains contribute to the local architecture of the cofactor-binding region. We also included R132, a residue implicated in Ac-CoA recognition and positioning by previous structural and mutational analyses ^[24]^ (Figure 4A). These residues were mutated to either glycine or alanine, and the catalytically inactive D157N mutant ^[25]^ was included as a control.

**Figure 4.**
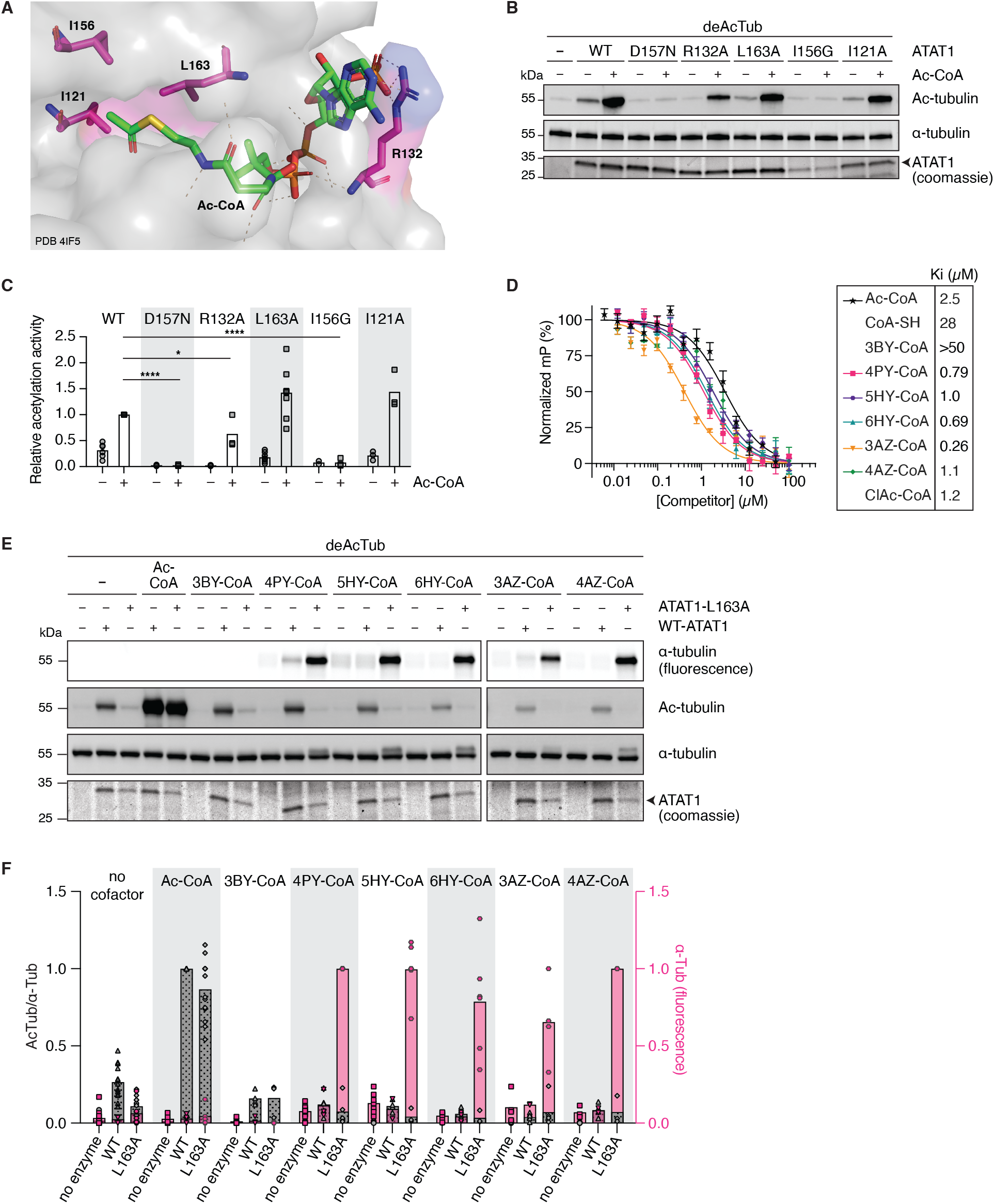
Active-site engineering of ATAT1 expands Ac-CoA mimic tolerance and enables tubulin acylation. (A) Structure-guided selection of bulky residues (pink) in the WT ATAT1 active site (PDB 4IF5)^[22]^ for substitution with alanine or glycine to expand the cofactor-binding site. The bound Ac-CoA molecule is shown in green. (B) Reduced steric bulk in the ATAT1 active site alters *α*-tubulin acetylation efficiency in vitro. Deacetylated tubulin (deAcTub; 10 *µ*M) was incubated with or without Ac-CoA (100 *µ*M) and 2 *µ*M of the indicated purified ATAT1 variant at room temperature for 24 h. *α*-Tubulin acetylation was assessed by western blot, and ATAT1 levels were assessed by coomassie staining. (C) Quantification of the data shown in (B), presented as relative acetylation activity, calculated by normalizing the acetylated tubulin/*α*-tubulin signal ratio to the corresponding ATAT1 level. *N* = 8 for ATAT1-L163A, *N* = 6 for WT-ATAT1, and *N* = 3 for all other ATAT1 variants. Statistical significance was assessed using complete datasets (*N* = 3 independent experiments) with a mixed-effects model (REML) followed by Dunnett’s multiple-comparisons test to compare each mutant with WT-ATAT1 under each reaction condition. **** *p* ≤ 0.0001; * *p* ≤ 0.05. (D) Representative FP-based competition assays (mean ± SEM of technical triplicates) measuring ATAT1-L163A binding to Ac-CoA mimics. Assays were performed using 500 nM ATAT1-L163A and 50 nM p11-CoA-TAMRA tracer. Mean *K*_*i*_ values were calculated from *N* = 2 independent experiments; individual fitted parameters are provided in Table S4. (E, F) ATAT1-L163A exhibits greater tolerance for Ac-CoA mimics than WT-ATAT1 and mediates tubulin acylation in vitro. DeAcTub (10 *µ*M) was incubated with or without Ac-CoA or Ac-CoA mimics (100 *µ*M) and 2 *µ*M ATAT1 (WT or L163A) at room temperature for 24 h, followed by click chemistry labeling. Acylated tubulin was detected by in-gel fluorescence scanning, and acetylated tubulin was detected by western blot. (E) representative gel/blot and (F) quantification, presented as acetylated *α*-tubulin band intensity or *α*-tubulin fluorescence relative to total *α*-tubulin, with western blot data (grey) normalized to WT-ATAT1 + Ac-CoA and fluorescence data (pink) normalized to ATAT1-L163A + 4PY-CoA. Bars: mean, symbols: independent experiments. See Table S6 for detailed *N* values.

As shown in Figure 4B, C, I156G purified poorly due to protein precipitation, which was not rescued by Ac-CoA addition (Figure S11). The small amount of I156G recovered was catalytically inactive. Both I121A and L163A acetylated *α*-tubulin to the same extent as the WT enzyme, whereas D157N was confirmed to be catalytically inactive and R132A showed about 62% WT activity, as expected. We were also unable to purify I121G in its expected form; it consistently eluted as a prominent, higher-molecular-weight band of unknown identity (Figure S11). We evaluated our p11-CoA-TAMRA tracer against the purified ATAT1 mutants to determine whether they retained sufficient binding to enable fluorescence polarization competition assays (Figure S12). We selected L163A as the mutant with the most robust expression and tracer-binding profiles for subsequent evaluation with the cofactor mimics. We retained R132A for evaluation because the mimics might bind in an orientation different from that of Ac-CoA, which this mutant could potentially accommodate. Using fluorescence polarization competition experiments to evaluate the binding affinities of the different cofactors for the mutant enzymes, we found that L163A retained its affinity for Ac-CoA yet bound more strongly to the mimics, with cofactor mimic affinities for L163A exceeding those for WT-ATAT1 by up to 45-fold (observed for 4PY-CoA) (Figure 4D; Figure S13, Table S4). Except for ClAc-CoA, none of the cofactors bound strongly to R132A, and we therefore excluded this mutant from the next set of experiments (Figure S14; Table S5).

We next asked whether L163A could convert the cofactor recognition observed with WT-ATAT1 into productive acyl transfer. LC-MS analysis of the p11 peptide detected acylated products for most cofactor mimics (4PY-CoA, 5HY-CoA, 6HY-CoA, 3AZ-CoA, and 4AZ-CoA) at both 24 and 37°C (Figure S15; Figure S16), indicating that L163A enables transfer of the bioorthogonal acyl groups from these cofactors.

We then subjected deacetylated tubulin to ATAT1-L163A or WT-ATAT1 in the presence of the mimics and probed acylation by copper(I)-catalyzed azide-alkyne cycloaddition (CuAAC) click chemistry (Figure 4E, F). In parallel, we probed acetylation levels by western blot, confirming that the mimics outcompeted co-purifying Ac-CoA (Figure 4E, F). The WT enzyme did not acylate tubulin, consistent with the LC-MS assay, which did not yield detectable product peaks. The L163A mutant, however, transferred the acyl group to tubulin, as shown by the fluorescent bands for 4PY-CoA, 5HY-CoA, 6HY-CoA, 3AZ-CoA, and 4AZ-CoA.As expected based on the binding affinities, 3BY-CoA was not suitable as an acyl donor. We therefore selected 4PY-CoA and 3AZ-CoA for more detailed investigations. Thus, L163A converted several cofactor mimics that bound WT-ATAT1 nonproductively into functional acyl donors for peptide and tubulin substrates.

The acylation of *α*-tubulin by L163A was a promising initial result and so we next asked whether ATAT1-L163A could acylate its natural substrate, intact microtubules ^[11]^. For this, we first deacetylated *α*-tubulin using HDAC6 before polymerizing the free tubulin into microtubules. These deacetylated microtubules were incubated with L163A in the presence or absence of a cofactor mimic (Figure 5A) and subsequently subjected to MS analysis (Figure 5B). Low-level background acetylation in no-cofactor controls was observed, consistent with the previously observed co-purification of Ac-CoA with ATAT1 (Figure 4E, F, Figure S15). Only in the presence of L163A and 4PY-CoA could we detect a mass peak corresponding to modification of K40 of *α*-tubulin with a pent-4-ynoyl group, confirming that the mutant successfully acylated the intended residue. We then compared the efficiency of incorporation by in-gel fluorescence (Figure 5C, D; Figure S17) and found that the substrate preference of the L163A mutant had not changed, as microtubules were more efficiently acetylated (in the presence of Ac-CoA) or acylated (in the presence of 4PY-CoA or 3AZ-CoA) than free *α*-tubulin. Acylation was time-dependent for both 4PY-CoA and 3AZ-CoA, with specific signal appearing at the 1 h time point. Acylation was slower than acetylation under these conditions; because click-derived background limited measurements at earlier time points, 1 h was selected as a robust endpoint (Figure 5E; Figure S18).

**Figure 5.**
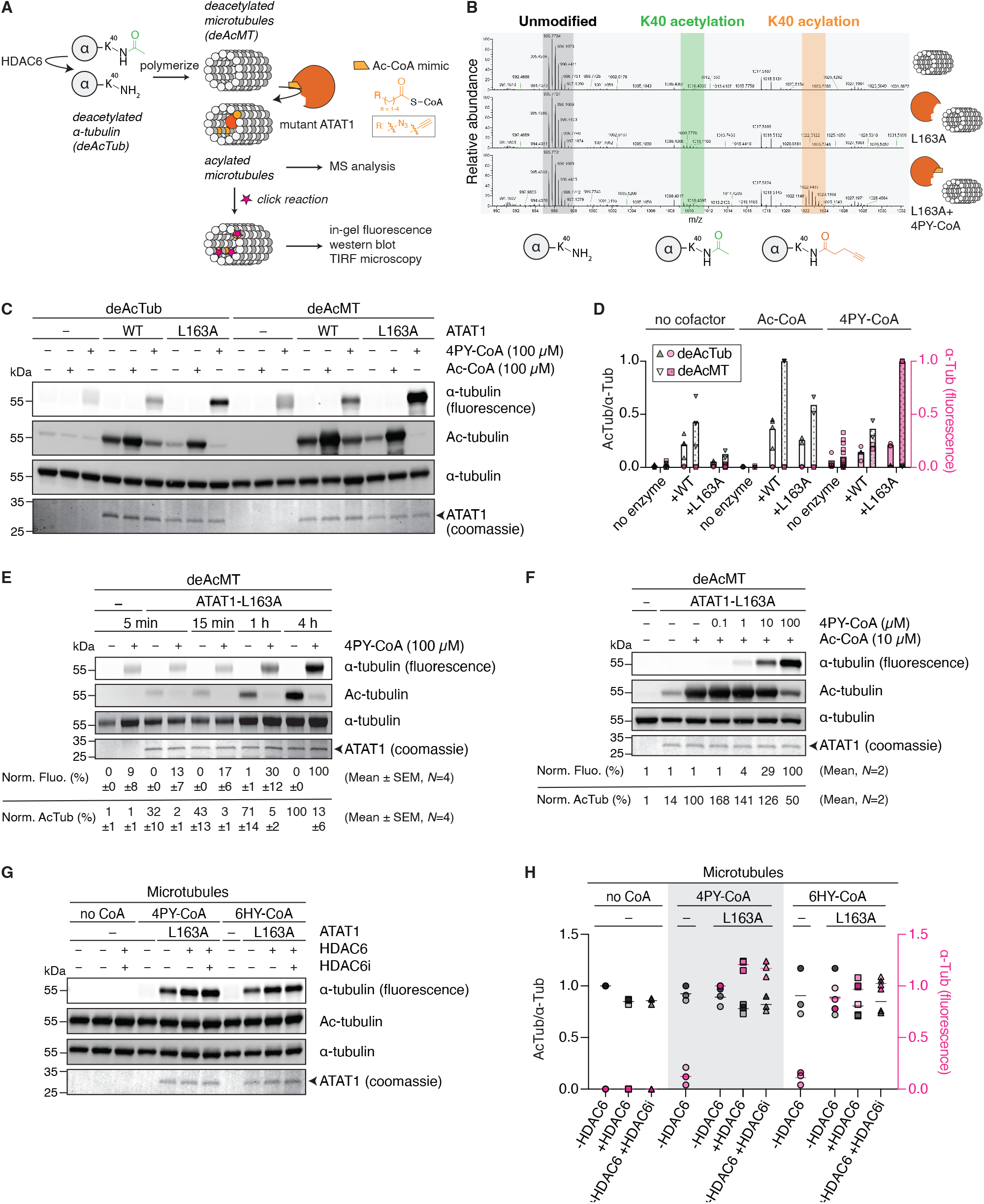
ATAT1-L163A enables time- and concentration-dependent microtubule acylation with Ac-CoA mimics and generates HDAC6-resistant modifications. (A) Workflow for detecting ATAT1-L163A-mediated acylation of deAcMT using complementary analytical readouts following incubation with Ac-CoA mimics. (B) LC–MS/MS analysis of site-specific *α*-tubulin K40 modification following incubation of deAcMT with ATAT1-L163A in the absence or presence of 4PY-CoA. Detection of the 4PY-acylated peptide (+80.0269 Da) confirms site-specific acylation at K40 by ATAT1-L163A. (C, D) ATAT1-L163A modification of soluble tubulin or microtubules. DeAcTub or deAcMT (10 *µ*M) was incubated with or without Ac-CoA or 4PY-CoA (100 *µ*M) and 2 *µ*M ATAT1 (WT or L163A) at room temperature for 1 h, followed by click chemistry labeling. Acylation was detected by in-gel fluorescence scanning, and acetylation by western blot. A representative gel/blot is shown in (C) and all data quantified in (D), presented as acetylated *α*-tubulin (grey) or *α*-tubulin fluorescence (pink) relative to total *α*-tubulin. Western blot band intensities were normalized to WT-ATAT1 + Ac-CoA and fluorescence intensities to ATAT1-L163A + 4PY-CoA. Bars: mean, symbols: independent experiments. See Table S7 for detailed *N* values. (E) Time-dependent microtubule acylation by ATAT1-L163A with 4PY-CoA in vitro. DeAcMT (10 *µ*M) was incubated with or without 4PY-CoA (100 *µ*M) and 2 *µ*M ATAT1-L163A at room temperature for the indicated times, quenched by boiling, and subjected to click chemistry labeling. Acylation was detected by in-gel fluorescence scanning and acetylation by western blot. A representative gel/blot is shown and data from *N* = 4 independent experiments are quantified. (F) Dose-dependent acylation with 4PY-CoA. DeAcMT (10 *µ*M), Ac-CoA (10 *µ*M), and ATAT1-L163A (2 *µ*M) were incubated with the indicated concentrations of 4PY-CoA at room temperature for 1 h, followed by click chemistry labeling. Acylation was detected by in-gel fluorescence scanning and acetylation by western blot. Representative gel/blot is shown and data from *N* = 2 independent experiments quantified. (G, H) Analysis of HDAC6-mediated removal of 4PY- and 6HY-acylation. Microtubules were incubated with 100 *µ*M 4PY-CoA or 6HY-CoA and 2 *µ*M ATAT1-L163A for 1 h at room temperature, followed by treatment with 1 *µ*M HDAC6, 1 *µ*M HDAC6 in the presence of 100 *µ*M Tubacin (HDAC6i), or no HDAC6 for 1.5 h. Acylated tubulin was detected after click chemistry labeling by in-gel fluorescence scanning, and acetylated tubulin by western blot. A representative blot/gel is shown in (G) and all data quantified in (H), presented as acetylated *α*-tubulin (grey) or *α*-tubulin fluorescence (pink) relative to total *α*-tubulin. Line represents the mean and symbols represent *N* = 3 independent experiments.

We next assessed whether acylation could nonetheless outcompete acetylation. Deacetylated microtubules were treated with ATAT1-L163A and increasing concentrations of 4PY-CoA or 3AZ-CoA at a fixed concentration of 10 *µ*M Ac-CoA (Figure 5F; Figure S19A). At equimolar concentrations, acylation was readily detected, and at a 10:1 ratio of mimic to Ac-CoA, acetylation was diminished in favor of acylation. To ensure that the anti-acetylated-tubulin antibody did not cross-react with the acylation marks, we split the samples post-acylation into two groups: one subjected to the click reaction and one run directly on SDS-PAGE. Reassuringly, acetylation levels were similar in both groups, confirming the specificity of the antibody for the acetyl group (Figure S19). Taken together, these results indicate that 4PY-CoA and 3AZ-CoA effectively compete with Ac-CoA for binding to ATAT1-L163A and function as acyl donors for microtubule acylation.

In contrast to ATAT1 ^[11]^, HDAC6 has been reported to preferentially deacetylate free tubulin ^[12]^, and indeed, acetylated microtubules were not deacetylated upon treatment with HDAC6 (Figure 5G, H; Figure S20), whereas *α*-tubulin was efficiently deacetylated in an HDAC6-dependent manner (Figure S21). To determine whether the acylation mark showed similar stability, we next treated acylated microtubules (Figure 5G, H; Figure S20) or acylated *α*-tubulin (Figure S21) with HDAC6 and found that all modifications from both substrate types remained intact and were thus not recognized by HDAC6.

To visualize acylation directly on microtubules, we turned to TIRF microscopy. We first treated deacetylated, taxol- and GMPCPP-stabilized microtubules containing ATTO-488- and biotin-labeled tubulin with 100 *µ*M 4PY-CoA in the presence or absence of L163A (1:5 molar ratio of microtubules to enzyme), and the resulting acylation reaction was subjected to a click reaction in solution with TAMRA-N_3_ (30 min at 37°C), before immobilizing the microtubules in a neutravidin-functionalized flow chamber, washing away unreacted dye, and imaging by TIRF microscopy. Click reactions with TAMRA-N_3_ compromised microtubule integrity, yielding fluorescently labeled tubulin aggregates only in the presence of L163A, and we therefore concluded that this dye was incompatible with our TIRF setup (Figure S22). We then decided to use an alternative fluorophore, CalFluor647-N_3_, which, by contrast, preserved microtubule integrity but produced no specific fluorescence signal under these in-solution click conditions (Figure S22). We therefore next tested whether performing the click reaction after immobilization, directly within the flow chamber, would yield specific signal. To this end, we performed acylation in solution as before, immobilized the microtubules in the flow chamber, and then carried out the click reaction with CalFluor647-N_3_ (30 min at 37°C), followed by washing and TIRF imaging. Immobilization alone did not compromise microtubule integrity, as intact (green) microtubules were readily observed prior to the click reaction (Figure S23). However, following the in-chamber click reaction, we could not detect sufficient signal above background (Figure S23), which was unexpected given that CalFluor azides have been reported to be quenched prior to conjugation and to only become fluorescent upon click reaction ^[26]^, a design intended to obviate the need for washing steps. After washing, however, we achieved an acceptable signal-to-noise ratio and readily detected dual 488/647-labeled microtubules (Figure 6A; Figure S23; Figure S24). This signal was observed exclusively in samples treated with both L163A and 4PY-CoA, confirming that the assay reported specifically on enzymatic acylation.

**Figure 6.**
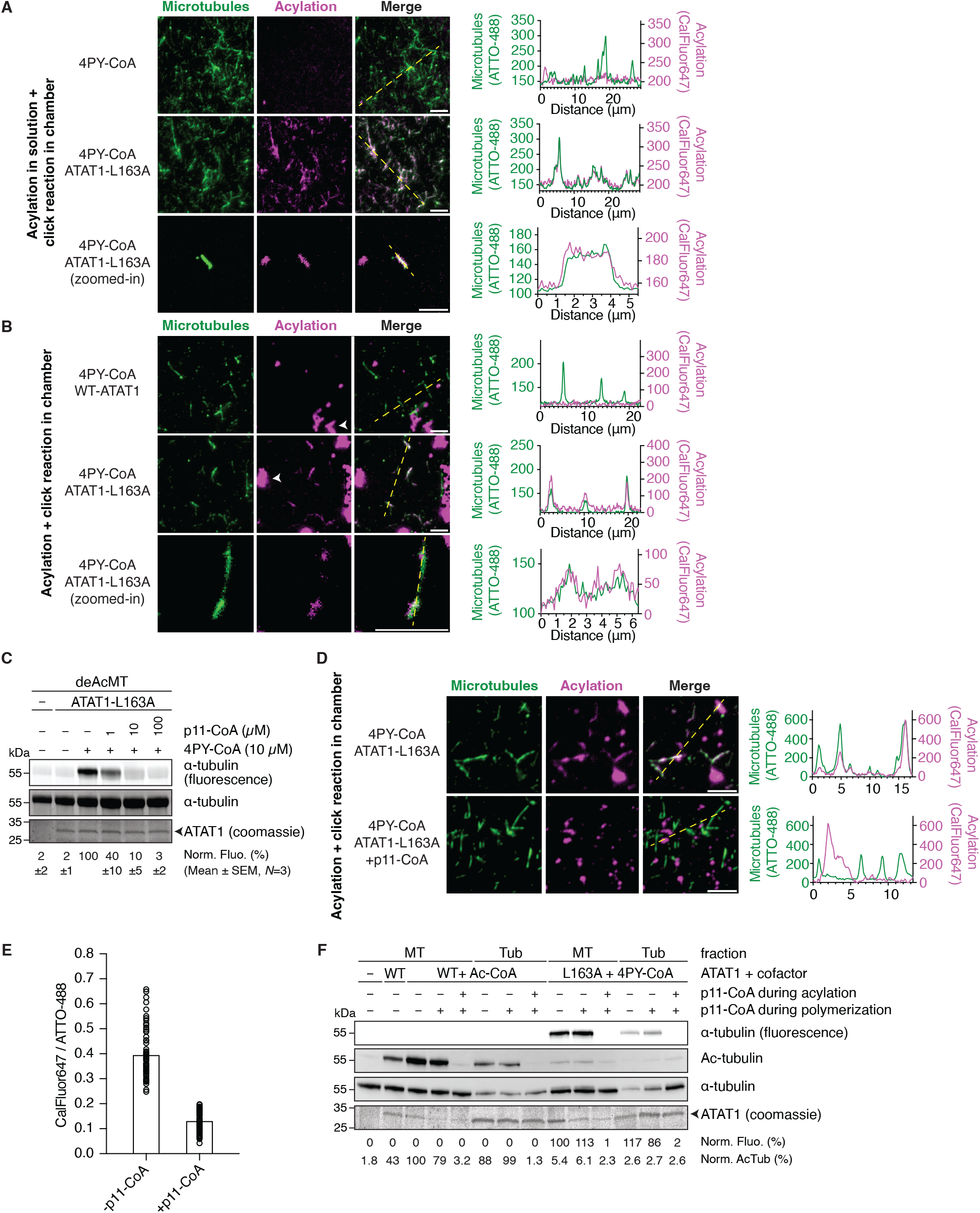
ATAT1-L163A-mediated acylation along the microtubule shaft can be inhibited by the bisubstrate probe. (A, B) TIRF microscopy reveals acylation along the microtubules. In (A), deAcMT were acylated with 4PY-CoA in solution in the presence or absence of ATAT1-L163A before immobilization, followed by in-chamber click chemistry and TIRF microscopy. In (B), deAcMT were immobilized before acylation with 4PY-CoA in the presence of WT-ATAT1 or ATAT1-L163A, followed by in-chamber click chemistry and TIRF microscopy. Representative TIRF images show ATTO-488-labeled microtubules and CalFluor647-labeled acylation signal. Arrowheads in (B) indicate large ATAT1 + 4PY-CoA–related precipitates. Scale bars, 5 *µ*m. Line profiles of the CalFluor647 fluorescence signal (640 nm channel) and corresponding ATTO-488-labeled microtubules (488 nm channel) were extracted along the regions indicated by the dashed yellow line in the merged images. *N* = 4 for (A) and *N* = 7 for (B). (C) In-gel fluorescence analysis of the competition between p11-CoA and 4PY-CoA in an ATAT1-L163A-dependent microtubule acylation assay. DeAcMT (10 *µ*M) were incubated with 4PY-CoA (10 *µ*M) and the indicated concentrations of p11-CoA in the presence of 2 *µ*M ATAT1-L163A for 1 h at room temperature, followed by click chemistry labeling and in-gel fluorescence scanning. A representative gel of *N* = 3 independent experiments is shown. (D, E) TIRF microscopy experiment evaluating the inhibition by p11-CoA of ATAT1-L163A-mediated acylation of immobilized microtubules. DeAcMT (10 *µ*M) were immobilized in flow chambers and incubated with 4PY-CoA (500 *µ*M) and ATAT1-L163A (5 *µ*M) in the presence or absence of p11-CoA (1 mM) for 1 h at room temperature, followed by in-chamber click chemistry labeling and TIRF microscopy. Representative images of ATTO-488-labeled microtubules and CalFluor647-labeled acylation signal are shown in (D). Scale bars, 5 *µ*m. Line profiles were extracted as in (A, B) and quantified in (E). Data are shown as the mean (bar) with symbols representing individual microtubule measurements (n = 62 microtubules per condition, one independent experiment). (F) Microtubule incorporation assay of acylated tubulin. DeAcTub (10 *µ*M) was incubated with Ac-CoA or 4PY-CoA (100 *µ*M) and 2 *µ*M ATAT1 (WT or L163A) for 4 h at room temperature in the presence or absence of 250 *µ*M p11-CoA. Modified tubulin was subsequently polymerized into stable microtubules in the presence or absence of inhibitor, and separated from soluble tubulin by centrifugation. Acylated tubulin in the soluble and microtubule pellet fractions was detected by click chemistry and in-gel fluorescence scanning, and acetylated *α*-tubulin was detected by western blot. A representative gel/blot is shown from *N* = 2 independent experiments.

Having developed an in-chamber click labeling protocol, we next asked whether acylation could also be performed directly on immobilized microtubules, as a step toward real-time visualization of the acylation event (Figure 6B). Microtubules were first immobilized in the flow chamber, and a solution containing 500 *µ*M 4PY-CoA in the presence or absence of 5 *µ*M ATAT1 (WT or L163A) was then introduced into the chamber and incubated for 1 h at 37°C, followed by in-chamber click labeling, washing, and TIRF imaging. This approach successfully reported on acylation, but we also observed large precipitates in reactions containing 4PY-CoA (though not Ac-CoA) together with either WT-ATAT1 or L163A. These precipitates likely correspond to cofactor-bound enzyme aggregates that were not efficiently removed by washing and were consequently also labeled during the click reaction (Figure 6B, arrowheads; Figure S25). Critically, microtubules treated with WT-ATAT1 and 4PY-CoA showed no fluorescent labeling, confirming that the wild-type enzyme remains unable to install the acyl group onto the microtubules (Figure 6B, upper panels; Figure S25). Across both click-in-chamber experimental setups, analysis of individual microtubules confirmed specific and robust co-localization of the 488 and 647 signals, with the CalFluor647 signal dispersed along the microtubule lattice (Figure 6A, B, bottom panels; Figure S24; Figure S26; Figure S27, Table S8). Although the signal appeared more intense near microtubule ends in some cases, the limited stability of microtubules under the click and wash conditions precluded any firm conclusions regarding preferential sites of acylation.

In further confirmation of the specific acylation mediated by ATAT1-L163A, we co-incubated microtubules with a fixed concentration of 4PY-CoA and ATAT1-L163A in the presence of increasing concentrations of the bisubstrate inhibitor p11-CoA. As shown in Figure 6C, p11-CoA potently inhibited acylation of deacetylated microtubules, diminishing acylation by 40% already at substoichiometric amounts (1 *µ*M p11-CoA, 2 *µ*M ATAT1-L163A, 10 *µ*M 4PY-CoA), and reducing acylation to only 3% of control levels at 100 *µ*M p11-CoA, indicating near-complete inhibition. Consistent with this, we next assessed the effect of p11-CoA by TIRF microscopy. Microtubules immobilized in the flow chamber were incubated with 500 *µ*M 4PY-CoA and 5 *µ*M ATAT1-L163A, with or without 1 mM p11-CoA, for 1 h at room temperature. In the absence of p11-CoA, microtubules treated with ATAT1-L163A and 4PY-CoA displayed dual ATTO-488/CalFluor647 labeling, whereas in the presence of p11-CoA, microtubules remained ATTO-488-positive but showed no specific CalFluor647 signal, indicating complete inhibition of acylation (Figure 6D, E).

Having established that ATAT1-L163A successfully installs a bioorthogonal ligation handle onto intact, immobilized microtubules, and that this reaction can be inhibited by p11-CoA, we next asked whether acylation affects the ability of tubulin to polymerize into microtubules. To this end, we incubated deacetylated tubulin with ATAT1-L163A and 4PY-CoA, or with WT-ATAT1 and Ac-CoA, either in the presence or absence of 250 *µ*M p11-CoA, before polymerizing the tubulin into microtubules, pelleting the resulting microtubules, and separating the supernatant containing non-polymerized tubulin. TIRF microscopy confirmed the presence of intact microtubules upon immobilization (Figure S28), and subsequent in-chamber click chemistry revealed that only tubulin incubated with L163A and 4PY-CoA had polymerized into acylated microtubules (Figure S29; Figure S30). Consistent with this, in-gel fluorescence and western blot analyses showed that the majority of both acetylated and acylated tubulin was successfully incorporated into microtubules (Figure 6F). As an additional control, we tested samples in which p11-CoA was added only after the modification reaction, immediately prior to polymerization, to determine whether acylation could continue–or even be favored, given ATAT1’s preference for microtubules over free tubulin–during this step. No consistent differences in acylation levels were observed between these samples and the non-inhibited control, however, indicating that acylation occurred prior to polymerization, on non-polymerized tubulin, and that the presence of the acyl group does not prevent tubulin from polymerizing into microtubules.

## Discussion

Here, we establish an integrated chemical toolbox that links quantitative ATAT1 ligand engagement to functional inhibition and catalytic output. These approaches provide complementary strategies to inhibit ATAT1 activity and reprogram its cofactor use. Structure-guided engineering further enabled installation of chemically addressable acyl marks at K40 on intact microtubules.

Previous ATAT1 assays have primarily monitored net acetyl transfer to peptide, tubulin, or microtubule substrates and therefore conflate ligand binding with downstream catalytic steps ^[11,22,25,27,28]^. To address ligand binding directly, we synthesized a fluorescent bisubstrate analog, p11-CoA-TAMRA, which allowed quantitative measurements of tracer binding to ATAT1 variants (Figure S12), with a *K*_*d*_ of 220 nM for WT-ATAT1 (Figure 2C). The assay further enabled competitive determination of apparent affinities for structurally diverse ligands (Figure 2D; Figure 3B; Figure 4D; Figure S14). Beyond providing a platform for ligand profiling and inhibitor discovery, the assay enabled ligand recognition to be evaluated independently of catalytic output, thereby providing insights into the molecular mechanisms of ATAT1-mediated tubulin acetylation.

Indeed, while we showed that p11 peptide is a poor but functional substrate, no binding was observed up to 100 *µ*M (Figure 1B,C; Figure 2D). Possibly, and in line with mechanisms found for other Gcn5-like acetyltransferases ^[29,30]^, ATAT1 operates with an ordered Ac-CoA first mechanism. As the tracer in our FP assay is based on a bisubstrate analog and thus occupies both pockets, its displacement would leave the CoA pocket unoccupied, which in turn could impact the affinity of p11. While such detailed mechanistic investigations are beyond the scope of this work, they nonetheless present exciting future applications for the developed assay. Enhanced affinity for substrate peptides in the presence of CoA also presents an additional argument in favor of inhibitors based on bisubstrate analogs. Indeed, neither the free peptides nor CoA alone measurably competed for tracer binding, whereas covalent peptide–CoA linkage generated ligands with low- to submicromolar apparent affinities (Figure 2D; Figure 3B), consistent with simultaneous engagement of cofactor- and substrate-recognition determinants ^[31]^. p11-CoA was the highest-affinity bisubstrate ligand and inhibited both native microtubule acetylation and L163A-dependent bioorthogonal acylation (Figure 2E,F; Figure 6C–E), providing a means to chemically suppress both native and engineered ATAT1 activity.

As a further illustration that protein engagement does not necessarily translate to catalytic activity, we found that several Ac-CoA mimics bound WT-ATAT1 with low micromolar apparent affinities and competitively inhibited acetylation, yet supported little or no detectable acyl transfer to p11 or tubulin (Figure 3B–D; Figure S6; Figure S7; Figure 4E,F). Thus, cofactor recognition and competition with Ac-CoA were insufficient to predict productive donor utilization by WT-ATAT1. In contrast, L163A retained WT-like acetylation activity toward tubulin and apparent affinity for Ac-CoA while increasing the apparent affinity of several cofactor mimics, most notably 4PY-CoA (Figure 4B–D). L163A also enabled transfer of five of seven tested mimics to p11 (Figure S15; Figure S16) and supported click-detectable tubulin acylation with the same alkyne- and azide-bearing donors (Figure 4E,F). These data identify Leu163 as an important determinant of ATAT1 permissiveness toward sterically extended acyl donors and are consistent with its proximity to the Ac-CoA-binding pocket (PDB 4IF5 ^[22]^; Figure 4A) and with previous engineering of lysine acetyltransferases by relieving steric constraints in the cofactor-binding site ^[19]^. The precise structural basis of the L163A effect remains unresolved and will require structural and kinetic analysis of the mutant with compatible Ac-CoA mimics.

A key validation of the engineered system was confirming that ATAT1-L163A retains activity toward the relevant polymeric substrate. MS directly identified pent-4-ynoylation of *α*-tubulin K40 in reactions containing L163A and 4PY-CoA (Figure 5B), and acetylation and bioorthogonal acylation produced stronger endpoint signals with microtubules than with soluble tubulin (Figure 5C,D; Figure S17), consistent with the reported preference of ATAT1 for polymerized microtubules ^[11]^. Because K40 is located within the microtubule lumen, productive modification requires access to a confined compartment not represented in peptide or soluble-tubulin assays ^[32,33]^. Both 4PY-CoA and 3AZ-CoA also functioned as acyl donors in the presence of Ac-CoA, with bioorthogonal acylation increasing when the mimics were supplied in excess (Figure 5F; Figure S19A). Together, these findings show that the engineered cofactor specificity is retained on the relevant polymeric substrate while preserving access to luminal K40.

4PY-CoA further enabled direct visualization of L163A-installed microtubule acylation. Following click labeling with CalFluor647-N_3_, the fluorescent signal colocalized with pre-labeled intact microtubules only in reactions containing ATAT1-L163A and 4PY-CoA; it was absent with WT ATAT1 and abolished by p11-CoA (Figure 6A,B,D,E). These controls extend the platform from bulk biochemical measurements to visualization of individual microtubules. Although the workflow currently provides an endpoint rather than real-time readout because fluorescence detection requires a subsequent click reaction, it establishes a tractable platform for examining engineered ATAT1 activity at the single-micro-tubule level.

The TIRF workflow also revealed assay-specific limitations. TAMRA-N_3_ disrupted microtubule integrity, whereas CalFluor647-N_3_ preserved microtubules but required washing to achieve adequate signal-to-noise (Figure S22; Figure S23). These effects currently limit quantitative spatial and kinetic interpretation, and apparent non-uniform labeling should therefore be interpreted cautiously. Improved click reagents or directly fluorogenic donors may enable quantitative mapping of ATAT1 activity on individual microtubules ^[26]^.

Finally, the installed unnatural K40 acyl marks were not detectably removed by HDAC6 and remained compatible with tubulin polymerization (Figure 5G,H; Figure 6F; Figure S20; Figure S21). HDAC6 resistance may reflect poor accommodation of the extended acyl groups in the deacetylase active site^[34,35]^. Incorporation of acylated tubulin into microtubules establishes compatibility with assembly, although effects on polymerization kinetics, lattice mechanics, and interactions with motors or microtubule-associated proteins remain to be determined ^[21,36]^.

## Conclusion

In conclusion, we here present complementary chemical tools that collectively were able to interrogate and reprogram ATAT1 activity. We show that quantitative measurements of ligand engagement - either inhibitors or cofactors - provide important insights in how binding relates to biochemical activity. Several synthetic cofactor mimics bound WT-ATAT1 and competed with Ac-CoA, thereby inhibiting native acetylation, yet were themselves inefficiently transferred. In contrast, the engineered L163A variant converted several of these bound mimics into functional acyl donors, enabling installation of a clickable acyl mark on intact microtubules and subsequent visualization. Extension to cellular settings will require validation of cofactor delivery, competition with endogenous Ac-CoA, and selectivity in complex proteomes; even so, the reconstituted system already supports mechanistic studies of K40-directed modifications on intact microtubules. More broadly, this work provides a framework for developing integrated chemical biology toolkits tailored to other lysine acetyltransferases, combining quantitative ligand-binding assays, bisubstrate-based inhibition, and engineered enzyme–cofactor pairs for site-selective substrate acylation.

## Supporting Information

Supplementary Information contains Supplementary Figures S1-S30 and Tables S1-S8, Supplementary Schemes S1-S6, full experimental procedures (biology and chemistry), analytical data (NMR spectra) and uncropped western blots.

## Acknowledgements

The authors thank Marie-Claire Velluz for help with HDAC6 protein purification, Mikhail Anisimov for the preparation of flow chambers, and Lorenzo Bizarri from the ChemBioMS mass spectrometry core facility (Faculty of Science, University of Geneva) for bottom-up proteomics experiments. This research was funded by the University of Geneva, the Swiss National Science Foundation (project grants 189246 and 10000608 to S. H., and TMSGI3_211433 to C. A.) and supported by funding from the European Research Council (ERC) under the European Union’s Horizon 2020 research and innovation programme (grant agreement n° 948750, DestCilia, to S.H.).

