## Supplemental Information for "Chemical Interrogation and Reprogramming of ATAT1-Mediated Tubulin Acetylation"

### SUPPORTING INFORMATION

#### SUPPLEMENTARY FIGURES

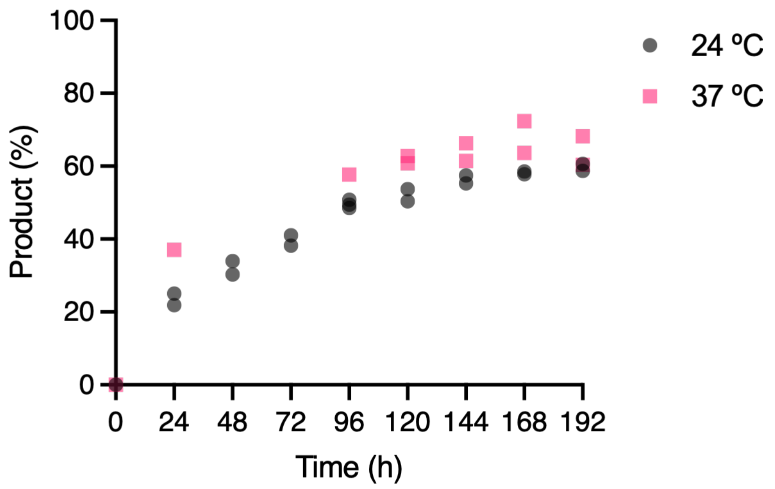

**Figure S1.** Quantification of the LC–MS-based ATAT1 acetylation assay monitoring the time-dependent acetylation of p11 at 24 and 37 °C (375 $\mu$M p11, 1.5 mM Ac-CoA, 75 $\mu$M WT-ATAT1). Data obtained at 24 °C are reproduced from Figure 1C for visualization. The numbers of independent experiments were as follows: at 24 °C, $N = 3$ at 96 h and $N = 2$ at all other time points; at 37 °C, $N = 2$ at 120–192 h and $N = 1$ at 24 and 96 h.

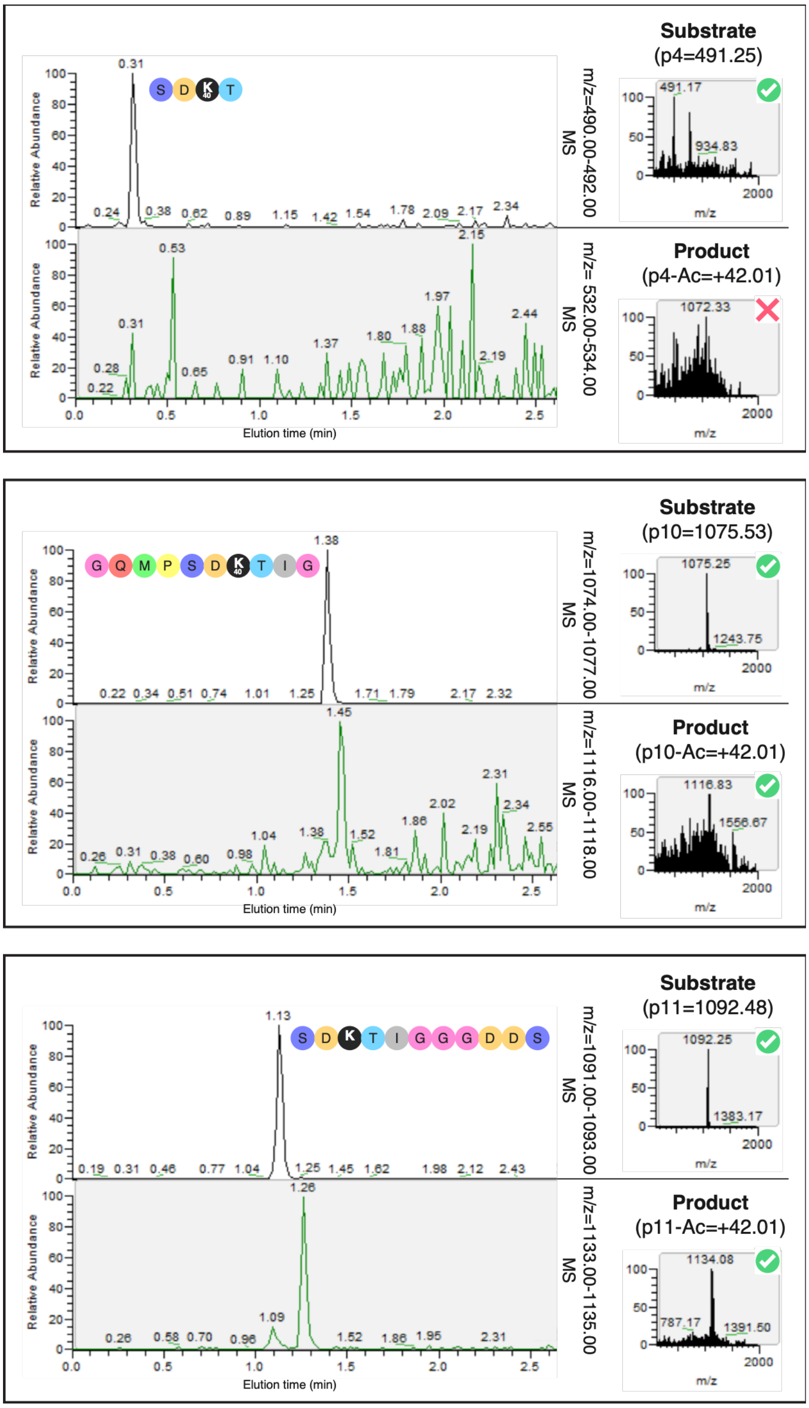

**Figure S2.** Extracted ion chromatograms showing the indicated m/z ranges for WT-ATAT1-mediated acetylation of α-tubulin peptide p4, p10, and p11 following 72 h of incubation at 24 °C (100 $\mu$M α-tubulin peptide, 500 $\mu$M Ac-CoA, 50 $\mu$M WT-ATAT1). Theoretical m/z values are indicated. Check marks (✓) indicate agreement between observed and theoretical m/z values; crosses (✗) indicate discrepancies, as observed for p4 acetylation. The numbers of independent experiments were $N = 1$ for p4 and $N = 2$ for p10 and p11.

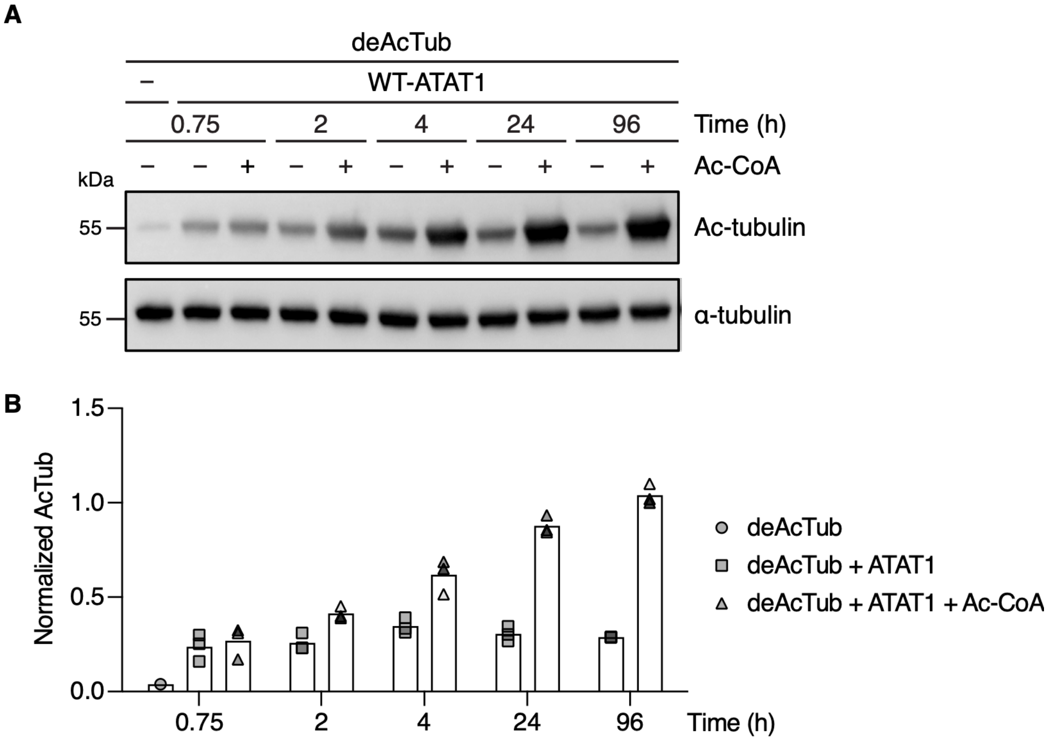

**Figure S3.** Detection of ATAT1-mediated acetylation from co-purifying Ac-CoA using the WB-based assay. (A) Representative WB-based ATAT1 activity assay showing enzymatic acetylation activity and the contribution of co-purifying Ac-CoA over time. Deacetylated tubulin (deAcTub, 10 µM) was incubated with or without externally added Ac-CoA (100 µM) and WT-ATAT1 (2 µM) at room temperature for the indicated times. (B) Quantification of the data shown in (A), presented as normalized acetylated α-tubulin relative to total α-tubulin. *N* = 3 independent experiments.

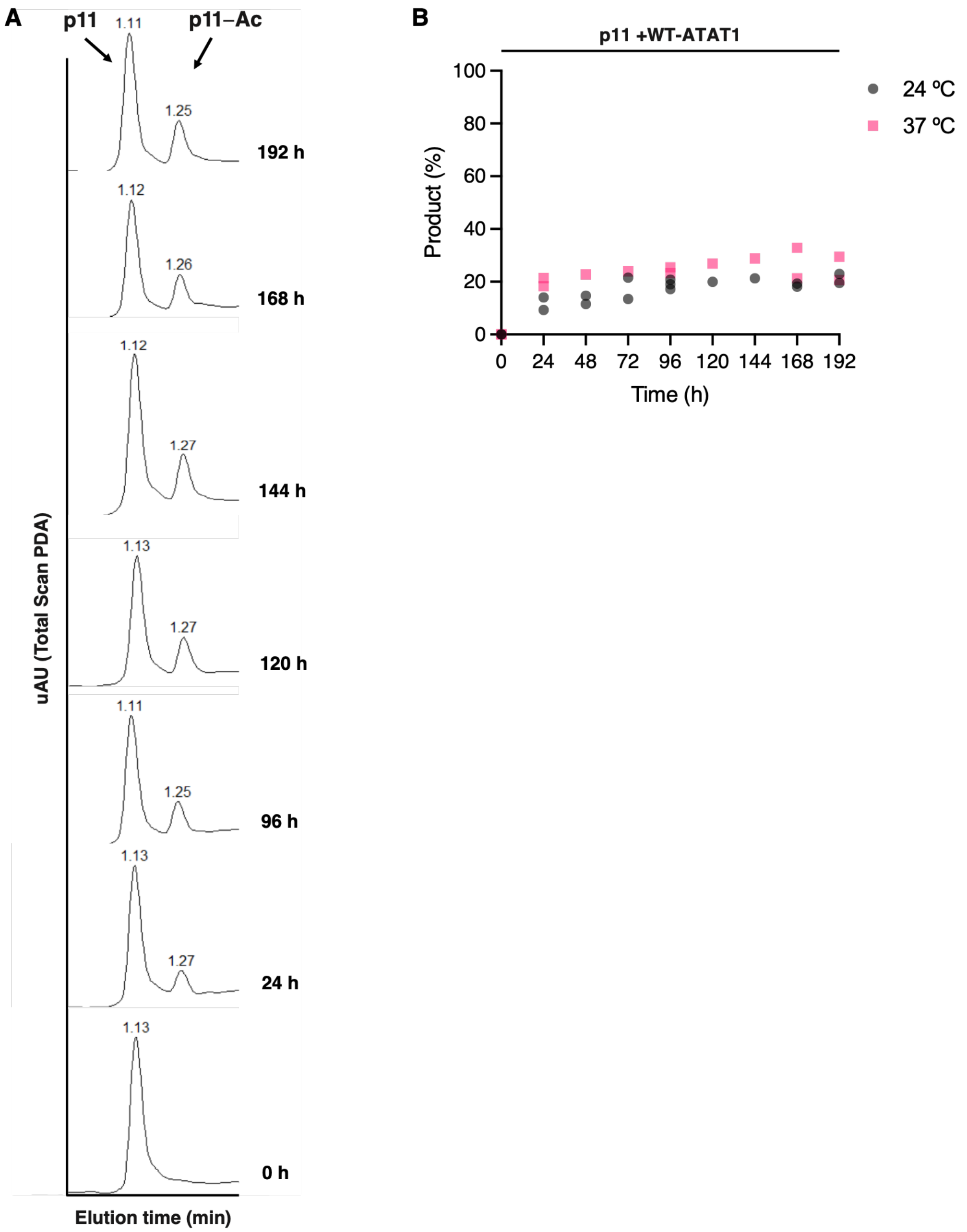

**Figure S4.** (A) Representative LC–MS traces monitored by UV absorbance showing the time-dependent acetylation of p11 at 24 ºC in the absence of externally added Ac-CoA (375 µM p11 and 75 µM WT-ATAT1). (B) LC–MS quantification of the p11-Ac product peak in the absence of externally added Ac-CoA over time at 24 and 37 ºC (375 µM p11 and 75 µM WT-ATAT1). The numbers of independent experiments were as follows: at 24 ºC, $N = 3$ at 96 h, $N = 2$ at 0, 24, 48, 72, 168, and 192 h, and $N = 1$ at 120 and 144 h; at 37 ºC, $N = 2$ at 0, 24, 96, 168, and 192 h, and $N = 1$ at 48, 72, 120, and 144 h.

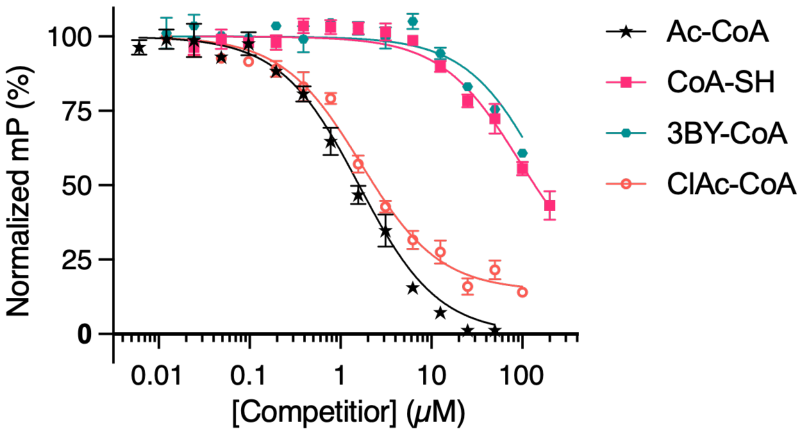

**Figure S5.** Representative fitted curves from fluorescence polarization (FP)-based competition assays measuring WT-ATAT1 binding to Ac-CoA mimics not shown in Figure 3B (500 nM WT-ATAT1 and 50 nM p11-CoA-TAMRA tracer). The Ac-CoA curve represents the same dataset shown in Figure 3B and is reproduced for visualization purposes. Individual *K*i values and fitted parameters are provided in Table S3.

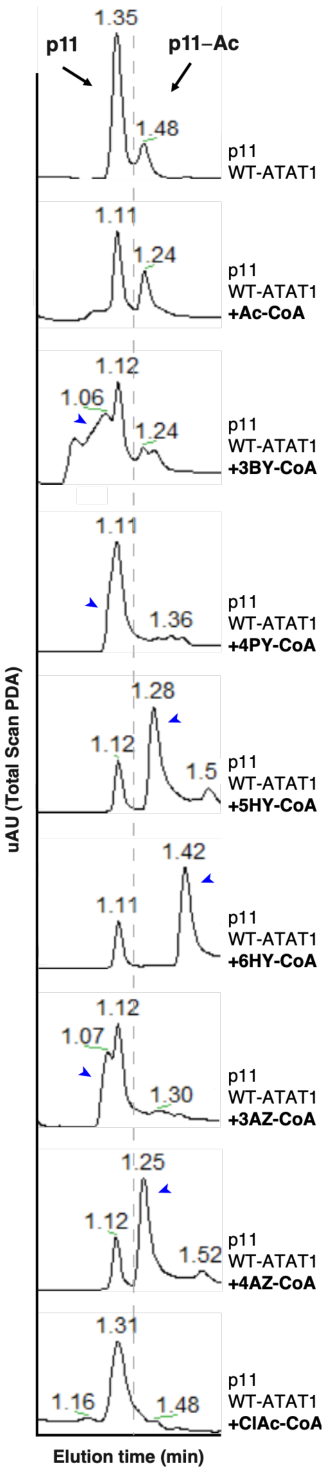

**Figure S6.** LC–MS-based in vitro screen of Ac-CoA mimics with WT-ATAT1. Representative LC–MS traces monitored by UV absorbance showing Ac-CoA mimics competing with co-purifying Ac-CoA, as indicated by reduced p11-Ac product formation relative to the p11 + WT-ATAT1 control. Reactions were incubated for 96 h at 24 °C (375 $\mu$M p11, 1.5 mM cofactor, 75 $\mu$M WT-ATAT1). The blue arrowhead indicates residual Ac-CoA mimic. The dashed gray line separates substrate and product regions; peaks are aligned across traces for visualization. $N$ = 7 independent experiments (ClAc-CoA); $N$ = 4 independent experiments (all other Ac-CoA mimics).

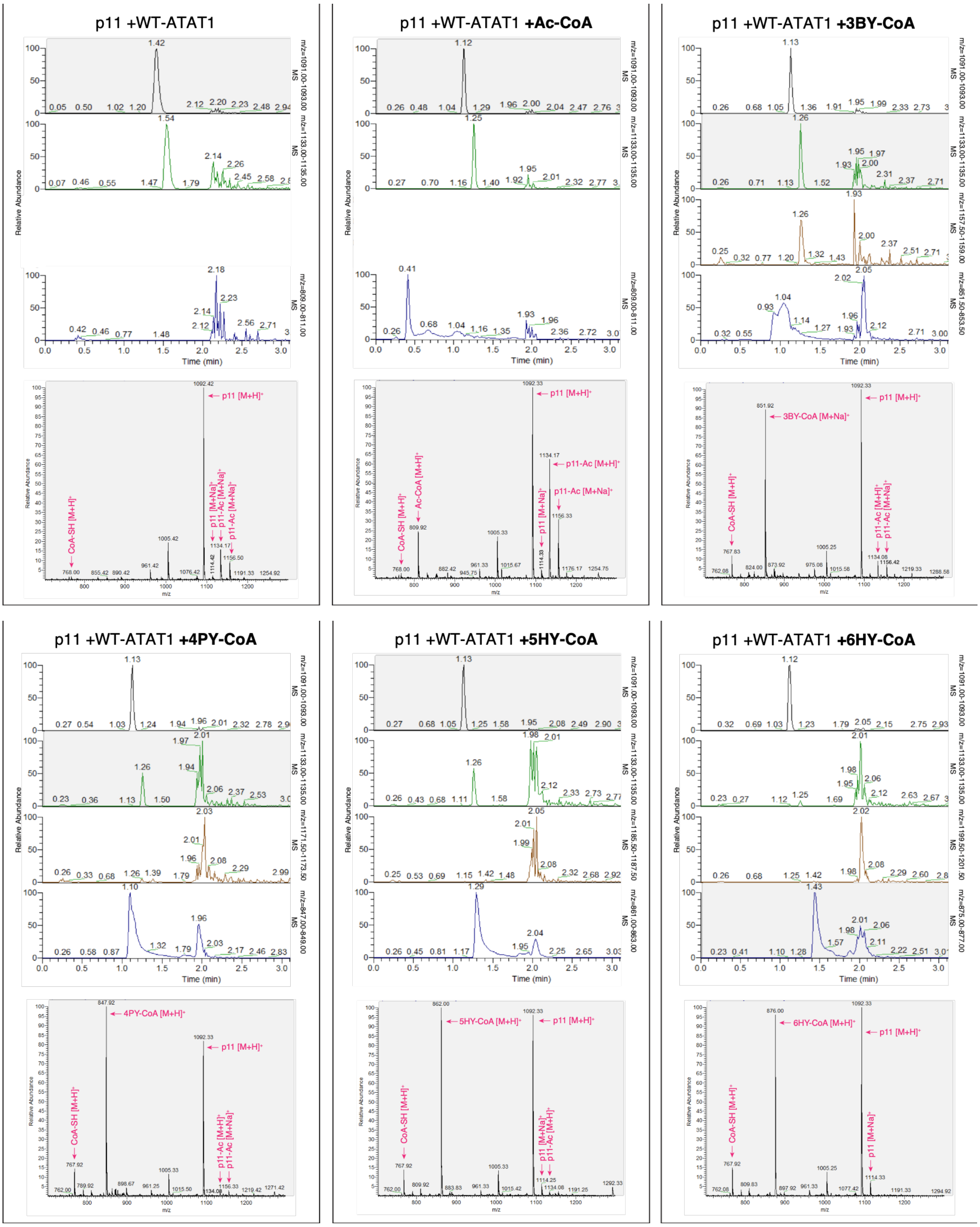
p11 +WT-ATAT1
p11 +WT-ATAT1 +Ac-CoA
p11 +WT-ATAT1 +3BY-CoA
p11 +WT-ATAT1 +4PY-CoA
p11 +WT-ATAT1 +5HY-CoA
p11 +WT-ATAT1 +6HY-CoA
Relative Abundance
Time (min)
m/z=1091.00-1093.00 MS
m/z=1133.00-1135.00 MS
m/z=809.00-811.00 MS
m/z=1157.50-1159.00 MS
m/z=851.50-853.50 MS
m/z=1171.50-1173.50 MS
m/z=847.00-849.00 MS
m/z=1185.50-1187.50 MS
m/z=861.00-863.00 MS
m/z=1199.50-1201.50 MS
m/z=875.00-877.00 MS
p11 [M+H]+
p11 [M+Na]+
p11-Ac [M+H]+
p11-Ac [M+Na]+
CoA-SH [M+H]+
Ac-CoA [M+H]+
3BY-CoA [M+Na]+
4PY-CoA [M+H]+
5HY-CoA [M+H]+
6HY-CoA [M+H]+
m/z

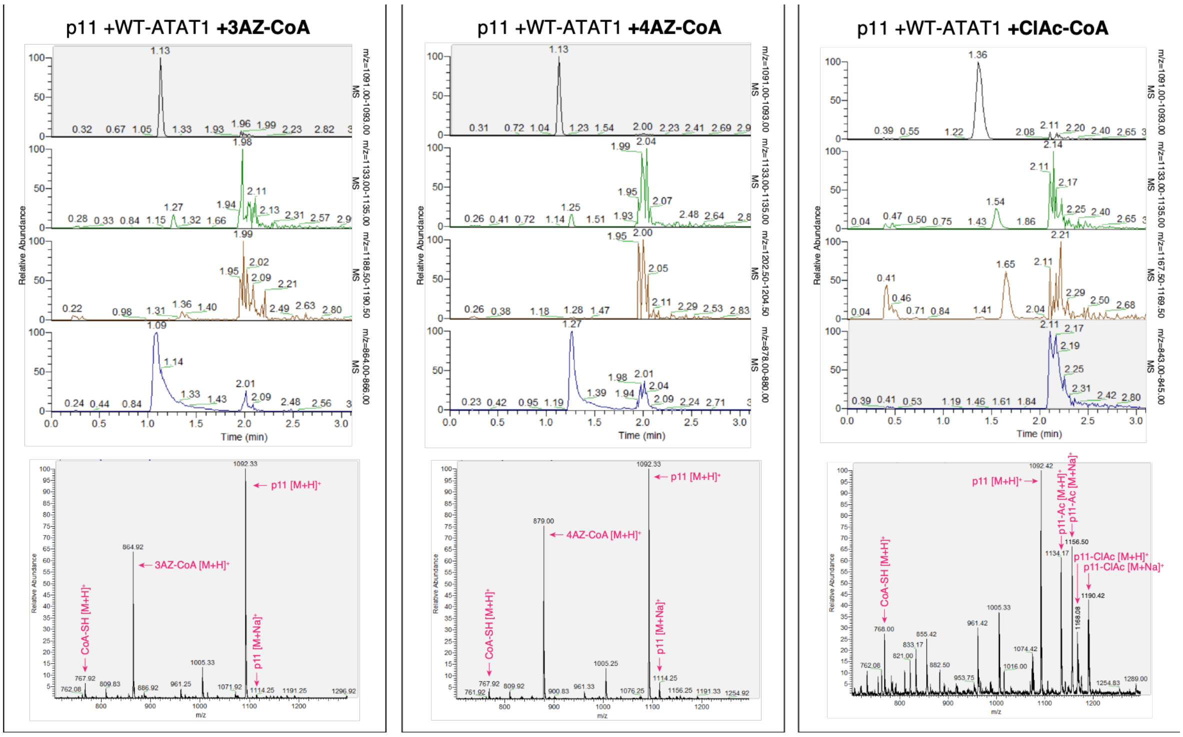

**Figure S7.** Extracted ion chromatograms (top) and extracted ESI–MS spectra (bottom) for the same WT-ATAT1 reactions shown in Figure S6, displaying the indicated m/z ranges and masses corresponding to the p11 substrate (black), p11-Ac (green), acylated p11 products (orange), and the corresponding cofactors (blue). Reactions were performed in the absence of externally added Ac-CoA or in the presence of Ac-, 3BY-, 4PY-, 5HY-, 6HY-, 3AZ-, 4AZ-, or ClAc-CoA, as indicated, and incubated for 96 h at 24 °C (375 $\mu$M p11, 1.5 mM cofactor, 75 $\mu$M WT-ATAT1). Theoretical m/z values are 1134.49 (p11-Ac), 1158.49 (p11-3BY), 1172.51 (p11-4PY), 1186.52 (p11-5HY), 1200.54 (p11-6HY), 1189.51 (p11-3AZ), 1203.52 (p11-4AZ), and 1168.45 (p11-ClAc). The mass peak around Rt = ~2 min corresponds to ATAT1. $N$ = 7 independent experiments (ClAc-CoA); $N$ = 4 independent experiments (all other Ac-CoA mimics).

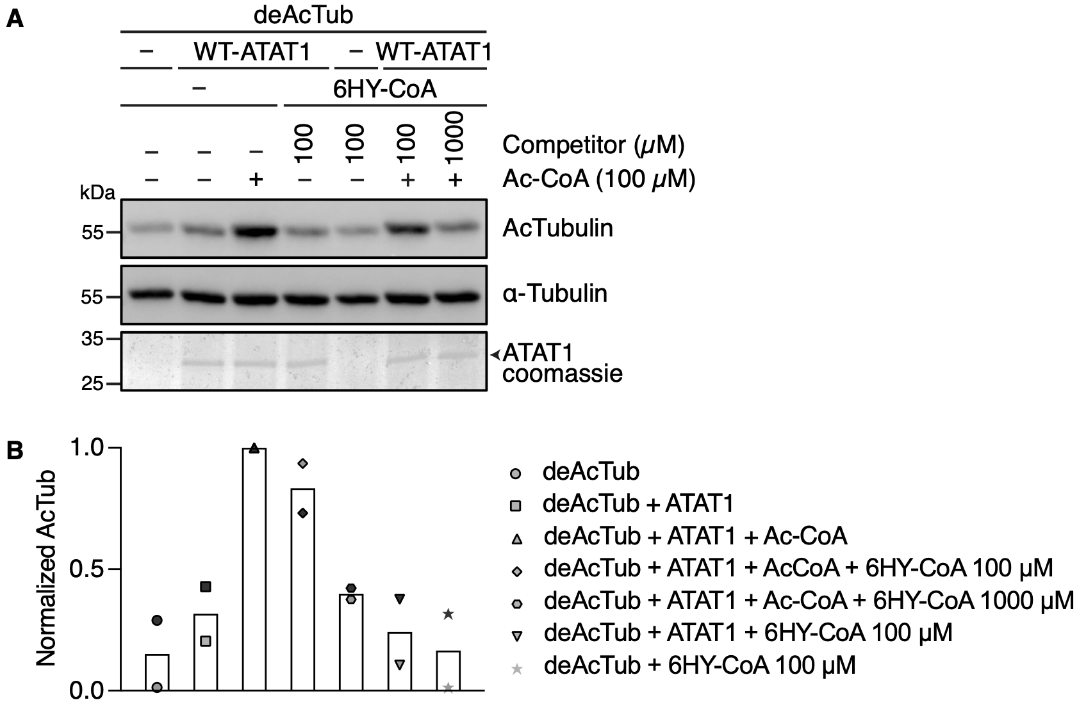

**Figure S8.** Representative in vitro competition assay showing dose-dependent inhibition of ATAT1-mediated acetylation by 6HY-CoA. (A) DeAcTub (10 µM), Ac-CoA (100 µM), and WT-ATAT1 (2 µM) were incubated with the indicated concentrations of 6HY-CoA at room temperature for 24 h. α-Tubulin acetylation was assessed by WB. (B) Quantification of the data shown in (A), presented as normalized acetylated α-tubulin relative to total α-tubulin. *N* = 2 independent experiments.

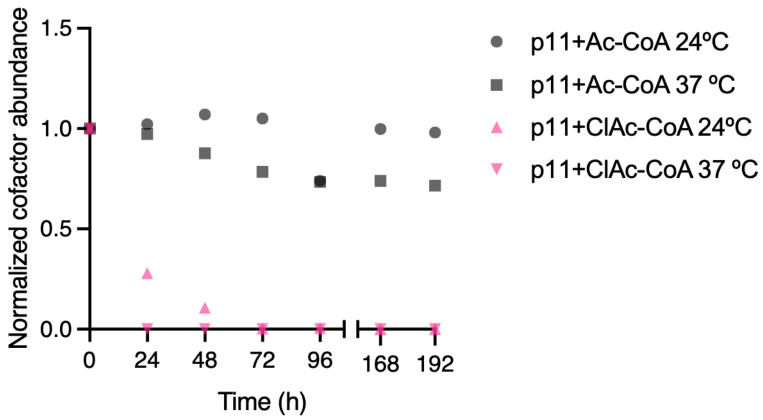

**Figure S9.** Comparison of ClAc-CoA and Ac-CoA stability at 24 and 37 °C over time (375 *µ*M p11 and 1.5 mM cofactor). Relative cofactor abundance was determined by LC–MS UV peak integration and normalized to the corresponding t=0 time point (0 h = 1). For Ac-CoA at 24 °C, *N* = 2 independent experiments (24, 48, 72, and 96 h time points) and *N* = 1 independent experiment for all other conditions.

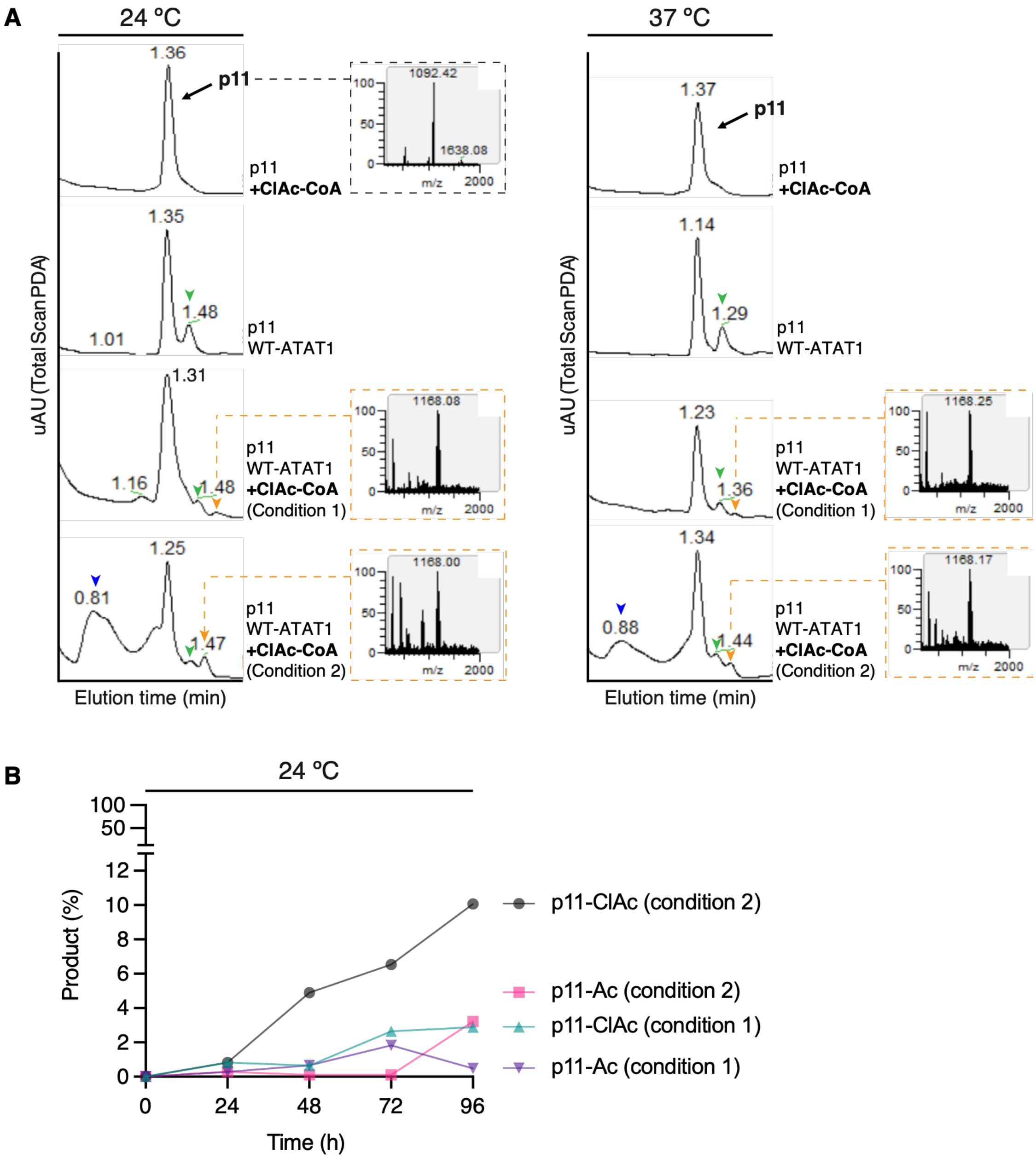

**Figure S10.** ClAc-CoA-mediated modification of p11 by WT-ATAT1. (A) Representative LC–MS traces monitored by UV absorbance showing competition between ClAc-CoA and co-purifying Ac-CoA and enzymatic transfer of the chloroacetyl group to p11 by WT-ATAT1 after 96 h at 24 °C or 37 °C, as indicated (375 $\mu$M p11, 1.5 mM ClAc-CoA, 75 $\mu$M WT-ATAT1). ClAc-CoA was added only at 0 h (Condition 1) or at 0 h and every 24 h thereafter (Condition 2). Insets show ESI-MS spectra of the p11 substrate and p11-ClAc product. The blue arrowhead indicates residual ClAc-CoA, the green arrowhead indicates acetylated p11 (p11-Ac), and the orange arrowhead indicates chloroacylated p11 (p11-ClAc). Substrate peaks are aligned for visualization. Theoretical m/z values are 1168.45 (p11-ClAc) and 1092.48 (p11). (B) Quantification of the LC–MS traces from reactions containing p11, WT-ATAT1, and ClAc-CoA at 24 °C under the two conditions described in (A). ClAc-CoA was added only at 0 h (Condition 1) or at 0 h and every 24 h thereafter (Condition 2). $N = 1$ independent experiment.

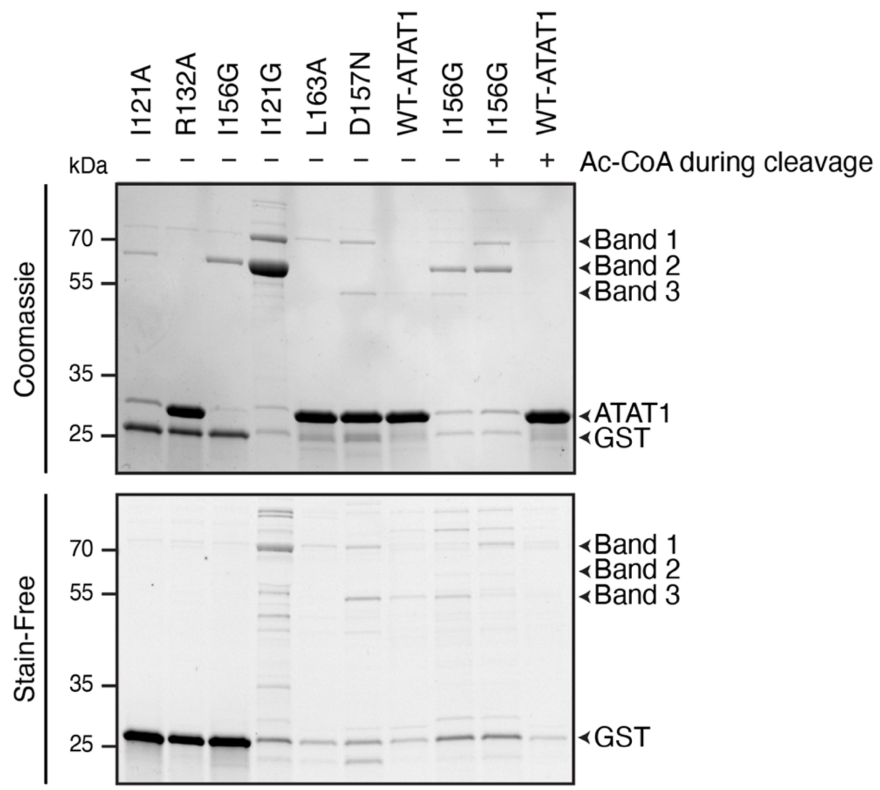

**Figure S11.** Coomassie staining and Stain-Free imaging of purified ATAT1 variants after expression and purification. To assess rescue of enzyme stability by Ac-CoA, 350 $\mu$M Ac-CoA was included during the cleavage incubation. ATAT1 and higher-molecular-weight unknown bands are indicated. Stain-Free imaging does not detect ATAT1 or the higher-molecular-weight unknown band 2.

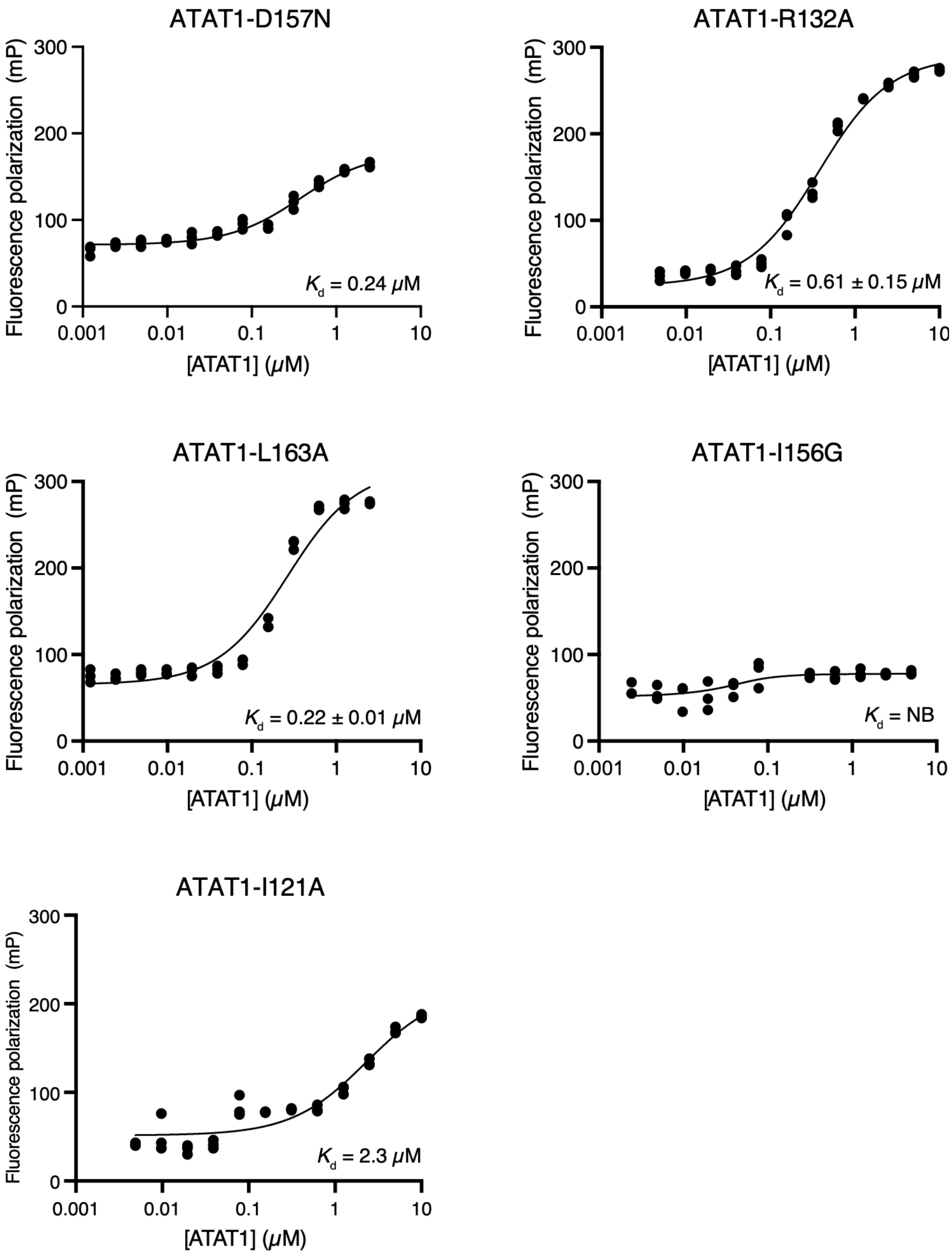

**Figure S12.** Representative FP titrations of 50 nM p11-CoA-TAMRA tracer with the indicated ATAT1 variants after 1 h of incubation at room temperature. Mean *K*d ± SEM values are shown for variants that exhibited detectable binding. The numbers of independent experiments were as follows: *N* = 2 for D157N, *N* = 3 for R132A, *N* = 4 for L163A, *N* = 2 for I156G, and *N* = 1 for I121A. NB, no binding detected.

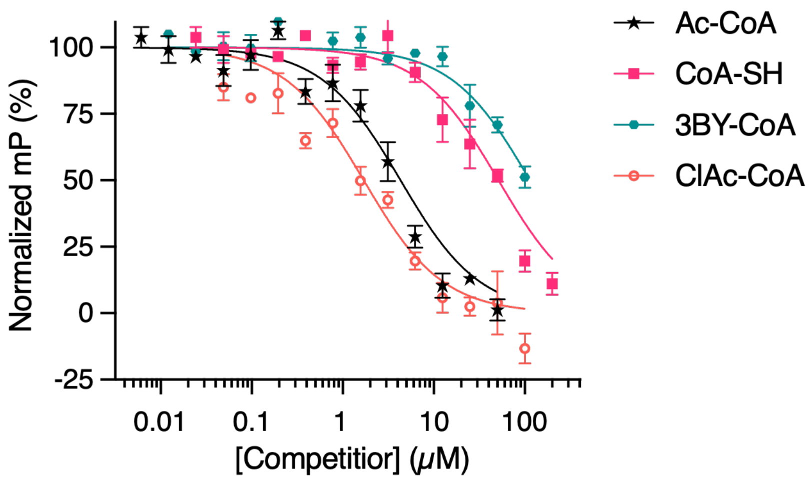

**Figure S13.** Representative fitted curves from FP-based competition assays measuring ATAT1-L163A binding to Ac-CoA mimics not shown in Figure 4D (500 nM ATAT1-L163A and 50 nM p11-CoA-TAMRA tracer). The Ac-CoA curve represents the same dataset shown in Figure 4D and is reproduced for visualization purposes. Individual *K*i values and fitted parameters are provided in Table S4.

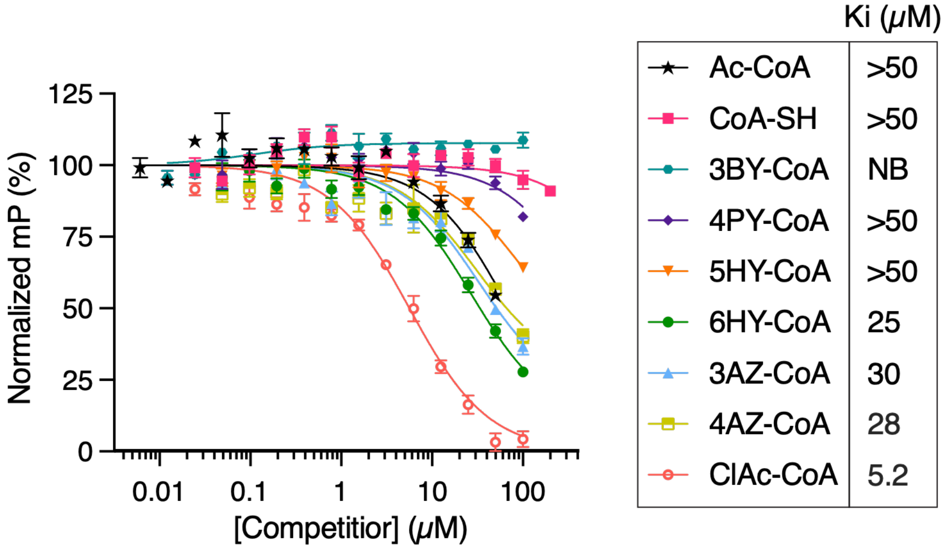

**Figure S14.** Representative FP-based competition assays (mean ± SEM of technical triplicates) measuring ATAT1-R132A binding to Ac-CoA mimics. Assays were performed using 500 nM ATAT1-R132A and 50 nM p11-CoA-TAMRA tracer. Mean Ki values were calculated from *N* = 1 independent experiment; individual fitted parameters are provided in Table S5. NB: no binding detected.

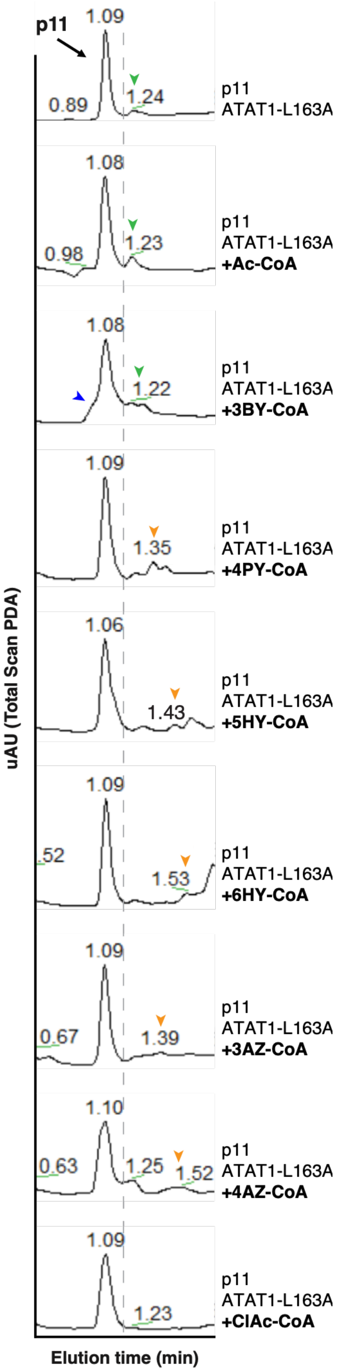

**Figure S15.** LC–MS-based in vitro screen of Ac-CoA mimics with ATAT1-L163A. Representative LC–MS traces monitored by UV absorbance showing low p11-Ac formation in the absence of externally added cofactor, consistent with co-purifying Ac-CoA, competition by Ac-CoA mimics, as indicated by the disappearance of the p11-Ac peak, and the appearance of new acylated p11 product peaks after 96 h of incubation with the cofactor mimic at 24 °C (375 $\mu$M p11, 1 mM cofactor, 75 $\mu$M ATAT1-L163A). The blue arrowhead indicates residual Ac-CoA mimic, the green arrowhead indicates p11-Ac, and the orange arrowhead indicates acylated p11 products. The dashed gray line separates substrate and product regions; peaks are aligned across traces for visualization. The absence of residual Ac-CoA mimic is consistent with its consumption during enzymatic acylation of p11, except for ClAc-CoA, which degraded under the assay conditions.

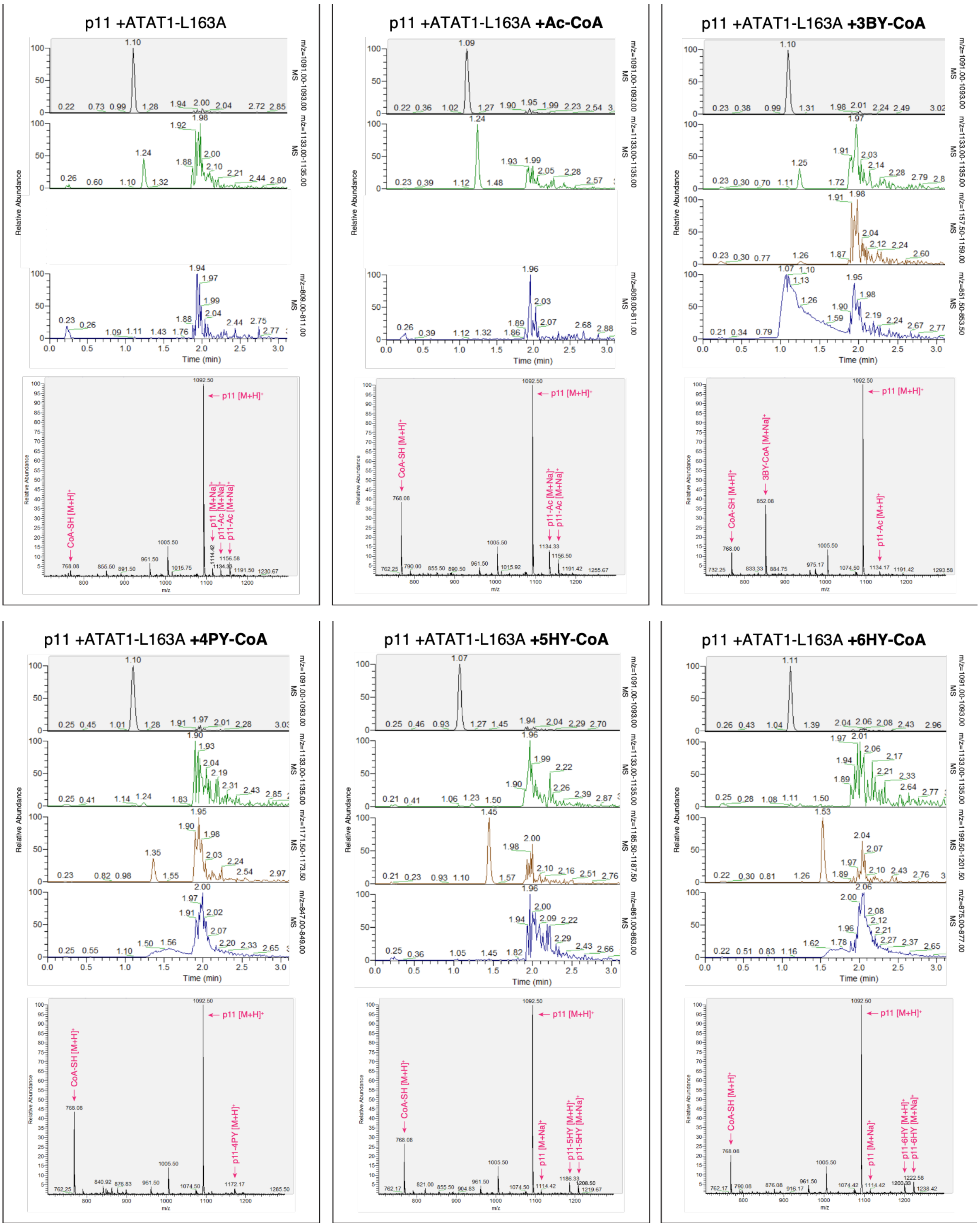
p11 +ATAT1-L163A
p11 +ATAT1-L163A +Ac-CoA
p11 +ATAT1-L163A +3BY-CoA
p11 +ATAT1-L163A +4PY-CoA
p11 +ATAT1-L163A +5HY-CoA
p11 +ATAT1-L163A +6HY-CoA
Relative Abundance
Time (min)
m/z=1091.00-1093.00
m/z=1133.00-1135.00
m/z=809.00-811.00
m/z=1157.50-1159.00
m/z=851.50-853.50
m/z=1171.50-1173.50
m/z=847.00-849.00
m/z=1185.50-1187.50
m/z=861.00-863.00
m/z=1199.50-1201.50
m/z=875.00-877.00
MS
p11 [M+H]+
CoA-SH [M+H]+
p11 [M+Na]+
p11-Ac [M+Na]+
p11-Ac [M+H]+
3BY-CoA [M+Na]+
p11-4PY [M+H]+
p11-5HY [M+H]+
p11-5HY [M+Na]+
p11-6HY [M+H]+
p11-6HY [M+Na]+
m/z

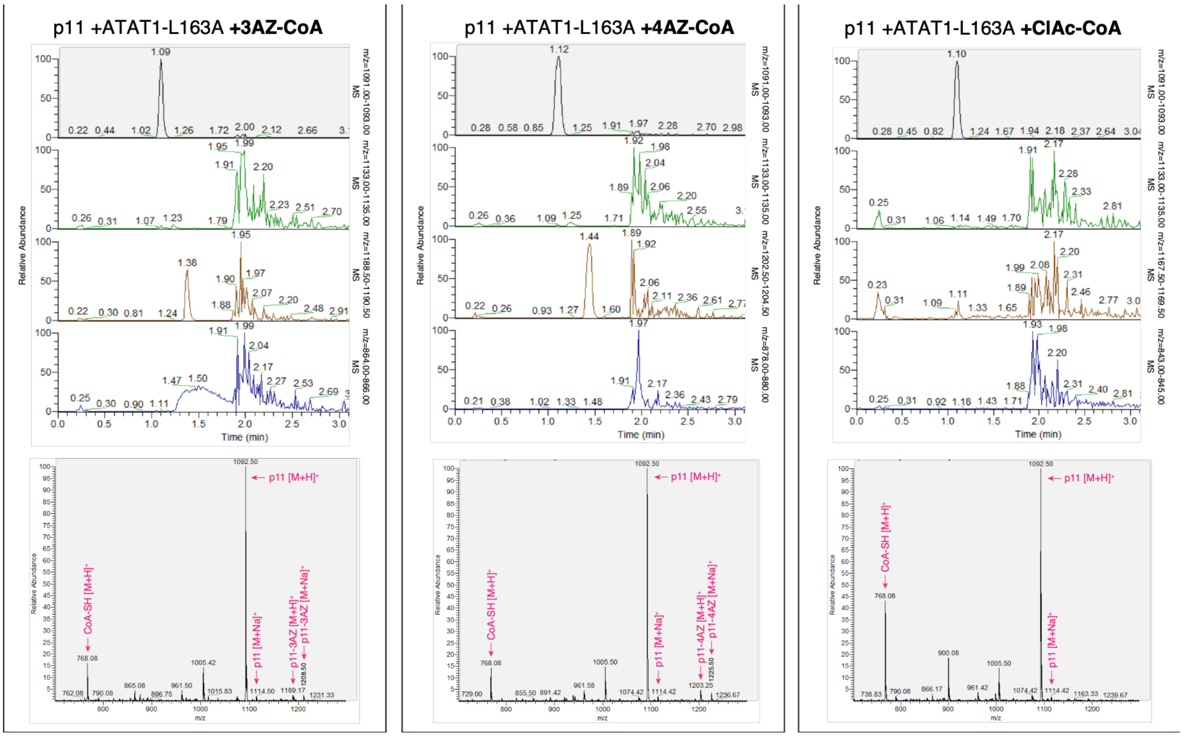

**Figure S16.** Extracted ion chromatograms (top) and extracted ESI–MS spectra (bottom) for the same ATAT1-L163A reactions shown in Figure S15, displaying the indicated m/z ranges and masses corresponding to the p11 substrate (black), p11-Ac (green), acylated p11 products (brown), and the corresponding cofactors (blue). Reactions were performed in the absence of externally added Ac-CoA or in the presence of Ac-, 3BY-, 4PY-, 5HY-, 6HY-, 3AZ-, 4AZ-, or ClAc-CoA, as indicated, and incubated for 96 h at 24 °C (375 $\mu$M p11, 1 mM cofactor, 75 $\mu$M WT-ATAT1). Theoretical m/z values are 1134.49 (p11-Ac), 1158.49 (p11-3BY), 1172.51 (p11-4PY), 1186.52 (p11-5HY), 1200.54 (p11-6HY), 1189.51 (p11-3AZ), 1203.52 (p11-4AZ), and 1168.45 (p11-ClAc). The mass peak around Rt = ~2 min corresponds to ATAT1.

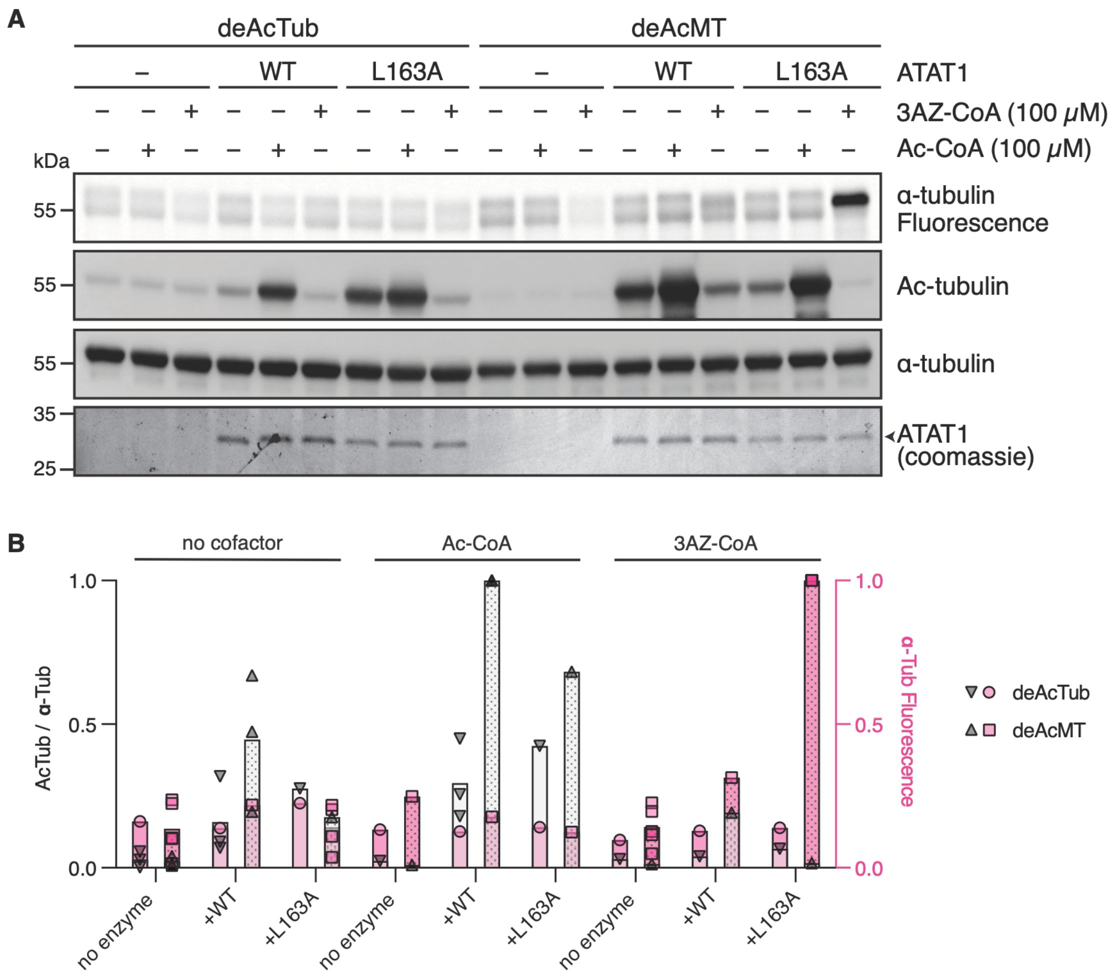

**Figure S17.** ATAT1-L163A modification of soluble tubulin or microtubules. **(A, B)** DeAcTub or deAcMT (10 μM) was incubated with or without Ac-CoA or 3AZ-CoA (100 μM) and 2 μM ATAT1 (WT or L163A) at room temperature for 1 h, followed by click chemistry labeling. Acylation was detected by in-gel fluorescence scanning, and acetylation by western blot. A representative gel/blot is shown in (A) and all data quantified in (B), presented as acetylated α-tubulin (grey) or α-tubulin fluorescence (pink) relative to total α-tubulin. Western blot band intensities were normalized to WT-ATAT1 + Ac-CoA and fluorescence intensities to ATAT1-L163A + 3AZ-CoA. Bars: mean, symbols: independent experiments (*N*). For WB, *N* = 3 for deAcTub, deAcTub + WT-ATAT1, deAcTub + WT-ATAT1 + Ac-CoA, deAcMT, deAcMT + WT-ATAT1, and deAcMT + WT-ATAT1 + Ac-CoA, and *N* = 1 for all other conditions. For fluorescence, *N* = 5 for deAcMT conditions and *N* = 1 for deAcTub conditions.

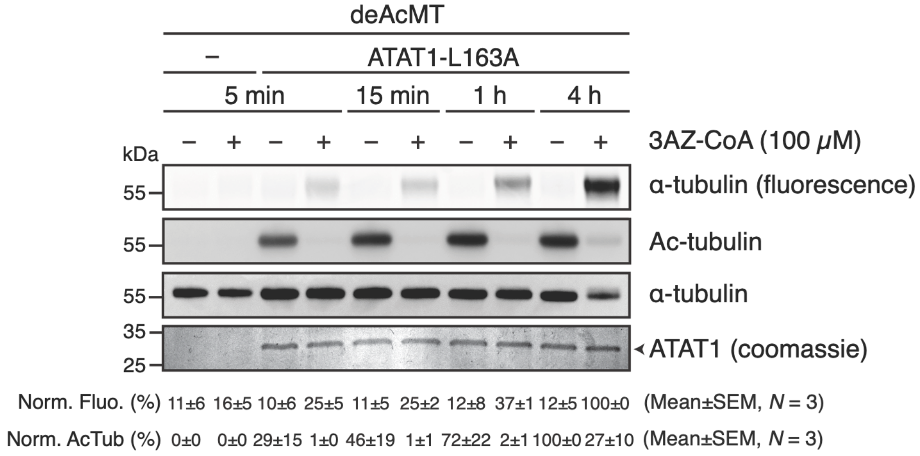

**Figure S18.** Time-dependent microtubule acylation by ATAT1-L163A with 3AZ-CoA in vitro. DeAcMT (10 $\mu$M) was incubated with or without 3AZ-CoA (100 $\mu$M) and 2 $\mu$M ATAT1-L163A at room temperature for the indicated times, quenched by boiling, and subjected to click chemistry labeling. Acylation was detected by in-gel fluorescence scanning and acetylation by western blot (WB). Representative gel/blot is shown and data from $N = 3$ independent experiments quantified. Quantification is presented as the percentage of α-tubulin fluorescence or acetylated α-tubulin relative to total α-tubulin, with fluorescence data normalized to the 4 h time point with 3AZ-CoA and WB data to the 4 h time point without 3AZ-CoA.

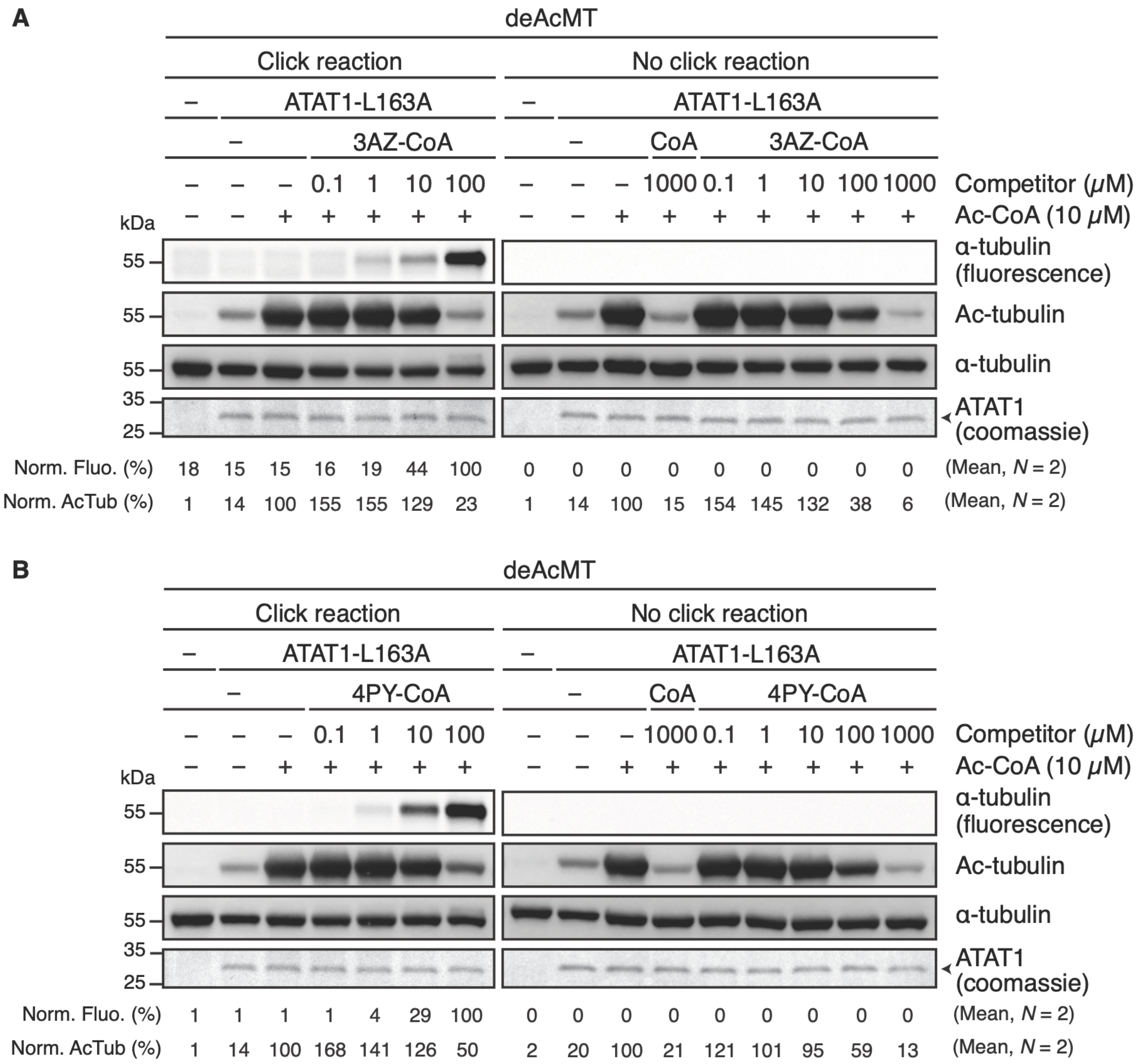

**Figure S19.** (A, B) Concentration-dependent competition and acylation assays with 3AZ-CoA (A) or 4PY-CoA (B), performed as described for Figure 5F. Acylation was detected by in-gel fluorescence scanning after click chemistry labeling. Parallel reactions analyzed by WB without click chemistry labeling assessed recognition of 4PY- or 3AZ-acylated tubulin by the anti-acetylated α-tubulin antibody. Quantification is presented as the percentage of α-tubulin fluorescence or acetylated α-tubulin relative to total α-tubulin, with fluorescence data normalized to ATAT1-L163A + 4PY- or 3AZ-CoA (100 μM) and WB data to ATAT1-L163A + Ac-CoA without 4PY- or 3AZ-CoA. The click reaction data in (B) are reproduced from Figure 5F for visualization purposes.

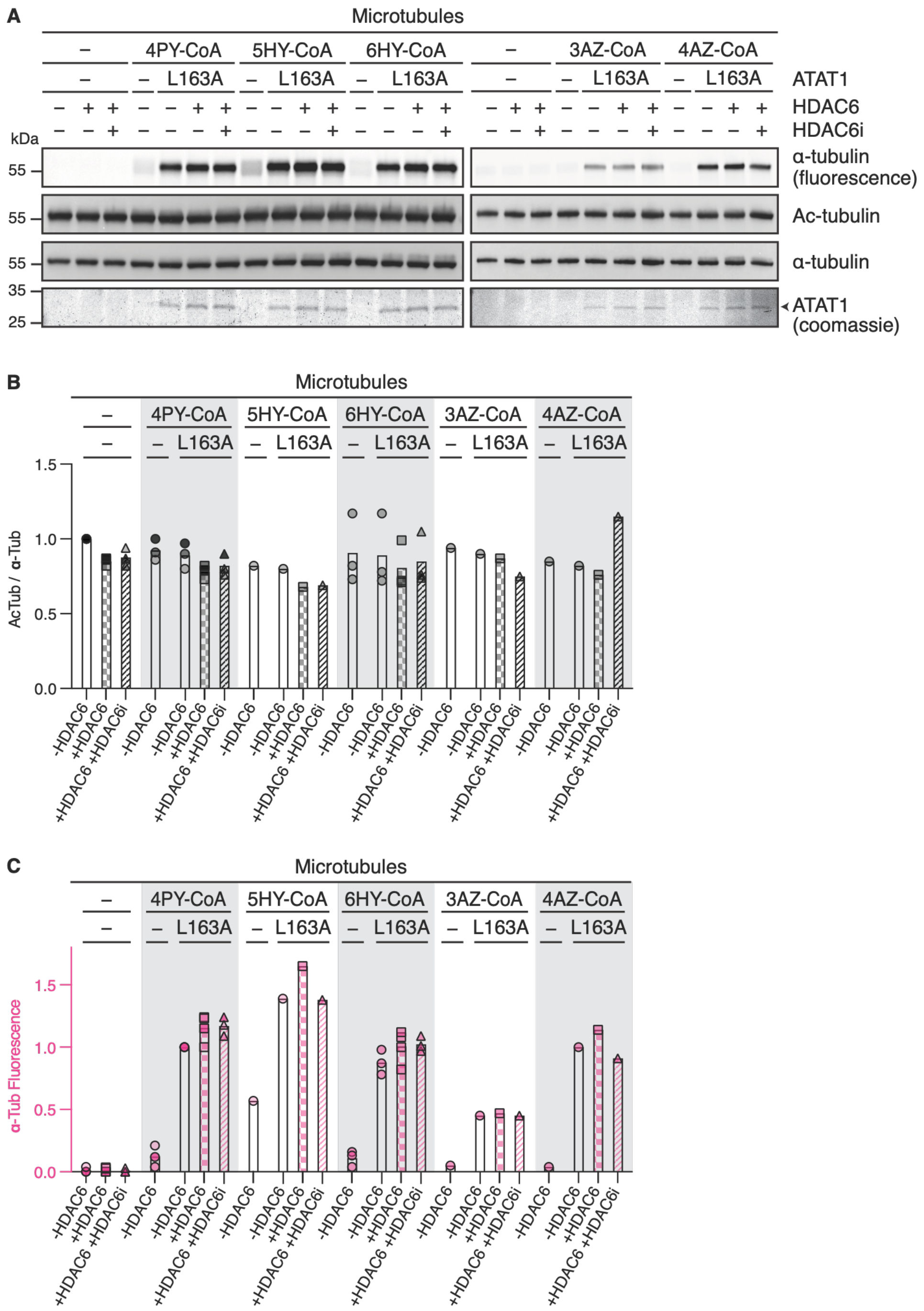

**Figure S20.** Analysis of HDAC6-mediated removal of Ac-CoA mimic-acylation on microtubules. (A–C) Microtubules were incubated with 100 $\mu$M Ac-CoA mimic and 2 $\mu$M ATAT1-L163A for 1 h at room temperature, followed by treatment with 1 $\mu$M HDAC6, 1 $\mu$M HDAC6 in the presence of 100 $\mu$M Tubacin (HDAC6i), or no HDAC6 for 1.5 h. Acylated tubulin was detected after click chemistry labeling by in-gel fluorescence scanning, and acetylated tubulin by WB. A representative blot/gel is shown in (A) and all data quantified in (B, C), presented as acetylated α-tubulin (B) or α-tubulin fluorescence (C) relative to total α-tubulin. WB data were normalized to microtubules without ATAT1, Ac-CoA mimics, or HDAC6, and fluorescence data to microtubules with ATAT1-L163A + 4PY-CoA without HDAC6. Bar represents the mean and symbols represent $N$ independent experiments. $N$ = 1 for conditions with 5HY-, 3AZ-, or 4AZ-CoA, and $N$ = 3 for all other conditions.

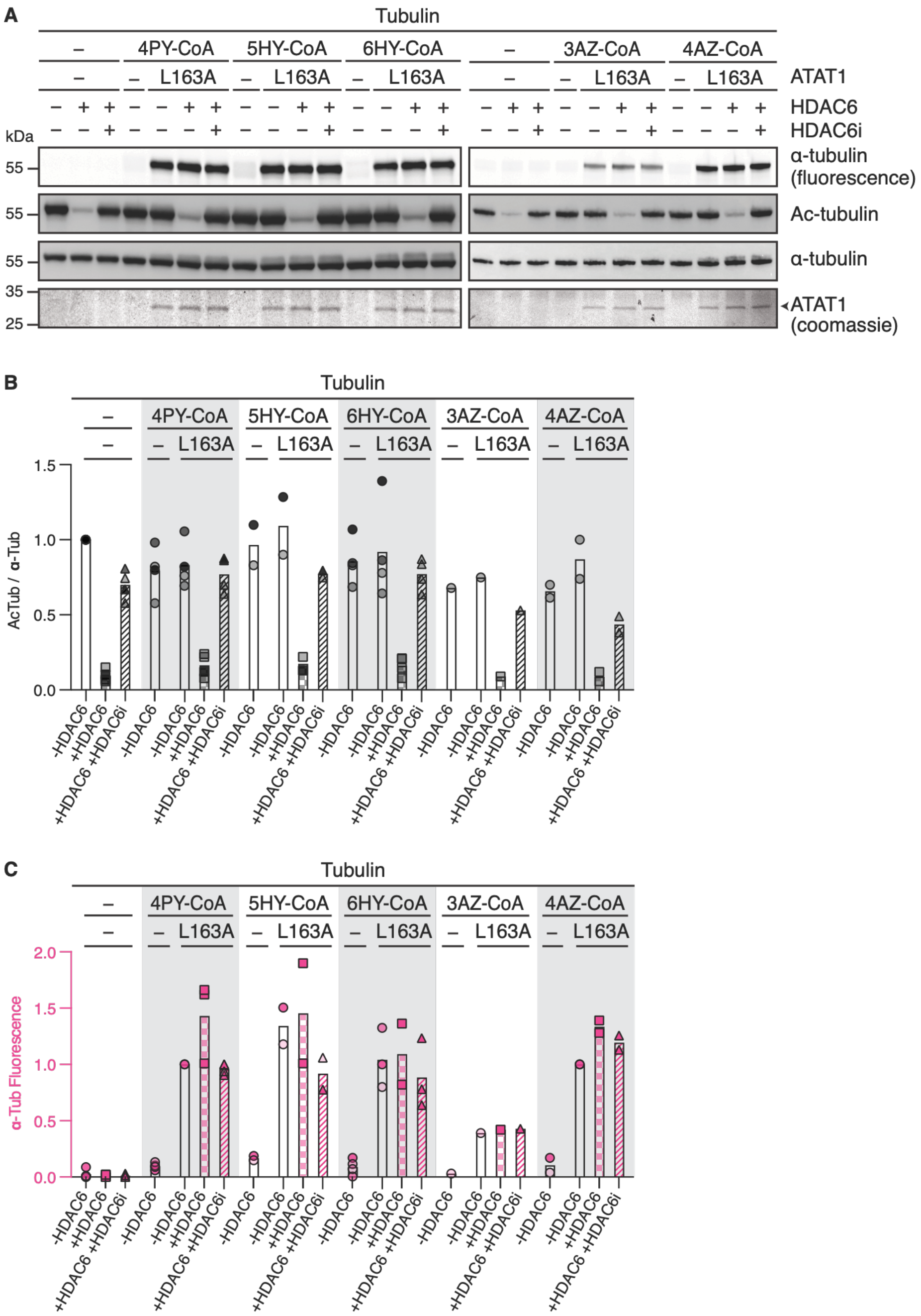

**Figure S21.** Analysis of HDAC6-mediated removal of Ac-CoA mimic-acylation on soluble tubulin. (A–C) Soluble tubulin was incubated with 100 $\mu$M Ac-CoA mimic and 2 $\mu$M ATAT1-L163A for 1 h at room temperature, followed by treatment with 1 $\mu$M HDAC6, 1 $\mu$M HDAC6 in the presence of 100 $\mu$M Tubacin (HDAC6i), or no HDAC6 for 1.5 h. Acylated tubulin was detected after click chemistry labeling by in-gel fluorescence scanning, and acetylated tubulin by WB. A representative blot/gel is shown in (A) and all data quantified in (B, C), presented as acetylated α-tubulin (B) or α-tubulin fluorescence (C) relative to total α-tubulin. WB data were normalized to tubulin without ATAT1, Ac-CoA mimics, or HDAC6, and fluorescence data to tubulin with ATAT1-L163A + 4PY-CoA without HDAC6. Bar represents the mean and symbols represent $N$ independent experiments. $N$ = 1 for conditions with 3AZ-CoA, $N$ = 2 for conditions with 5HY- or 4AZ-CoA, and $N$ = 4 for all other conditions.

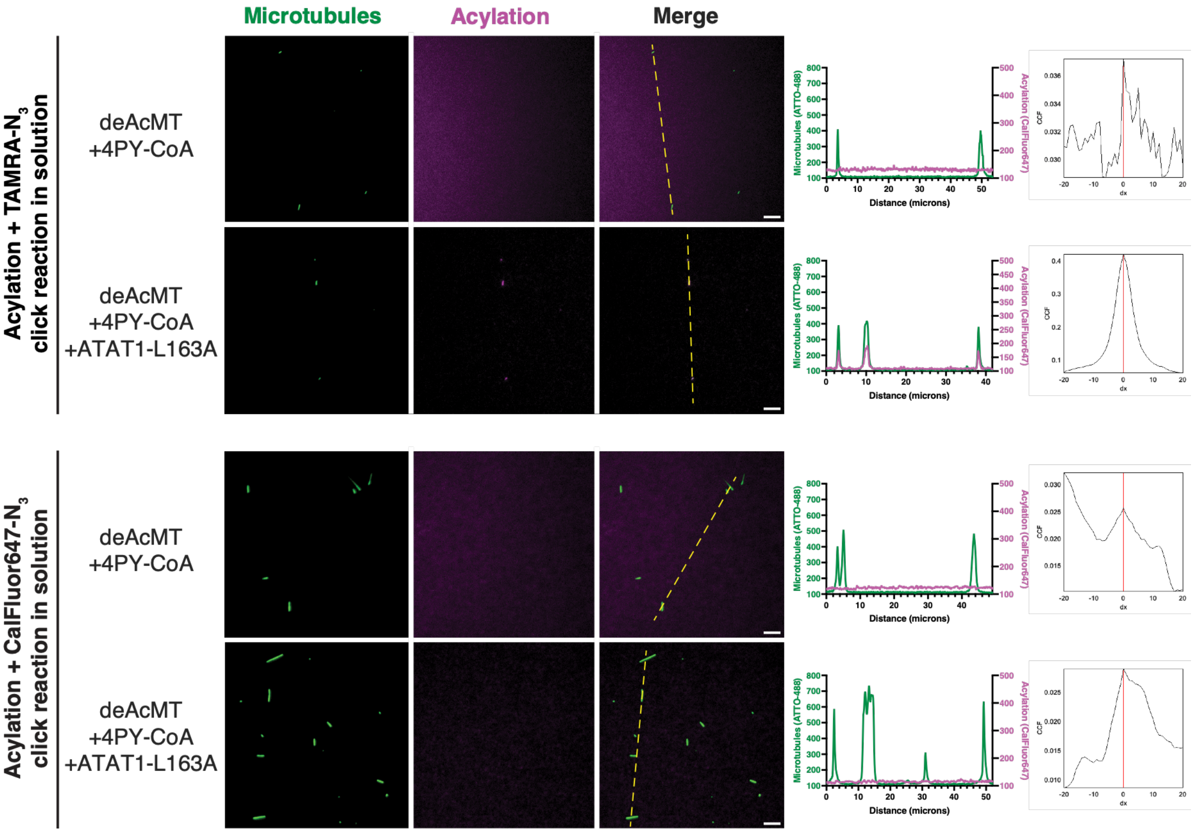

**Figure S22.** Representative TIRF images of microtubules following solution-phase acylation and click chemistry before immobilization. DeAcMT (5 $\mu$M) were acylated with 4PY-CoA (100 $\mu$M) in the presence or absence of ATAT1-L163A (1 $\mu$M), followed by in-solution click chemistry with TAMRA-$N_3$ or CalFluor647-$N_3$, as indicated. Microtubules were immobilized prior to TIRF microscopy analysis. Line profiles of the CalFluor647 fluorescence signal (640 nm channel) and corresponding ATTO-488-labeled microtubules (488 nm channel) were extracted along the regions indicated by the dashed yellow line in the merged images. Van Steensel cross-correlation function (CCF) analysis is shown alongside the line profiles. Scale bars, 5 µm. $N$ = 6 independent experiments for TAMRA-$N_3$ and $N$ = 2 independent experiments for CalFluor647-$N_3$.

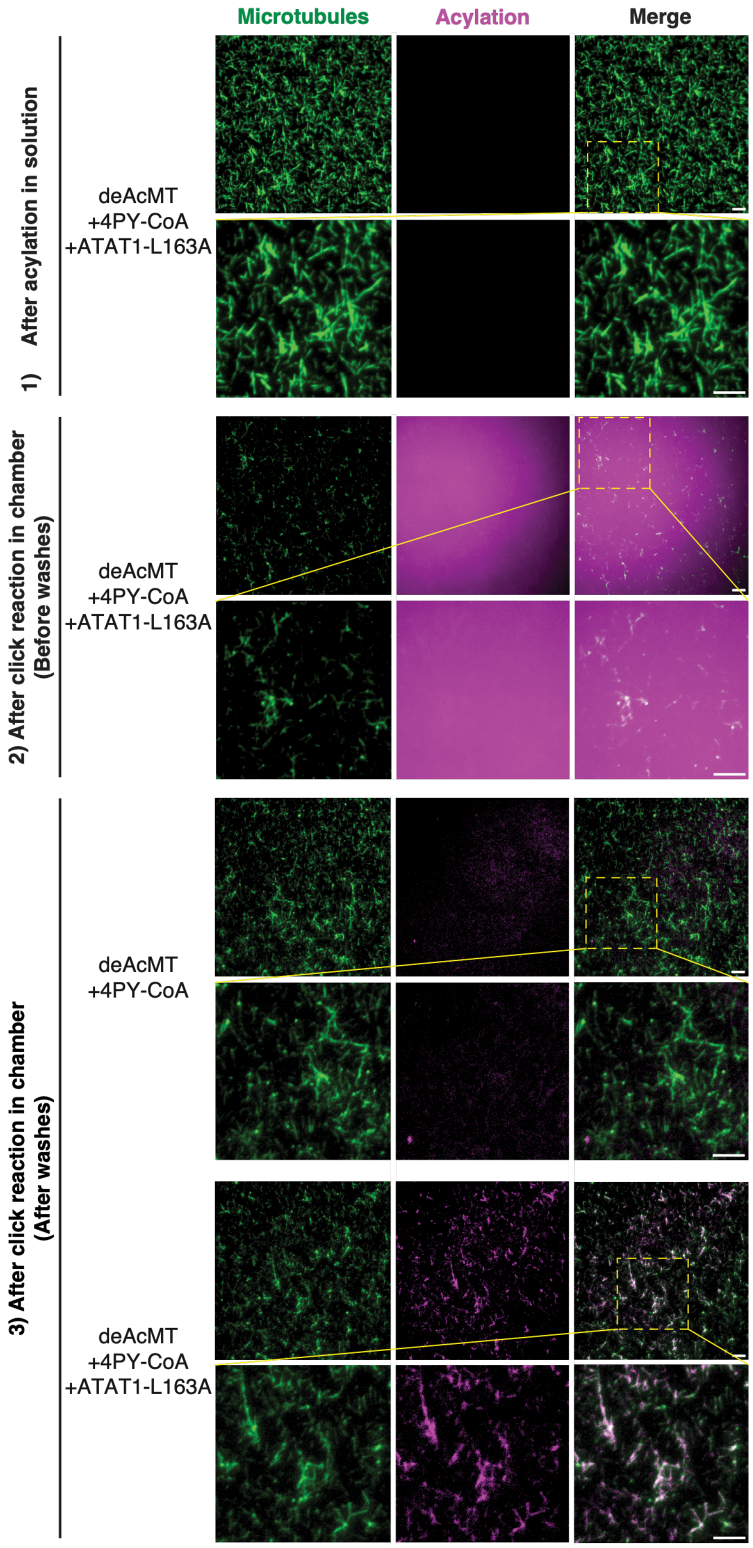

**Figure S23.** Representative TIRF images of immobilized microtubules following acylation in solution (5 $\mu$M deAcMT, 100 $\mu$M 4PY-CoA, 1 $\mu$M ATAT1-L163A) and in-chamber click chemistry with CalFluor647-$N_3$, imaged before or after washing, as indicated. Zoomed-out views of the representative images shown in Figure 6A are shown, with the indicated regions of interest (ROIs) reproduced as insets. Images show ATTO-488-labeled microtubules and CalFluor647-labeled acylation signal. Scale bars, 5 µm. *N* = 4 independent experiments.

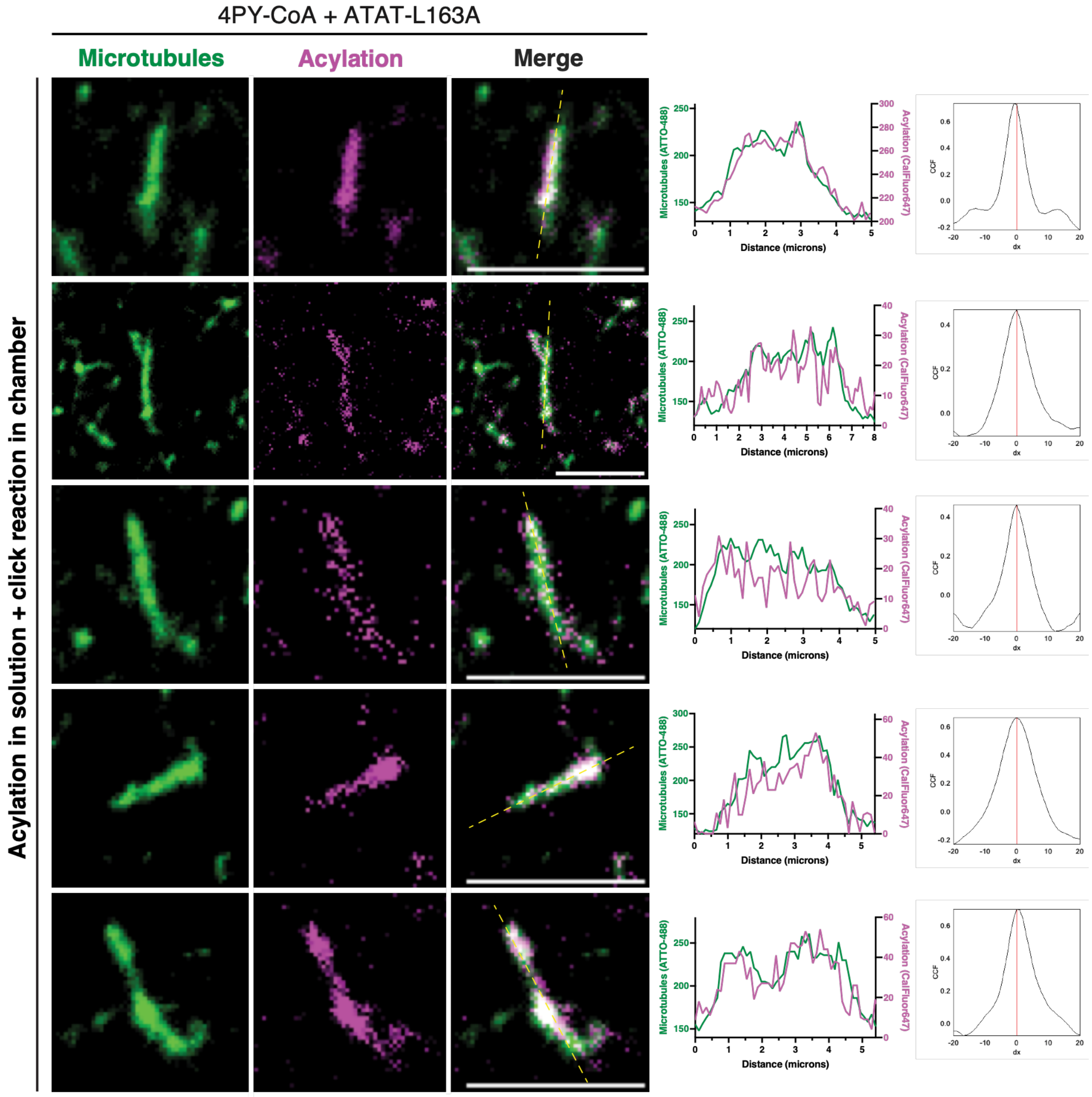

4PY-CoA + ATAT-L163A
Microtubules
Acylation
Merge
Acylation in solution + click reaction in chamber
Microtubules (ATTO-488)
Acylation (CalFluor647)
Distance (microns)
CCF
dx

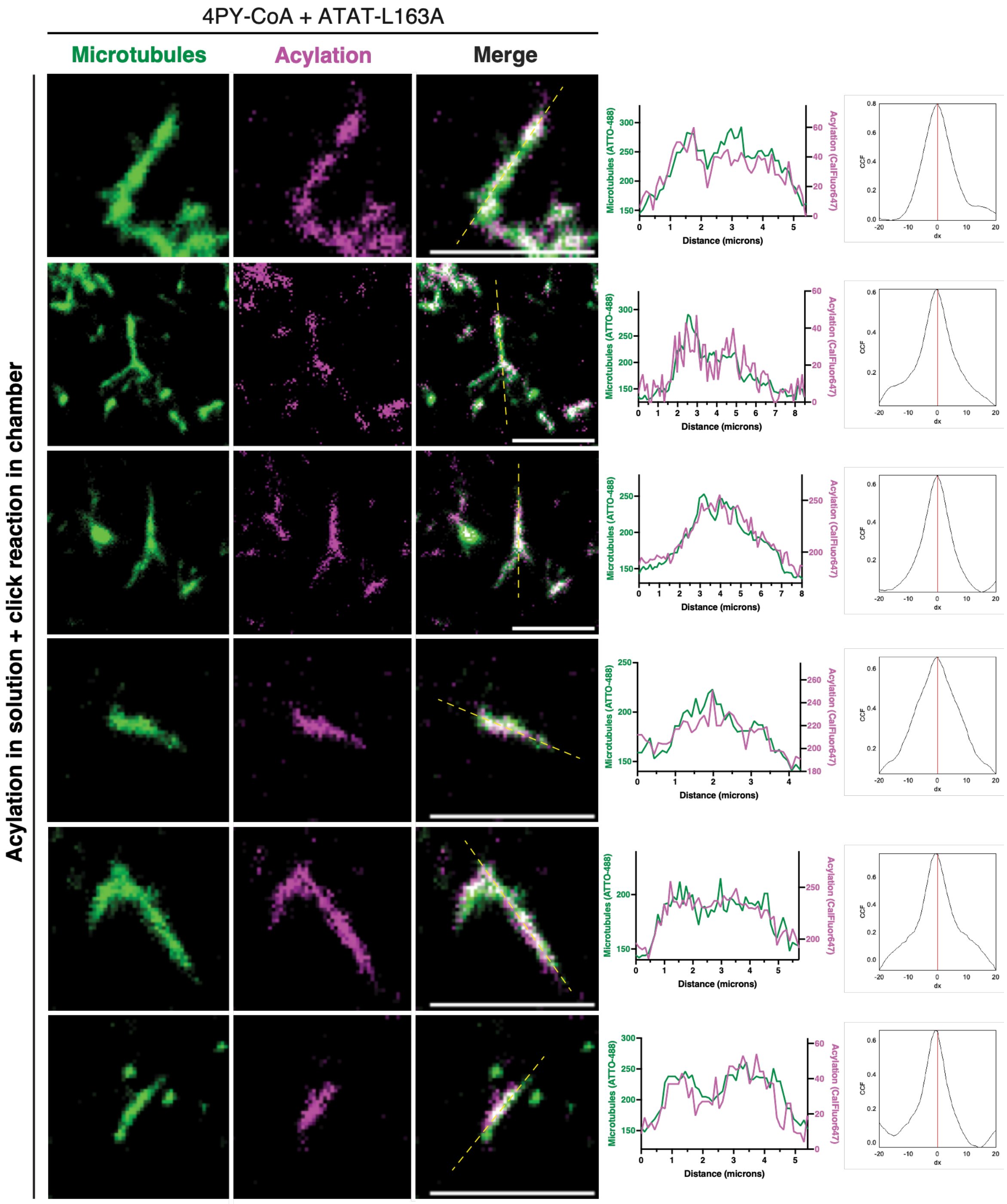

4PY-CoA + ATAT-L163A
Microtubules
Acylation
Merge
Acylation in solution + click reaction in chamber
Microtubules (ATTO-488)
Acylation (CalFluor647)
Distance (microns)
CCF
dx

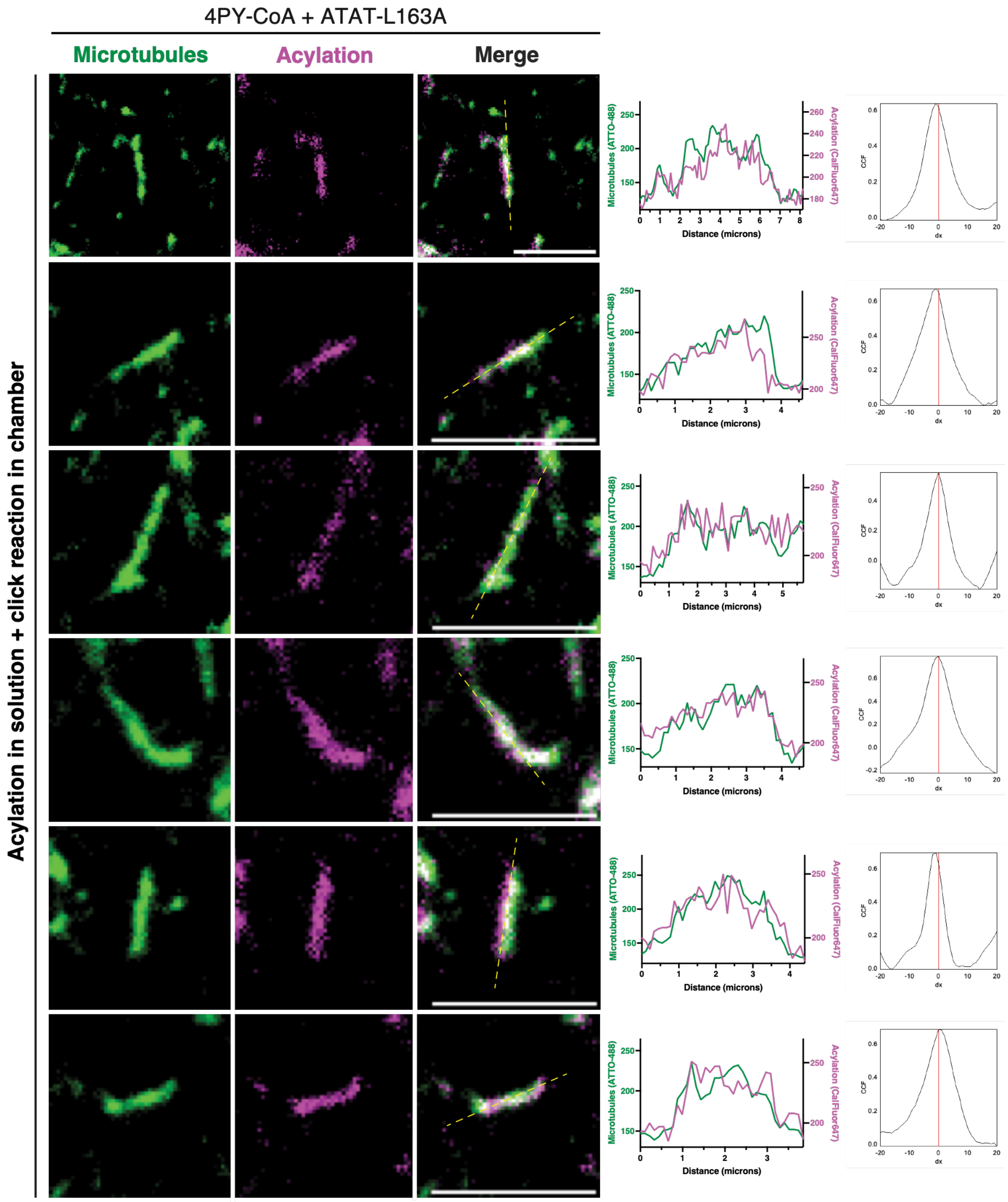

4PY-CoA + ATAT-L163A
Microtubules
Acylation
Merge
Acylation in solution + click reaction in chamber
Microtubules (ATTO-488)
Acylation (CalFluor647)
Distance (microns)
CCF
dx

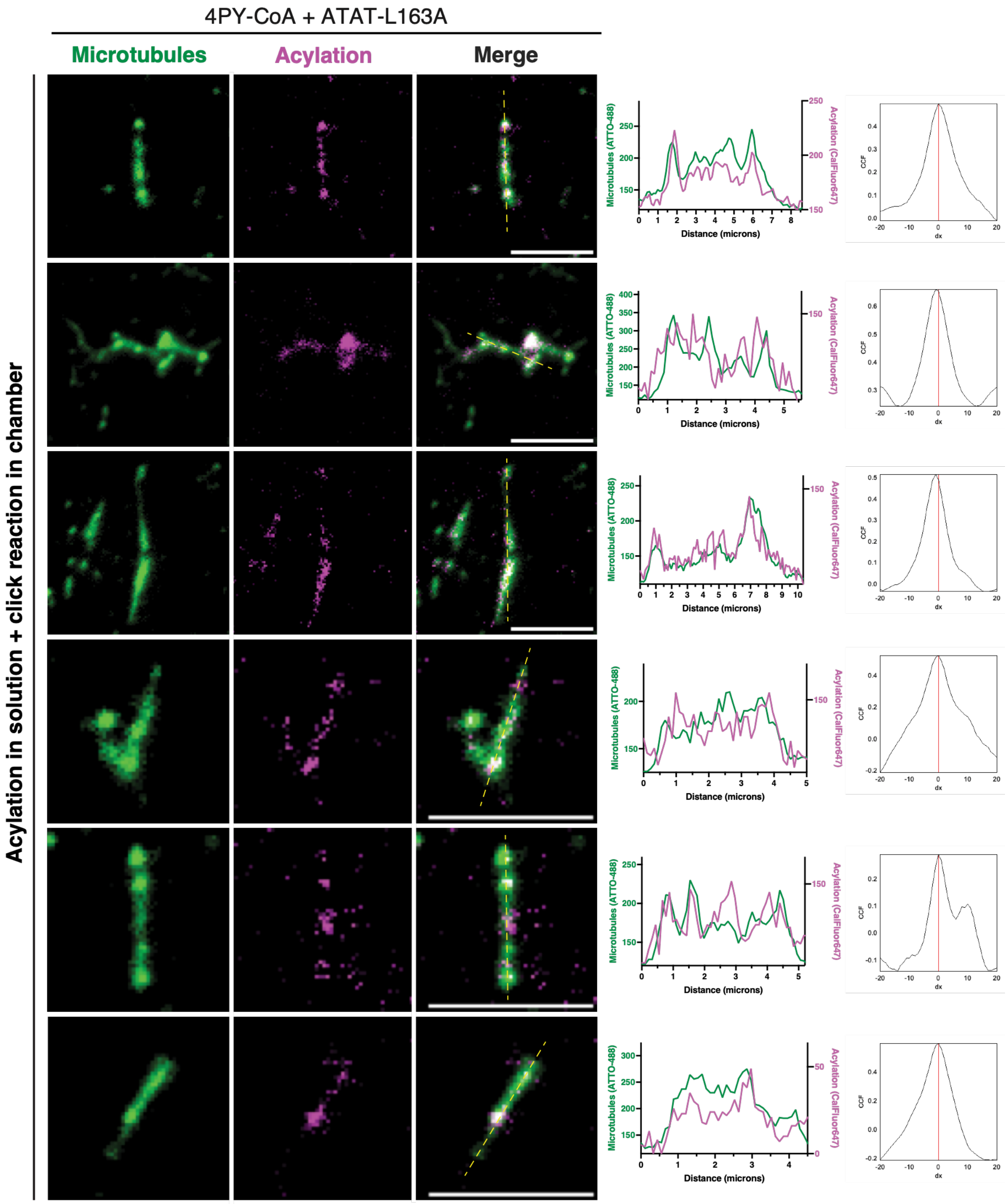
4PY-CoA + ATAT-L163A
Microtubules
Acylation
Merge
Acylation in solution + click reaction in chamber
Microtubules (ATTO-488)
Acylation (CalFluor647)
Distance (microns)
CCF
dx

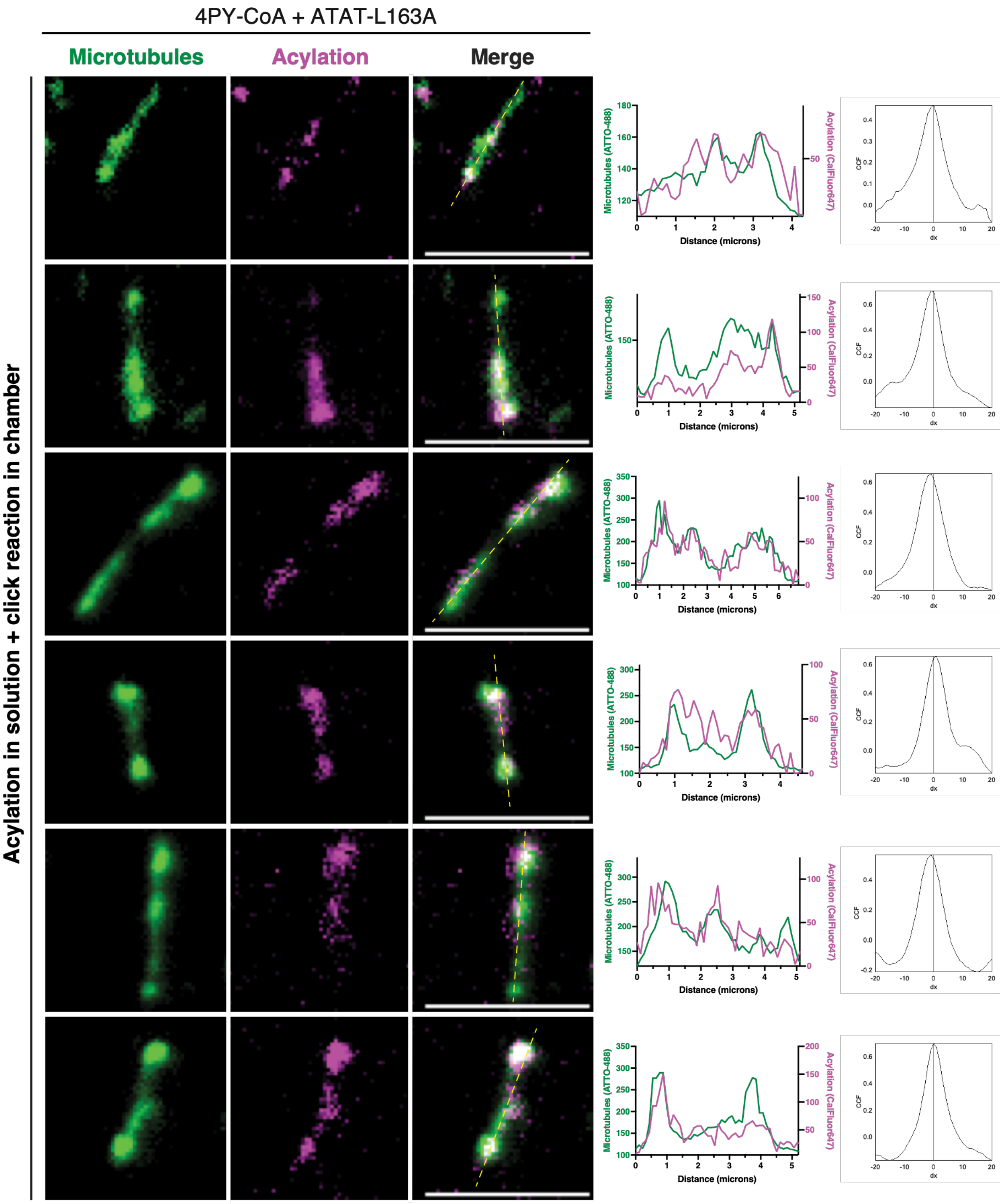

**Figure S24.** Representative individual microtubules from *N* = 4 independent experiments following solution-phase acylation by ATAT1-L163A, immobilization, and in-chamber click chemistry with CalFluor647-$N_3$, as described in Figure S23. Images show ATTO-488-labeled microtubules and CalFluor647-labeled acylation signal. Line profiles and Van Steensel CCF analysis are shown as described in Figure S22. Complementary colocalization metrics are provided in Table S6. Scale bars, 5 µm.

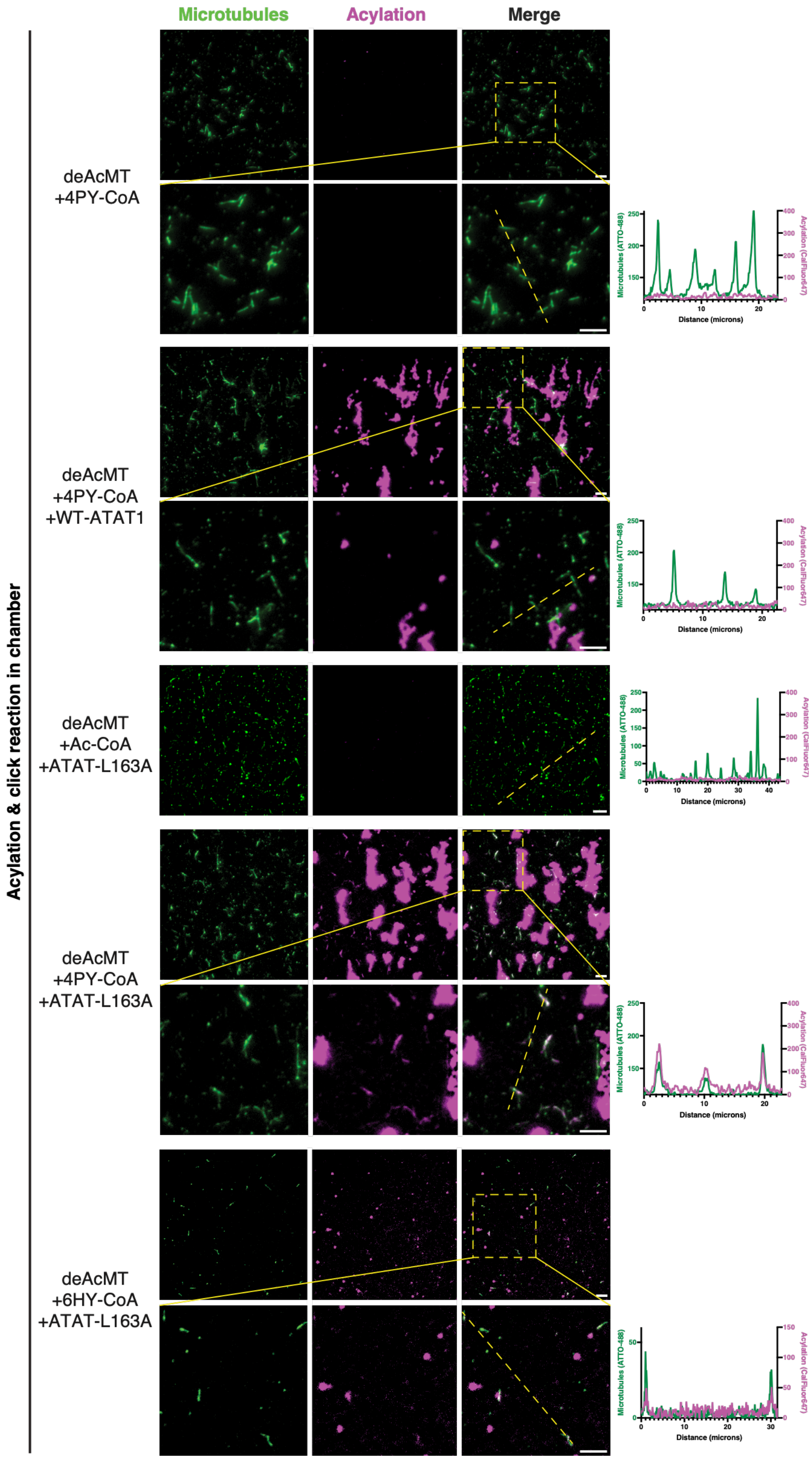

**Figure S25.** Representative TIRF images of immobilized microtubules following in-chamber acylation and click chemistry with CalFluor647-$N_3$ under the indicated conditions (500 $\mu$M cofactor and 5 $\mu$M ATAT1). Zoomed-out views are shown, with the indicated ROIs shown as insets. Insets for deAcMT + 4PY-CoA + WT-ATAT1 and deAcMT + 4PY-CoA + ATAT1-L163A are reproduced from Figure 6B for visualization purposes. Images show ATTO-488-labeled microtubules and CalFluor647-labeled acylation signal. Line profiles were extracted as described in Figure S22. Scale bars, 5 µm. The number of independent experiments were as follows: $N = 1$ for deAcMT + 6HY-CoA + ATAT1-L163A, $N = 2$ for deAcMT + Ac-CoA + ATAT1-L163A, and $N = 7$ for all other conditions.

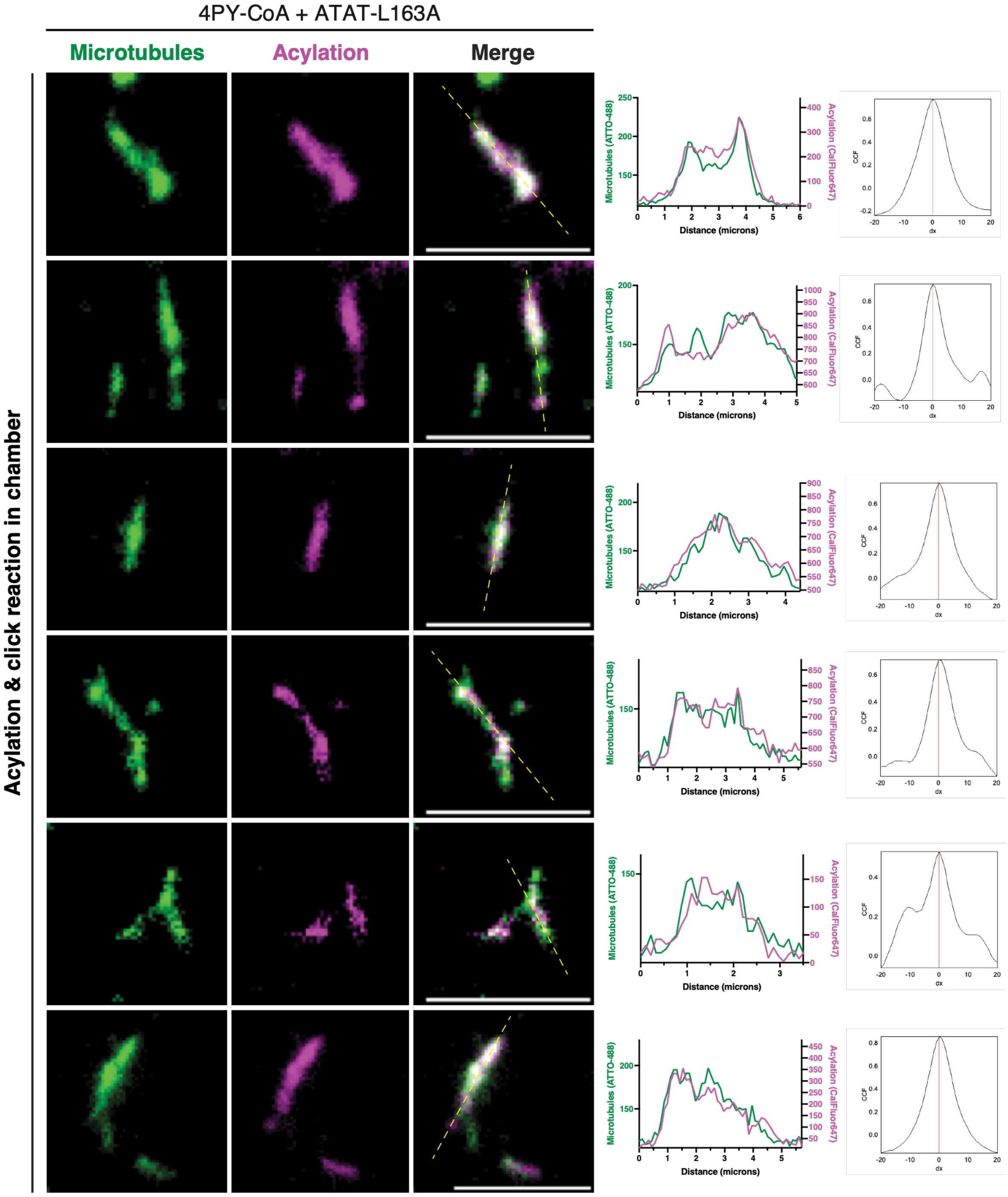

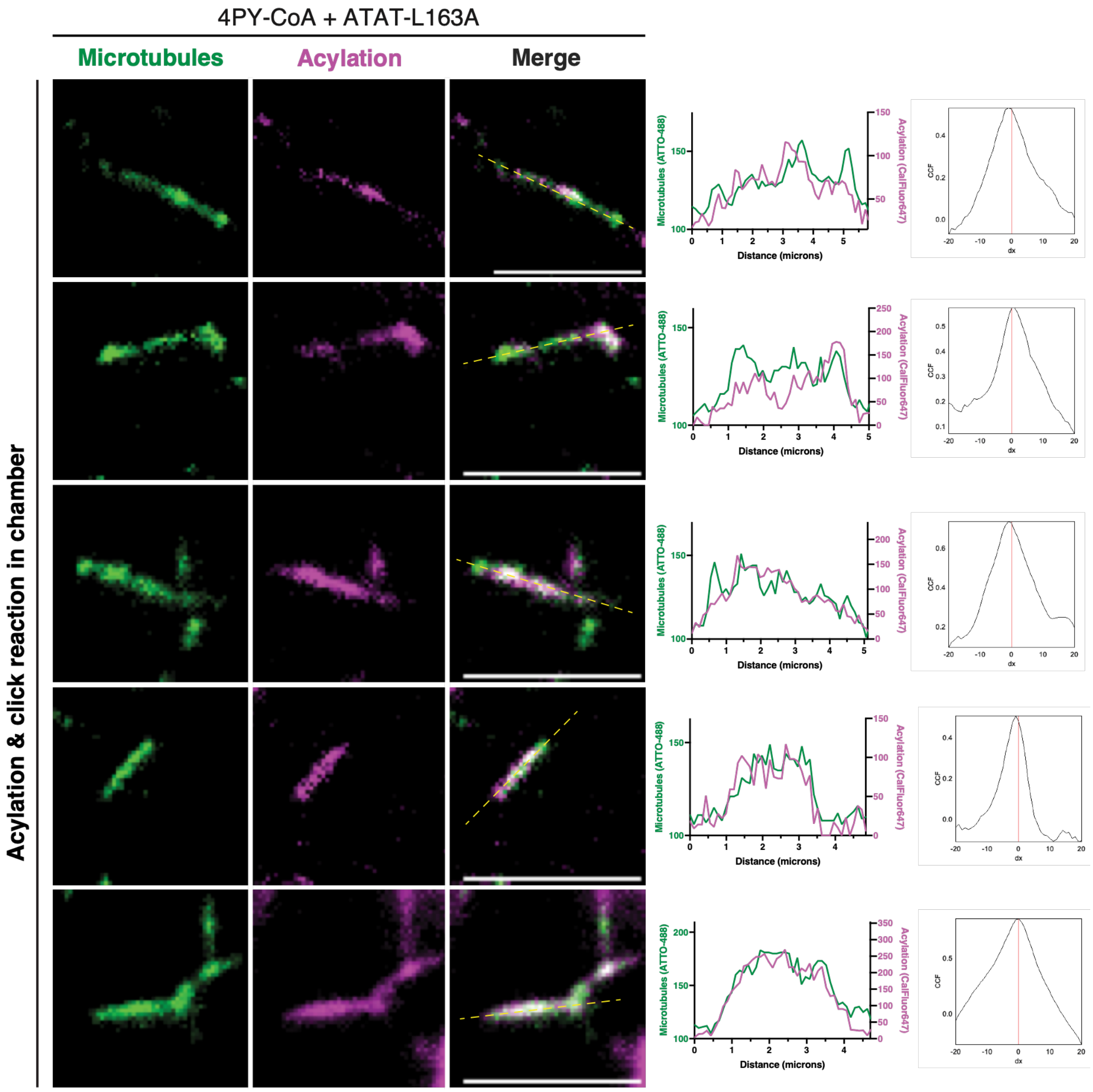

4PY-CoA + ATAT-L163A
Microtubules
Acylation
Merge
Acylation & click reaction in chamber
Microtubules (ATTO-488)
Acylation (CalFluor647)
Distance (microns)
CCF
dx

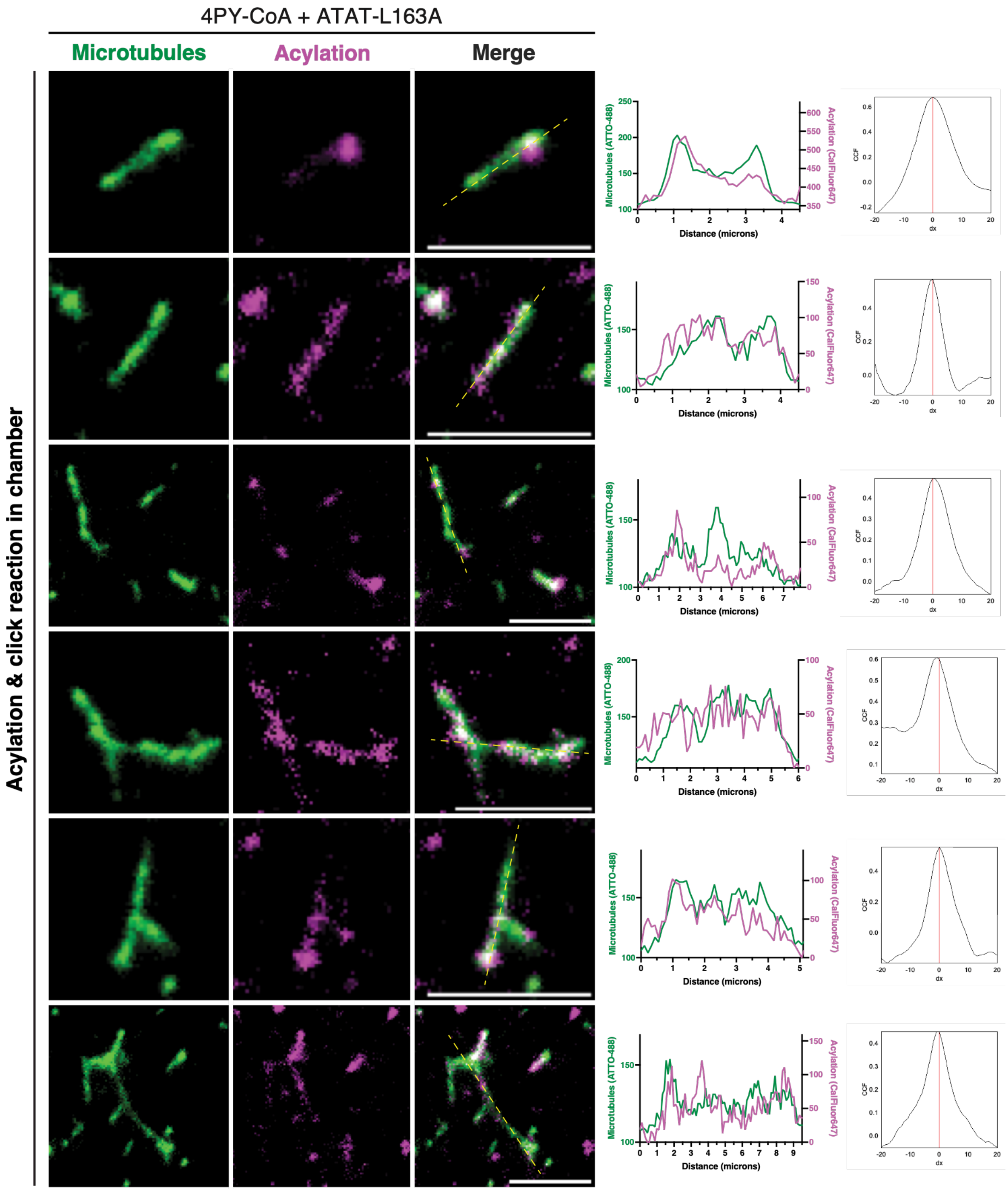
4PY-CoA + ATAT-L163A
Microtubules
Acylation
Merge
Acylation & click reaction in chamber
Microtubules (ATTO-488)
Acylation (CalFluor647)
Distance (microns)
CCF
dx

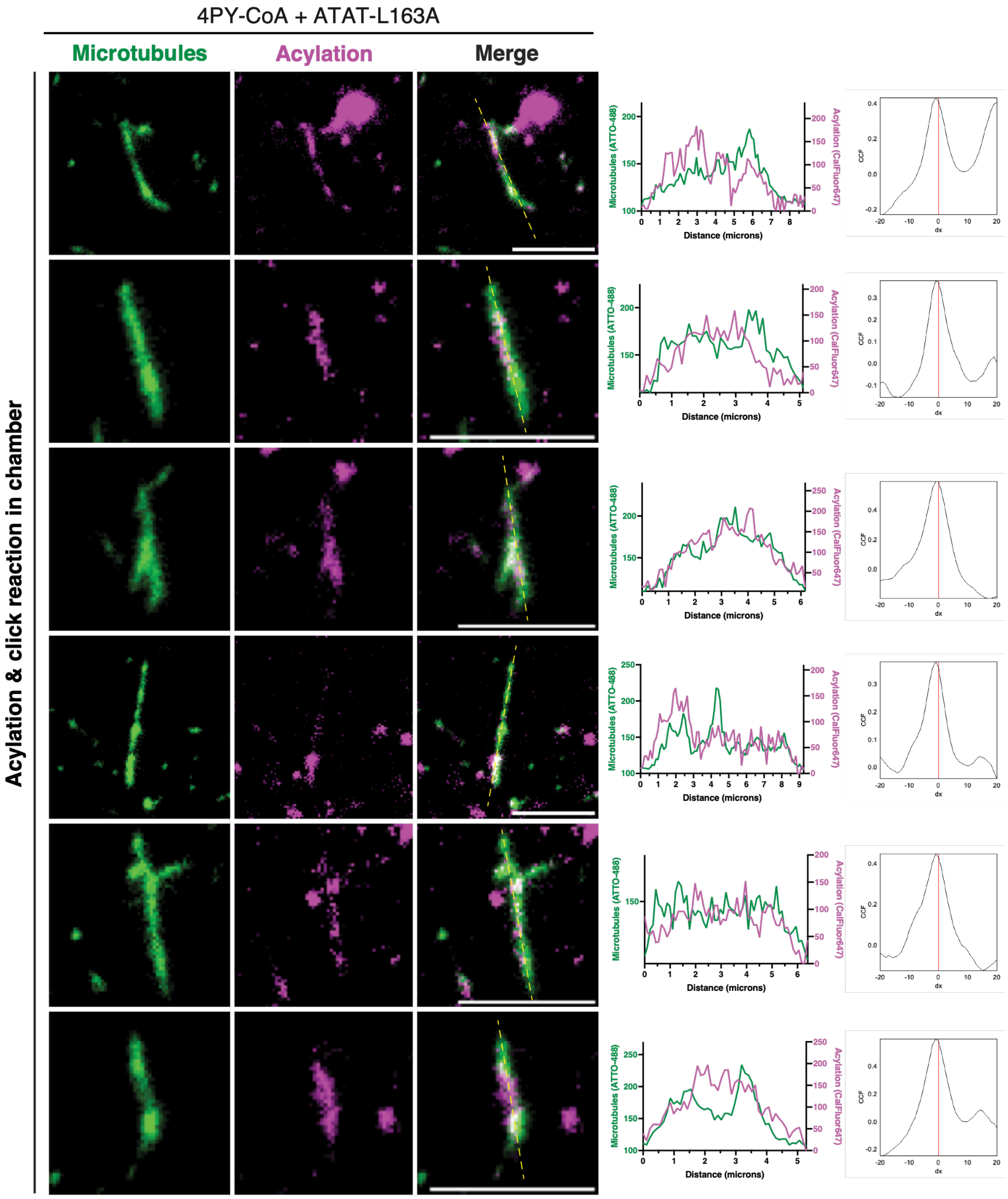

4PY-CoA + ATAT-L163A
Microtubules
Acylation
Merge
Acylation & click reaction in chamber
Microtubules (ATTO-488)
Acylation (CalFluor647)
Distance (microns)
CCF
dx

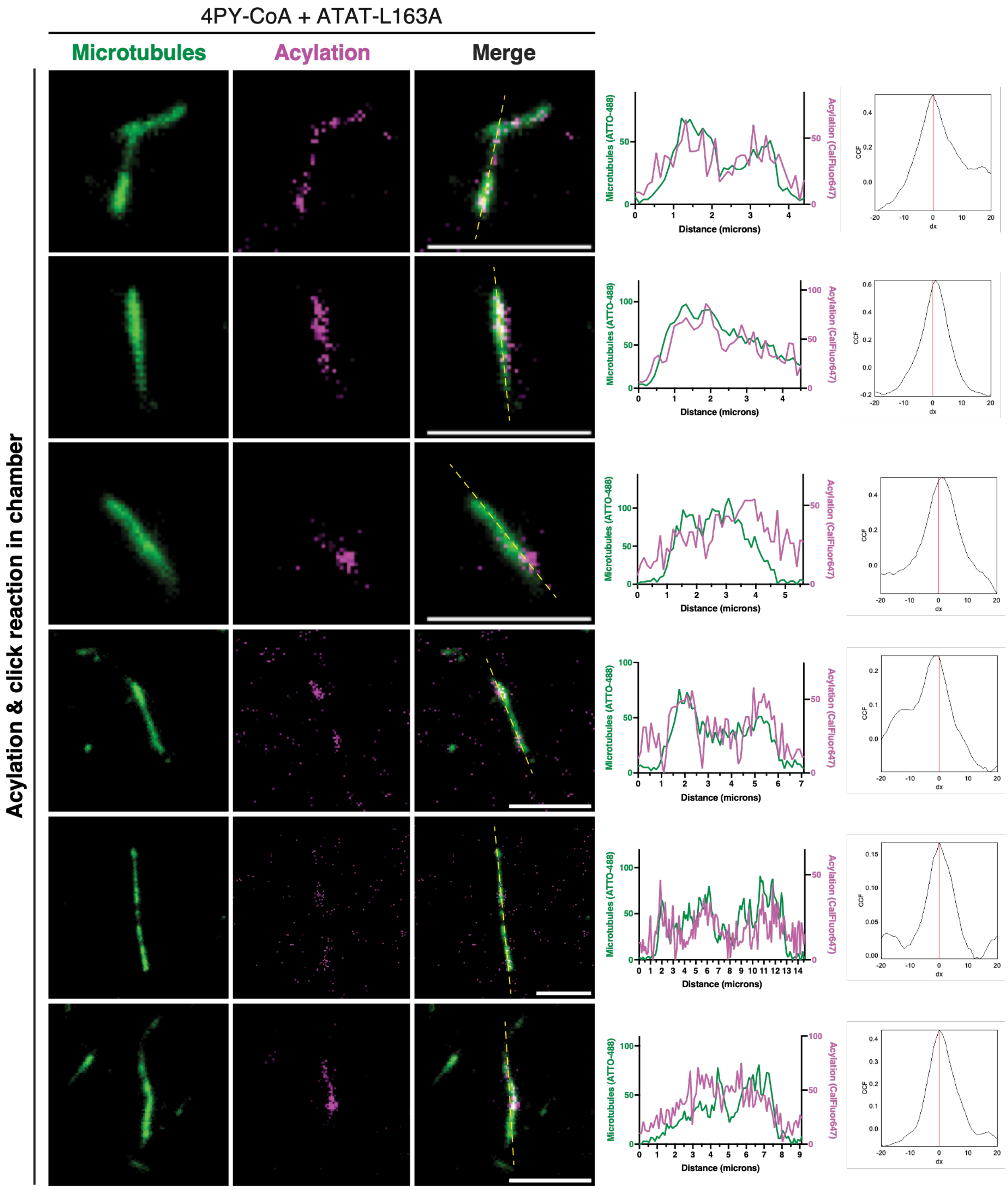

**Figure S26.** Representative individual microtubules from $N = 7$ independent experiments following in-chamber acylation by 500 $\mu$M 4PY-CoA + 5 $\mu$M ATAT1-L163A and click chemistry with CalFluor647-$N_3$. Images show ATTO-488-labeled microtubules and CalFluor647-labeled acylation signal. Line profiles and Van Steensel CCF analysis are shown as described in Figure S22. Complementary colocalization metrics are provided in Table S6. Scale bars, 5 µm.

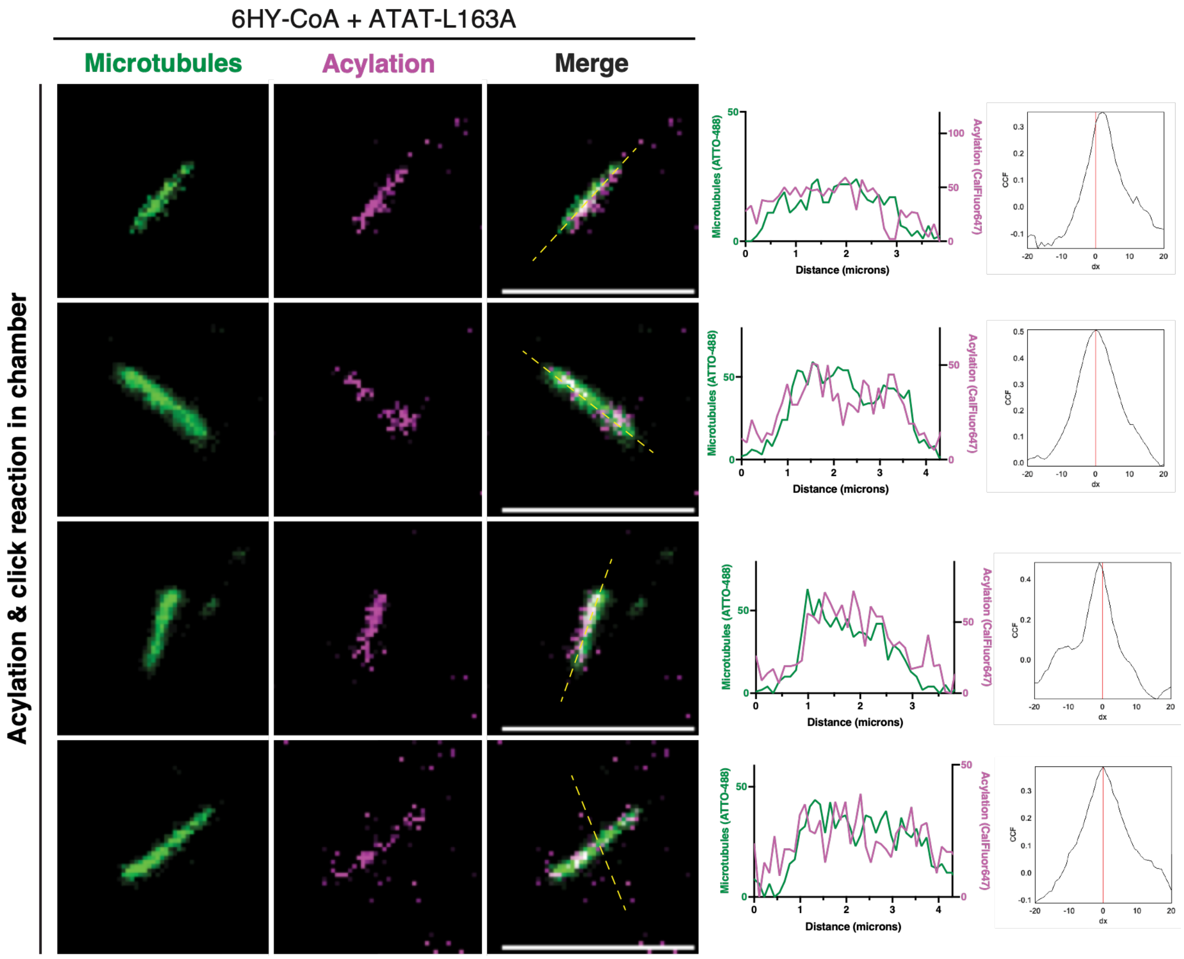

**Figure S27.** Representative individual microtubules from $N = 1$ independent experiment following in-chamber acylation by 500 $\mu$M 6HY-CoA + 5 $\mu$M ATAT1-L163A and click chemistry with CalFluor647-$N_3$. Images show ATTO-488-labeled microtubules and CalFluor647-labeled acylation signal. Line profiles and Van Steensel CCF analysis are shown as described in Figure S22. Complementary colocalization metrics are provided in Table S6. Scale bars, 5 μm.

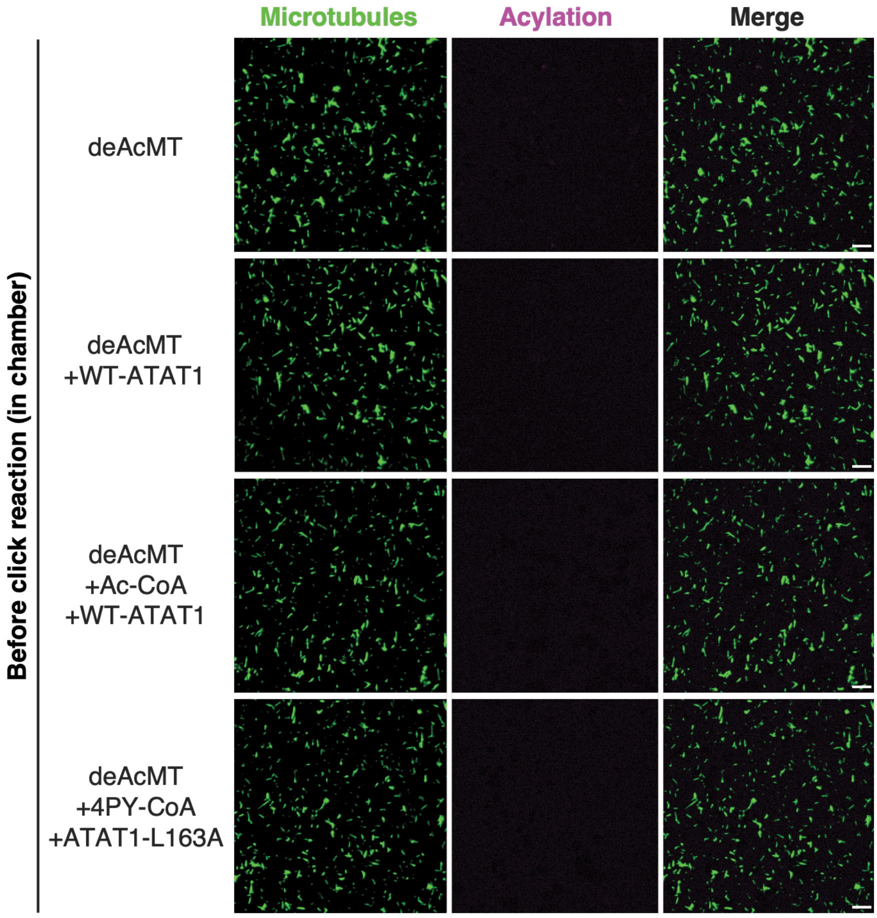

**Figure S28.** Representative TIRF images of immobilized microtubules before in-chamber click chemistry labeling with CalFluor647-$N_3$. Microtubules were generated from deAcTub (10 $\mu$M) modified with Ac-CoA or 4PY-CoA (100 $\mu$M) in the presence of WT-ATAT1 or ATAT1-L163A (2 $\mu$M), as indicated, and subsequently polymerized into stable microtubules. Images show ATTO-488-labeled microtubules. Scale bars, 5 µm. $N$ = 2 independent experiments.

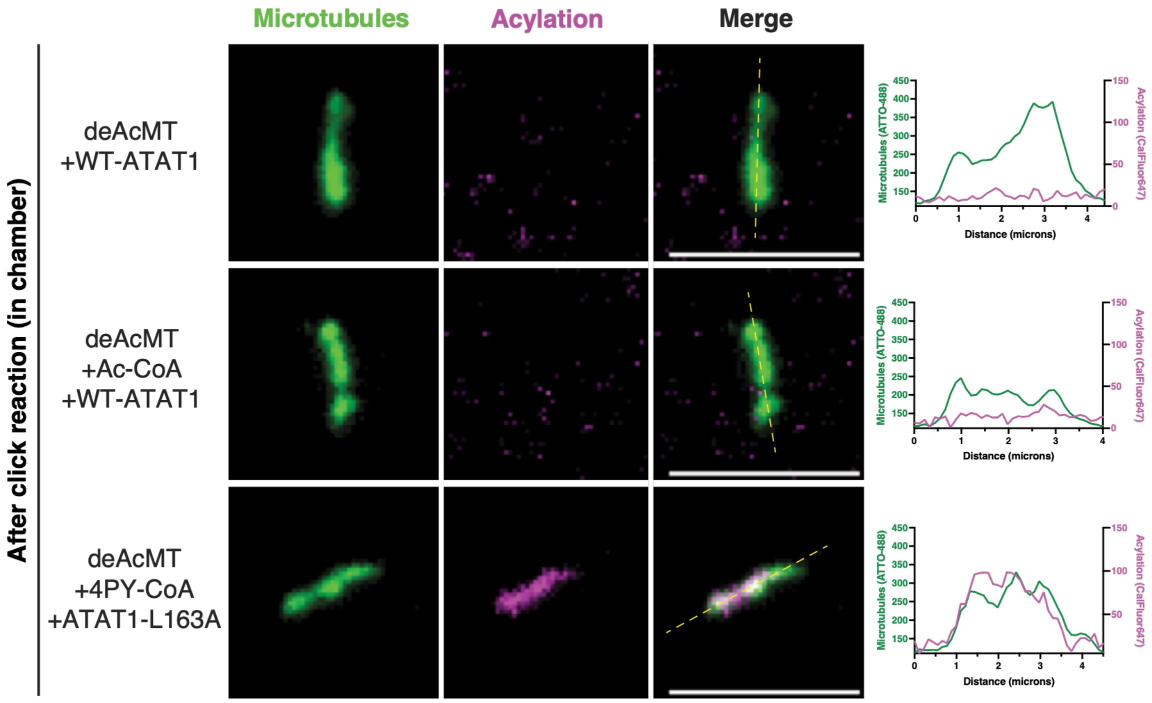

**Figure S29.** Representative TIRF images of the same conditions shown in Figure S28 following in-chamber click chemistry labeling with CalFluor647-$N_3$. Images show ATTO-488-labeled microtubules and CalFluor647-labeled acylation signal. Line profiles were extracted as described in Figure S22. Scale bars, 5 µm.

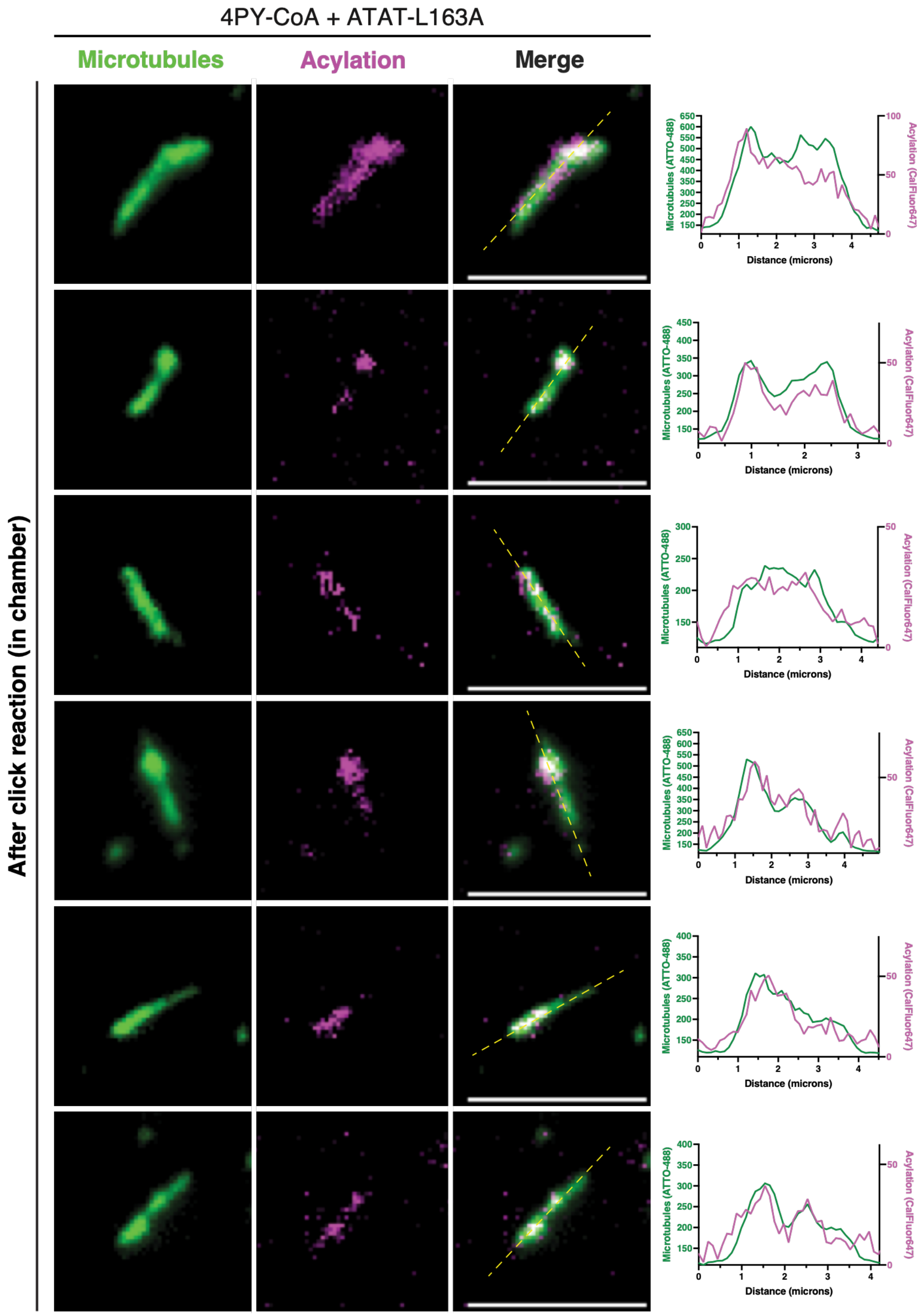
4PY-CoA + ATAT-L163A
Microtubules
Acylation
Merge
After click reaction (in chamber)
Microtubules (ATTO-488)
Acylation (CalFluor647)
Distance (microns)

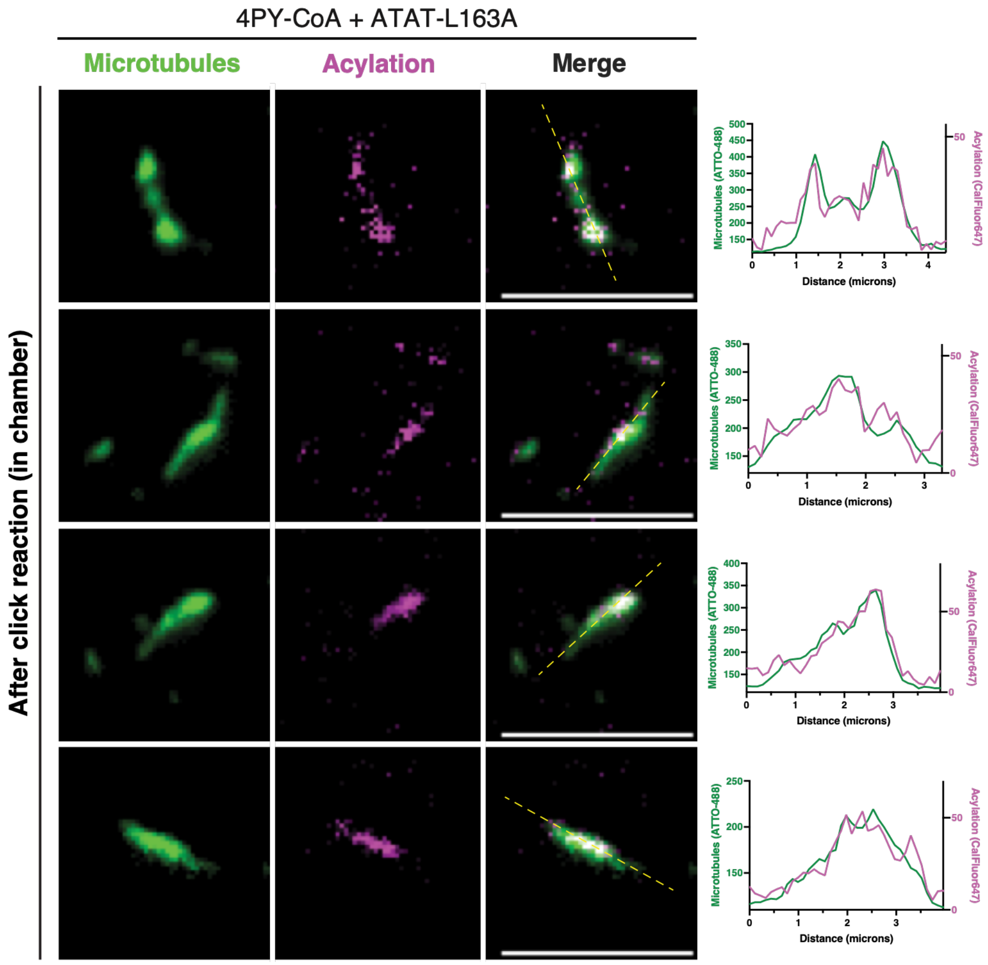

**Figure S30.** Representative TIRF images and line-profile analysis of individual microtubules from the deAcMT + 100 μM 4PY-CoA + 2 μM ATAT1-L163A condition shown in Figure S29 following in-chamber click chemistry labeling with CalFluor647-$N_3$. Scale bars, 5 μm.

#### SUPPLEMENTARY TABLES

**Table S1.** Sequences of forward and reverse PCR primers used for site-directed mutagenesis of *Hs*ATAT1.

| ATAT1 mutation | Primer | DNA sequence (5′→3′)* |
|---|---|---|
| **D157N** | Forward | ACTGGCAATT**AAT**CGACCCTCACAGAAG |
| | Reverse | TGGTGCGGTTCCACTCGC |
| **R132A** | Forward | GTCTGTGCAA**GCC**CATGGCCATG |
| | Reverse | TCATGGATGTAAAAGTCC |
| **L163A** | Forward | CTCACAGAAG**GCC**CTGAAATTCCTGAATAAGCAC |
| | Reverse | GGTCGGTCAATTGCCAGT |
| **I156G** | Forward | CCAACTGGCA**GGC**GACCGACCCTCACAG |
| | Reverse | TGCGGTTCCACTCGCTCC |
| **I121A** | Forward | ACCACTTTGC**GCG**CTGGACTTTTAC |
| | Reverse | TCTACCTCATTATGAGCC |
| **I121G** | Forward | ACCACTTTGC**GGC**CTGGACTTTTAC |
| | Reverse | TCTACCTCATTATGAGCC |

*The mutated codons are highlighted in orange.

**Table S2.** Individual *K*i values and fitted lower plateaus from FP competition assays of α-tubulin peptide substrates and bisubstrate analogs with WT-ATAT1.

| Experiment | Parameter | p4 | p10 | p11 | p4-CoA | p10-CoA | p11-CoA |
|---|---|---|---|---|---|---|---|
| 1 | ***K*i (μM)** | — | — | — | — | — | **0.77** |
| | Bottom | — | — | — | — | — | 3.6 |
| 2 | ***K*i (μM)** | — | — | **NB** | — | — | **0.30** |
| | Bottom | — | — | 87 | — | — | 27 |
| 3 | ***K*i (μM)** | — | — | — | — | — | **1.1** |
| | Bottom | — | — | — | — | — | 0.42 |
| 4 | ***K*i (μM)** | **NB** | **NB** | **NB** | **5.5** | **1.5** | **0.43** |
| | Bottom | — | — | — | 0 | 0 | 3.1 |
| 5 | ***K*i (μM)** | **NB** | **NB** | **NB** | **0.50** | — | **0.31** |
| | Bottom | — | — | — | 0 | — | 2.4 |
| 6 | ***K*i (μM)** | — | — | — | — | **0.41** | — |
| | Bottom | — | — | — | — | 0 | — |

**NB** = no binding detected. *K*i values were determined from individual FP competition experiments. Bottom values represent the lower plateau of the fitted competition curves (minimum fluorescence polarization). Em dashes (—) indicate that the parameter was not determined.

**Table S3.** Individual *K*i values and fitted lower plateaus from FP competition assays of Ac-CoA, CoA-SH, and Ac-CoA mimics with WT-ATAT1.

| Experiment | Parameter | Ac-CoA | CoA-SH | 3BY-CoA | 4PY-CoA | 5HY-CoA | 6HY-CoA | 3AZ-CoA | 4AZ-CoA | ClAc-CoA |
|---|---|---|---|---|---|---|---|---|---|---|
| 1 | **Ki (μM)** | **1.9** | — | — | — | — | — | — | — | — |
| | Bottom | 0 | — | — | — | — | — | — | — | — |
| 2 | **Ki (μM)** | **2.2** | **29** | — | — | — | — | — | **4.7** | — |
| | Bottom | 2 | 0 | — | — | — | — | — | 0 | — |
| 3 | **Ki (μM)** | **1.4** | **82** | **169** | **16** | **6.5** | **1.5** | **1.1** | **1.8** | **1.4** |
| | Bottom | 0 | 16 | 0 | 21 | 22 | 16 | 16 | 3.6 | 15 |
| 4 | **Ki (μM)** | **1.7** | **56** | **251** | **55** | **14** | **4.6** | **4.3** | **4.1** | **1.6** |
| | Bottom | 4.7 | 10 | 0 | 22 | 21 | 11 | 17 | 9.3 | 7.3 |

*K*i values were determined from individual FP competition experiments. Bottom values represent the lower plateau of the fitted competition curves (minimum fluorescence polarization). Em dashes (—) indicate that the parameter was not determined.

**Table S4.** Individual *K*i values and fitted lower plateaus from FP competition assays of Ac-CoA, CoA-SH, and Ac-CoA mimics with ATAT1-L163A.

| Experiment | Parameter | Ac-CoA | CoA-SH | 3BY-CoA | 4PY-CoA | 5HY-CoA | 6HY-CoA | 3AZ-CoA | 4AZ-CoA | ClAc-CoA |
|---|---|---|---|---|---|---|---|---|---|---|
| 1 | **Ki (μM)** | **3.4** | **41** | **90** | **0.97** | **1.33** | **1.05** | **0.26** | **1.45** | **1.3** |
| | Bottom | 0 | 0 | 0 | 0 | 3.3 | 0 | 7.1 | 0.39 | 0 |
| 2 | **Ki (μM)** | **1.6** | **15** | **145** | **0.60** | **0.67** | **0.34** | **0.27** | **0.76** | **1.0** |
| | Bottom | 0 | 0 | 0 | 2.6 | 9.4 | 3.6 | 8.7 | 0 | 0 |

*K*i values were determined from individual FP competition experiments. Bottom values represent the lower plateau of the fitted competition curves (minimum fluorescence polarization). Em dashes (—) indicate that the parameter was not determined.

**Table S5.** Individual *K*i values and fitted lower plateaus from FP competition assays of Ac-CoA, CoA-SH, and Ac-CoA mimics with ATAT1-R132A.

| Experiment | Parameter | Ac-CoA | CoA-SH | 3BY-CoA | 4PY-CoA | 5HY-CoA | 6HY-CoA | 3AZ-CoA | 4AZ-CoA | ClAc-CoA |
|---|---|---|---|---|---|---|---|---|---|---|
| 1 | **Ki (μM)** | **65** | **2112** | **NB** | **542** | **89** | **25** | **30** | **28** | **5.2** |
| | Bottom | 0 | 0 | 108 | 0 | 30 | 10 | 18 | 27 | 0 |

**NB** = no binding detected. *K*i values were determined from individual FP competition experiments. Bottom values represent the lower plateau of the fitted competition curves (minimum fluorescence polarization). Em dashes (—) indicate that the parameter was not determined.

**Table S6.** Number of independent experiments contributing to the fluorescence and western blot quantifications shown in Figure 4F.

| Substrate | ATAT1 variant | Cofactor | Fluorescence (*N*) | Western blot (*N*) |
|---|---|---|---|---|
| deAcTub | — | — | 10 | 17 |
| | WT-ATAT1 | — | 3 | 13 |
| | ATAT1-L163A | — | 5 | 14 |
| | — | Ac-CoA | 4 | 5 |
| | — | 3BY-CoA | 4 | 5 |
| | — | 4PY-CoA | 11 | 6 |
| | — | 5HY-CoA | 3 | 5 |
| | — | 6HY-CoA | 7 | 6 |
| | — | 3AZ-CoA | 4 | 7 |
| | — | 4AZ-CoA | 3 | 5 |
| | WT-ATAT1 | Ac-CoA | 4 | 15 |
| | WT-ATAT1 | 3BY-CoA | 3 | 4 |
| | WT-ATAT1 | 4PY-CoA | 3 | 5 |
| | WT-ATAT1 | 5HY-CoA | 2 | 4 |
| | WT-ATAT1 | 6HY-CoA | 3 | 5 |
| | WT-ATAT1 | 3AZ-CoA | 2 | 6 |
| | WT-ATAT1 | 4AZ-CoA | 2 | 4 |
| | ATAT1-L163A | Ac-CoA | 5 | 14 |
| | ATAT1-L163A | 3BY-CoA | 3 | 3 |
| | ATAT1-L163A | 4PY-CoA | 11 | 5 |
| | ATAT1-L163A | 5HY-CoA | 7 | 3 |
| | ATAT1-L163A | 6HY-CoA | 6 | 3 |
| | ATAT1-L163A | 3AZ-CoA | 4 | 5 |
| | ATAT1-L163A | 4AZ-CoA | 3 | 3 |

*N* values represent the number of independent experiments contributing to each readout. Em dashes (—) indicates that no ATAT1 variant or exogenous cofactor was added. All experiments used deacetylated tubulin (deAcTub).

**Table S7.** Number of independent experiments contributing to the fluorescence and western blot quantifications shown in Figure 5D.

| Substrate | ATAT1 variant | Cofactor | Fluorescence (*N*) | Western blot (*N*) |
|---|---|---|---|---|
| **deAcTub** | — | — | 10 | 4 |
| | — | 4PY-CoA | 4 | 2 |
| | WT-ATAT1 | — | 3 | 4 |
| | WT-ATAT1 | Ac-CoA | 2 | 4 |
| | ATAT1-L163A | — | 5 | 2 |
| | ATAT1-L163A | 4PY-CoA | 4 | 2 |
| **deAcMT** | — | — | 16 | 4 |
| | — | 4PY-CoA | 13 | 2 |
| | WT-ATAT1 | — | 2 | 4 |
| | WT-ATAT1 | Ac-CoA | 2 | 4 |
| | ATAT1-L163A | — | 13 | 2 |
| | ATAT1-L163A | 4PY-CoA | 11 | 2 |

*N* values represent the number of independent experiments contributing to each readout. Em dashes (—) indicates that no ATAT1 variant or exogenous cofactor was added. All experiments used deacetylated tubulin (deAcTub).

**Table S8.** Complementary colocalization metrics for ATTO-488 and CalFluor647 fluorescence signals in immobilized microtubules.

| Acylation/click workflow | Condition | *n* (ROI) | rCostes | M1 (Costes) | M2 (Costes) | ICQ* |
|---|---|---|---|---|---|---|
| Acylation in solution + click reaction in chamber | **+4PY-CoA** | 5 | 0.29 ± 0.18 | 0.27 ± 0.18 | 0.27 ± 0.17 | 0.07 ± 0.02 |
| | **+4PY-CoA +ATAT1-L163A** | 31 | 0.87 ± 0.01 | 0.72 ± 0.05 | 0.77 ± 0.02 | 0.19 ± 0.01 |
| Acylation + click reaction in chamber | **+4PY-CoA** | 5 | 0.72 ± 0.05 | 0.64 ± 0.14 | 0.80 ± 0.03 | 0.09 ± 0.02 |
| | **+4PY-CoA +WT-ATAT1** | 5 | 0.74 ± 0.18 | 0.26 ± 0.17 | 0.77 ± 0.02 | 0.06 ± 0.02 |
| | **+4PY-CoA +ATAT1-L163A** | 25 | 0.85 ± 0.02 | 0.49 ± 0.05 | 0.74 ± 0.03 | 0.18 ± 0.01 |
| | **+6HY-CoA +ATAT1-L163A** | 4 | 0.85 ± 0.02 | 0.51 ± 0.11 | 0.66 ± 0.04 | 0.11 ± 0.02 |

*Data are presented as mean ± SEM. rCostes, Pearson's correlation coefficient above the Costes-determined threshold; M1 and M2, Costes-thresholded Manders' overlap coefficients; ICQ, Li's Intensity Correlation Quotient.

### EXPERIMENTAL PROCEDURES

#### General synthetic procedures

All reagents and solvents used for organic synthesis were purchased from commercial sources and used without further purification. Chemicals were obtained from Sigma-Aldrich, Acros, Fluorochem, and other commercial suppliers as indicated. Automated solid-phase peptide synthesis was carried out on an Intavis AG MultiPep RS instrument. Fmoc-protected amino acids, Fmoc-L-propargylglycine, and Rink Amide HL resin for solid-phase peptide synthesis were purchased from Sigma-Aldrich, Novabiochem, and Fluorochem. Coenzyme A (free acid) was obtained from Larodan AB, and 5-carboxytetramethylrhodamine azide (TAMRA-azide) was purchased from Carl Roth.

For the synthesis of acetyl coenzyme A (Ac-CoA) mimics, analytical thin-layer chromatography (TLC) was performed on silica gel 60 plates (Merck, 0.2 mm) and visualized by staining with iodine or bromocresol green. Purifications were performed by reverse-phase high-performance liquid chromatography (RP-HPLC). Ac-CoA mimics were purified on a Shimadzu HPLC system using a C18 column with a linear gradient of 10–50% buffer B over 60 min at a flow rate of 7 mL min$^{-1}$. Unless otherwise stated, buffer A consisted of 0.1% trifluoroacetic acid (TFA) in $H_2O$, and buffer B consisted of 0.1% TFA in acetonitrile (ACN). UV detection was performed at 260 nm. Peptide substrates and peptide-CoA conjugates were purified on an Agilent Technologies system equipped with a diode array detector using an Agilent Zorbax C18 column (5 µm, 9.4 × 250 mm).

Liquid chromatography–mass spectrometry (LC–MS) analyses were performed on a Dionex UltiMate 3000 UHPLC coupled to a Thermo LCQ Fleet mass spectrometer using electrospray ionization (ESI) in positive ion mode. Analyses were performed using a Pinnacle DB C18 column (1.9 µm, 50 × 2.1 mm). The elution gradient was 95:5 to 10:90 (A:B) over 4 min at a flow rate of 0.750 mL min$^{-1}$, where buffer A consisted of 0.01% TFA in $H_2O$ and buffer B consisted of 0.01% TFA in HPLC-grade acetonitrile. Xcalibur™ software (Thermo Fisher Scientific) was used to determine the nominal masses of the major peaks.

All nuclear magnetic resonance (NMR) spectra were recorded at 25 °C on Bruker spectrometers operating at 500 MHz for $^{1}H$ and $^{13}C$ NMR and 400 MHz for $^{31}P$ NMR using $D_2O$ as the solvent. Chemical shifts (δ) are reported in parts per million (ppm) relative to the internal standard tetramethylsilane (TMS, δ = 0 ppm). Spin multiplicities are reported as singlet (s), doublet (d), triplet (t), doublet of triplets (dt), quintet (p), or multiplet (m), and coupling constants (J) are reported in Hz.

High-resolution mass spectrometry (HRMS) analyses were performed at the Chemical Biology Mass Spectrometry Core Facility (ChemBioMS, University of Geneva, Switzerland) by chip-based nanospray infusion using a TriVersa NanoMate (Advion Interchim Scientific, Harlow, UK) coupled to a Q Exactive Plus hybrid quadrupole-Orbitrap high-resolution mass spectrometer (Thermo Fisher Scientific). For LC–MS and HRMS analyses, samples were prepared in methanol (MeOH) or distilled deionized $H_2O$ (dd$H_2O$) containing 0.1% formic acid (FA).

The concentrations of all CoA-containing compounds, including peptide-CoA conjugates and Ac-CoA mimics, were determined by UV–Vis spectroscopy using a Thermo Fisher NanoDrop 2000 spectrophotometer in dd$H_2O$. Concentrations were calculated using the molar extinction coefficient of coenzyme A (ε = 15,000 $M^{-1}$ $cm^{-1}$) at $\lambda_{max}$ = 260 nm.

**General procedure for automated SPPS of peptide sequences**

Fifty milligrams of Rink Amide HL resin (0.70–1.00 mmol $g^{-1}$, Novabiochem) was swollen in dichloromethane (DCM) for 10 min and washed twice with dimethylformamide (DMF). Peptides were synthesized by automated Fmoc-based solid-phase peptide synthesis (SPPS) using iterative cycles consisting of Fmoc deprotection (Procedure 1), amino acid coupling (Procedure 2), and capping of unreacted amines (Procedure 3). After completion of the peptide sequence, the N-terminus was deprotected and capped using the same procedures. When required, the ivDde protecting group was removed manually (Procedure 4), followed by coupling of (*R*)-2-bromopropionic acid as a conjugation handle (Procedure 5). Finally, the peptides were cleaved from the resin and purified by RP-HPLC (Procedure 6). The click chemistry procedure used for the synthesis of p11-CoA-TAMRA is described separately.

The Fmoc-protected amino acids used were Fmoc-Lys(ivDde)-OH (Novabiochem) and Fmoc-Gly-OH, Fmoc-Ser(tBu)-OH, Fmoc-Thr(tBu)-OH, Fmoc-Asp(tBu)-OH, Fmoc-Ile-OH, Fmoc-Gln(Trt)-OH, Fmoc-Met-OH, Fmoc-Pro-OH, and Fmoc-N-(propargyl)glycine (Fluorochem).

**Procedure 1: Fmoc deprotection.** The resin was treated with 20% piperidine in DMF for 5 min. The solution was removed by filtration, and the resin was washed twice with DMF. This deprotection step was repeated once. Finally, the resin was washed with DMF (2×), DCM (2×), and DMF (2×).

**Procedure 2: Amide coupling.** The appropriate Fmoc-protected amino acid (4 equiv.; 0.2 M in *N*-methyl-2-pyrrolidone, NMP) was preactivated for 5 min with HATU (3.5 equiv.; 0.5 M in NMP) in the presence of *N,N*-diisopropylethylamine (DIPEA, 4 equiv.; 1.2 M) and 2,6-lutidine (6 equiv.; 1.8 M) in NMP. The activated solution was added to the resin and allowed to react for 20 min. The resin was then filtered and washed twice with DMF.

**Procedure 3: Capping.** The resin was treated with a capping solution consisting of acetic anhydride (0.92 mL) and 2,6-lutidine (1.3 mL) in DMF (18 mL; 10 mL solution per gram of resin) for 5 min. The resin was filtered and washed with DMF (2×), DCM (2×), and DMF (2×). Following coupling of the final amino acid, the terminal Fmoc group was removed, and the N-terminus was capped using the same procedure. After completion of the automated SPPS, the resin was handled manually for subsequent modifications.

**Procedure 4: Removal of the ivDde protecting group.** Where required, ivDde-protected lysine residues were incorporated to enable orthogonal functionalization. The ivDde protecting group was removed by treating the resin with 3% hydrazine in DMF (5 mL) for 2 h at room temperature on a rotary mixer. The resin was then washed thoroughly with DMF (2×), DCM (2×), and DMF (2×; 3 × 5 mL).

**Procedure 5: Coupling of (*R*)-2-bromopropionic acid.** To introduce a conjugation handle, the free amine on the resin-bound peptide was coupled with (*R*)-2-bromopropionic acid. First, the corresponding NHS ester was prepared by adding N-hydroxysuccinimide (394.97 mg, 3.43 mmol, 1.05 equiv.) and 1-ethyl-3-(3-dimethylaminopropyl)carbodiimide hydrochloride (EDC·HCl, 532.79 mg, 3.43 mmol, 1.05 equiv.) to a solution of (*R*)-2-bromopropionic acid (500 mg, 3.27 mmol, 1.0 equiv.) in DCM (100 mL). The reaction mixture was stirred overnight at room temperature, washed three times with $H_2O$ (100 mL each) and twice with brine (100 mL), dried over $MgSO_4$, filtered, and concentrated under reduced pressure to afford the NHS ester of (*R*)-2-bromopropionic acid as a crude yellow oil (400 mg, 49%, approximately 70% purity), which was used without further purification. The resin-bound peptide was then treated with a solution of the NHS ester (150 mg, 0.60 mmol, 3 equiv.) in DMF (5 mL) for 1 h at room temperature. After coupling, the resin was washed thoroughly with DMF (2×), DCM (2×), and DMF (2×; 3 × 5 mL).

**Procedure 6: Global deprotection and cleavage.** Acid-labile *tert*-butyl groups were removed, and peptides were cleaved from the resin using a cleavage cocktail of TFA/TIPS/$H_2O$ (95:2.5:2.5, v/v/v) for 2 h at room temperature with gentle shaking. The cleavage solution was transferred to a 50 mL Falcon tube, and the peptide was precipitated by the addition of cold diethyl ether ($Et_2O$, –20 °C). The suspension was centrifuged (6000 × g, 5 min), and the supernatant was discarded. The peptide pellet was dissolved in DMSO/$H_2O$ containing 0.1% TFA and purified by RP-HPLC using one of three methods: method A (10–90% acetonitrile in $H_2O$ over 50 min), method B (5–90% acetonitrile in $H_2O$ over 50 min), or method C (1–90% acetonitrile in $H_2O$ over 50 min), each containing 0.1% TFA. Fractions containing the desired peptide were pooled, lyophilized, and stored at –20 °C until use.

##### Chemical synthesis and characterization of peptide substrates

For the synthesis of peptide substrates, Procedures 1–3 were used, followed by Procedure 6 to obtain the desired peptides. Synthetic αK40 peptide substrates p4 (Ac-SD**K**T-$NH_2$), p10 (Ac-GQMPSD**K**TIG-$NH_2$), and p11 (Ac-SD**K**TIGGGDDS-$NH_2$) were dissolved in $H_2O$ to a final concentration of 10 mM. All peptide substrates were N-terminally acetylated.

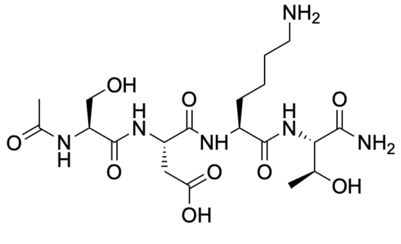

**p4**

Ac-SDKT-$NH_2$ (**p4**). HPLC method A (5.2 mg, 10.5 *μ*mol, 30%). LC–MS analysis: Rt = 0.28 min. HRMS (ESI): calcd for $C_{19}H_{35}N_6O_9$ $[M+H]^+$ *m/z* 491.2460; found 491.2463.

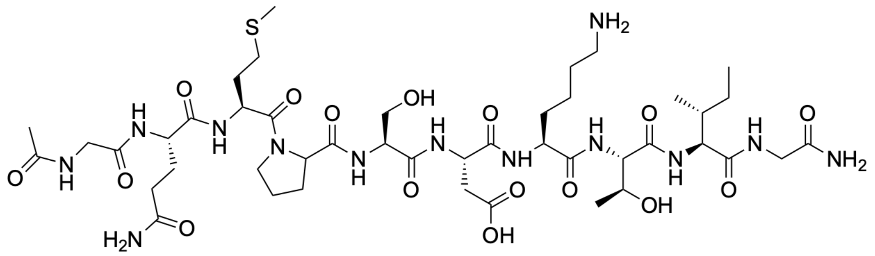

**p10**

Ac-GQMPSDKTIG-NH$_2$ (**p10**). HPLC method B (10.5 mg, 9.3 $\mu$mol, 26%) LC–MS analysis: Rt = 1.32 min. HRMS (ESI): calcd for $C_{44}H_{77}N_{13}O_{16}S$ $[M+2H]^{2+}$ *m/z* 537.7660; found 537.7788.

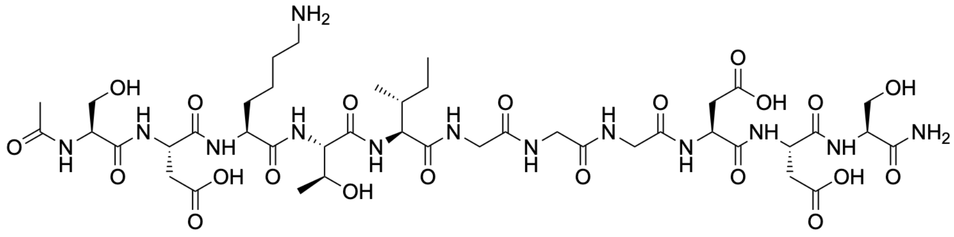

**p11**

Ac-SDKTIGGGDDS-NH$_2$ (**p11**). HPLC method B (13.5 mg, 12.4 $\mu$mol, 36%). LC–MS analysis: Rt = 1.09 min. HRMS (ESI): calcd for $C_{42}H_{70}N_{13}O_{21}$ $[M+H]^{+}$ *m/z* 1092.4804; found 1092.4809.

**Chemical synthesis and characterization of peptide-CoA conjugates**

Peptide-CoA conjugates were synthesized according to Scheme S1. The peptide backbones (**p4-PrBr**, **p10-PrBr**, and **p11-PrBr**) were synthesized using Procedures 1–6 described above. The general procedure for the coupling of CoA-SH to obtain p4-CoA, p10-CoA and p11-CoA can be found below.

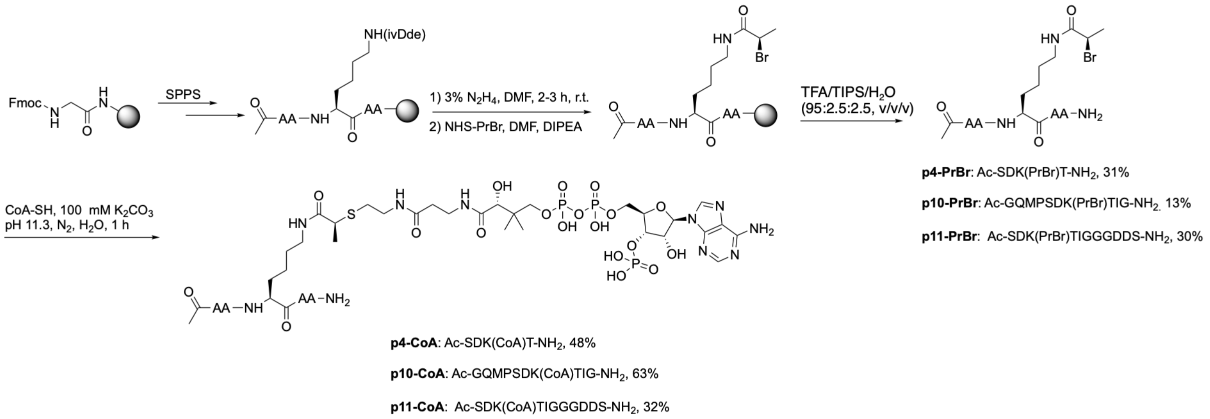

**Scheme S1.** General chemical synthesis of peptide-CoA conjugates.

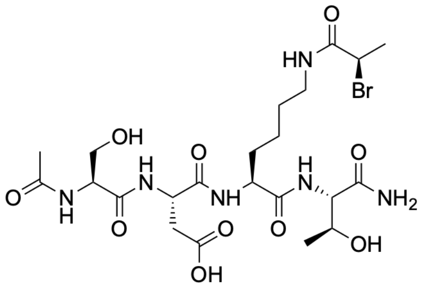

**p4-PrBr**

Ac-SDK(PrBr)T-NH$_2$ (**p4-PrBr**). HPLC method A (35.5 mg, 57 μmol, 31%). LC–MS analysis: Rt = 1.19 min. HRMS (ESI): calcd for $C_{22}H_{38}BrN_6O_{10}$ $[M+H]^+$ *m/z* 625.1827; found 625.1828.

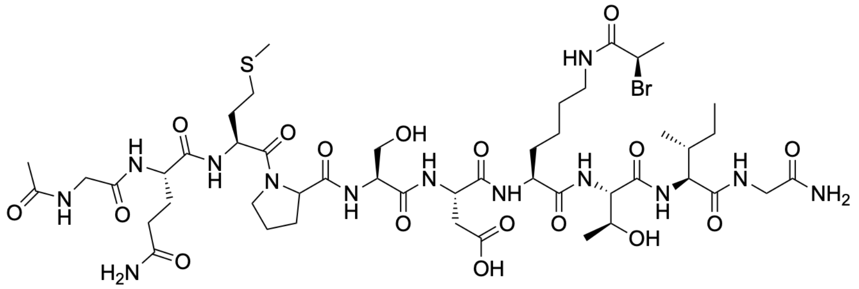

**p10-PrBr**

Ac-GQMPSDK(PrBr)TIG-NH$_2$ (**p10-PrBr**). HPLC method B (15.5 mg, 12.8 μmol, 13%). LC–MS analysis: Rt = 1.63 min. HRMS (ESI): calcd for $C_{47}H_{79}BrN_{13}O_{17}S$ $[M+H]^+$ *m/z* 1208.4615; found 1208.4845

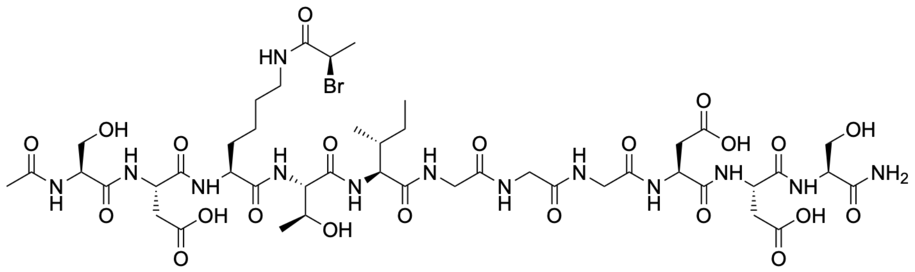

**p11-PrBr**

Ac-SDK(PrBr)TIGGGDDS-NH$_2$ (**p11-PrBr**). HPLC method B (12 mg, 9.8 μmol, 30%). LC–MS analysis: Rt = 1.45 min. HRMS (ESI): calcd for $C_{45}H_{73}BrN_{13}O_{22}$ $[M+H]^+$ *m/z* 1226.4171; found 1226.4171.

Exemplary synthesis of peptide-CoA conjugate based on p11: A small 5 mL flask was charged with CoA-SH (3.8 mg, 4.9 μmol, 2 equiv.) and purged with $N_2$. A second flask was charged with **p11-PrBr** (3 mg, 2.4 μmol, 1 equiv.), purged with $N_2$, and the compound was dissolved in previously degassed 100 mM $K_2CO_3$ buffer (pH = 11.3) (or 100 mM $NaHCO_3$ buffer, pH = 8.5, for Ac-p4/10-PrBr). The solution of **p11-PrBr** was transferred by syringe and added to the flask containing CoA-SH. The mixture was stirred for 1–2 h (or 24 h for Ac-p4/10-PrBr) at room temperature. The reaction was monitored by LC–MS. Upon completion, the reaction mixture was slowly neutralized with acetic acid (AcOH) and purified by HPLC.Fractions containing the desired product were flash-frozen immediately after purification to minimize degradation and lyophilized to afford the corresponding CoA-peptides as a white powder.

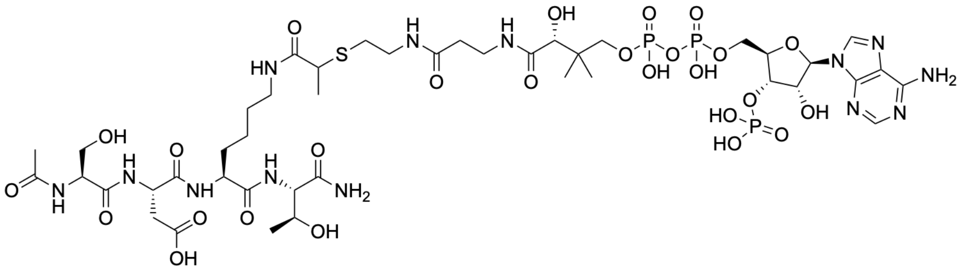

**p4-CoA**

Ac-SDK(CoA)T-$NH_2$ (**p4-CoA**). HPLC method A (2 mg, 3.2 μmol, 48%). LC–MS analysis: Rt = 0.45 min. HRMS (ESI): calcd for $C_{43}H_{74}N_{13}O_{26}P_3S$ $[M+2H]^{2+}$ *m/z* 656.6895; found 656.6891.

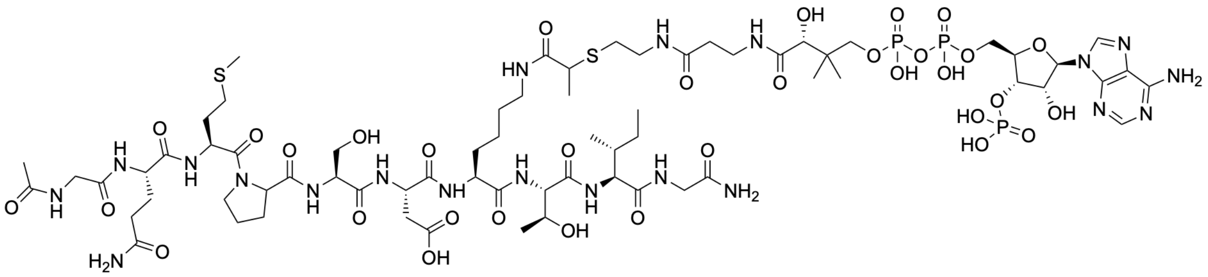

**p10-CoA**

Ac-GQMPSDK(CoA)TIG-$NH_2$ (**p10-CoA**). HPLC method B (2 mg, 1 μmol, 63%). LC–MS analysis: Rt = 1.32 min. HRMS (ESI): calcd for $C_{68}H_{115}N_{20}O_{33}P_3S_2$ $[M+2H]^{2+}$ *m/z* 948.3289; found 948.2887.

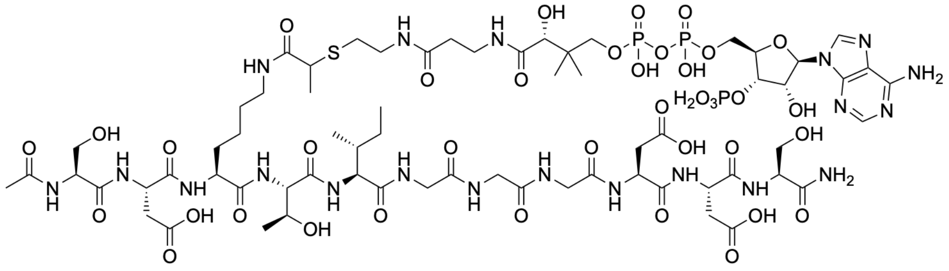

**p11-CoA**

Ac-SDK(CoA)TIGGGDDS-$NH_2$ (**p11-CoA**). HPLC method B (1.5 mg, 0.8 μmol, 32%). LC–MS analysis: Rt = 1.16 min. HRMS (ESI): calcd for $C_{66}H_{109}N_{20}O_{38}P_3S$ $[M+2H]^{2+}$ *m/z* 957.3067; found 957.3060.

#### Chemical synthesis and characterization of tracer p11-CoA-TAMRA

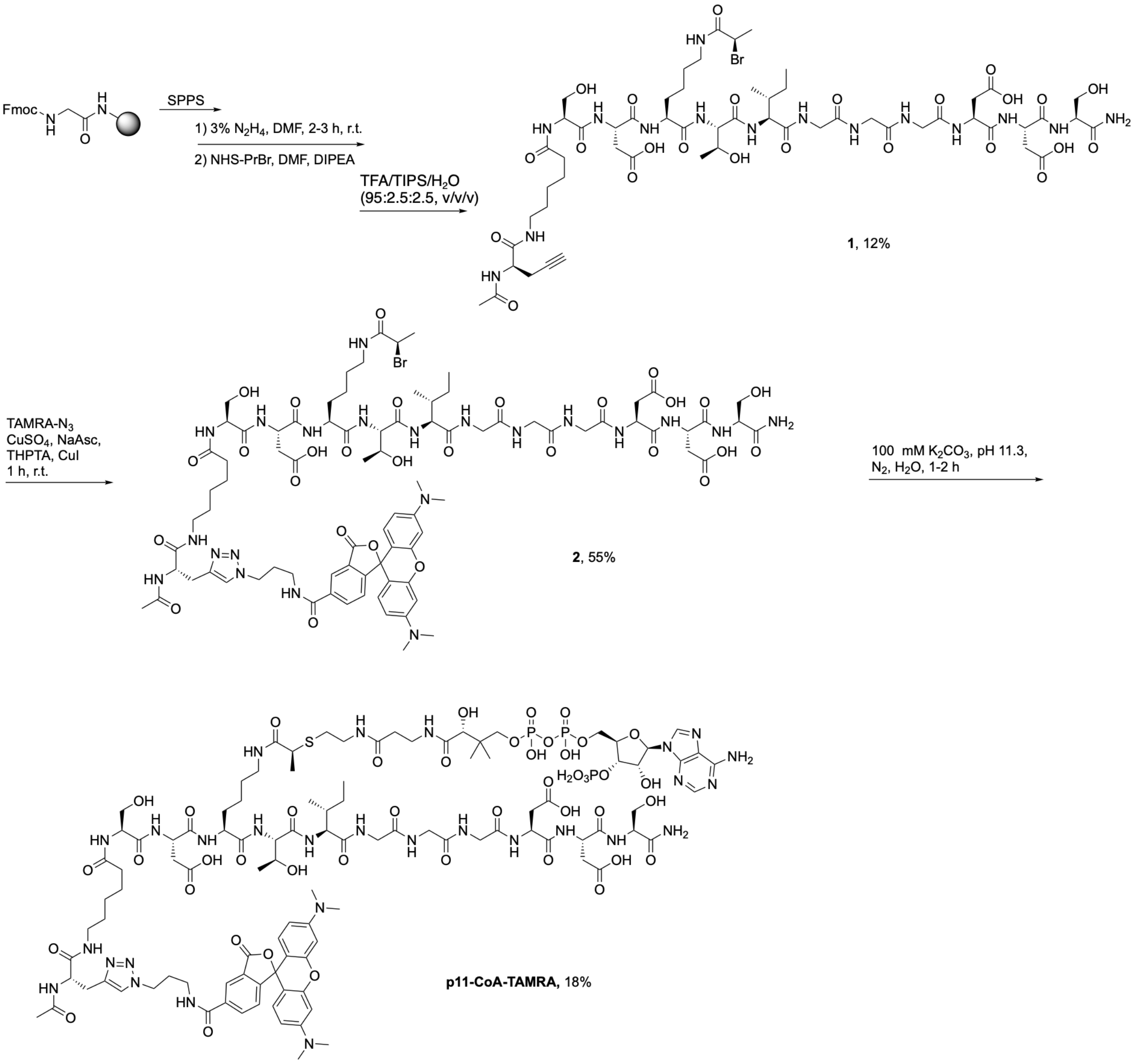

**Scheme S2**. Chemical synthesis of tracer p11-CoA-TAMRA.

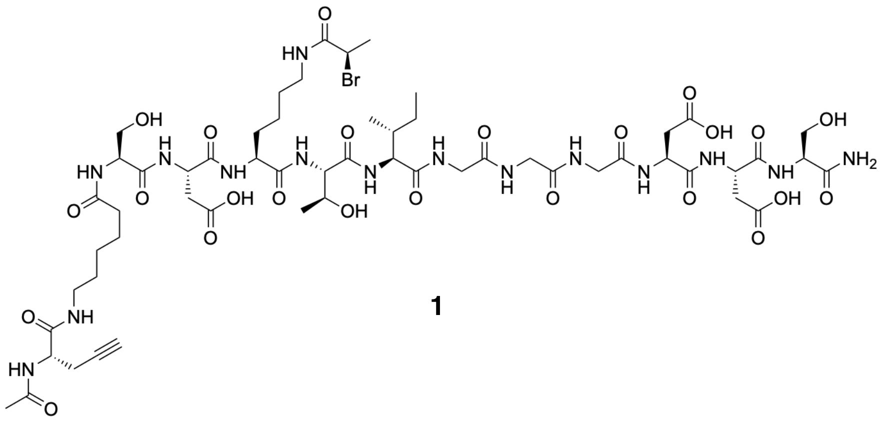

**1**

Ac-p11-Anx-Propargyl-PrBr (**1**). The peptide backbone was synthesized using the same SPPS procedure 1-3 described above. After assembly of p11, 6-aminohexanoic acid was introduced as a linker, followed by incorporation of propargylalanine as the clickable handle. After SPPS, (*R*)-2-bromopropionic acid was introduced as described above (General Procedure 4, 5). A portion of the resin (140 mg, 100 *μ*mol) was deprotected and cleaved using the cleavage mixture TFA/TIPS/$H_2O$ (95:2.5:2.5, v/v/v) for 2 h with stirring. The cleavage mixture was transferred to a 50 mL Falcon tube, and the peptide was precipitated by addition of cold $Et_2O$ (–20 °C). The precipitated peptide was dissolved in DMSO/$H_2O$ containing 0.1% TFA and purified by HPLC (method C, gradient $H_2O$/ACN 10–90%). Fractions containing the desired peptide were collected, pooled, and lyophilized to afford Ac-p11-Anx-Propargyl-PrBr (**1**) (18 mg, 12.5 *μ*mol, 12% yield). LC–MS analysis: Rt = 1.57 min. HRMS (ESI): calcd for $C_{56}H_{89}BrN_{15}O_{24}$ $[M+H]^+$ *m/z* 1434.5383; found 1434.5382.

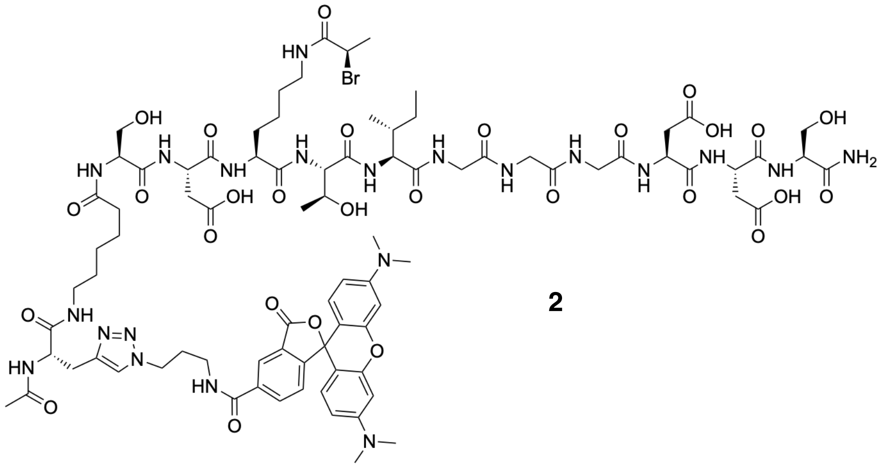

**2**

Ac-p11-TAMRA-PrBr (**2**). The obtained alkyne-functionalized peptide (**1**) was dissolved in DMSO/$H_2O$ (1 mL) and reacted with TAMRA-azide using CuAAC click chemistry. Briefly, TAMRA-azide (155 *μ*L, 10 mM, 1.1 equiv.) was added to a solution of peptide (**1**) (2 mg, 1.4 *μ*mol, 1 equiv.), followed by $CuSO_4$ solution (700 *μ*L, 10 mM, 5 equiv.), THPTA solution (77 *μ*L, 100 mM, 5 equiv.), and freshly prepared sodium ascorbate solution (28 *μ*L, 500 mM, 10 equiv.). CuI (1.3 mg, 7 *μ*mol, 5 equiv.) was added to ensure reaction completion. The reaction mixture was stirred for 1 h at room temperature. After completion, the mixture was acidified by addition of AcOH and purified by HPLC (method C, gradient $H_2O$/ACN 10–90%). Fractions containing the desired product were collected, pooled, and lyophilized to afford Ac-p11-TAMRA-PrBr (**2**) (1.5 mg, 0.77 *μ*mol, 55%) as a pink solid. LC–MS analysis: Rt = 1.86 min. HRMS (ESI): calcd for $C_{84}H_{117}BrN_{21}O_{28}$ $[M+H]^+$ *m/z* 1946.7555; found 1946.7498.

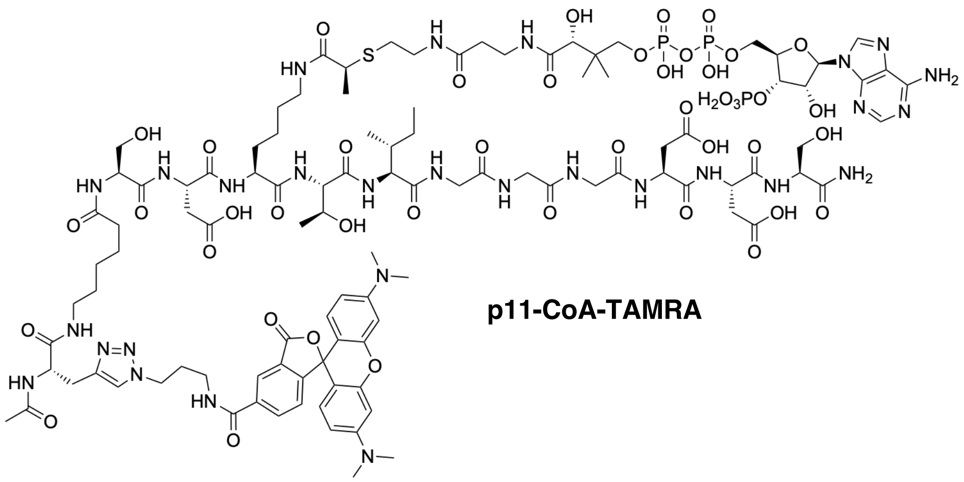

**p11-CoA-TAMRA**

Ac-p11-CoA-TAMRA (**p11-CoA-TAMRA**). A 5 mL flask was charged with CoA-SH (2.4 mg, 3.1 $\mu$mol, 3 equiv.) and purged with $N_2$. A second flask was charged with Ac-p11-TAMRA-PrBr (**2**) (3 mg, 1 $\mu$mol, 1 equiv.), purged with $N_2$, and dissolved in previously degassed 100 mM $K_2CO_3$ buffer (pH = 11.3). The solution of Ac-p11-TAMRA-PrBr (**2**) was transferred by syringe and added to the flask containing CoA-SH. The mixture was stirred for 1 h at room temperature. The reaction was monitored by LC–MS. Upon completion, the reaction mixture was slowly neutralized with AcOH and purified by HPLC (method B, gradient $H_2O$/ACN 5–90%). Fractions containing the desired product were flash-frozen immediately after purification to minimize degradation and lyophilized to afford **p11-CoA-TAMRA** as a pink powder (0.5 mg, 0.2 $\mu$mol, 18% yield). LC–MS analysis: Rt = 1.32 min. HRMS (ESI): calcd for $C_{105}H_{153}N_{28}O_{44}P_3S$ $[M+2H]^{2+}$ *m/z* 1317.4759; found 1317.4777.

##### Chemical synthesis and characterization of Ac-CoA mimics

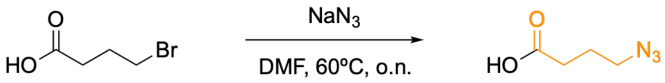

**3**

**Scheme S3**. Chemical synthesis of 4-azidobutanoic acid (**3**). The acyl ligation handle is highlighted in orange.

4-azidobutanoic acid (**3**). To a solution of 5 mmol of 4-bromobutanoic acid (1 equiv.) in dry DMF (5 mL) under argon was added 5 mmol of $NaN_3$ (1 equiv.) and the mixture was stirred at 60°C overnight. Reaction was monitored by TLC plate by using DCM/MeOH 9/1 + 0.1% acetic acid as solvent. After cooling to room temperature, ethyl acetate (20 mL) and aqueous HCl 0.1 M (20 mL) were successively added, and the aqueous phase was extracted twice with ethyl acetate. The combined organic extracts were washed with 10% LiCl and brine, dried over $MgSO_4$, filtered, and concentrated under vacuum. The product was then lyophilized to remove DMF and afford compound **3** as a brown slurry which was used in the next step without further purification. $^1$H NMR (400 MHz, $CDCl_3$) δ 3.39 (t, J = 6.7 Hz, 2H), 2.48 (t, J = 7.2 Hz, 2H), 1.98 – 1.87 (m, 2H). $^{13}$C NMR (400 MHz, $CDCl_3$) δ 50.61, 30.75.

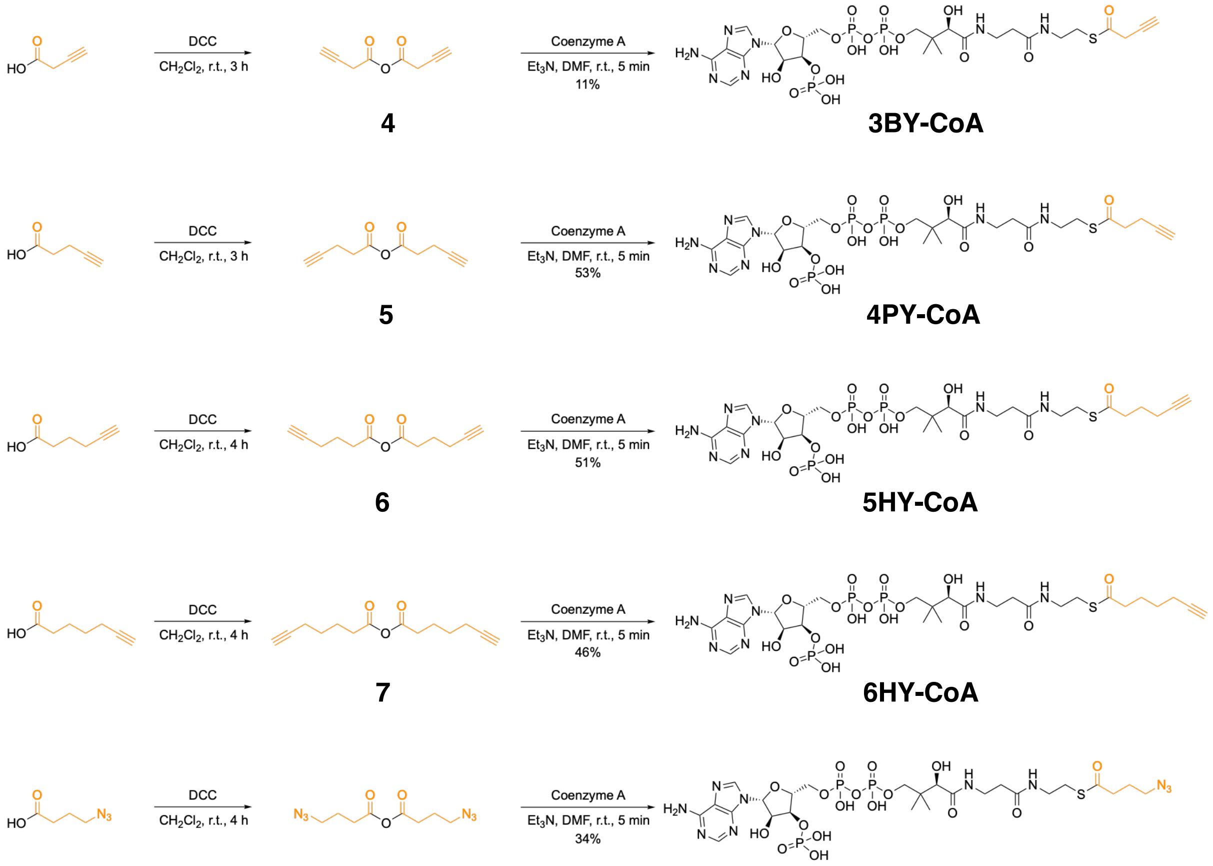

4 3BY-CoA 5 4PY-CoA 6 5HY-CoA 7 6HY-CoA 3 8 4AZ-CoA

**Scheme S4**. Chemical synthesis of 3BY-CoA, 4PY-CoA, 5HY-CoA, 6HY-CoA, and 4AZ-CoA. Acyl ligation handles are highlighted in orange.

<u>3-Butynoyl-CoA (**3BY-CoA**), 4-pentynoyl-CoA (**4PY-CoA**), 5-hexynoyl-CoA (**5HY-CoA**), 6-heptynoyl-CoA (**6HY-CoA**), and 4-azidobutanoyl-CoA (**4AZ-CoA**)</u> were synthesized based on a previously reported procedure.[1] 0.1 mmol of 3-butynoic acid, 4-pentynoic acid, 5-hexynoic acid, 6-heptynoic acid, or 4-azidobutanoic acid **3** (6 equiv. each) was dissolved in 500 $\mu$L of anhydrous DCM, under argon. To this solution was added 0.05 mmol of *N,N'*-dicyclohexylcarbodiimide (DCC, 3 equiv.), and the reaction was allowed to proceed at room temperature for 3–4 h under argon. The corresponding anhydride products (**4**, **5**, **6**, **7**, or **8**) were monitored by TLC using ethyl acetate/pentane (1:1) as the eluent. The reaction mixture was then filtered to remove dicyclohexylurea (DCU). DCM was removed by rotary evaporation at 30 °C. The dried crude material was redissolved in anhydrous DMF (500 $\mu$L) under argon and cooled in an ice bath to give solution A. To solution A was added coenzyme A (0.02 mmol, 1 equiv.) and 0.09 mmol of triethylamine ($Et_3N$, 5 equiv.). The reaction mixture was stirred for 5 min at room temperature under argon. Completion of the reaction was monitored by LC/MS. The reaction was quenched by addition of $H_2O$ (6 mL) and lyophilized. The lyophilizate was then dissolved in $H_2O$, filtered,

and subjected to two rounds of HPLC purification to obtain pure final products as white powders (**3BY-CoA** → 1.6 mg, 2 $\mu$mol, 11%; **4PY-CoA** → 8.3 mg, 10 $\mu$mol, 53%; **5HY-CoA** → 7.5 mg, 9 $\mu$mol, 51%; **6HY-CoA** → 6.9 mg, 8 $\mu$mol, 46%; or **4AZ-CoA** → 5 mg, 6 $\mu$mol, 34%). The fractions were collected and lyophilized, and the dried products were stored at −20°C for subsequent experiments. Stock solution concentrations were calculated based on the isolated mass and final solution volume and confirmed by UV–Vis spectroscopy before use.

All synthesized Ac-CoA mimics were characterized by NMR spectroscopy and HRMS:

**3BY-CoA:** $^{1}$H NMR (500 MHz, $D_2O$) δ 8.70 (s, 1H), 8.47 (s, 1H), 6.24 (d, *J* = 5.4 Hz, 1H), 5.63 (s, 1H), 4.95 – 4.89 (m, 2H), 4.63 (s, 1H), 4.36 – 4.23 (m, 2H), 4.06 (s, 1H), 3.94 – 3.84 (m, 1H), 3.68 – 3.60 (m, 1H), 3.50 (t, *J* = 6.8 Hz, 2H), 3.44 (t, *J* = 7.2 Hz, 2H), 2.97 (t, *J* = 6.7 Hz, 2H), 2.49 (t, *J* = 6.7 Hz, 2H), 2.30 (s, 2H), 0.97 (s, 3H), 0.85 (s, 3H). $^{13}$C NMR (126 MHz, $D_2O$) δ 174.73, 174.08, 168.52, 160.67, 149.91, 148.51, 144.66, 142.54, 118.59, 107.94, 87.53, 83.51, 74.21, 73.97, 71.92, 65.12, 42.26, 38.32, 37.24, 35.31, 30.15, 23.06, 20.83, 20.34, 18.31. $^{31}$P NMR (126 MHz, $D_2O$) δ -0.39, -11.04, -11.48. HRMS (ESI): calcd for $C_{25}H_{34}N_7NaO_{17}P_3S$ $[M+Na]^+$ *m/z* 852.0837; found 852.1381.

**4PY-CoA:** $^{1}$H NMR (500 MHz, $D_2O$) δ 8.70 (s, 1H), 8.47 (s, 1H), 6.26 (d, *J* = 5.9 Hz, 1H), 4.94 – 4.88 (m, 2H), 4.63 (s, 1H), 4.33 – 4.24 (m, 2H), 4.05 (s, 1H), 3.92 – 3.86 (m, 1H), 3.66 – 3.60 (m, 1H), 3.48 (t, *J* = 7.2 Hz, 2H), 3.37 (t, *J* = 6.5 Hz, 2H), 3.06 (t, *J* = 6.5 Hz, 2H), 2.86 (t, *J* = 7.0 Hz, 2H), 2.53 (dt, *J* = 7.1 Hz, 2H), 2.46 (t, *J* = 7.2 Hz, 2H), 2.39 – 2.36 (m, 1H), 0.96 (s, 3H), 0.84 (s, 3H). $^{13}$C NMR (126 MHz, $D_2O$) δ 201.90, 174.73, 174.02, 149.95, 148.55, 144.67, 142.56, 118.61, 87.49, 83.61, 83.15, 74.22, 74.14, 71.94, 70.05, 65.12, 41.70, 38.57, 35.40, 35.33, 28.05, 20.86, 18.18, 14.09. $^{31}$P NMR (126 MHz, $D_2O$) δ -0.43, -11.05, -11.51. HRMS (ESI): calcd for $C_{26}H_{41}N_7O_{17}P_3S$ $[M+H]^+$ *m/z* 848.1487; found 848.1465.

**5HY-CoA:** $^{1}$H NMR (500 MHz, $D_2O$) δ 8.67 (s, 1H), 8.45 (s, 1H), 6.22 (d, *J* = 5.9 Hz, 1H), 4.94 – 4.88 (m, 2H), 4.62 (s, 1H), 4.34 – 4.25 (m, 2H), 4.04 (s, 1H), 3.92 – 3.87 (m, 1H), 3.67 – 3.63 (m, 1H), 3.47 (t, *J* = 6.9 Hz, 2H), 3.35 (t, *J* = 6.5 Hz, 2H), 3.01 (t, *J* = 6.5 Hz, 2H), 2.74 (t, *J* = 7.3 Hz, 2H), 2.45 (t, *J* = 7.0 Hz, 2H), 2.37 – 2.35 (m, 1H), 2.24 (dt, *J* = 7.1 Hz, 2H), 1.82 (p, *J* = 7.2 Hz, 2H), 0.96 (s, 3H), 0.84 (s, 3H). $^{13}$C NMR (126 MHz, $D_2O$) δ 203.57, 174.71, 173.98, 149.86, 148.48, 144.69, 142.49, 118.53, 87.49, 84.53, 83.31, 74.15, 73.86, 72.08, 70.04, 65.14, 42.21, 38.54, 35.30, 28.04, 23.85, 20.81, 18.26, 16.88. $^{31}$P NMR (126 MHz, $D_2O$) δ -0.55, -11.14, -11.56. HRMS (ESI): calcd for $C_{27}H_{43}N_7O_{17}P_3S$ $[M+H]^+$ *m/z* 862.1644; found 862.1630.

**6HY-CoA:** $^{1}$H NMR (500 MHz, $D_2O$) δ 8.67 (s, 1H), 8.45 (s, 1H), 6.23 (d, *J* = 5.8 Hz, 1H), 4.97 – 4.88 (m, 2H), 4.65 – 4.60 (m, 1H), 4.34 – 4.25 (m, 2H), 4.04 (s, 1H), 3.93 – 3.87 (m, 1H), 3.68 – 3.62 (m, 1H), 3.47 (t, *J* = 6.7 Hz, 2H), 3.35 (t, *J* = 6.4 Hz, 2H), 3.01 (t, *J* = 6.4 Hz, 2H), 2.65 (t, *J* = 7.3 Hz, 2H), 2.45 (t, *J* = 7.0 Hz, 2H), 2.36 – 2.32 (m, 1H), 2.20 (dt, *J* = 7.2 Hz, 2H), 1.71 (p, *J* = 7.6 Hz, 2H), 1.50 (p, *J* = 7.6 Hz, 2H), 0.96 (s, 3H), 0.84 (s, 3H). $^{13}$C NMR (126 MHz, $D_2O$) δ 204.22, 174.69, 173.95, 149.86, 148.48, 144.68, 142.49, 118.53, 87.48, 85.53, 83.34, 74.14, 73.83, 72.07, 69.38, 65.09, 42.86, 38.58, 35.32, 27.98, 26.80, 24.31, 20.81, 18.24, 17.20. $^{31}$P NMR (126 MHz, $D_2O$) δ -0.53, -11.13, -11.55. HRMS (ESI): calcd for $C_{28}H_{45}N_7O_{17}P_3S$ $[M+H]^+$ *m/z* 876.1800; found 876.1780.

**4AZ-CoA:** $^1$H NMR (500 MHz, $D_2O$) δ 8.68 (s, 1H), 8.46 (s, 1H), 6.23 (d, $J$ = 5.3 Hz, 1H), 4.94 – 4.89 (m, 2H), 4.62 (s, 1H), 4.33 – 4.25 (m, 2H), 4.04 (s, 1H), 3.92 – 3.88 (m, 1H), 3.66 – 3.62 (m, 1H), 3.47 (t, $J$ = 6.8 Hz, 2H), 3.37 – 3.34 (m, 4H), 3.03 (t, $J$ = 6.5 Hz, 2H), 2.73 (t, $J$ = 7.3 Hz, 2H), 2.46 (t, $J$ = 6.8 Hz, 2H), 1.91 (p, $J$ = 7.1 Hz, 2H), 0.96 (s, 3H), 0.84 (s, 3H). $^{13}$C NMR (126 MHz, $D_2O$) δ 203.17, 174.72, 173.99, 149.89, 148.50, 144.67, 142.52, 118.56, 87.50, 83.36, 74.15, 72.04, 65.09, 50.16, 40.58, 38.54, 35.30, 28.09, 24.38, 20.81, 18.23. $^{31}$P NMR (126 MHz, $D_2O$) δ -0.42, -11.05, -11.51. HRMS (ESI): calcd for $C_{25}H_{42}N_{10}O_{17}P_3S$ [M+H]$^+$ *m/z* 879.1657; found 879.1683.

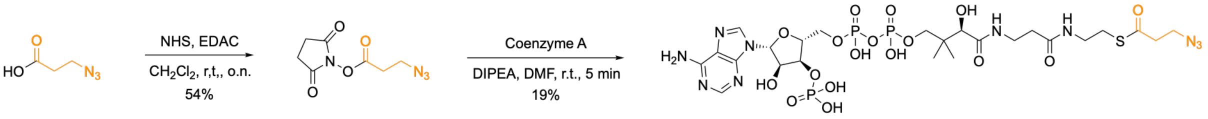

**9** **3AZ-CoA**

**Scheme S5**. Chemical synthesis of 3AZ-CoA. The acyl ligation handle is highlighted in orange.

<u>3-azidopropanoic NHS ester (**9**).</u> 0.46 mmol of N-hydroxysuccinimide (NHS, 1.05 equiv.) and 0.46 mmol of 1-ethyl-3-(3-dimethylaminopropyl)carbodiimide (EDAC, 1.05 equiv.) were added to a solution of 0.44 mmol of 3-azidopropanoic acid (1 equiv.) in anhydrous DCM (2 mL) and stirred overnight at room temperature under argon. The NHS ester product was monitored by TLC using ethyl acetate/pentane (1/1) as eluent. The reaction mixture was then washed three times with $H_2O$ and once with brine. The combined organic layers were dried over $MgSO_4$, filtered, and removed under reduced pressure to yield compound **9** as a pale-yellow oil (50 mg, 235.7 $\mu$mol, 54%), which was used in the next step without further purification. $^1$H NMR (400 MHz, $CDCl_3$) δ 3.68 (t, 2H), 2.91 – 2.82 (m, 6H). $^{13}$C NMR (400 MHz, $CDCl_3$) δ 168.93, 166.47, 46.18, 31.27, 25.70.

<u>3-azidopropanoyl-CoA (**3AZ-CoA**).</u> 0.03 mmol of DIPEA (5 equiv.) and 0.02 mmol of 3-azidopropanoic NHS ester (**9**, 1.5 equiv.) were added to an ice-cooled stirred solution of 0.006 mmol of coenzyme A (1 equiv.) in dry DMF (500 $\mu$L). The reaction mixture was then stirred for 5 min at room temperature. Completion of reaction was followed by LC/MS. The reaction was quenched by addition of $H_2O$ (5 mL) and lyophilized. The lyophilizate was then dissolved in $H_2O$, filtered, and subjected to two rounds of HPLC purification to obtain a pure final product **3AZ-CoA** as a white powder (1 mg, 1 $\mu$mol, 19%). The fractions were collected and lyophilized, and the dried product was stored at –20°C for subsequent experiments. Stock solution concentrations were calculated based on the isolated mass and final solution volume and confirmed by UV–Vis spectroscopy before use. $^1$H NMR (500 MHz, $D_2O$) δ 8.70 (s, 1H), 8.47 (s, 1H), 6.25 (d, $J$ = 6.0 Hz, 1H), 4.94 – 4.89 (m, 2H), 4.63 (s, 1H), 4.31 – 4.26 (m, 2H), 4.05 (s, 1H), 3.93 – 3.86 (m, 1H), 3.63 (t, $J$ = 6.0 Hz, 4H), 3.48 (t, $J$ = 6.9 Hz, 2H), 3.38 (t, $J$ = 6.4 Hz, 2H), 3.07 (t, $J$ = 6.9 Hz, 2H), 2.94 (t, $J$ = 6.5 Hz, 2H), 2.46 (t, $J$ = 6.9 Hz, 2H), 0.96 (s, 3H), 0.84 (s, 3H). $^{13}$C NMR (126 MHz, $D_2O$) δ 201.13, 174.74, 174.04, 149.99, 148.57, 144.69, 142.56, 118.61, 87.48, 83.62, 74.14, 71.89, 65.07, 46.69, 42.33, 38.48, 35.31, 28.15, 20.87, 18.16. $^{31}$P NMR (126 MHz, $D_2O$) δ -0.29, -10.89, -11.42. HRMS (ESI): calcd for $C_{24}H_{40}N_{10}O_{17}P_3S$ [M+H]$^+$ *m/z* 865.1501; found 865.1493.

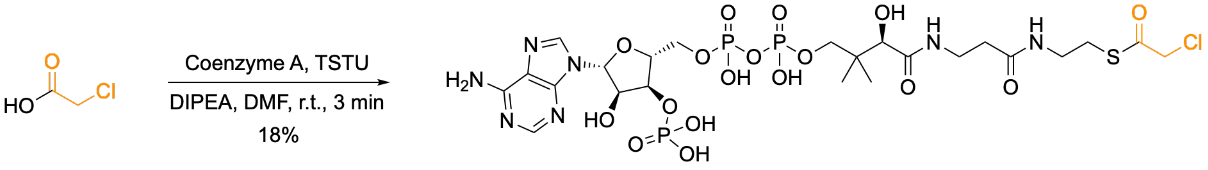

**ClAc-CoA**

**Scheme S6.** Chemical synthesis of ClAc-CoA. The acyl ligation handle is highlighted in orange.

Chloroacetyl-CoA (**ClAc-CoA**). A mixture of dry DMF (500 $\mu$L), 0.06 mmol of chloroacetic acid (3 equiv.), 0.09 mmol of DIPEA (5 equiv.) and 0.07 mmol of TSTU (3.6 equiv.) was stirred under an inert atmosphere for 15 min. The mixture was then added to a solution of 0.019 mmol of coenzyme A (1 equiv.) in dry DMF (500 $\mu$L). The reaction mixture was then stirred for 3 min at room temperature. Completion of reaction was followed by LC/MS. The reaction was quenched by addition of $H_2O$ (10 mL) and lyophilized. The lyophilizate was redissolved in $H_2O$, filtered, and subjected to two rounds of HPLC purification to obtain a pure final product **ClAc-CoA** as a white powder (2.9 mg, 3.4 $\mu$mol, 18%). The fractions were collected and lyophilized, and the dried product was stored at –20°C for subsequent experiments. Stock solution concentrations were calculated based on the isolated mass and final solution volume and confirmed by UV–Vis spectroscopy before use. $^{1}$H NMR (500 MHz, $D_2O$) δ 8.69 (s, 1H), 8.46 (s, 1H), 6.24 (d, $J$ = 5.3 Hz, 1H), 4.93 – 4.88 (m, 2H), 4.62 (s, 1H), 4.42 (s, 2H), 4.32 – 4.24 (m, 2H), 4.05 (s, 1H), 3.92 – 3.86 (m, 1H), 3.67 – 3.61 (m, 1H), 3.48 (t, $J$ = 13.2 Hz, 2H), 3.40 (t, $J$ = 13.1 Hz, 2H), 3.10 (t, $J$ = 12.8 Hz, 2H), 2.46 (t, $J$ = 13.7 Hz, 2H), 0.96 (s, 3H), 0.84 (s, 3H). $^{13}$C NMR (126 MHz, $D_2O$) δ 197.41, 174.73, 174.10, 149.92, 148.52, 144.67, 142.54, 118.59, 87.49, 83.48, 74.16, 73.89, 71.97, 65.12, 47.89, 38.26, 35.39, 28.66, 20.83, 18.23. $^{31}$P NMR (126 MHz, $D_2O$) δ -0.42, -11.04, -11.50. HRMS (ESI): calcd for $C_{23}H_{38}ClN_7O_{17}P_3S$ ([M+H]$^+$) $m/z$ 844.0941; found 844.0932.

#### Site-directed mutagenesis

Site-directed mutagenesis was performed using the Q5 Site-Directed Mutagenesis Kit (NEB). A plasmid encoding the catalytic domain of *Hs*ATAT1 (residues 2–236) fused to a GST tag (pGEX-GST-ATAT1 (2–236)) was used as the template to introduce the selected Ala, Gly, or Asn substitutions. The forward and reverse primers used to generate the single-point mutants are listed in Table S1 and were synthesized by Microsynth AG. Polymerase chain reaction (PCR) amplification was performed as follows: initial denaturation of plasmid DNA at 98 °C for 30 s, followed by annealing at 56–68 °C for 30 s depending on the melting temperature of the respective primers, and extension at 72 °C for 3 min. The amplification cycle was repeated 25 times. The PCR products were then subjected to Kinase, Ligase & DpnI (KLD) treatment at room temperature for 5 min and analyzed by 1% agarose gel electrophoresis. The treated PCR products were used to transform *E. coli* DH5α competent cells. Transformed cells were grown overnight at 37 °C in LB medium supplemented with ampicillin (100 μg/mL). Plasmids were isolated using the QIAprep Spin Miniprep Kit (QIAGEN). All mutations were confirmed by DNA sequencing performed by Genesupport SA Life Sciences (Fasteris).

**Protein expression and purification**

*Hs*ATAT1 (WT or mutants; residues 2–236) was expressed in *E. coli* BL21(DE3)pLysS cells as a GST fusion protein using the pGEX-GST-ATAT1(2–236) plasmid. Protein expression was induced with 0.2 mM IPTG for 16 h at 16 °C. Bacterial pellets were resuspended in lysis buffer containing 50 mM Tris (pH 7.5), 250 mM NaCl, 2 mM $MgCl_2$, and 5–10% glycerol, supplemented with 1 mM PMSF, protease inhibitor cocktail (Roche), and 0.25 mg/mL lysozyme, and subsequently lysed by sonication. Cell lysates were clarified by centrifugation at 17,000 rpm for 45 min at 4 °C. Proteins were purified using Glutathione Sepharose 4B resin (Cytiva). After overnight cleavage with PreScission protease (GE Healthcare) at 4 °C, proteins were eluted using wash/elution buffer containing 50 mM Tris (pH 7.6), 250 mM NaCl, 1 mM tris(2-carboxyethyl)phosphine (TCEP), and 5–10% glycerol. To assess rescue of enzyme stability by Ac-CoA, 350 μM Ac-CoA was included during the cleavage incubation. Eluted fractions were analyzed by 4–15% SDS-PAGE. Fractions containing the purified protein were pooled and concentrated using Amicon Ultra 10K centrifugal filters (Millipore). The concentrated protein was dialyzed overnight against dialysis buffer (50 mM Tris, pH 7.6, 250 mM NaCl, 1 mM TCEP, and 5–10% glycerol). Protein concentration was determined using the Pierce 660 nm Protein Assay. Purified enzyme was aliquoted into single-use aliquots, flash-frozen in liquid nitrogen, and stored at –80 °C until use.

The HDAC6 sequence from HDAC6-Flag (Addgene #13823) was subcloned into a pFastBacHTA vector using PluTI/NotI restriction enzymes for baculovirus generation and subsequent Sf9 cell infection. HDAC6 was purified from insect cells by resuspending the cell pellet in lysis buffer containing 100 mM Tris (pH 8.0), 10 mM NaCl, 5 mM KCl, 2 mM $MgCl_2$, 10% glycerol, and 0.2% Triton X-100, supplemented with protease inhibitor cocktail. Following cell lysis, the NaCl concentration was adjusted to 150 mM, and the lysate was clarified by ultracentrifugation at 50,000 rpm for 20 min at 4 °C. The supernatant was loaded onto a 1 mL HisTrap HP affinity column. After washing with buffer containing 25 mM imidazole, HDAC6 was eluted with buffer containing 250 mM imidazole over five column volumes. The eluate was further purified by size-exclusion chromatography using the lysis buffer. Pooled fractions were concentrated, and protein concentration was determined using the Pierce 660 nm Protein Assay. Purified HDAC6 was supplemented with additional glycerol to a final concentration of 10%, flash-frozen in liquid nitrogen, and stored at –80 °C until use.

**Tubulin purification and labeling**

Tubulin was purified from bovine brain by two cycles of polymerization and depolymerization in high-molarity PIPES buffer as previously described.[2] Briefly, the first polymerization/depolymerization cycle was performed using high-molarity PIPES buffer (1 M PIPES, pH 6.9, 10 mM $MgCl_2$, 20 mM EGTA, 1.5

mM ATP, and 0.5 mM GTP) supplemented with an equal volume of glycerol for polymerization, and depolymerization buffer (50 mM MES, pH 6.6, 1 mM $CaCl_2$) for depolymerization. A second polymerization/depolymerization cycle was performed using high-molarity PIPES buffer for polymerization and 0.25× BRB80 buffer for depolymerization, followed after 15 min by the addition of 5× BRB80 buffer to obtain a final concentration of 1× BRB80 buffer (80 mM PIPES, pH 6.8, 1 mM $MgCl_2$, and 1 mM EGTA).

Purified tubulin was either used unlabeled (black tubulin) or labeled with ATTO-488 (black/green tubulin), ATTO-565 (black/red tubulin) (ATTO-TEC GmbH), or biotin (biotinylated tubulin) following an adapted version of a previously published protocol.[3] Briefly, tubulin was polymerized in glycerol PB buffer (80 mM PIPES, pH 6.8, 5 mM $MgCl_2$, 1 mM EGTA, 1 mM GTP, and 33% glycerol) for 30 min at 37 °C and layered onto cushions containing 0.1 M NaHEPES (pH 8.6), 1 mM $MgCl_2$, 1 mM EGTA, and 60% glycerol, followed by centrifugation. The resulting pellet was resuspended in resuspension buffer (0.1 M NaHEPES, pH 8.6, 1 mM $MgCl_2$, 1 mM EGTA, and 40% glycerol) and incubated for 10 min at 37 °C with one-tenth of the reaction volume of 100 mM ATTO-488- or ATTO-565-NHS ester, or for 20 min at 37 °C with 2 mM NHS-biotin. Labeled tubulin was sedimented through cushions of 1× BRB80 buffer supplemented with 60% glycerol, resuspended in BRB80 buffer, and subjected to a second polymerization/depolymerization cycle. The labeling ratios were 11% for ATTO-488 and 15% for ATTO-565.

**LC–MS-based ATAT1 acetylation and acylation activity assays**

Reaction mixtures containing 1× acetylation buffer (40 mM PIPES, pH 6.9, 0.8 mM EGTA, 0.4 mM $MgS_4$, and 30% glycerol), 0.5, 1 or 1.5 mM Ac-CoA (Cayman Chemical) or Ac-CoA mimics, and 100 or 375 $\mu$M α-tubulin peptide substrate (p4, p10, or p11) were prepared. *Hs*ATAT1(2–236) (WT or mutant) was added to a final concentration of 50 or 75 $\mu$M, and the reactions were incubated at either 24 or 37 °C for 0, 24, 48, 72, 96, 120, 144, 168, or 192 h, as indicated in the corresponding figures. A control without added cofactor was included in each experiment to assess baseline acetylation arising from co-purifying Ac-CoA. Where indicated, reactions were supplemented with 1.5 mM ClAc-CoA every 24 h. Following incubation, aliquots were diluted with 2.4 volumes of 0.1% formic acid in $ddH_2O$ and analyzed by LC–MS. Peptides containing a single acetyl or acyl modification eluted later than the corresponding unmodified substrates because of their increased hydrophobicity, enabling chromatographic separation of substrate and product.

Remaining peptide substrate and modified product at each time point were quantified by integration of the corresponding peaks in the UV chromatograms recorded with the PDA detector. Relative product formation was calculated as the percentage of the total peptide signal corresponding to the modified product, according to the following equation:

$$Product\ (\%) = \frac{Peak\ area_{product}}{Peak\ area_{product} + Peak\ area_{substrate}} \cdot 100$$

Cofactor stability was quantified by integration of the UV absorbance peak corresponding to Ac-CoA or the respective Ac-CoA mimic relative to the peptide substrate peak. Relative cofactor abundance was calculated as the cofactor-to-substrate peak area ratio and normalized either to the initial time point (0 h = 1) or to the Ac-CoA control, as indicated.

The mass-to-charge ratios (m/z) of the peptide substrates, modified peptides, and cofactors were determined by ESI-MS. Data were processed using Xcalibur™ software.

**Fluorescence polarization assay**

A solution containing 50 nM p11-CoA-TAMRA was titrated with increasing concentrations of *Hs*ATAT1(2–236) (WT or mutant) dissolved in FP buffer (50 mM Tris pH 7.5, 250 mM NaCl, 1 mM TCEP, and 1% DMSO) in a black, flat-bottom 96-well plate (Greiner Bio-One). The plate was incubated for 1 h at room temperature prior to fluorescence polarization measurements. All measurements were performed in triplicate. Fluorescence polarization was measured using a plate reader (Tecan Spark® 20M) with an excitation wavelength of 531 nm (20 nm bandwidth) and an emission wavelength of 580 nm (25 nm bandwidth). The measured polarization values (mP) were plotted against the protein concentration, and the apparent $K_d$ of p11-CoA-TAMRA was determined by fitting the data to the following equation:[4,5]

$$FP = FP_{min} + \frac{(FP_{max} - FP_{min}) \cdot \left(C_t + Kd + C_p - \sqrt{-4 \cdot C_t \cdot C_p + \left(C_t + Kd + C_p\right)^2}\right)}{2 \cdot C_t}$$

where $F_{min}$ is the minimum polarization value, $F_{max}$ is the maximum polarization value, $C_t$ is the concentration of the tracer (p11-CoA-TAMRA), and $C_p$ is the concentration of *Hs*ATAT1(2–236) (WT or mutant).

In the competition assay, 50 nM p11-CoA-TAMRA and 500 nM *Hs*ATAT1(2–236) (WT or mutant) were used. The tracer–protein solution was titrated with increasing concentrations of the synthesized peptides, peptide-CoA conjugates, or Ac-CoA mimics. Plates were incubated for 1 h at room temperature, and fluorescence polarization measurements were performed under the same conditions as described above. The measured mP values were converted to the concentration of free protein, which was then plotted against the concentration of the competing ligand. The resulting data were fitted to a single-site binding isotherm to determine the $K_d$ of the competing ligand using the following equation:

$$logEC_{50} = \log\left(10^{logKi} \cdot \left(\frac{HotNM}{HotKdNM} + 1\right)\right)$$

$$Y = Bottom + \frac{Top - Bottom}{10^{-LogEC_{50}+X} + 1}$$

where *X* is the base-10 logarithm of the molar concentration of the competing (unlabeled) ligand, *Y* is the fluorescence polarization signal (mP), *Top* is the upper plateau of the fitted binding curve (maximum fluorescence polarization), *Bottom* is the lower plateau of the fitted binding curve (minimum fluorescence polarization), *logEC₅₀* is the base-10 logarithm of the competitor concentration resulting in 50% displacement under the assay conditions, $logK_i$ is the base-10 logarithm of the inhibition constant ($K_i$) of the competing ligand, *HotNM* is the concentration of tracer used in the competition assay (nM), and *HotKdNM* is the equilibrium dissociation constant ($K_d$) of the tracer–protein interaction (nM).

**SDS-PAGE, in-gel fluorescence scanning, and western blotting**

Samples were diluted to 1× SDS sample buffer (50 mM Tris-HCl, pH 6.8, 8% v/v glycerol, 2% w/v SDS, 100 mM DTT, and 0.1 mg/mL bromophenol blue), boiled, and loaded onto 4–15% or 8–16% Criterion TGX Stain-Free gels (Bio-Rad). Electrophoresis was performed at 200 V in 1× Tris/Glycine/SDS buffer (Bio-Rad).

For in-gel fluorescence experiments, fluorescently labeled proteins were visualized directly in the gel using a Fusion FX Gel Doc imaging system (Vilber). Following electrophoresis, gels were transferred to the imaging tray and scanned for 5–15 s using the appropriate fluorescence acquisition program, with excitation and emission settings optimized for each fluorophore. Images were acquired using identical acquisition parameters for all samples within each experiment. Following fluorescence imaging, gels were either stained with Coomassie Brilliant Blue to assess total protein loading or transferred onto PVDF membranes for subsequent western blot (WB) analysis, as described below.

For non-fluorescent experiments, gels were irradiated for 1 min and imaged using the stain-free imaging mode before being transferred onto PVDF membranes with iBlot™ 3 Transfer Stacks (Thermo Fisher Scientific) using an iBlot™ 3 Western Blot Transfer Device (Thermo Fisher Scientific). Membranes were blocked with 5% milk in TBST (0.1% Tween-20 in Tris-buffered saline) for 1 h. Following blocking, membranes were incubated with a primary antibody against α-tubulin (mouse monoclonal anti-TUBA4A (TUBA1), clone B-5-1-2, Sigma-Aldrich, T5168; 1:4000) diluted in blocking solution (5% milk in TBST) for 16 h at 4 °C. Membranes were washed with TBST (5 × 5 min), incubated with an HRP-conjugated secondary antibody (donkey anti-mouse IgG-HRP, Jackson ImmunoResearch Europe Ltd., 1:5000) diluted in blocking solution for 1 h at room temperature, washed again, and developed using Western BLoT Chemiluminescence HRP Substrate (Takara). Chemiluminescent signals were detected using a Fusion FX Gel Doc imaging system (Vilber). Membranes were stripped using Blue Clear SB antibody stripping solution (SERVA) for 45 min. The stripping conditions were validated to ensure efficient removal of the primary and secondary antibodies while maintaining protein detectability and allowing subsequent detection of acetylated tubulin. Following stripping, membranes were blocked for 1 h in blocking solution before incubation with a mouse monoclonal anti-acetylated tubulin antibody (clone 6-11B-1, Sigma-Aldrich, T7451; 1:5000) for 16 h at 4 °C. Following washing steps, membranes were incubated with the HRP-conjugated secondary antibody, developed using Western BLoT Chemiluminescence HRP Substrate (Takara), and imaged using the Fusion FX Gel Doc imaging system (Vilber).

Band intensities from fluorescence scans and WB were quantified using Fiji (ImageJ2). For fluorescence experiments, α-tubulin fluorescence signals were normalized to the corresponding α-tubulin loading signal determined by Coomassie staining or α-tubulin immunoblotting. For WB-based experiments, acetylated tubulin signals were normalized to total α-tubulin levels determined by α-tubulin immunoblotting or stain-free imaging. The resulting normalized values were subsequently expressed relative to the control condition within each experiment.

**Preparation of deacetylated tubulin and microtubules for enzymatic assays**

Deacetylated tubulin (deAcTub) was prepared by HDAC6 treatment of purified tubulin dimers prior to microtubule polymerization, based on the reported preference of HDAC6 for soluble tubulin dimers over assembled microtubules.[6]

To remove the acetyl modification from α-tubulin K40 (K40) and generate deAcTub, purified acetylated tubulin was incubated with recombinant HDAC6 at a 1:10 molar ratio of HDAC6:tubulin. Tubulin was diluted to 35 μM in 1× BRB80 buffer and incubated with recombinant HDAC6 (3.5 μM) at room temperature for 30 min, followed by incubation on ice for 10 min to prevent tubulin polymerization, and a final incubation at room temperature for 0.5–1 h. The reaction was terminated by addition of 100 μM Tubacin (HDAC6 inhibitor, Biorbyt). Deacetylation efficiency was assessed by WB using the anti-acetylated tubulin antibody.

GMPCPP-stabilized microtubules were polymerized by diluting deAcTub to 30 μM in 1× BRB80 buffer supplemented with 0.5 mM GMPCPP and incubating for 20 min at 37 °C. To increase microtubule length, two subsequent additions of 1 μM deAcTub were performed, each followed by incubation for 10 min at 37 °C. The polymerized deAcTub mixture was centrifuged at 12,700 rpm for 15 min at room temperature to pellet deacetylated microtubules (deAcMT). The supernatant containing soluble tubulin, excess GMPCPP, HDAC6, and Tubacin was discarded, and the microtubule pellet was resuspended in 1× BRB80 buffer. Deacetylated tubulin (deAcTub) and deacetylated microtubules (deAcMT) were used as substrates for enzymatic assays. Unless otherwise stated, the same microtubule polymerization conditions were used for all experiments.

**Western blot-based ATAT1 acetylation activity assays**

Reactions were performed in a final volume of 5 μL containing 10 μM deAcTub (derived from black/red tubulin), 100 μM Ac-CoA, and 2 μM *Hs*ATAT1(2–236) (WT or mutant) in 1× acetylation buffer. Unless otherwise indicated for time-course experiments, reactions were incubated at room temperature for 24 h. Reactions were quenched with SDS sample buffer and divided into two aliquots, which were loaded onto separate SDS–PAGE gels for WB analysis and Coomassie Brilliant Blue staining, respectively. Stain-free imaging was used to determine deAcTub loading levels, acetylated tubulin levels were determined by WB using the antibody conditions and quantification procedure described above, and Coomassie Brilliant Blue staining was used to determine ATAT1 levels. Relative ATAT1 acetylation activity was calculated by normalizing the acetylated tubulin/α-tubulin signal ratio to the corresponding ATAT1 level.

Statistical analyses were performed using GraphPad Prism. Acetylation activities of *Hs*ATAT1(2–236) mutants were compared with wild-type under each reaction condition using complete datasets (*N* = 3 independent experiments). A mixed-effects model (REML) followed by Dunnett's multiple comparisons test was used to compare each mutant with the WT-ATAT1 control under each reaction condition. Statistical analyses were performed on the raw acetylation/α-tubulin and ATAT1 measurements prior to normalization to avoid bias introduced by normalization. *P* values <0.05 were considered statistically significant.

**Competitive inhibition assays of ATAT1-mediated acetylation**

For tubulin substrates, 5 μL reactions containing 10 μM deAcTub (derived from black/red tubulin), 100 μM Ac-CoA, increasing concentrations of 6HY-CoA, and 2 μM *Hs*ATAT1(2–236) (WT) were prepared in 1× acetylation buffer. After incubation for 24 h at room temperature, reactions were quenched with SDS sample buffer and divided into two aliquots for Coomassie Brilliant Blue staining and WB analysis. Coomassie Brilliant Blue staining was used to determine ATAT1 levels, and acetylated tubulin levels were determined by WB using the same antibody conditions and quantification procedure described above. Activity was normalized to the control reaction containing Ac-CoA in the absence of 6HY-CoA.

For microtubule substrates, 5 μL reactions containing 10 μM deAcMT (derived from black/red tubulin) and a fixed concentration of Ac-CoA (10 μM) were assembled in 1× acetylation buffer, and increasing concentrations of p11-CoA or Ac-CoA mimics were added. Reactions were initiated by addition of 2 μM *Hs*ATAT1(2–236) (WT). After incubation for 1 h at room temperature, reactions were quenched with SDS sample buffer and divided into two aliquots for Coomassie Brilliant Blue staining and WB analysis. Stain-free imaging was used to determine deAcMT loading levels, Coomassie Brilliant Blue staining was used to determine ATAT1 levels, and acetylated tubulin levels were determined by WB using the same antibody conditions and quantification procedure described above. Activity was normalized to the control reaction containing Ac-CoA in the absence of p11-CoA or Ac-CoA mimics. A dose–response curve was fitted using GraphPad Prism, and the $pIC_{50}$ value was calculated.

**In-gel fluorescence-based ATAT1 acylation activity and competition assays**

For screening Ac-CoA mimics for ATAT1-mediated acylation, 10 μM deAcTub (derived from black tubulin) was incubated with 100 μM Ac-CoA or Ac-CoA mimic in the presence of 2 μM *Hs*ATAT1(2–236) (WT or L163A) in EGTA-free 1× acetylation buffer for 24 h at room temperature. Reactions were performed in a final volume of 5 μL. Following incubation, reactions were subjected to CuAAC click chemistry using conditions adapted from a previously reported protocol.[7] For reactions containing alkyne-functionalized Ac-CoA mimics, a click reaction mixture containing 100 μM L-cysteine, 100 μM TAMRA-azide, 3 mM TCEP, and 100 μM tris[(1-benzyl-4-triazolyl)methyl]amine (TBTA) in EGTA-free 1x acetylation buffer was added. For reactions containing azido-functionalized Ac-CoA mimics, a click reaction mixture containing 100 μM L-cysteine, 100 μM TAMRA-alkyne (Sigma-Aldrich), 3 mM TCEP, and 100 μM TBTA in EGTA-free 1x acetylation buffer was used. Click reactions were initiated by addition of 1 mM copper sulfate and incubated for 1–1.5 h in the dark at room temperature or 37 °C. Samples were separated on 8–16% SDS–PAGE gels and analyzed by in-gel fluorescence scanning as described above, using 532 nm excitation and a 580/30 nm band-pass emission filter. Coomassie Brilliant Blue staining was used to determine α-tubulin and ATAT1 levels. Tubulin acetylation levels were determined by WB using the same antibody conditions and quantification procedure described above.

For assessment of tubulin versus microtubule acylation, 10 μM deAcTub or deAcMT (derived from black tubulin) was incubated with 100 μM Ac-CoA, 4PY-CoA, or 3AZ-CoA in the presence of 2 μM *Hs*ATAT1(2–236) (WT or L163A) in EGTA-free 1× acetylation buffer for 1 h at room temperature. For time-course experiments using deAcMT, reactions were incubated for 5, 15, 60, or 240 min. At each time point, reactions were quenched by boiling and subsequently subjected to click chemistry using the same procedure described above.

For competition assays between Ac-CoA and Ac-CoA mimics, reactions containing 10 μM deAcMT (derived from black tubulin), a fixed concentration of Ac-CoA (10 μM), and increasing concentrations of 4PY-CoA or 3AZ-CoA were assembled in EGTA-free 1× acetylation buffer. For competition assays between 4PY-CoA and p11-CoA, reactions containing 10 μM deAcMT (derived from black tubulin), a fixed concentration of 4PY-CoA (10 μM), and increasing concentrations of p11-CoA were assembled in EGTA-free 1× acetylation buffer. Both sets of reactions were initiated by addition of 2 μM *Hs*ATAT1(2–236) (L163A) and incubated for 1 h at room temperature before being subjected to click chemistry using the same procedure described above. To assess the ability of the anti-acetylated tubulin antibody to recognize acyl-modified tubulin, an equivalent set of reactions was processed without the click chemistry step and analyzed by WB.

For all click chemistry–based experiments, reactions were quenched with SDS sample buffer and divided into two aliquots for Coomassie Brilliant Blue staining and WB analysis, and samples were subsequently processed and quantified as described above. For competition assays between 4PY-CoA and p11-CoA, quantification was presented as α-tubulin fluorescence relative to total α-tubulin, as determined by Coomassie staining, and normalized to ATAT1-L163A + 4PY-CoA without p11-CoA.

**HDAC6-mediated deacylation analysis**

Reactions containing 10 μM purified α-tubulin or polymerized microtubules (prepared from black tubulin) in a final volume of 5 μL were first incubated with 100 μM Ac-CoA mimic in the presence of 2 μM *Hs*ATAT1(2–236) (L163A) in EGTA-free 1× acetylation buffer for 1 h at room temperature. To assess HDAC6-mediated deacylation of labeled substrates, reactions were then treated with either 1 μM HDAC6, 1 μM HDAC6 in the presence of 100 μM Tubacin, or a no-HDAC6 control for 1.5 h at room temperature. Following treatment, reactions were subjected to click chemistry using the same procedure described above. Reactions were quenched with SDS sample buffer and divided into two aliquots for Coomassie Brilliant Blue staining and WB analysis. Samples were resolved on 8–16% SDS–PAGE gels and analyzed by in-gel fluorescence scanning as described above. Coomassie Brilliant Blue staining was used to determine ATAT1 levels. Acetylated tubulin levels were determined by WB using the same antibody conditions and quantification procedure described above.

**Site-specific mapping of α-tubulin K40 acylation in microtubules by LC–MS/MS**

To identify site-specific modifications at K40 of α-tubulin, 50 $\mu$L reactions containing deAcMT (10 $\mu$M, derived from black tubulin) alone, deAcMT with *Hs*ATAT1(2–236) (L163A, 2 $\mu$M), or deAcMT with *Hs*ATAT1(2–236) (L163A, 2 $\mu$M) and 4PY-CoA (100 $\mu$M) were prepared in duplicate in 1× acetylation buffer and incubated at room temperature for 24 h. Reactions were quenched by freezing at –20 °C. Samples were processed by the Mass Spectrometry Core Facility (ChemBio MS, University of Geneva, Switzerland) using the single-pot, solid-phase-enhanced sample preparation (SP3) protocol, including disulfide reduction, cysteine alkylation, on-bead proteolytic digestion, and desalting. Because modification of K40 prevents tryptic cleavage at this residue, digestion was performed with endoproteinase Glu-C (1:50 enzyme-to-substrate ratio) to generate peptides suitable for site-specific analysis.

Peptides were separated on an Easy-nLC 1000 system coupled online to an Orbitrap Fusion mass spectrometer using a 65 min method with a 30 min active gradient and analyzed in data-dependent acquisition (DDA) mode. An inclusion list encompassing the expected m/z values of the K40-containing peptide in its unmodified, acetylated (+42.0106 Da), and 4PY-acylated (+80.0269 Da) forms across 2+, 3+, and 4+ charge states was used to enhance detection of low-abundance modified species. Raw files were processed in MaxQuant (v2.7.3.0) against a custom FASTA database comprising bovine α- and β-tubulin isoforms, *Hs*ATAT1, and common contaminants. Methionine oxidation, N-terminal acetylation, lysine acetylation (+42.0106 Da), and 4PY-acylation (+80.0269 Da) were set as variable modifications. Protein identifications required at least one unique peptide match.

Relative occupancy of K40 modification states was calculated from the corresponding peptide intensity ratios. Because peptide ionization efficiencies may differ between modified and unmodified species, these values represent relative changes in K40 modification occupancy rather than absolute modification stoichiometry.

**Flow chamber preparation**

Glass slides and coverslips were cleaned by sonication in 1 M NaOH for 40 min, rinsed with ddH$_2$O, and subsequently sonicated in 96% ethanol for 40 min, followed by repeated rinsing with ddH$_2$O. Slides and coverslips were dried with compressed air and plasma-treated in a plasma cleaner (Diener Electronic, Plasma Surface Technology). Slides were functionalized by incubation for 2 days at room temperature with gentle agitation in triethoxysilane-PEG (1 mg/mL in 96% ethanol containing 0.02% HCl; Creative PEGWorks). Coverslips were functionalized under the same conditions using a 1:5

mixture of triethoxysilane-PEG-biotin and triethoxysilane-PEG. After functionalization, slides and coverslips were washed with 96% ethanol and $ddH_2O$, dried with compressed air, and stored in a sealed container at 4 °C. Flow chambers were assembled by attaching a functionalized coverslip to a functionalized glass slide with double-sided adhesive tape, yielding a chamber volume of approximately 15 $\mu$L.

**TIRF-based analysis of acylated immobilized microtubules**

Deacetylated microtubules (deAcMT) were prepared from deAcTub by mixing 70% ATTO-488-labeled tubulin with 30% biotinylated tubulin to a final tubulin concentration of 30 $\mu$M in 1× BRB80 buffer. Recombinant HDAC6 (3 $\mu$M) was added following the deacetylation procedure described above. For microtubule polymerization, 5 $\mu$M deAcTub was supplemented with 0.5 mM GMPCPP and 1 $\mu$M paclitaxel (Taxol) in 1× BRB80 buffer and incubated using the same polymerization and elongation procedure described above. The resulting microtubule pellet was resuspended in 1× BRB80 buffer supplemented with 10 $\mu$M Taxol to obtain a final deAcMT concentration of 7 $\mu$M.

For acylation reactions in solution, 5 $\mu$M deAcMT was incubated with 100 $\mu$M 4PY-CoA in EGTA-free 2× acetylation buffer in the presence or absence of 1 $\mu$M *Hs*ATAT1(2–236) (L163A) for 1 h at room temperature before transfer to a flow chamber. For acylation reactions in the flow chamber, a solution containing 5 $\mu$M deAcMT in EGTA-free 2× acetylation buffer was prepared.

Flow chambers were first incubated with 20 $\mu$L neutravidin (50 $\mu$g/mL; Thermo Fisher Scientific) for 5 min and then washed with 20 $\mu$L EGTA-free 1× acetylation buffer. Microtubules (10 $\mu$L) were diluted into 90 $\mu$L EGTA-free 1× acetylation buffer and introduced into the flow chamber in five sequential 20 $\mu$L additions. The first addition was incubated for 10 min, whereas subsequent additions were incubated for 1 min each. Unbound microtubules were removed by washing with 20 $\mu$L EGTA-free 1× acetylation buffer.

For acylation reactions in the flow chamber, a solution containing 500 $\mu$M 4PY-CoA, 6HY-CoA or Ac-CoA in the presence or absence of 5 $\mu$M *Hs*ATAT1(2–236) (WT or L163A) in EGTA-free 1× acetylation buffer was introduced into the chamber and incubated for 1 h at 37 °C. For competition between 4PY-CoA and p11-CoA, a solution containing 500 $\mu$M 4PY-CoA, 5 $\mu$M *Hs*ATAT1(2–236) (L163A), and with or without 1 mM p11-CoA in EGTA-free 1× acetylation buffer was added to the flow chamber and incubated for 1 h at room temperature.

Following acylation either in solution (followed by transfer to the flow chamber and microtubule immobilization) or directly in the flow chamber, immobilized microtubules were washed and subjected to click chemistry directly in the flow chamber. A click reaction mixture containing 100 $\mu$M L-cysteine, 100 $\mu$M CalFluor647-azide (Vector Labs), 3 mM TCEP, 100 $\mu$M TBTA, and 1 mM copper sulfate in EGTA-free 1× acetylation buffer was introduced into the chamber and incubated for 30 min in the dark at 37 °C. The click reaction mixture was removed by several washes with EGTA-free 1× acetylation buffer before imaging by total internal reflection fluorescence (TIRF) microscopy.

For samples subjected to click reaction in solution, acylated microtubules were labeled by CuAAC click chemistry directly in the reaction mixture using the procedure described above before transfer to the flow chamber for immobilization and TIRF microscopy imaging.

**TIRF microscopy imaging and image analysis**

Control and experimental conditions were processed in parallel, and microtubules were randomly selected during image acquisition. Images were acquired using an Axio Observer inverted TIRF microscope (Zeiss, 3i) equipped with a Prime 95B camera (Photometrics). Imaging was performed using a Plan-Apochromat 100×/1.46 Oil DIC (UV) VIS-IR objective (Zeiss). Image acquisition was controlled using SlideBook 6X64 software. Temperature during imaging was maintained using a Tokai Hit stage-top incubator (STXG-SP400NX-SET). Image analysis was performed using Fiji (ImageJ2).

For visualization of acylation incorporation along microtubules, fluorescence intensity line profiles were generated in Fiji (ImageJ2) following background subtraction. Line profiles of the CalFluor647 fluorescence signal (640 nm channel) and the corresponding ATTO-488-labeled microtubules (488 nm channel) were extracted along the regions indicated in the figure insets. The ATTO-488 and CalFluor647 fluorescence intensity profiles were overlaid after aligning each dataset to its respective minimum intensity value.

To quantify the association between the ATTO-488 and CalFluor647 fluorescence signals on immobilized microtubules, colocalization analysis was performed using the JACoP plugin in Fiji (ImageJ2). Multiple regions of interest (ROIs) were analyzed from the positive condition of each experiment together with ROIs from the corresponding negative control conditions. Van Steensel cross-correlation function (CCF) analysis was performed, and three complementary colocalization metrics were extracted: Pearson's correlation coefficient above the Costes-determined threshold ($r_{Costes}$), Costes-thresholded Manders' overlap coefficients (M1 and M2), and Li's Intensity Correlation Quotient (ICQ).

For competition assays between 4PY-CoA and p11-CoA, fluorescence intensities in the ATTO-488 and CalFluor647 channels were measured for individual microtubules. Regions of interest (ROIs) were defined using the polygon selection tool in Fiji (ImageJ2). Acylation levels were quantified as the ratio of CalFluor647 to ATTO-488 fluorescence intensity and compared between conditions in the absence or presence of p11-CoA.

**Analysis of incorporation of acylated tubulin into stable microtubules**

Reactions (10 $\mu$L) containing deAcTub (10 $\mu$M), Ac-CoA or 4PY-CoA (100 $\mu$M), and *Hs*ATAT1(2–236) (WT or L163A, as indicated, 2 $\mu$M) in EGTA-free 2× acetylation buffer were incubated for 4 h at room temperature to generate acetylated or acylated tubulin, respectively. As an inhibition control, ATAT1 was pre-incubated with 250 $\mu$M p11-CoA for 30 min prior to addition of deAcTub and cofactor. For the remaining reactions, 250 $\mu$M p11-CoA was added only after the 4 h modification reaction to inhibit ATAT1 activity during subsequent microtubule polymerization. A non-inhibited control was performed without p11-CoA addition. Modified tubulin samples were subsequently polymerized into GMPCPP- and Taxol-stable microtubules using the polymerization procedure described above. Following centrifugation, the microtubule pellet and soluble tubulin supernatant were separated and retained for subsequent click chemistry analysis. The microtubule pellet was resuspended in 1× acetylation buffer and divided into separate aliquots. One aliquot was subjected to CuAAC click chemistry with TAMRA-azide using the conditions described above, followed by SDS–PAGE, in-gel fluorescence scanning, and WB analysis. A second aliquot was analyzed by TIRF microscopy under the imaging conditions described above, followed by in-chamber click reaction with CalFluor647-azide.

Quantification of gel/blot was presented as α-tubulin fluorescence or acetylated α-tubulin relative to total α-tubulin, with fluorescence data normalized to the microtubule fraction + ATAT1-L163A + 4PY-CoA without p11-CoA and WB data normalized to the microtubule fraction + WT-ATAT1 + Ac-CoA without p11-CoA.

### NMR SPECTRA

**[1]H (top), [13]C (middle) and [31]P (bottom) NMR spectra of 3BY-CoA.**

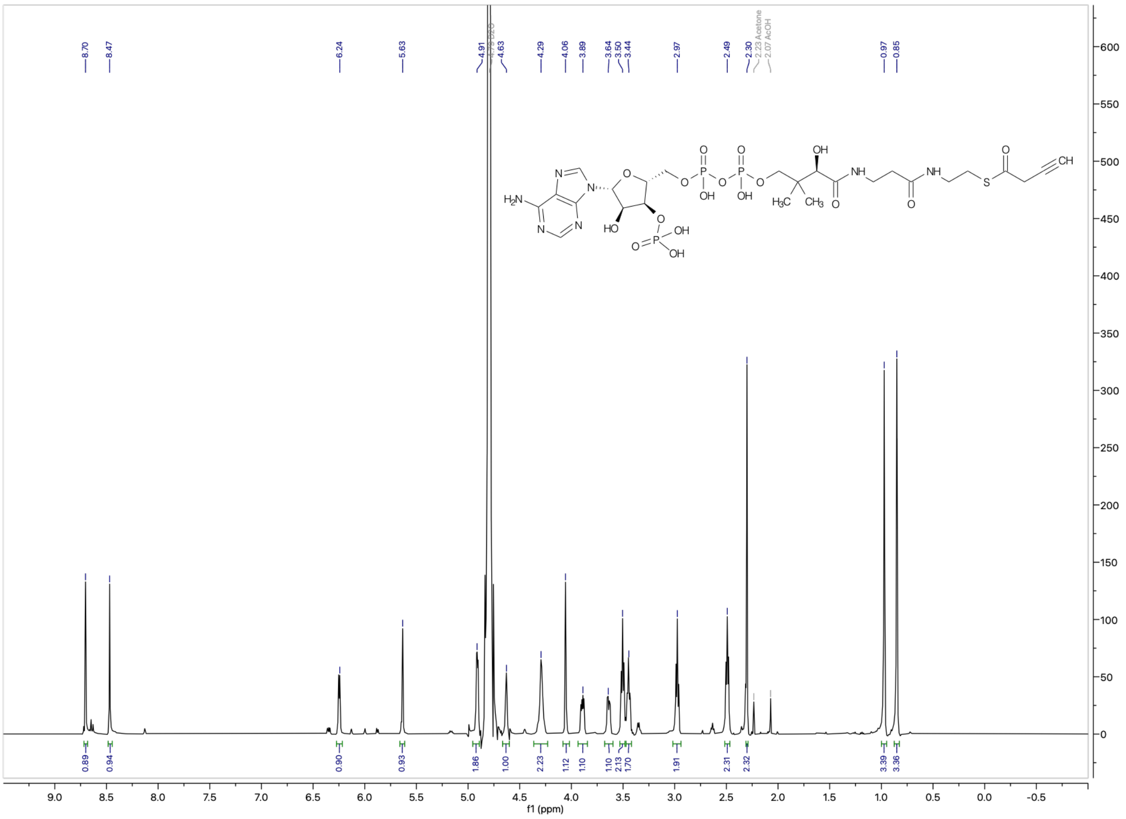

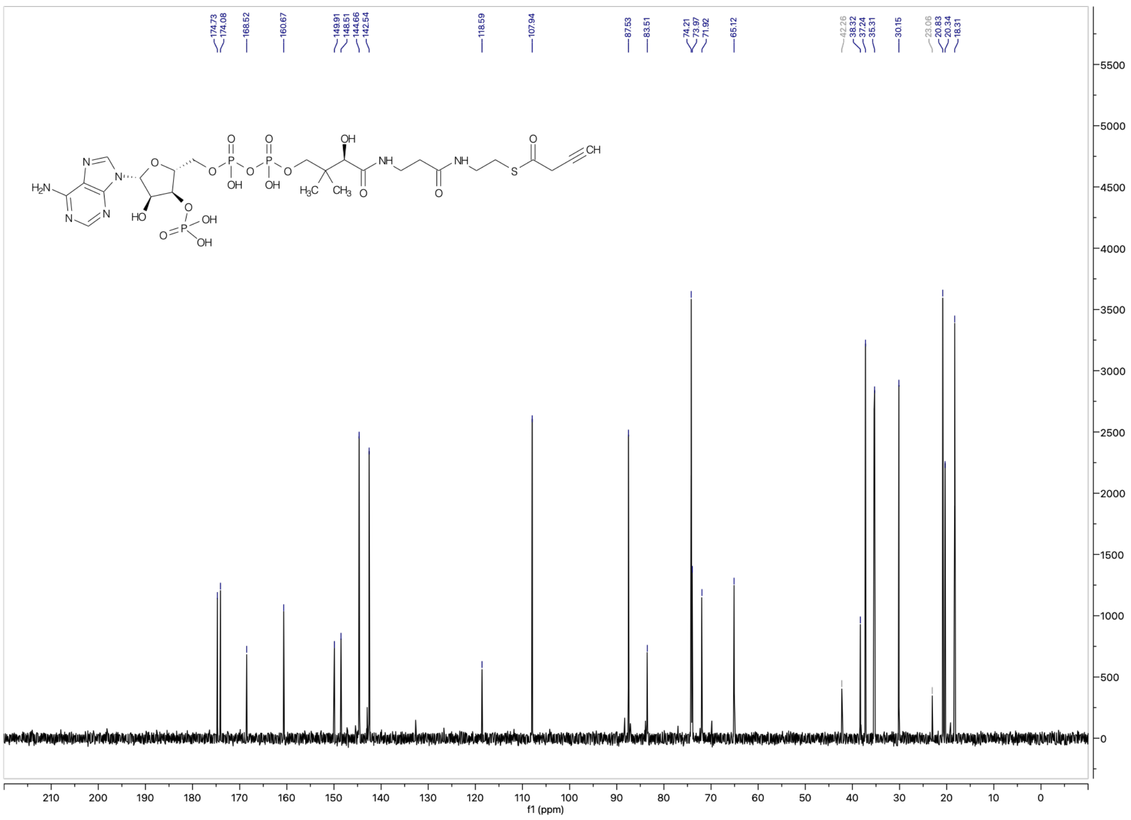
174.73
174.08
168.52
160.67
149.91
148.51
144.66
142.54
118.59
107.94
87.53
83.51
74.21
73.97
71.92
65.12
42.26
38.32
37.24
35.31
30.15
23.06
20.83
20.34
18.31
f1 (ppm)

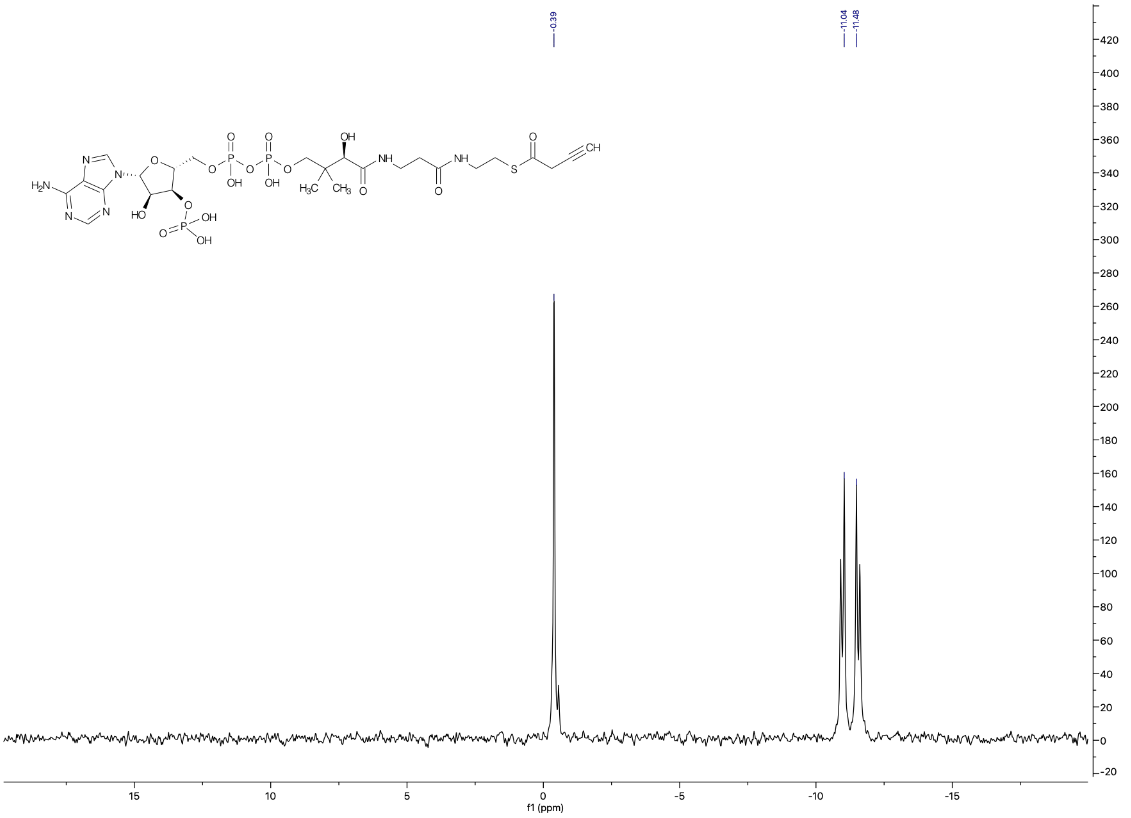
-0.39
-11.04
-11.48
f1 (ppm)

**$^{1}$H (top), $^{13}$C (middle) and $^{31}$P (bottom) NMR spectra of 4PY-CoA.**

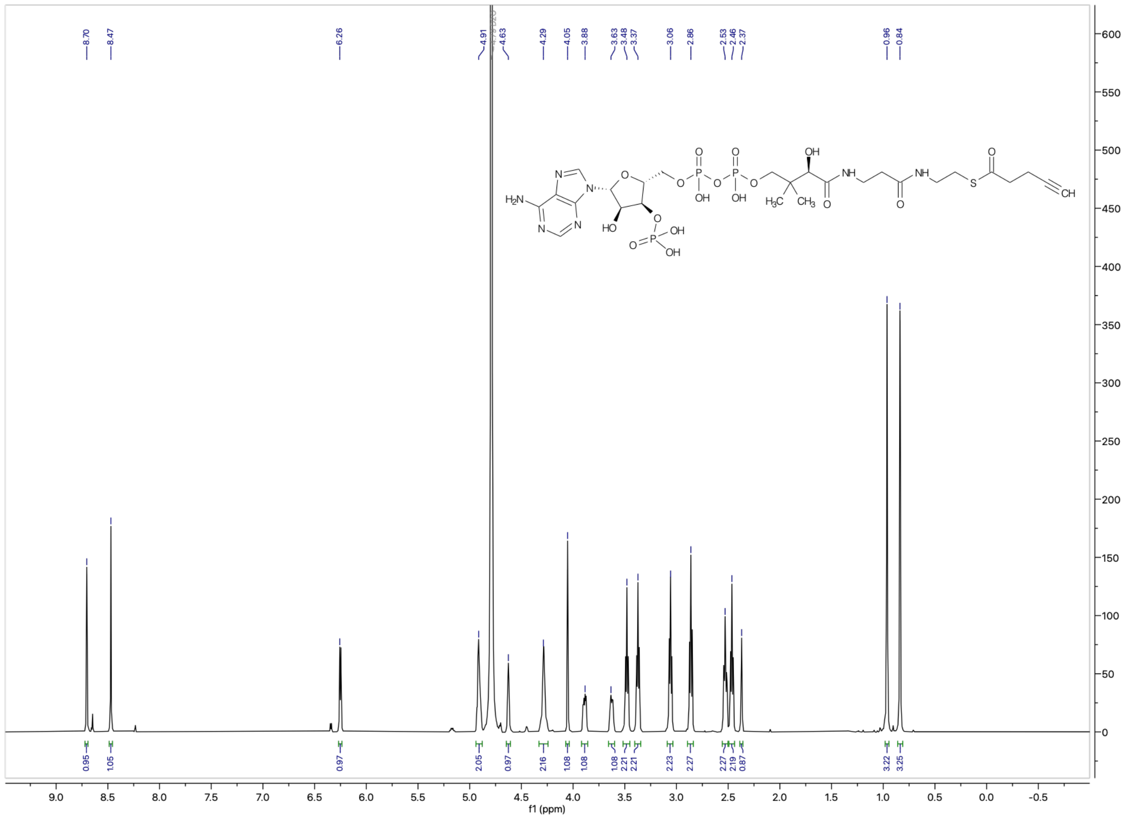

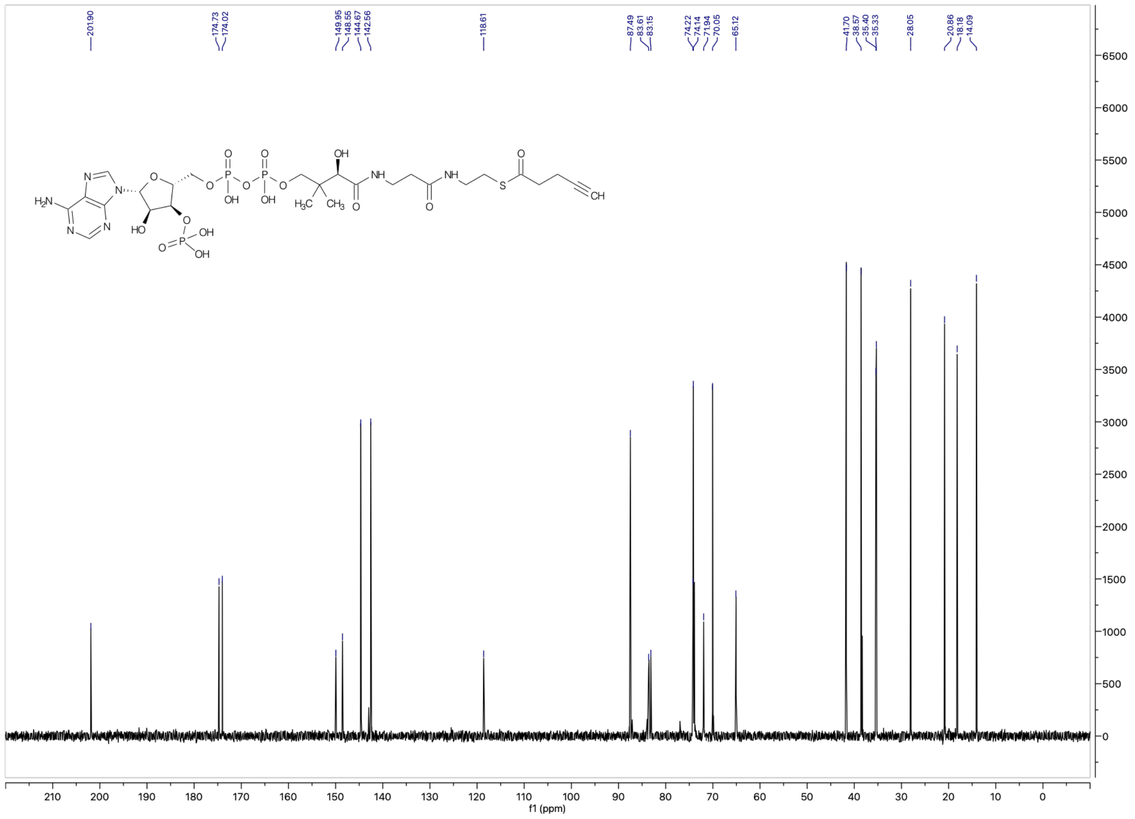
201.90
174.73
174.02
149.95
148.55
144.67
142.56
118.61
87.49
83.61
83.15
74.22
74.14
71.94
70.05
65.12
41.70
38.57
35.40
35.33
28.05
20.86
18.18
14.09
$H_2N$
N
O
OH
HO
O
P
NH
S
CH
$H_3C$
$CH_3$
f1 (ppm)

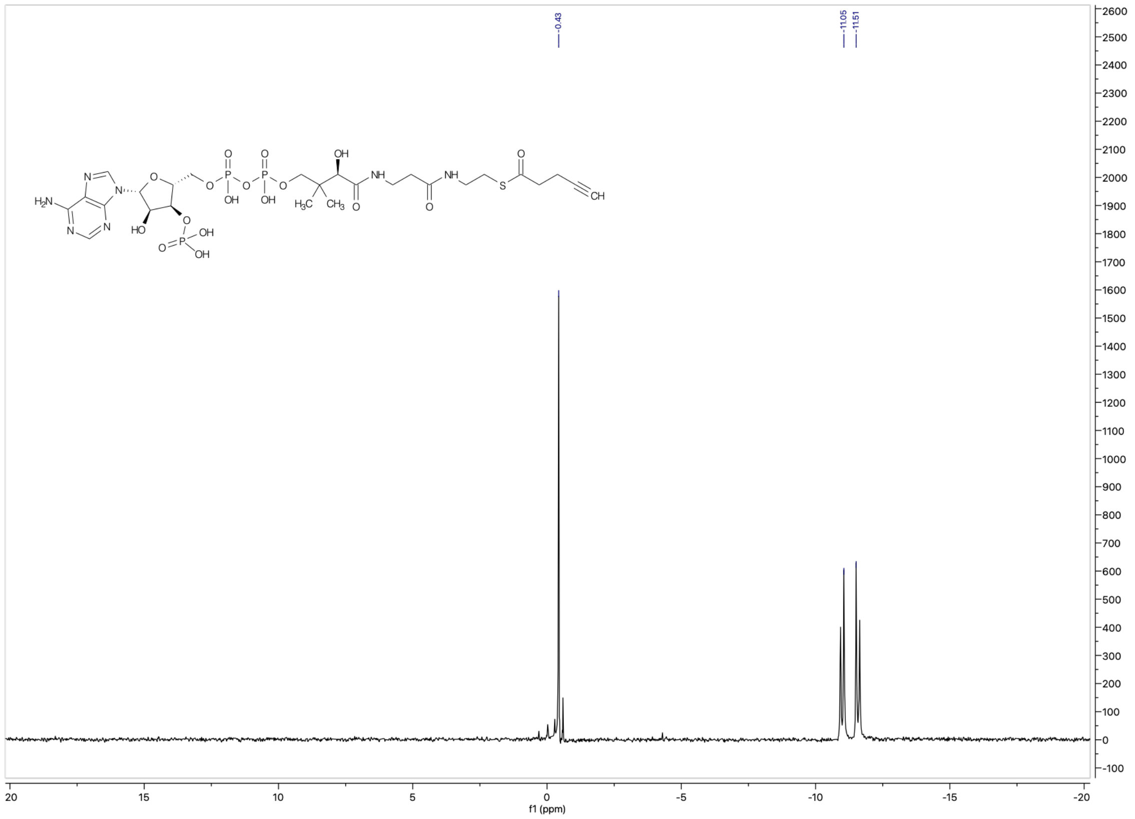
-0.43
-11.05
-11.51
$H_2N$
N
O
OH
HO
O
P
NH
S
CH
$H_3C$
$CH_3$
f1 (ppm)

**$^{1}H$ (top), $^{13}C$ (middle) and $^{31}P$ (bottom) NMR spectra of 5HY-CoA.**

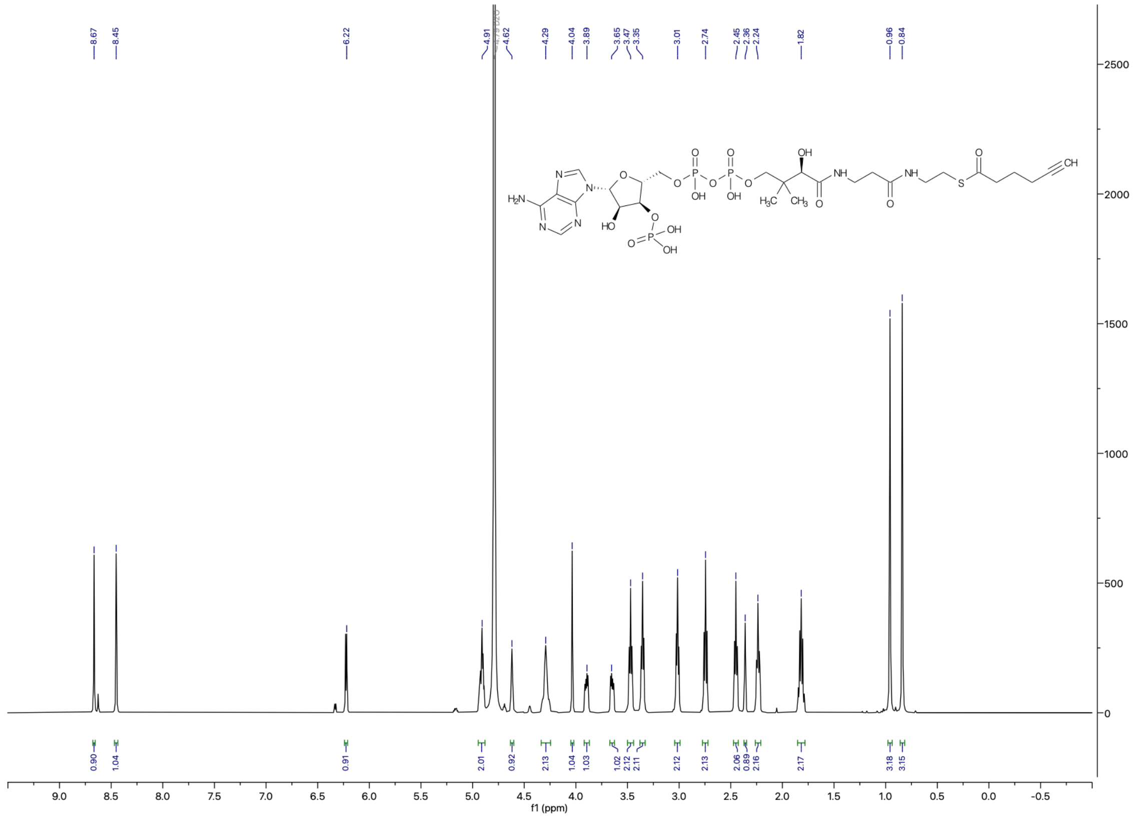

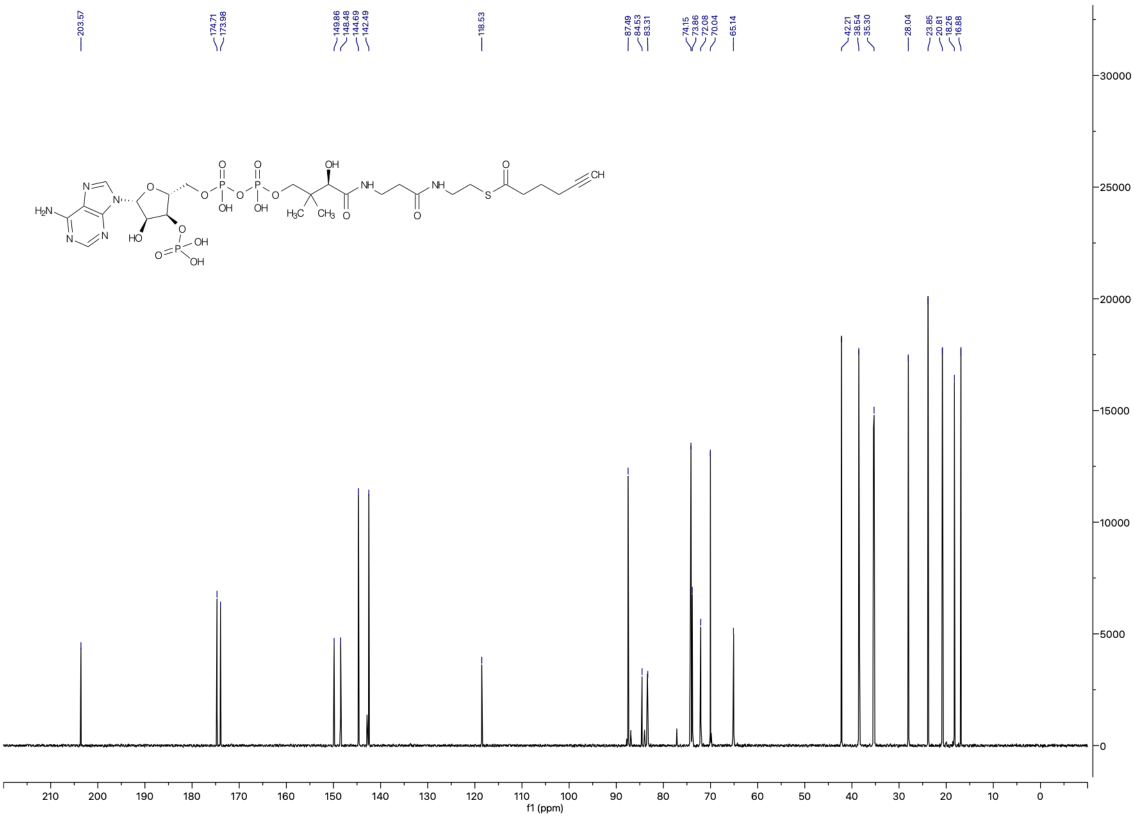

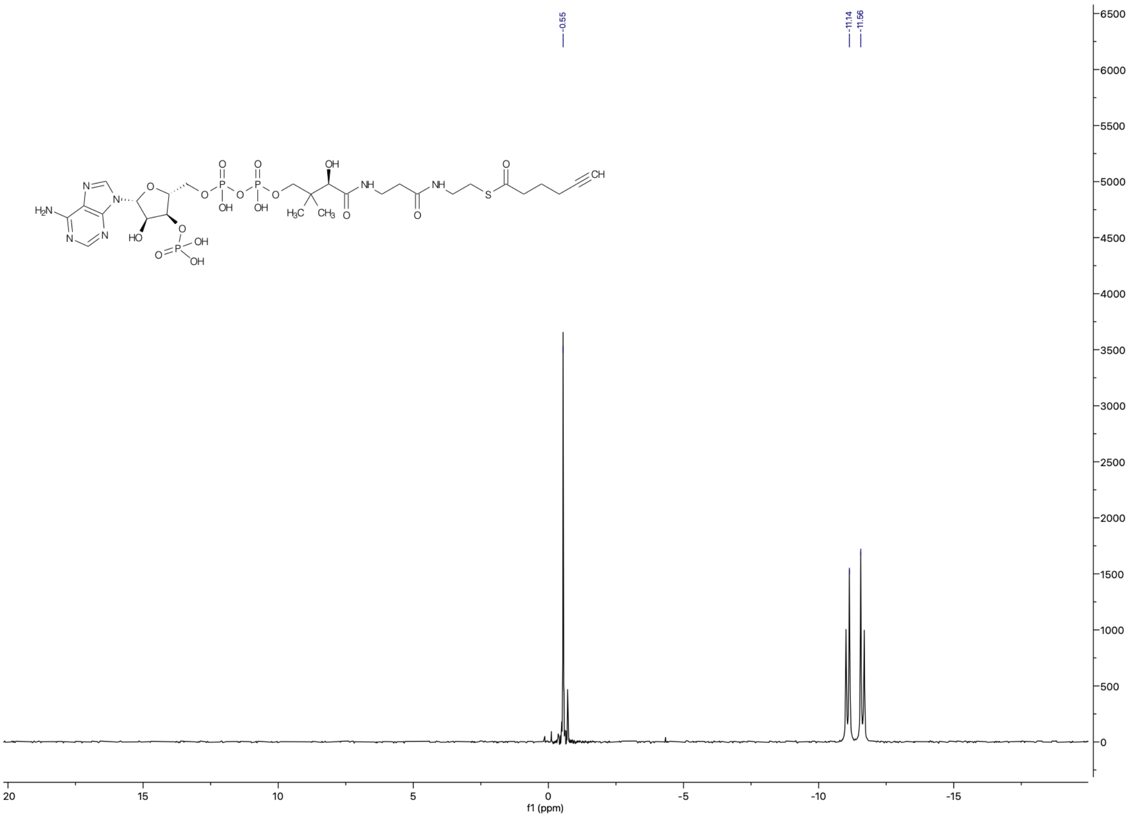

**[1]H (top), [13]C (middle) and [31]P (bottom) NMR spectra of 6HY-CoA.**

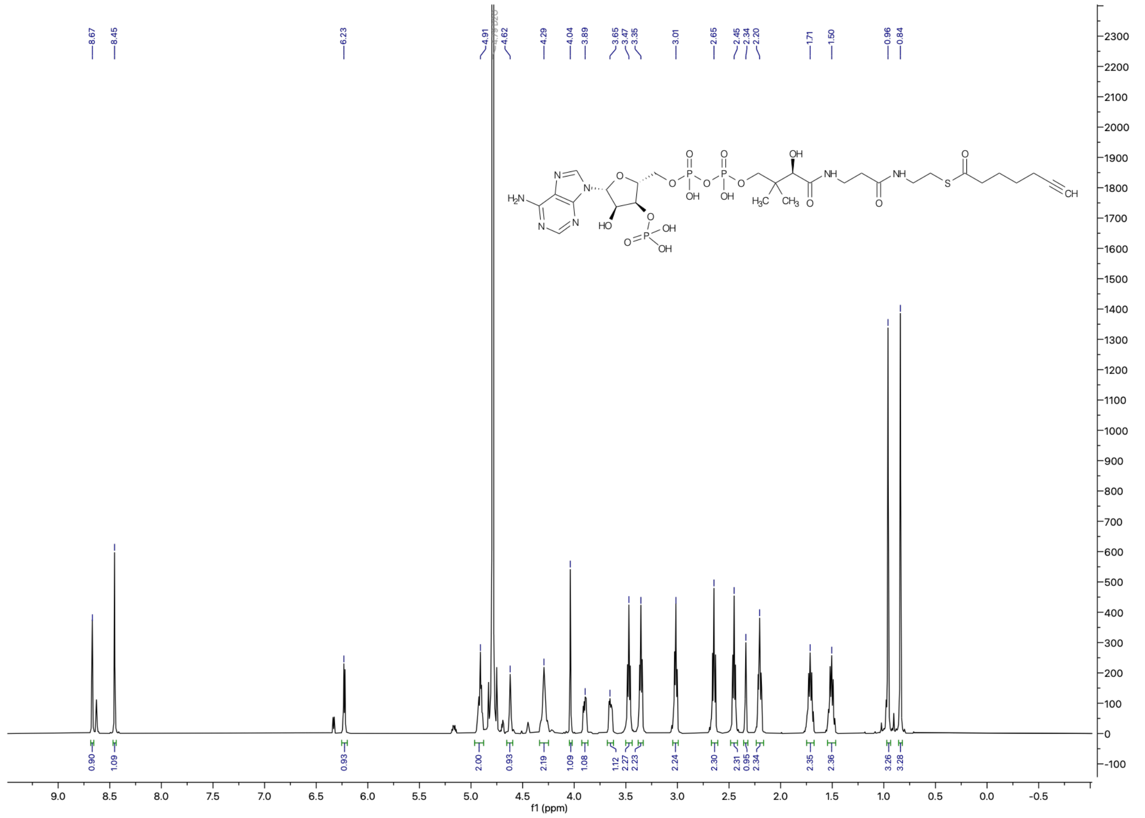

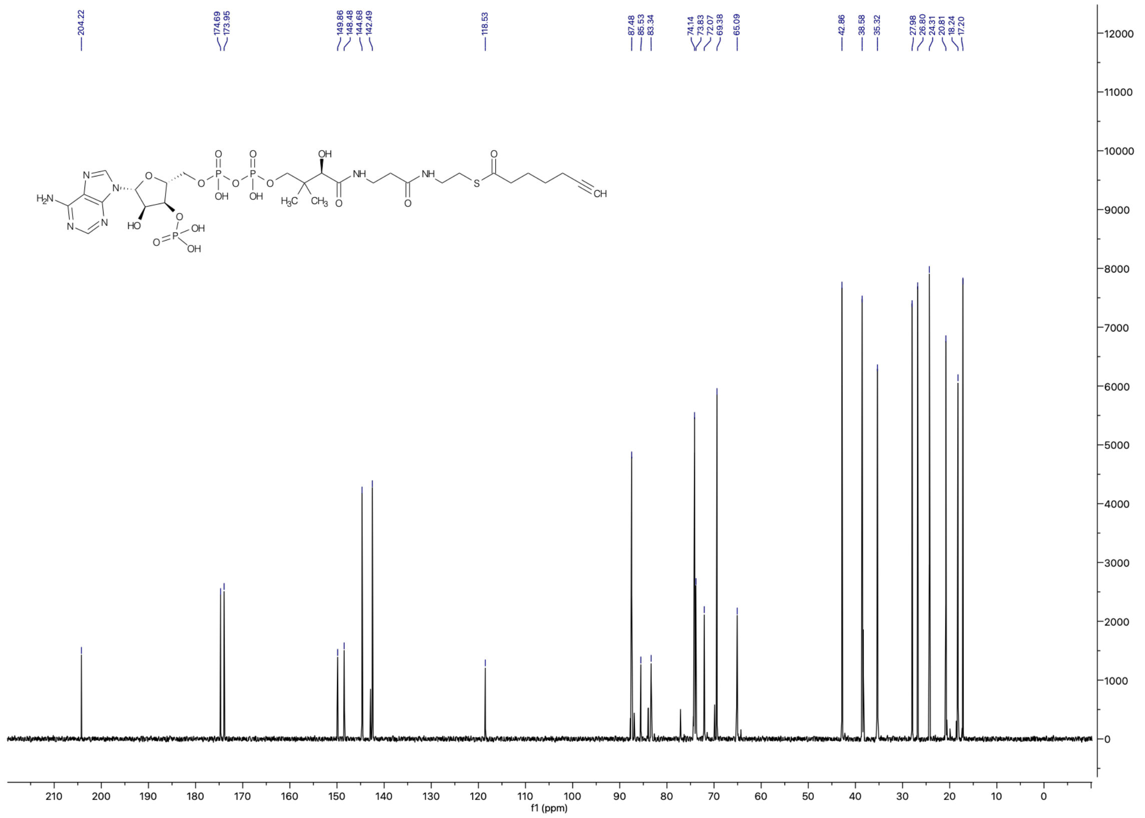

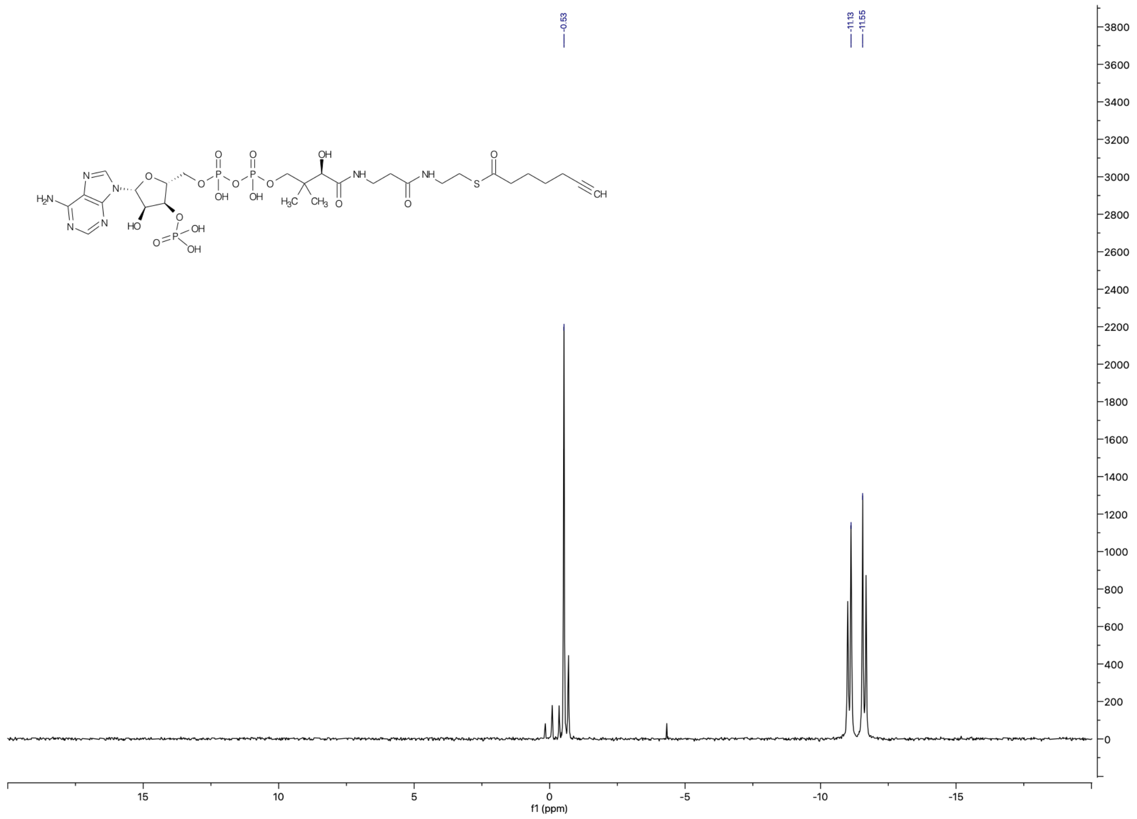

**[1]H (top), [13]C (middle) and [31]P (bottom) NMR spectra of 4AZ-CoA).**

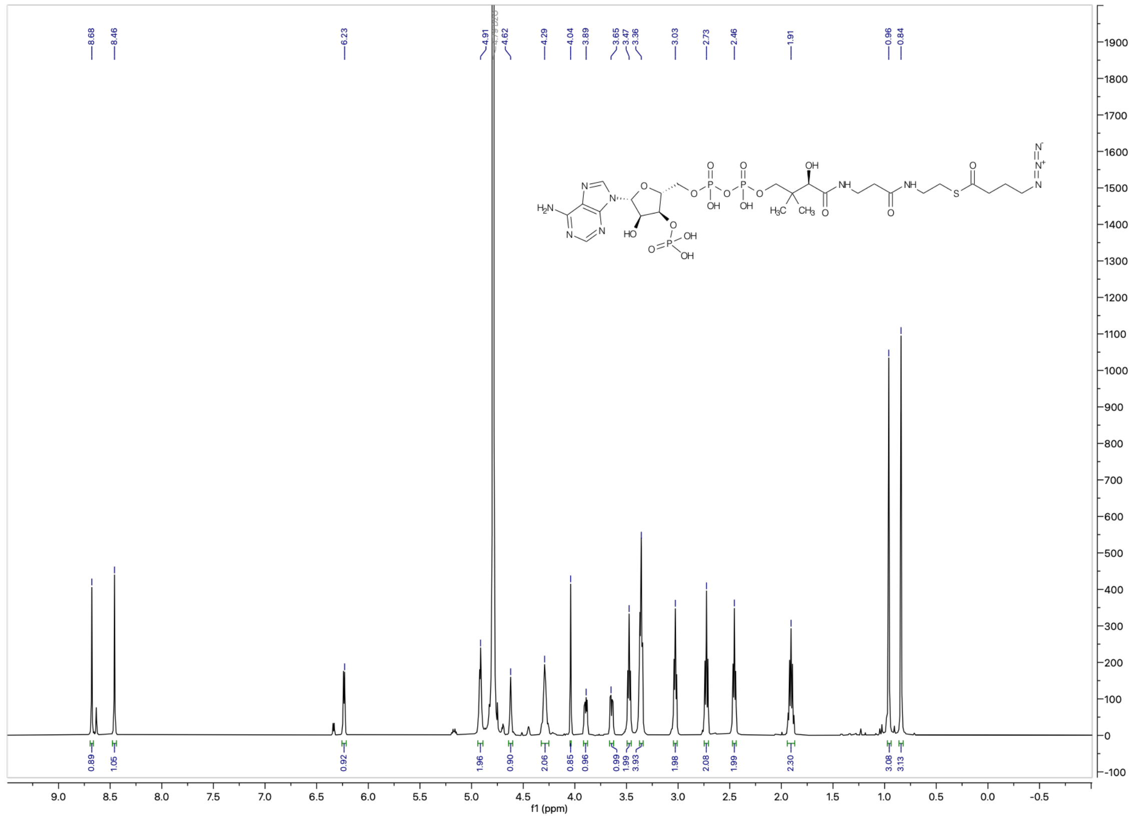

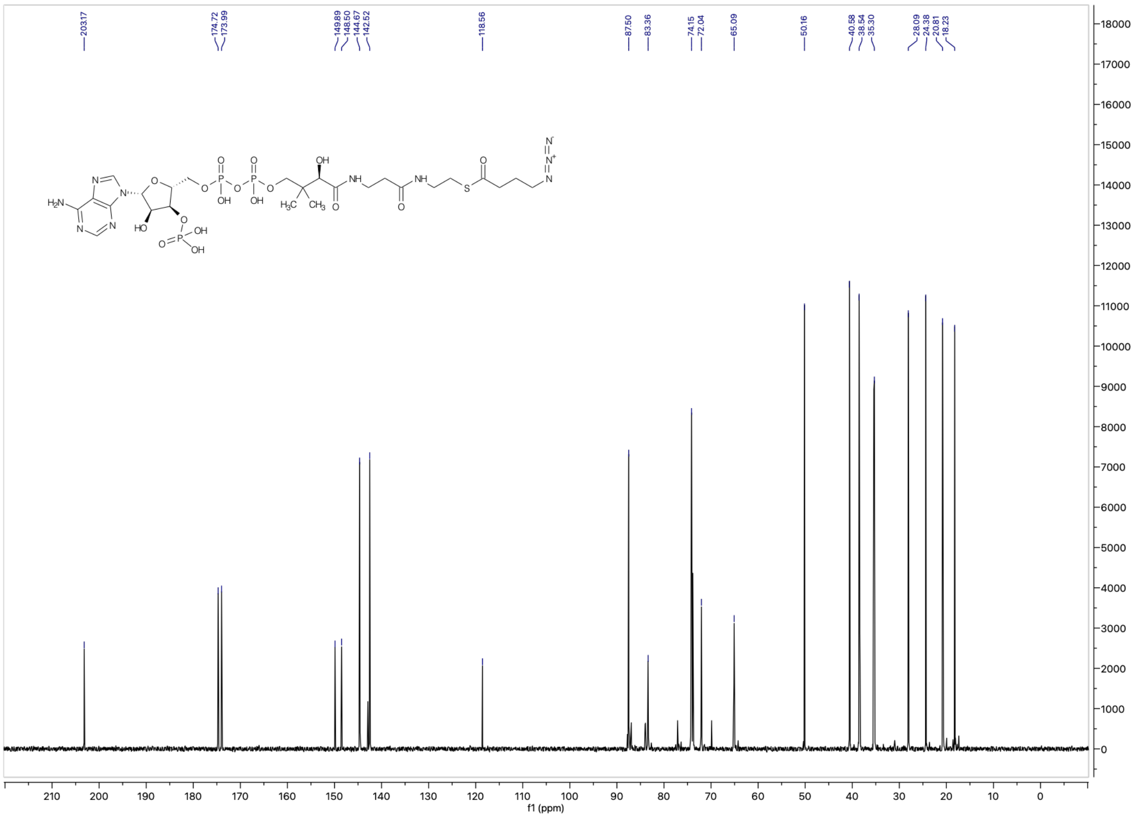
203.17
174.72
173.99
149.89
148.50
144.67
142.52
118.56
87.50
83.36
74.15
72.04
65.09
50.16
40.58
38.54
35.30
28.09
24.38
20.81
18.23
f1 (ppm)

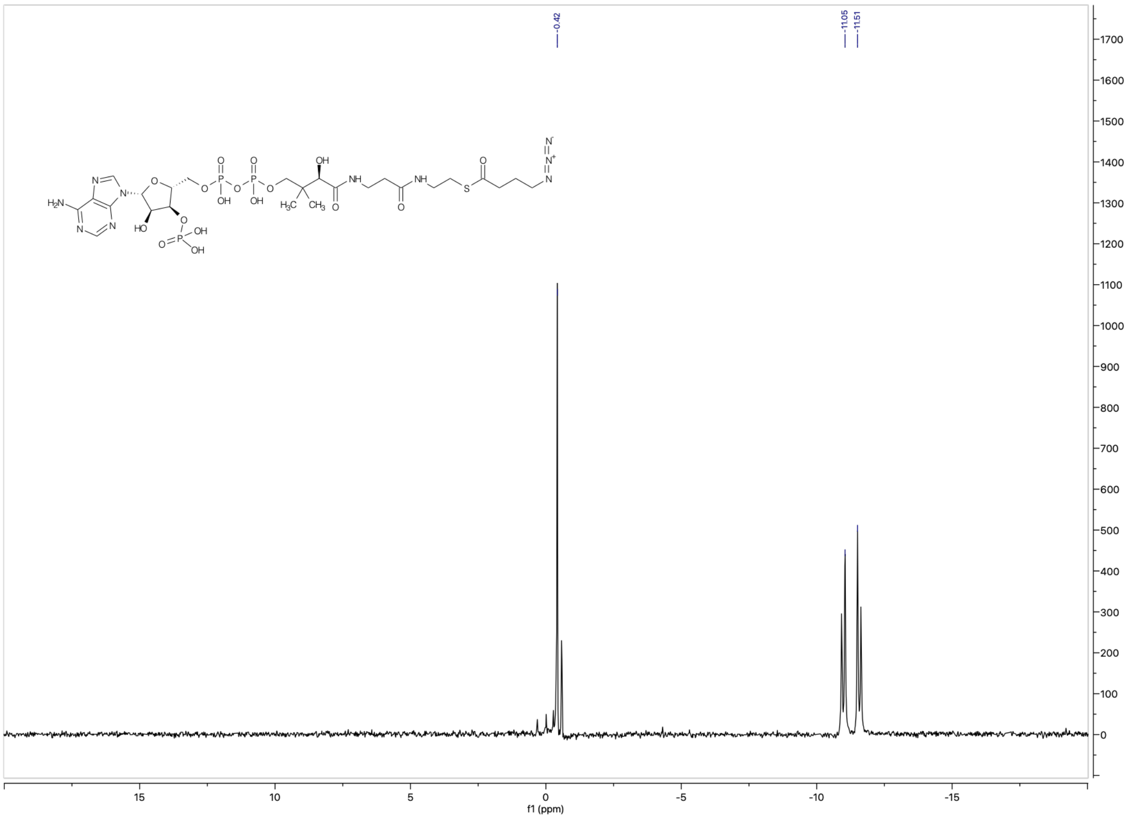
-0.42
-11.05
-11.51
f1 (ppm)

**¹H (top), ¹³C (middle) and ³¹P (bottom) NMR spectra of 3AZ-CoA.**

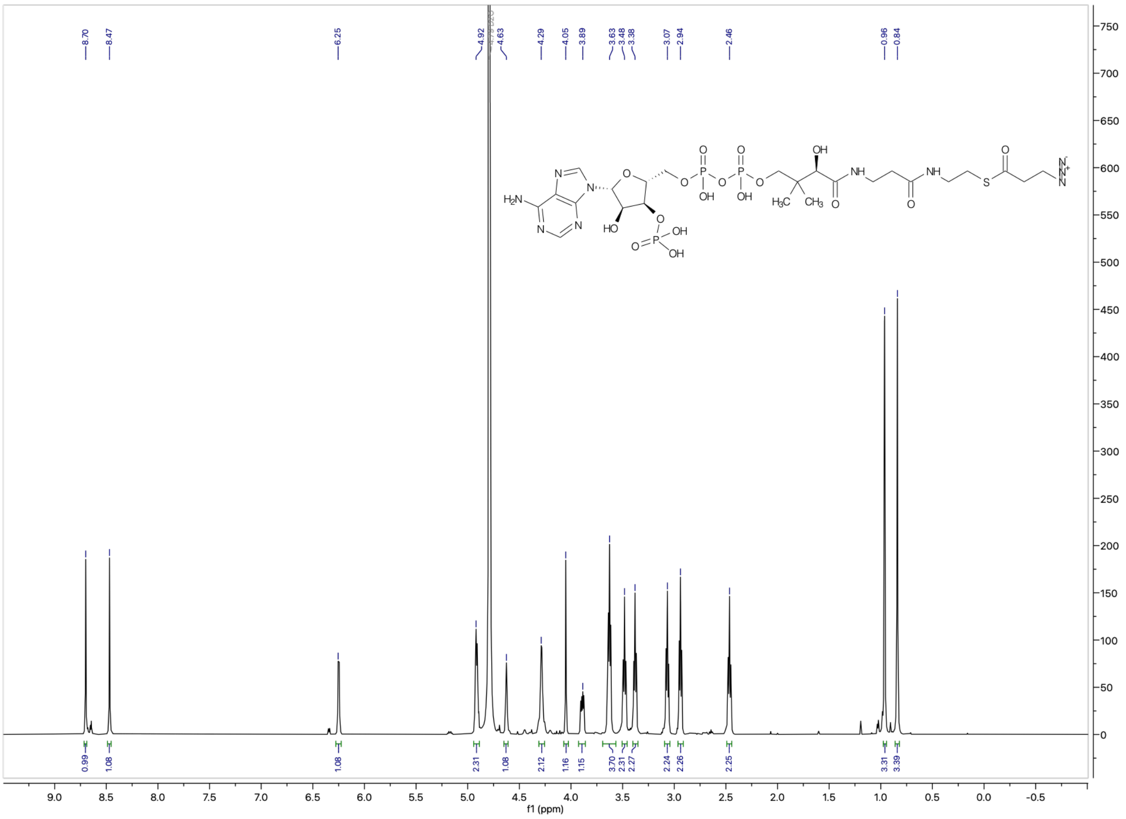

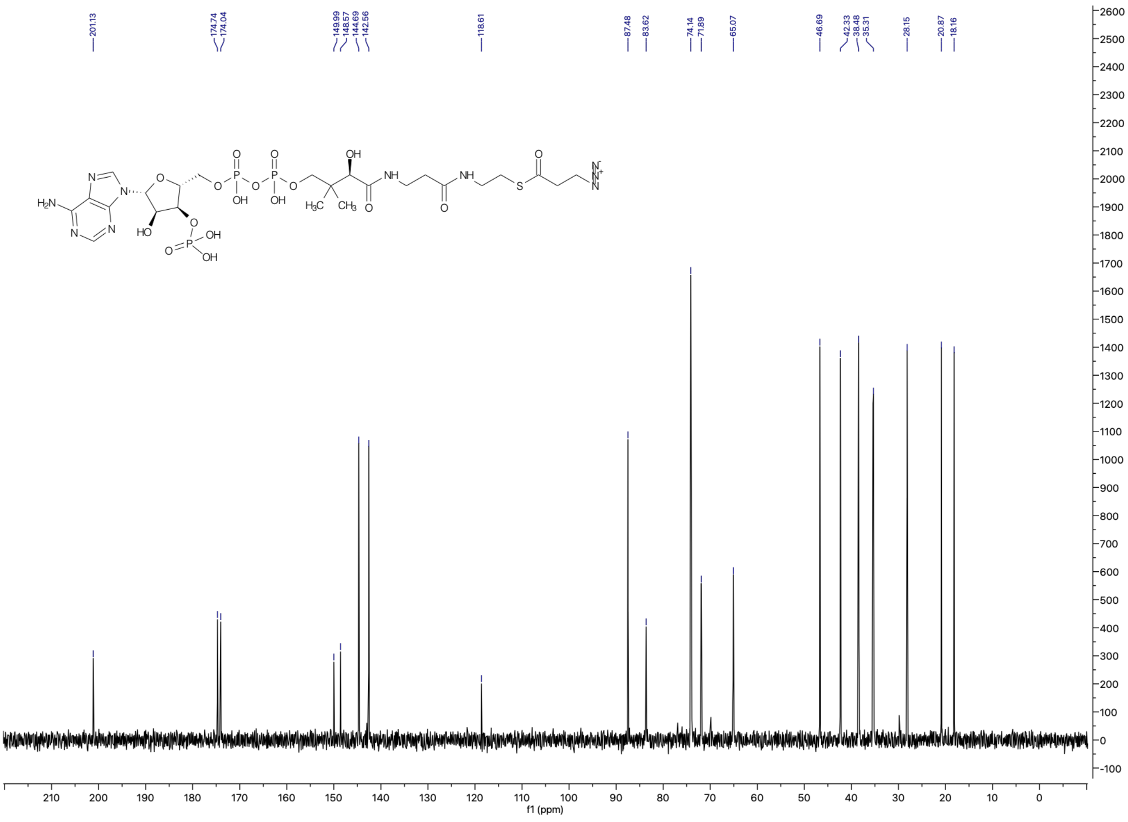

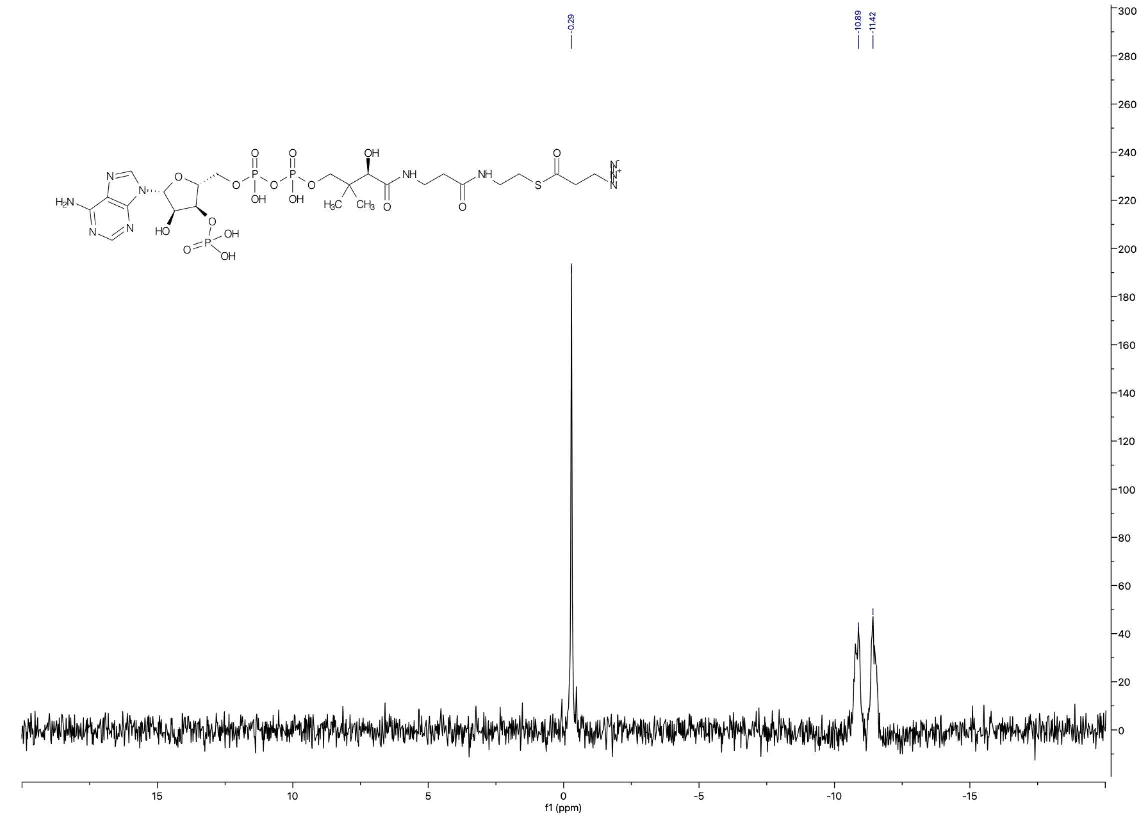

**$^{1}H$ (top), $^{13}C$ (middle) and $^{31}P$ (bottom) NMR spectra of ClAc-CoA .**

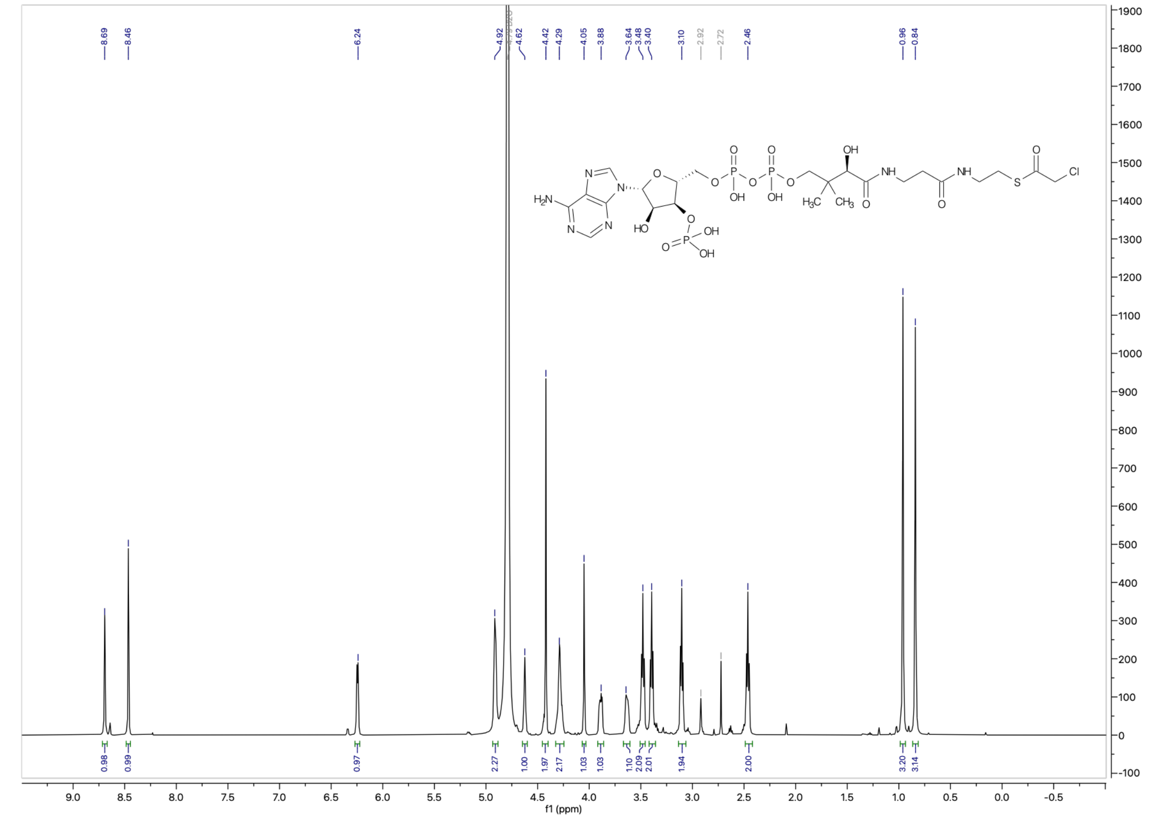

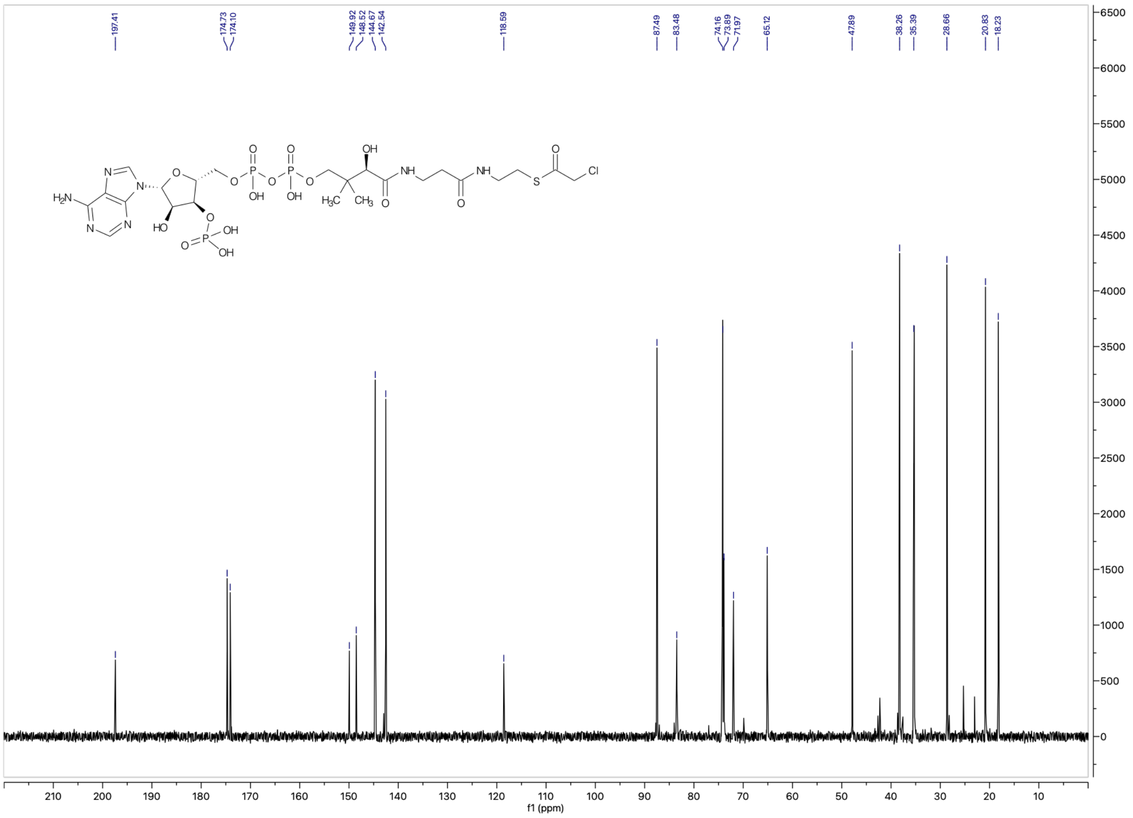
197.41
174.73
174.10
149.92
148.52
144.67
142.54
118.59
87.49
83.48
74.16
73.89
71.97
65.12
47.89
38.26
35.39
28.66
20.83
18.23
f1 (ppm)

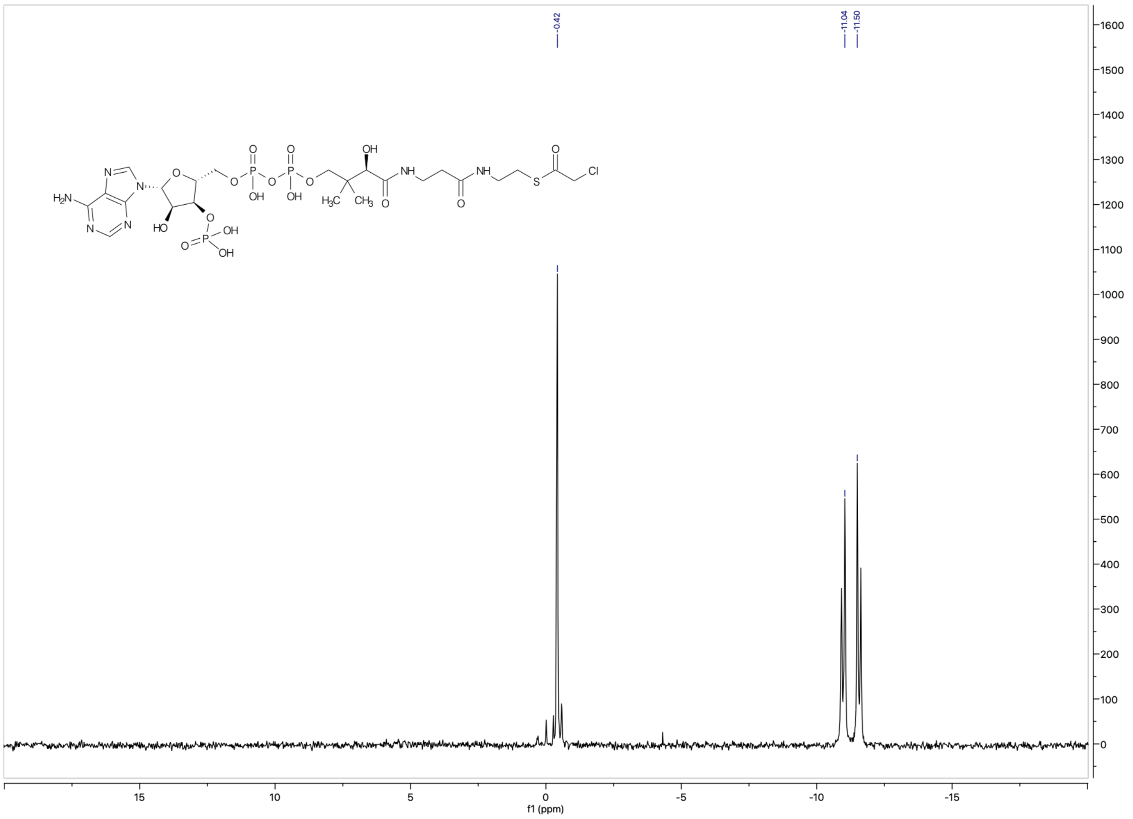
0.42
-11.04
-11.50
f1 (ppm)

### UNCROPPED GELS AND WESTERN BLOTS

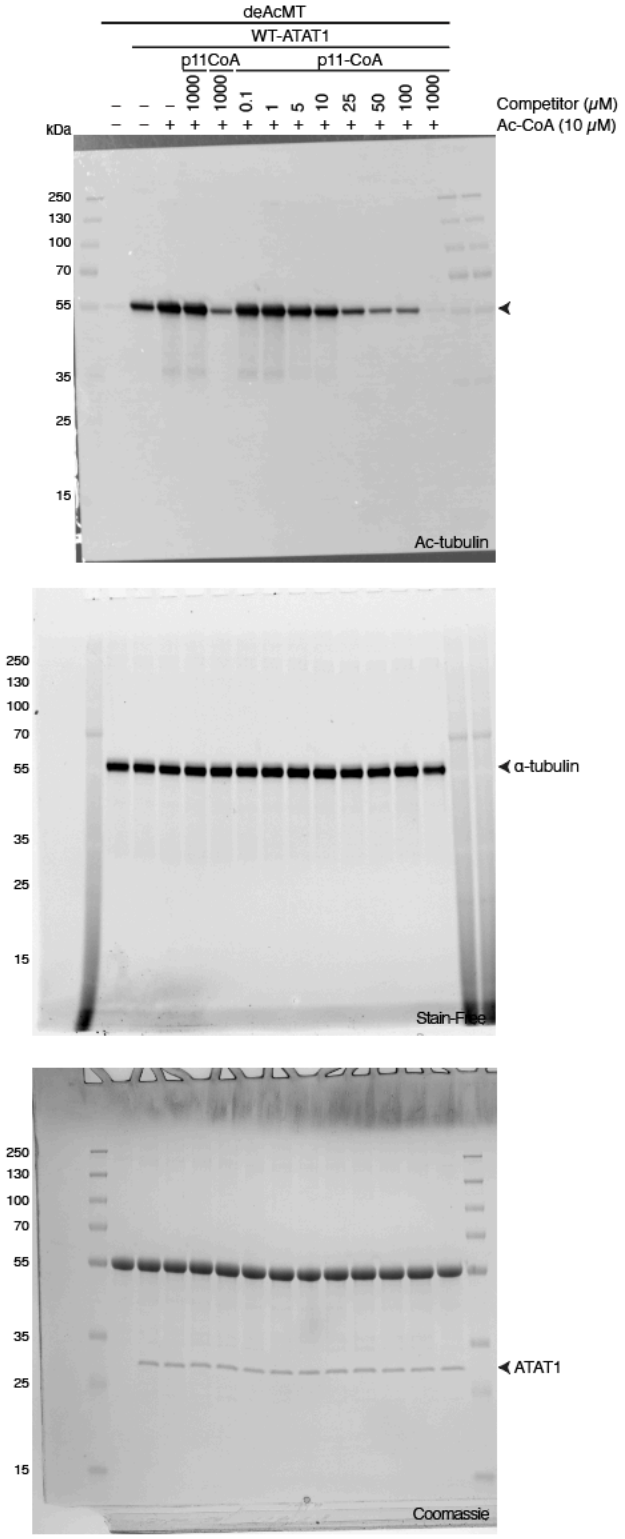

**Figure 2E.**

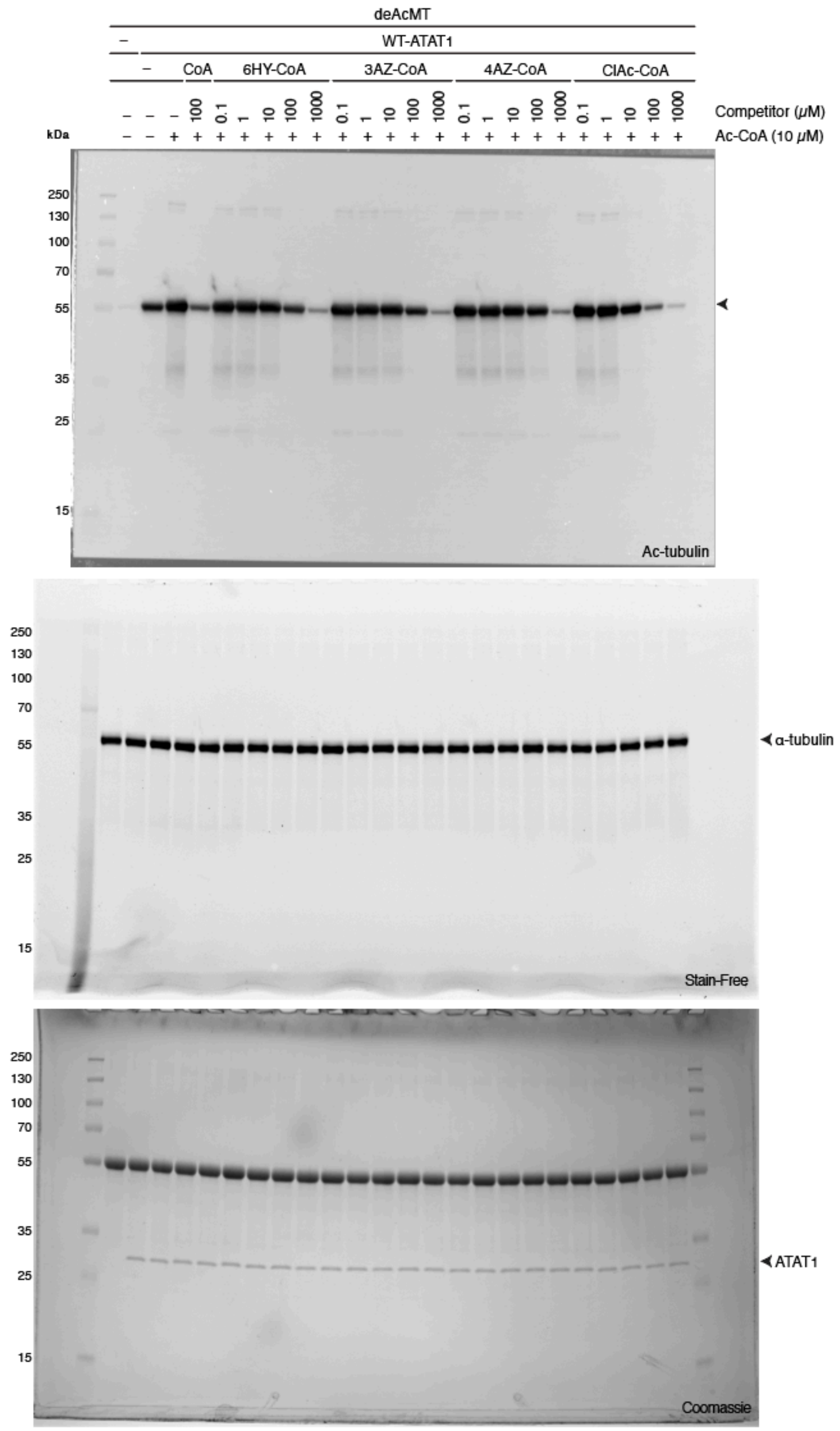
deAcMT
WT-ATAT1
CoA
6HY-CoA
3AZ-CoA
4AZ-CoA
ClAc-CoA
Competitor (μM)
Ac-CoA (10 μM)
kDa
Ac-tubulin
α-tubulin
Stain-Free
ATAT1
Coomassie

**Figure 3C.**

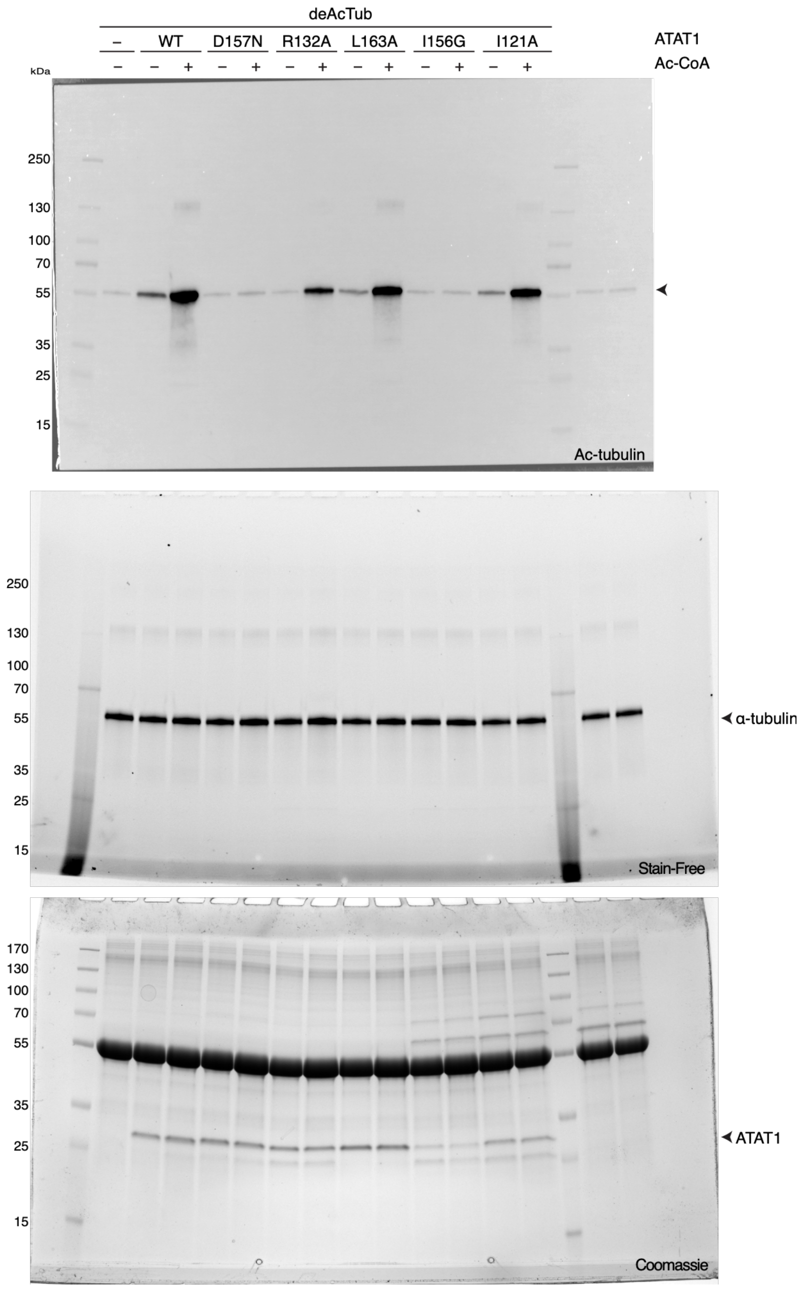
deAcTub
–
WT
D157N
R132A
L163A
I156G
I121A
ATAT1
Ac-CoA
kDa
250
130
100
70
55
35
25
15
Ac-tubulin
α-tubulin
Stain-Free
170
ATAT1
Coomassie

**Figure 4B.**

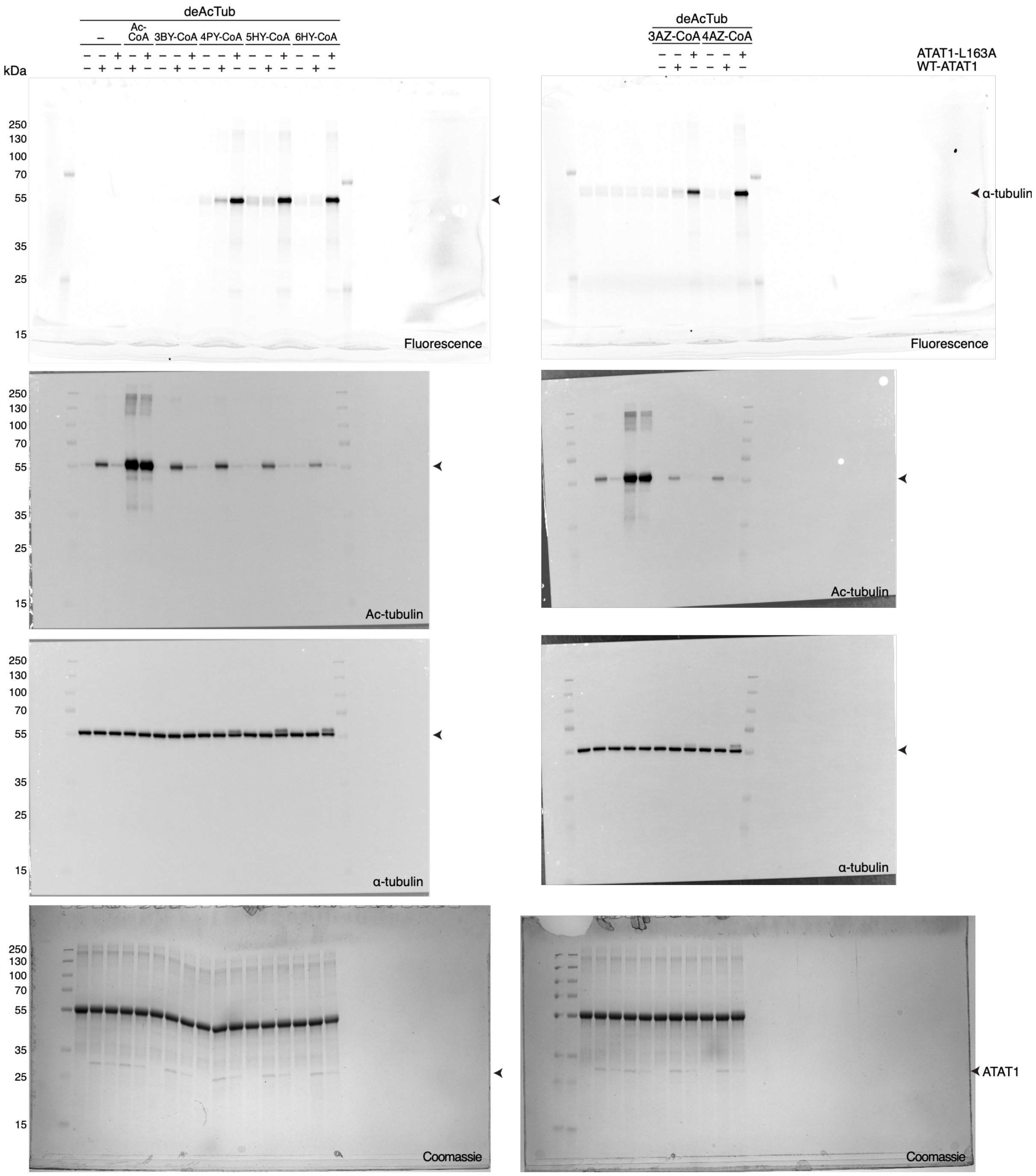
deAcTub
–
Ac-CoA
3BY-CoA
4PY-CoA
5HY-CoA
6HY-CoA
3AZ-CoA
4AZ-CoA
ATAT1-L163A
WT-ATAT1
kDa
250
130
100
70
55
35
25
15
α-tubulin
ATAT1
Fluorescence
Ac-tubulin
α-tubulin
Coomassie

**Figure 4E.**

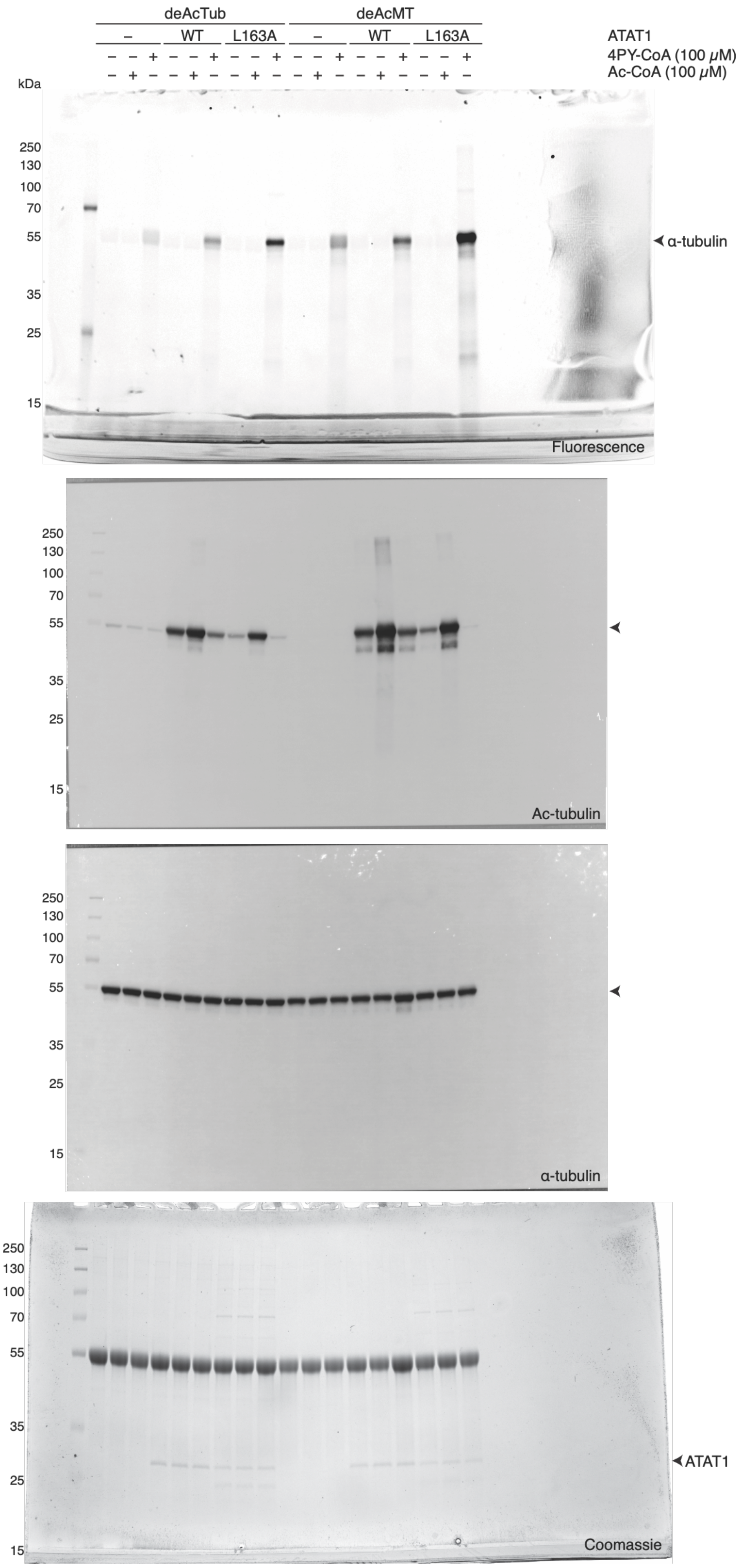
deAcTub
deAcMT
– WT L163A – WT L163A
ATAT1
– – + – – + – – + – – + – – + – – +
4PY-CoA (100 μM)
– + – – + – – + – – + – – + – – + –
Ac-CoA (100 μM)
kDa
250
130
100
70
55
35
25
15
α-tubulin
Fluorescence
Ac-tubulin
α-tubulin
Coomassie
ATAT1

**Figure 5C.**

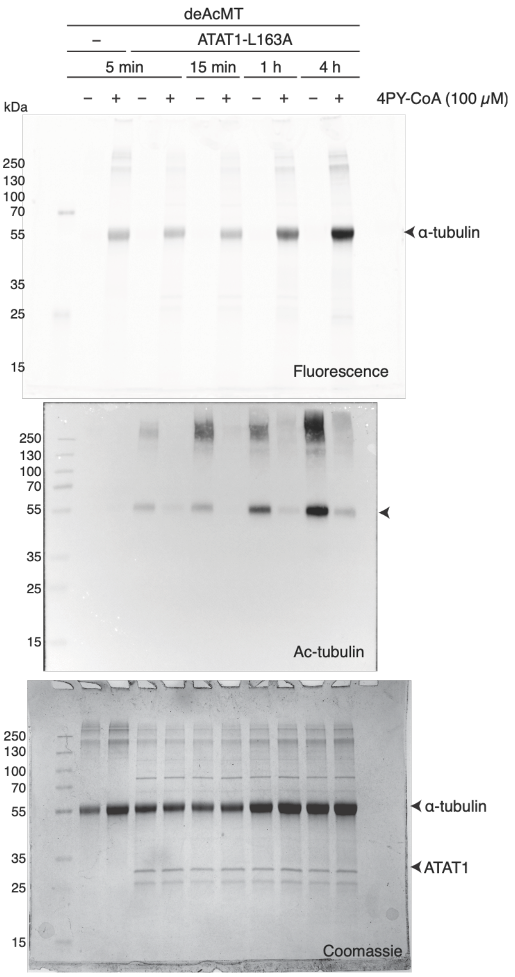
deAcMT
–
ATAT1-L163A
5 min
15 min
1 h
4 h
4PY-CoA (100 μM)
kDa
250
130
100
70
55
35
25
15
α-tubulin
Fluorescence
Ac-tubulin
Coomassie
ATAT1

**Figure 5E.**

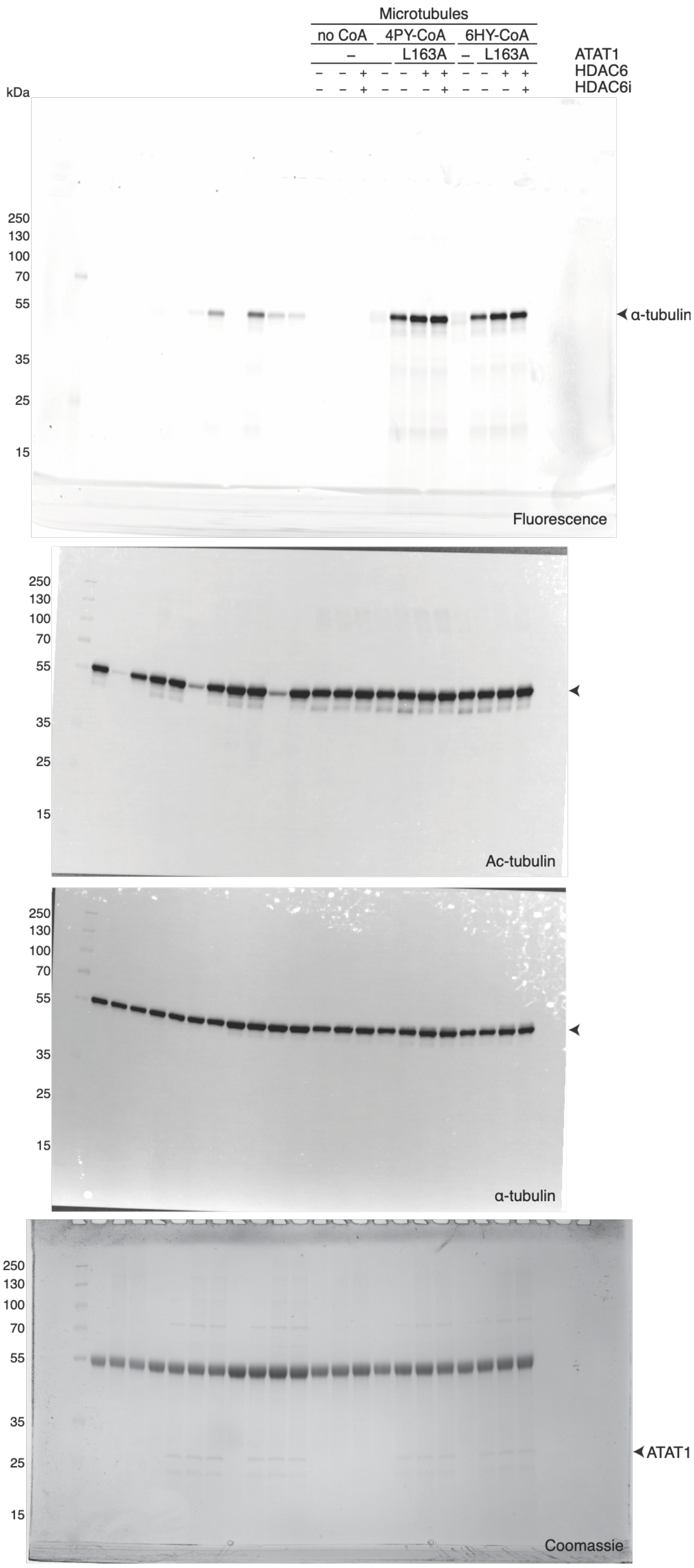

Microtubules
no CoA
4PY-CoA
6HY-CoA
–
L163A
–
L163A
ATAT1
HDAC6
HDAC6i
kDa
250
130
100
70
55
35
25
15
α-tubulin
Fluorescence
Ac-tubulin
α-tubulin
ATAT1
Coomassie

**Figure 5G.**

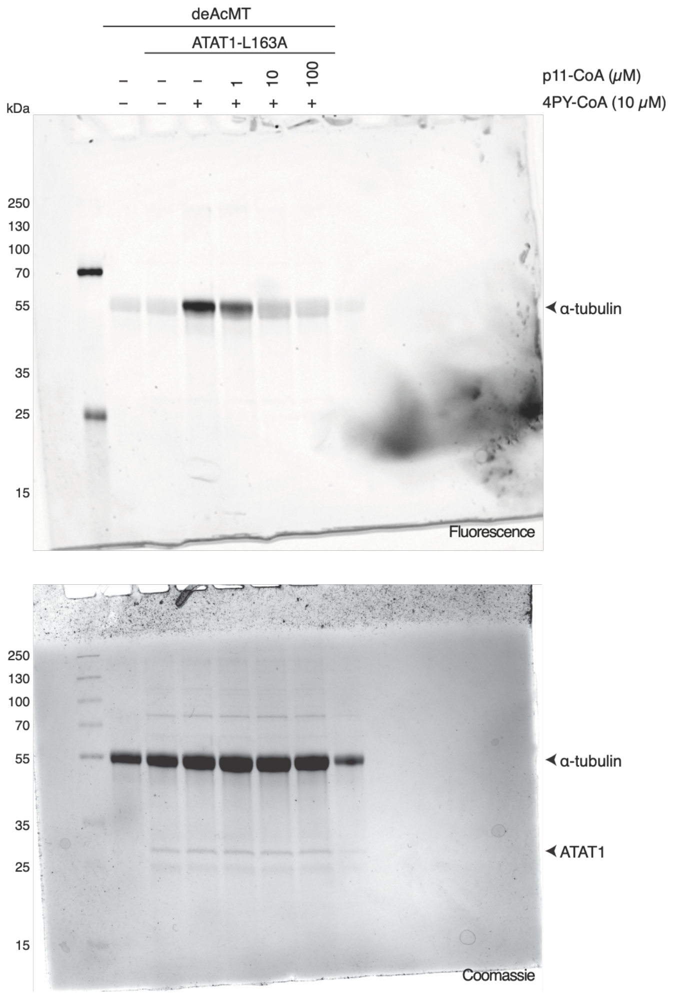
deAcMT
ATAT1-L163A
– – – 1 10 100
– – + + + +
p11-CoA (μM)
4PY-CoA (10 μM)
kDa
250
130
100
70
55
35
25
15
α-tubulin
Fluorescence
250
130
100
70
55
35
25
15
α-tubulin
ATAT1
Coomassie

**Figure 6C.**

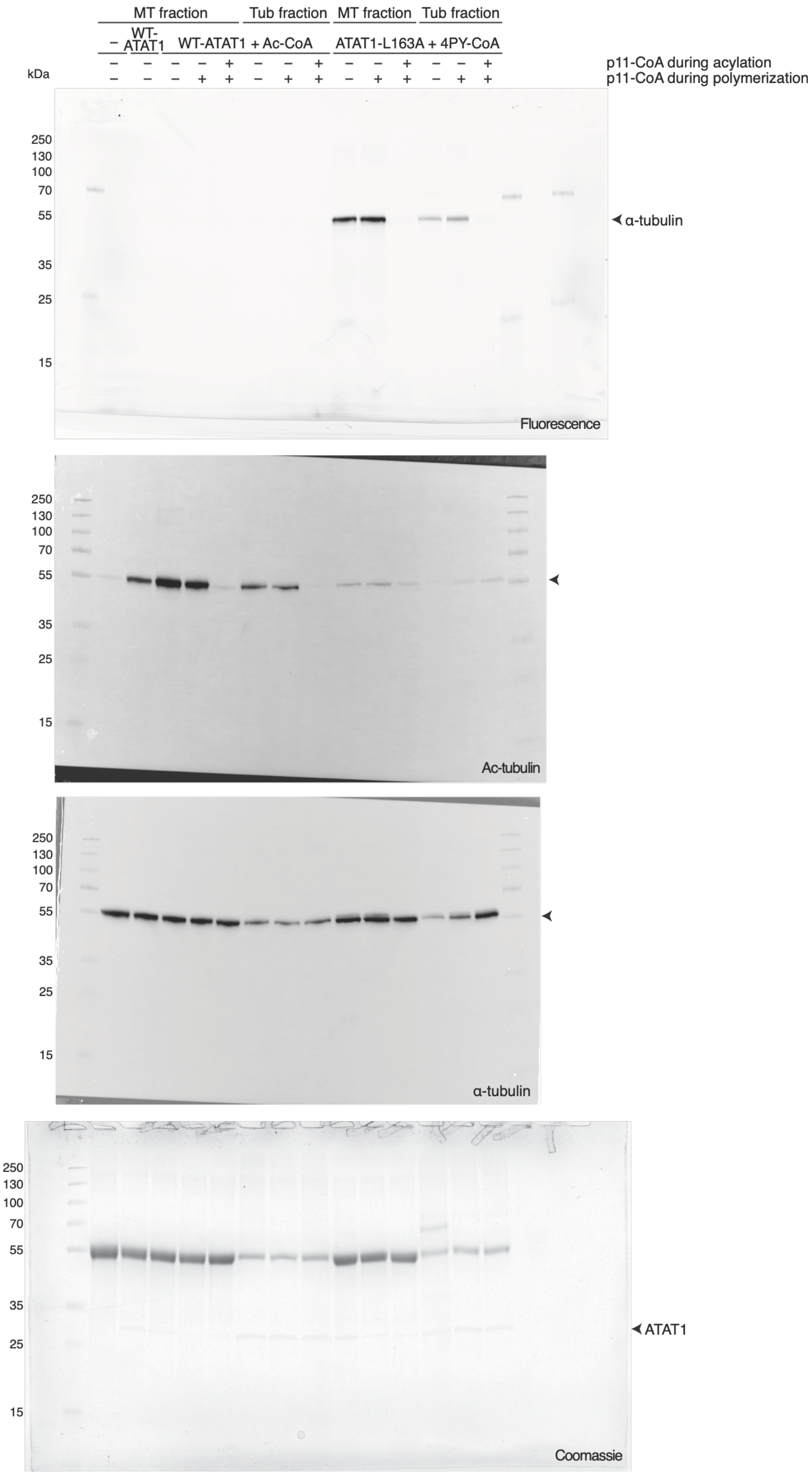
MT fraction
Tub fraction
MT fraction
Tub fraction
WT-ATAT1
WT-ATAT1 + Ac-CoA
ATAT1-L163A + 4PY-CoA
p11-CoA during acylation
p11-CoA during polymerization
kDa
250
130
100
70
55
35
25
15
α-tubulin
Fluorescence
Ac-tubulin
α-tubulin
Coomassie
ATAT1

**Figure 6F.**

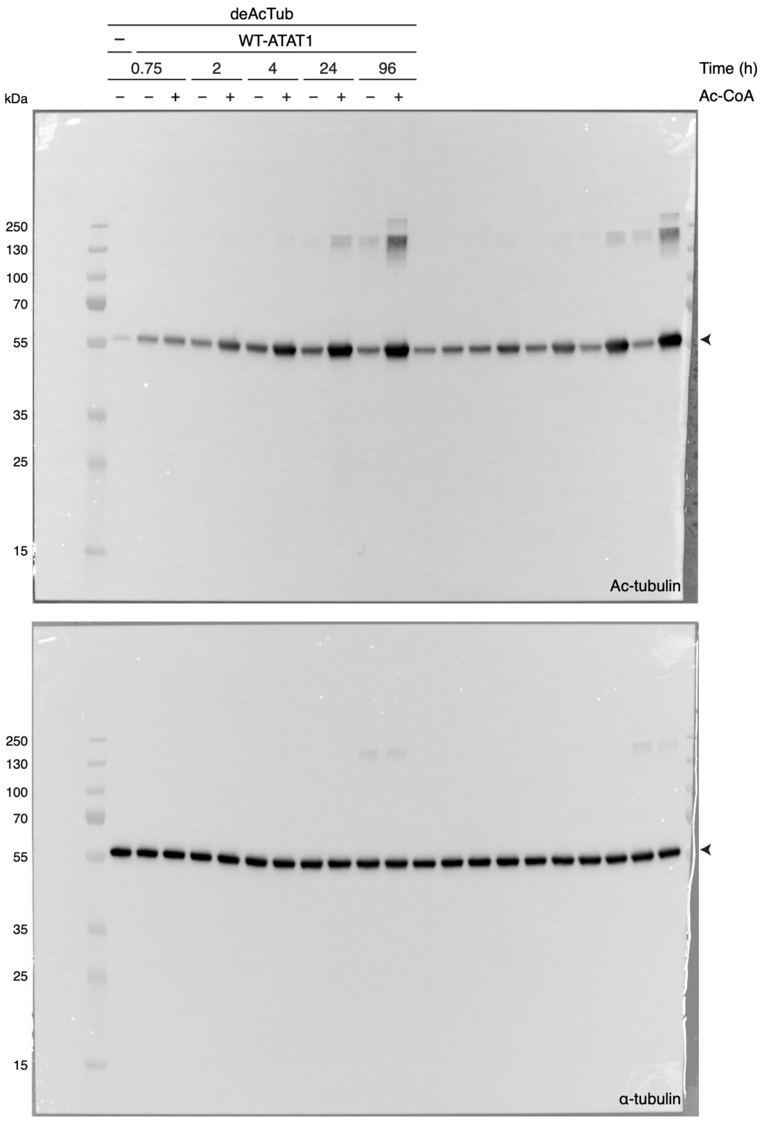
deAcTub
–
WT-ATAT1
0.75
2
4
24
96
Time (h)
kDa
– – + – + – + – + – +
Ac-CoA
250
130
100
70
55
35
25
15
Ac-tubulin
250
130
100
70
55
35
25
15
α-tubulin

**Figure S3.**

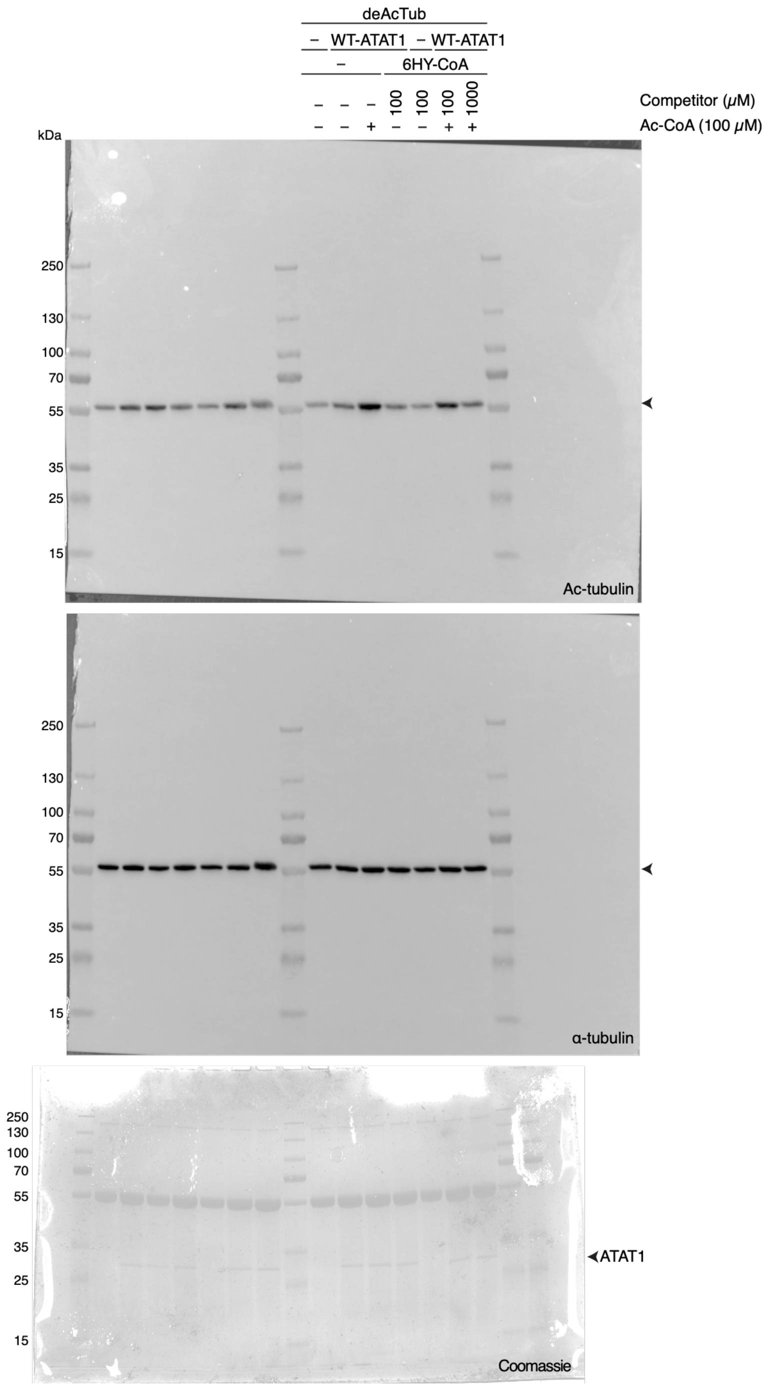

deAcTub
– WT-ATAT1 – WT-ATAT1
– 6HY-CoA
– – – 100 100 100 1000 Competitor (μM)
– – + – – + + Ac-CoA (100 μM)
kDa
250
130
100
70
55
35
25
15
Ac-tubulin
α-tubulin
ATAT1
Coomassie

**Figure S8.**

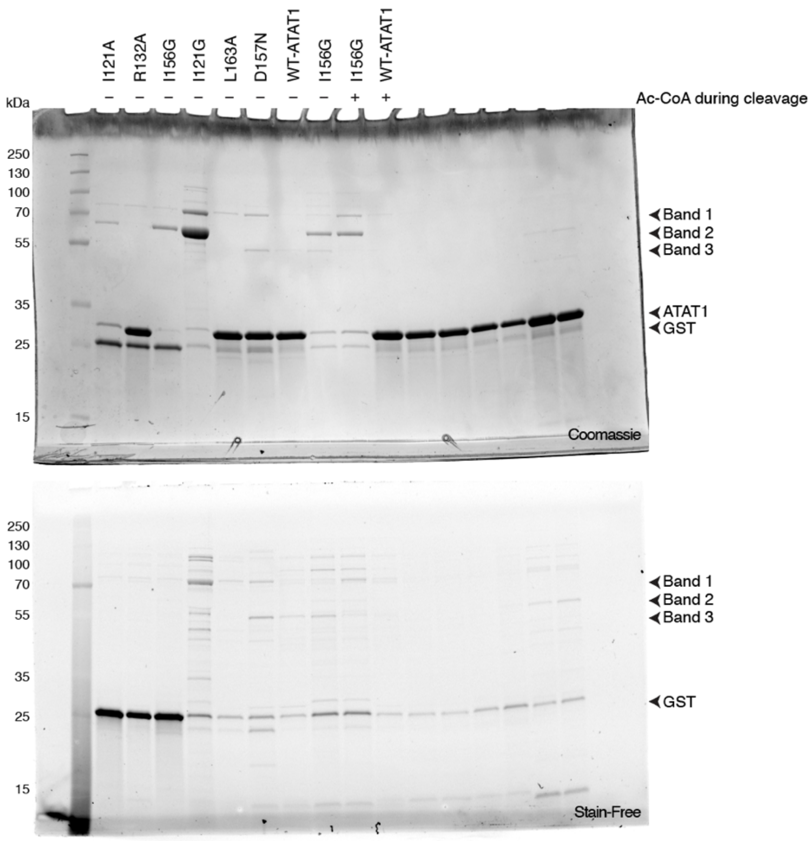
I121A
R132A
I156G
I121G
L163A
D157N
WT-ATAT1
I156G
I156G
WT-ATAT1
Ac-CoA during cleavage
kDa
250
130
100
70
55
35
25
15
Band 1
Band 2
Band 3
ATAT1
GST
Coomassie
Stain-Free

**Figure S11.**

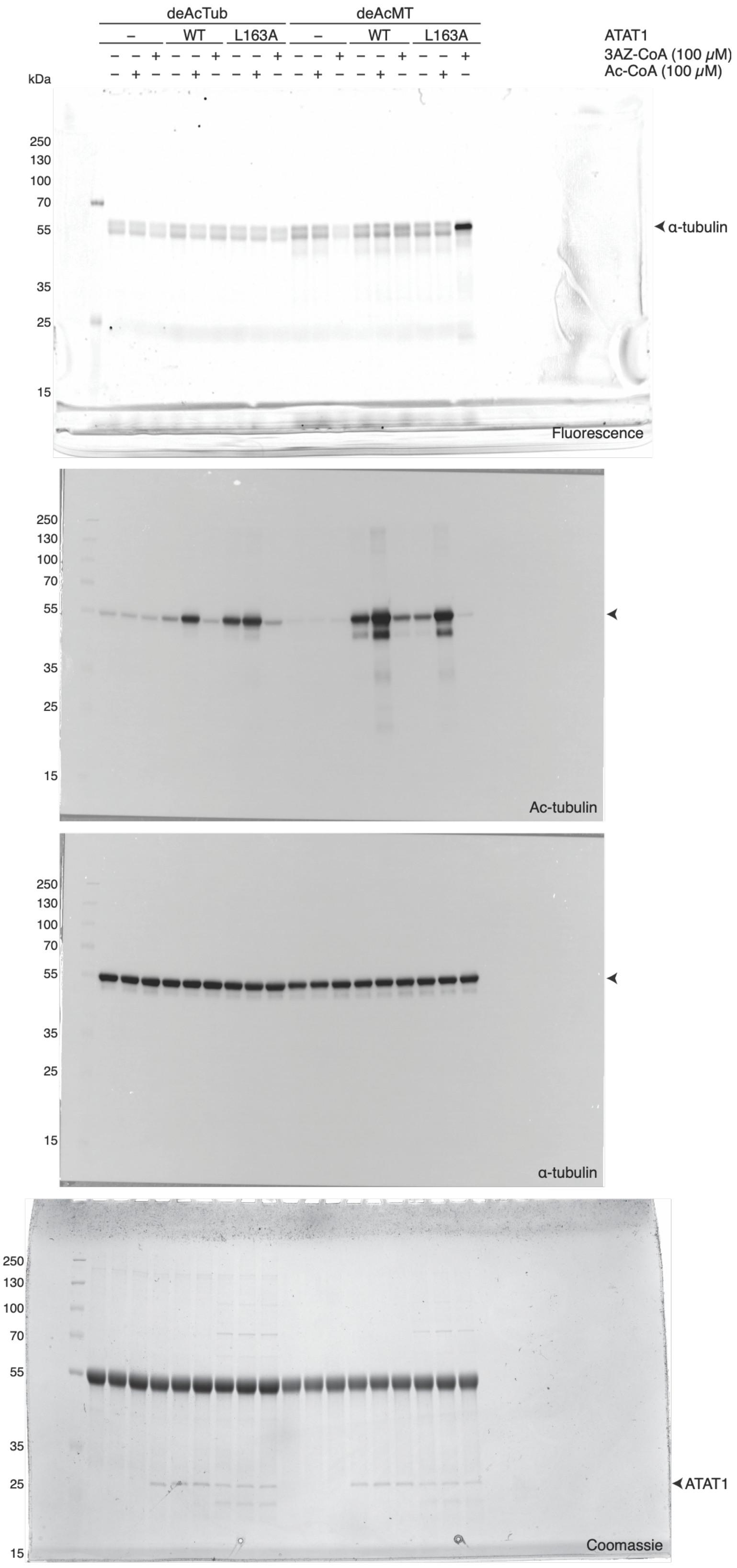

deAcTub
deAcMT
–
WT
L163A
ATAT1
3AZ-CoA (100 μM)
Ac-CoA (100 μM)
kDa
250
130
100
70
55
35
25
15
α-tubulin
Fluorescence
Ac-tubulin
Coomassie
ATAT1

**Figure S17.**

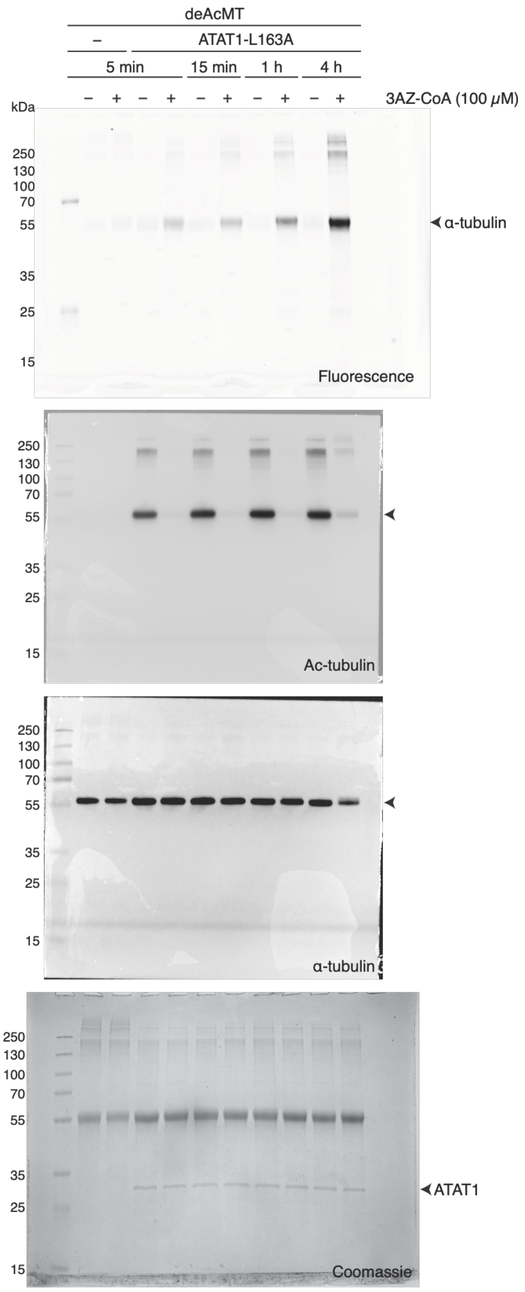
deAcMT
–
ATAT1-L163A
5 min
15 min
1 h
4 h
– + – + – + – + – +
3AZ-CoA (100 μM)
kDa
250
130
100
70
55
35
25
15
α-tubulin
Fluorescence
Ac-tubulin
α-tubulin
ATAT1
Coomassie

**Figure S18.**

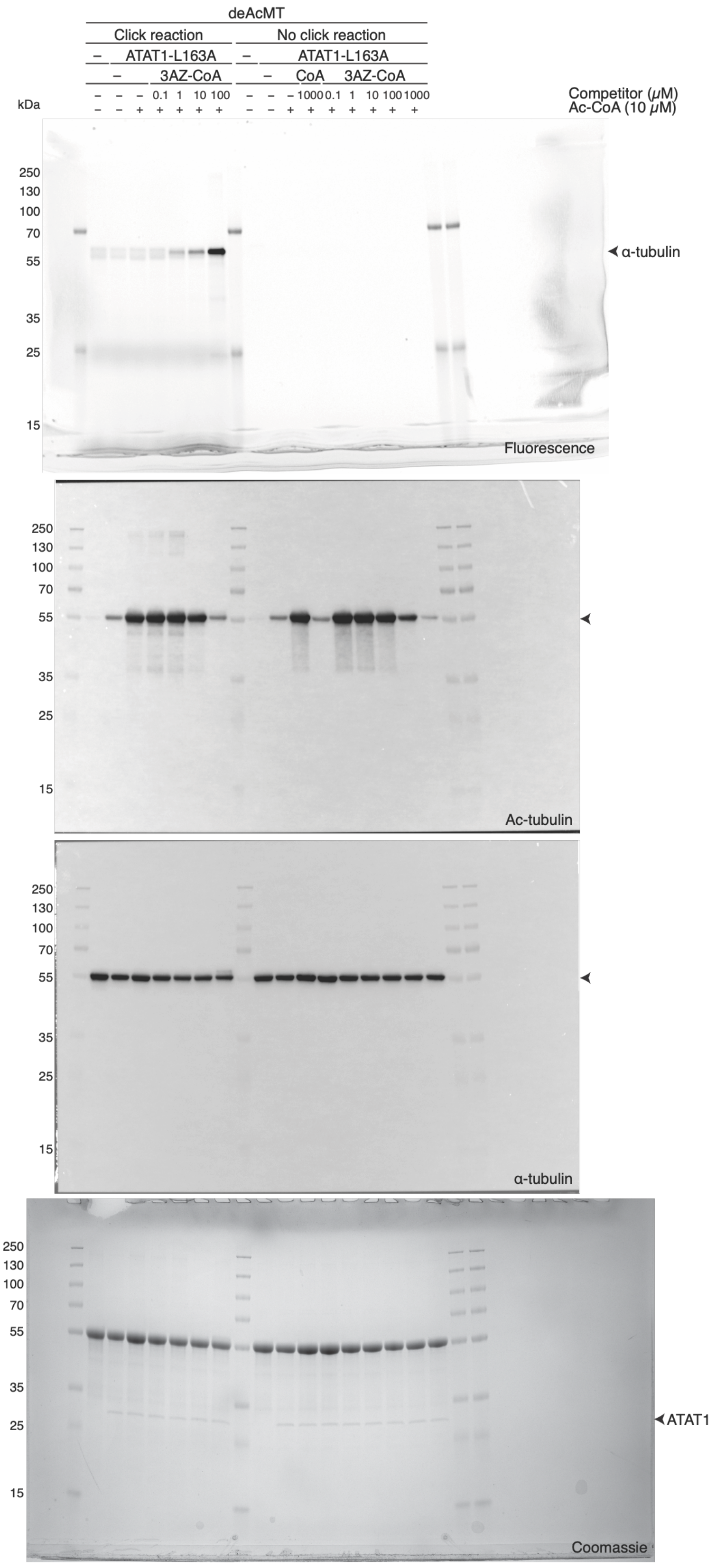
deAcMT
Click reaction
No click reaction
ATAT1-L163A
ATAT1-L163A
3AZ-CoA
CoA
3AZ-CoA
Competitor (μM)
Ac-CoA (10 μM)
kDa
α-tubulin
Fluorescence
Ac-tubulin
α-tubulin
ATAT1
Coomassie

**Figure S19A.**

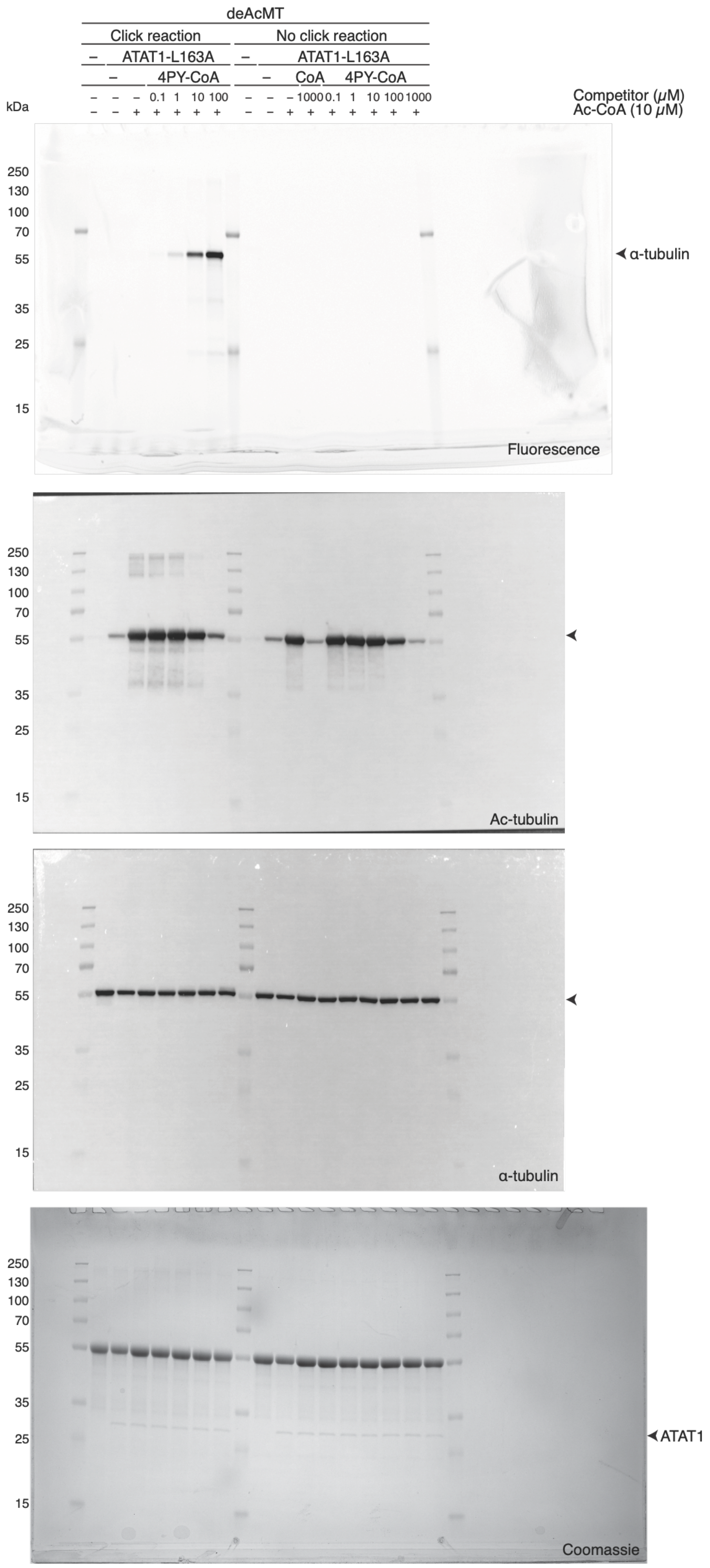

deAcMT
Click reaction
No click reaction
ATAT1-L163A
4PY-CoA
CoA
Competitor (μM)
Ac-CoA (10 μM)
kDa
250
130
100
70
55
35
25
15
α-tubulin
Fluorescence
Ac-tubulin
Coomassie
ATAT1

**Figure S19B.**

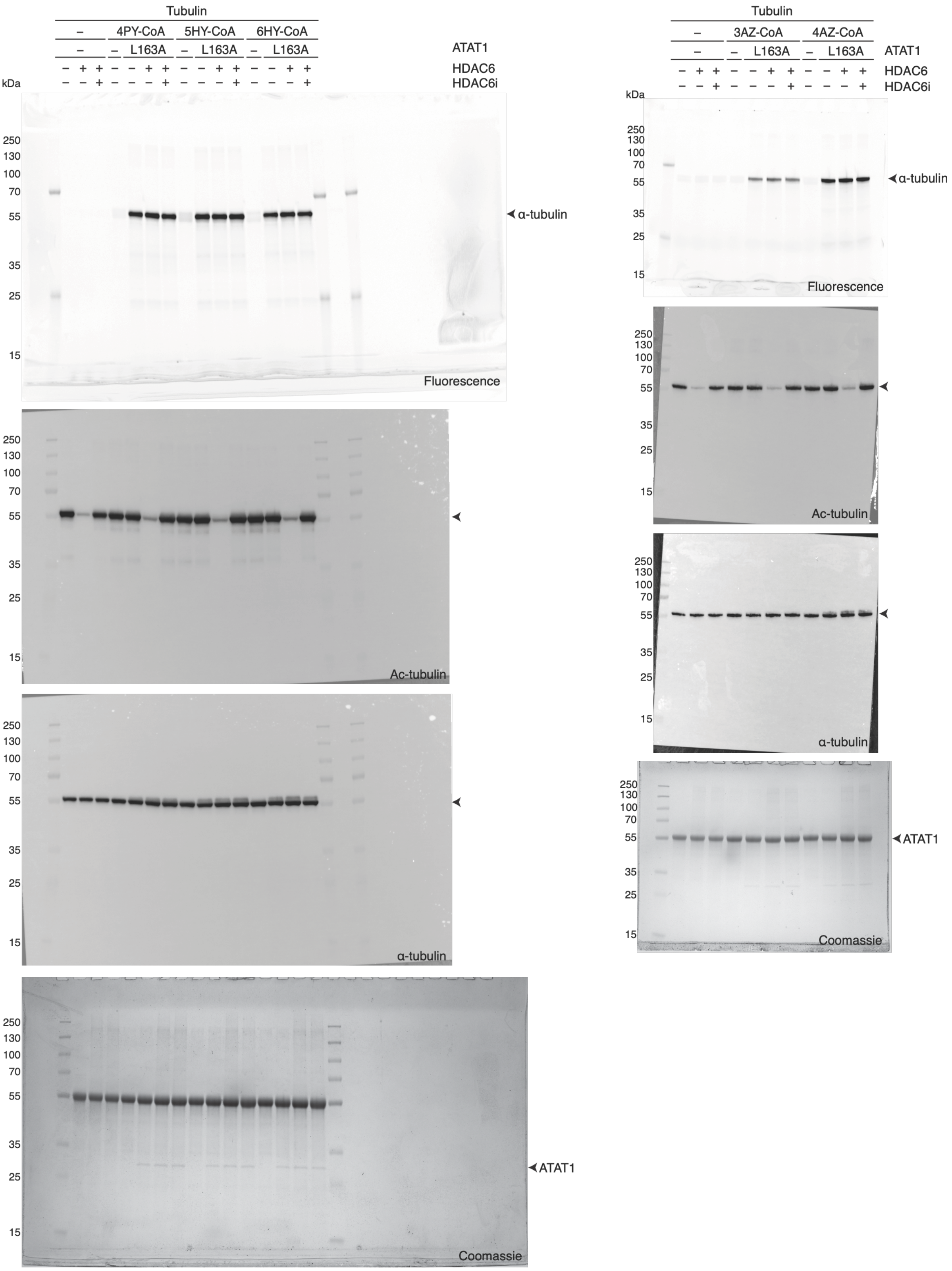
Tubulin
–
4PY-CoA
5HY-CoA
6HY-CoA
L163A
ATAT1
HDAC6
HDAC6i
kDa
250
130
100
70
55
35
25
15
α-tubulin
Fluorescence
Ac-tubulin
α-tubulin
ATAT1
Coomassie
3AZ-CoA
4AZ-CoA

**Figure S20.**

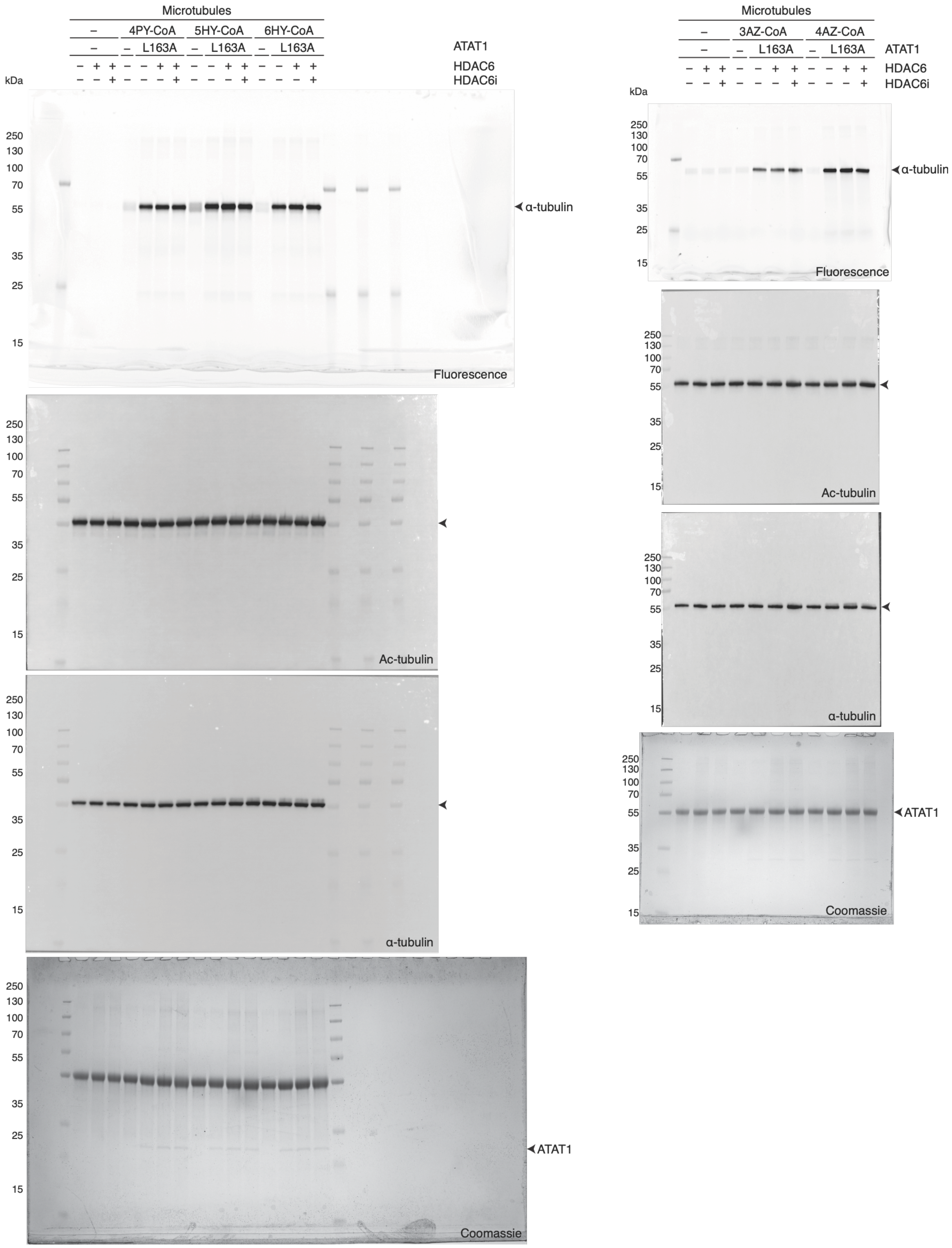
Microtubules
–
4PY-CoA
5HY-CoA
6HY-CoA
L163A
ATAT1
HDAC6
HDAC6i
kDa
α-tubulin
Fluorescence
Ac-tubulin
α-tubulin
Coomassie
ATAT1
3AZ-CoA
4AZ-CoA
250
130
100
70
55
35
25
15

**Figure S21.**
